# *Nighthawk*: acoustic monitoring of nocturnal bird migration in the Americas

**DOI:** 10.1101/2023.05.22.541336

**Authors:** Benjamin M. Van Doren, Andrew Farnsworth, Kate Stone, Dylan M. Osterhaus, Jacob Drucker, Grant Van Horn

**Affiliations:** Cornell Lab of Ornithology, Cornell University, Ithaca, New York, USA; MPG Ranch Missoula, Missoula, Montana, USA; New Mexico State University, Department of Biology, Las Cruces, New Mexico, USA; University of Chicago, Committee on Evolutionary Biology, Chicago, Illinois, USA

**Keywords:** bird migration, bioacoustics, acoustic monitoring, machine learning, machine listening, movement ecology, artificial intelligence

## Abstract

1. Animal migration is one of nature’s most spectacular phenomena, but migratory animals and their journeys are imperiled across the globe. Migratory birds are among the most well-studied animals on Earth, yet relatively little is known about in-flight behavior during nocturnal migration. Because many migrating bird species vocalize during flight, passive acoustic monitoring shows great promise for facilitating widespread monitoring of bird migration.
2. Here, we present Nighthawk, a deep learning model designed to detect and identify the vocalizations of nocturnally migrating birds. We trained Nighthawk on the in-flight vocalizations of migratory birds using a diverse dataset of recordings from across the Americas.
3. Our results demonstrate that Nighthawk performs well as a nocturnal flight call detector and classifier for dozens of avian taxa, both at the species level and for broader taxonomic groups (e.g., orders and families). The model accurately quantified nightly nocturnal migration intensity and species phenology and performed well on data from across North America. Incorporating modest amounts of additional annotated audio (50-120 h) into model training yielded high performance on target datasets from both North and South America.
4. By monitoring the vocalizations of actively migrating birds, Nighthawk provides a detailed window onto nocturnal bird migration that is not presently attainable by other means (e.g., radar or citizen science). Scientists, managers, and practitioners could use acoustic monitoring with Nighthawk for a number of applications, including: monitoring migration passage at wind farms; studying airspace usage during migratory flights; monitoring the changing migrations of species susceptible to climate change; and revealing previously unknown migration routes and behaviors. Overall, this work will empower diverse stakeholders to efficiently monitor migrating birds across the Western Hemisphere and collect data in aid of science and conservation. Nighthawk is freely available at https://github.com/bmvandoren/Nighthawk.

## Introduction

Animal migration is important to science and society. Migrating animals have captured human imagination for millennia, and movement is a key mechanism by which organisms can adjust to rapid environmental change (Van Doren et al. 2021). Movement is a fundamental mediator of organisms’ interactions with their environment and with each other. As anthropogenic pressures increasingly influence these interactions, knowledge of movement is increasingly necessary for guiding conservation action (Davy et al. 2017, Fraser et al. 2018). This is particularly true for animals that use the atmosphere—a key habitat where volant organisms interact, forage, and even rest (Diehl 2013, Liechti et al. 2013). For species constantly on the move, their life histories and conservation threats are inextricably tied to ever-changing spatiotemporal distributions. Despite the need for movement data to inform science and conservation, migratory animals are often challenging to monitor. They may be rare, secretive, sensitive to disturbance, too small to carry a tracking device, or too expensive to monitor at sufficient numbers or resolution (Kays et al. 2015).

Migratory birds, for instance, are among the world’s most well-studied moving organisms, but scientists still lack important information about the movements of most species. There is great urgency to develop better methods for monitoring migratory birds, as populations of these global travelers have declined precipitously over the last half-century (Rosenberg et al. 2019). Available tools for monitoring migratory birds include Doppler radar networks (Gauthreaux et al. 2003, Dokter et al. 2011, Bauer et al. 2019) and widespread citizen science projects such as eBird (Sullivan et al. 2014). These tools are invaluable: radar can measure the fluxes of moving birds even at continental extents (Van Doren and Horton 2018, Nussbaumer et al. 2021), and data from citizen scientists can support detailed spatial models used by scientists and practitioners globally (Reynolds et al. 2017, Fink et al. 2020). However, these tools also carry major shortcomings. Doppler weather radars cannot discern individuals or species identities, and they are generally stationary, expensive, and easily limited by mountainous topography. Scientific use of radar data is also frequently impeded by bureaucratic, political, or technical issues (Shamoun-Baranes et al. 2022), and radar is still a developing tool outside of North America and the Western Palearctic. Citizen science databases such as eBird (Sullivan et al. 2014) can provide information at finer taxonomic scales, but survey coverage can be highly variable away from human population centers. In addition, most bird species migrate at night, and so crowdsourced human observations have limited ability to actively monitor nocturnal avian movements. Fortunately, radar, citizen science, and other data sources are highly complementary, and a growing number of studies integrate these and other multimodal data to infer unseen behavior (Shipley et al. 2018, Bota et al. 2020, Van Doren et al. 2023). However, no current approaches provide a scalable and affordable path towards monitoring the in-flight behavior of billions of nocturnally migrating birds at species and individual resolution.

Acoustic methods are used frequently by bat researchers to monitor species presence and activity and increasingly by ornithologists to detect breeding species (Sugai et al. 2019), but their potential to study animals on the move is often overlooked. Many bird species actively vocalize during nocturnal flights, and ground-based microphones can capture the “flight calls” of migrating individuals (Farnsworth 2005). Because flight calls often encode species identity, acoustic monitoring provides an accessible path towards portable, inexpensive, and widely deployable systems for species- and individual-level monitoring (Evans and Rosenberg 2000). Recent work has demonstrated the potential of acoustic monitoring to document the nightly passage rates of migratory birds (Van Doren et al. 2023), and acoustic information could enable the study of intra- and interspecies interactions during migration (Gayk and Mennill 2023). The primary impediment to the widespread adoption of acoustic monitoring of bird migration is the large time investment needed to transform hours of recorded audio into counts of identified vocalizations. In recent years, however, the outlook has greatly improved due to advances in machine learning technology for sound detection and classification (Lostanlen et al. 2018, Kahl et al. 2021, Van Doren et al. 2023). However, important challenges remain that prevent broad application of acoustic systems for avian migration monitoring. Existing systems are rarely trained on large numbers of nocturnal flight calls and are frequently based on datasets that are restricted in geographic scope, and therefore these systems may not generalize well to other areas.

*Merlin* is a smartphone application that uses machine learning to identify bird vocalizations (https://merlin.allaboutbirds.org/). The sound identification module of Merlin is powered by a convolutional neural network that applies computer vision principles to audio spectrograms. Merlin Sound ID is a high-performing system with great promise for a range of acoustic monitoring applications, but the current system is not ideal for migration monitoring applications for several reasons. Merlin outputs species detections on 3-second chunks of audio, but flight calls are much shorter than this duration (often lasting only 50-250 ms) (Evans and O’Brien 2002, Lanzone et al. 2009). Temporal resolution is therefore limited to 3 seconds; the use of smaller chunk sizes could increase temporal resolution. Furthermore, many flight calls cannot be easily identified to species, and Merlin does not capture this taxonomic uncertainty (but see Cramer et al. 2020). However, because neural networks are highly configurable, adjustments to model architecture and training data could address these challenges.

Here, we present *Nighthawk*, a deep learning model based on Merlin Sound ID that is designed to detect and identify the vocalizations of nocturnally migrating birds. We trained Nighthawk on in-flight vocalizations from a diverse collection of recordings collected across the Americas. In this research paper, we examine the performance of the Nighthawk model across spatial and taxonomic scales. We evaluate model performance across a taxonomic hierarchy of 82 species, 18 families, and 4 orders of birds that vocalize during nocturnal migration. We apply Nighthawk to an entire season of data and investigate how well the model generalizes across spatial scales, including acoustic monitoring data from South America. Finally, we discuss best practices for using Nighthawk to monitor bird migration in the Americas.

## Methods

### Overview

Our analysis comprised three parts (Figure 1). In Part 1, we assembled a dataset of annotated recordings and trained Nighthawk to predict the presence of flight calls of different avian taxa in an audio segment. We refer to this dataset as the “Core dataset”. In Part 2, we trained additional versions of Nighthawk and applied them to data from locations not included in training (hereafter, “target datasets”). We tested target recordings from regions well-represented in the Core dataset (“in-domain” data) and regions poorly-represented (“out-of-domain” data). In Part 3, we applied Nighthawk to continuous recordings collected across an entire migration season—the intended use case of Nighthawk.

**Figure 1:**
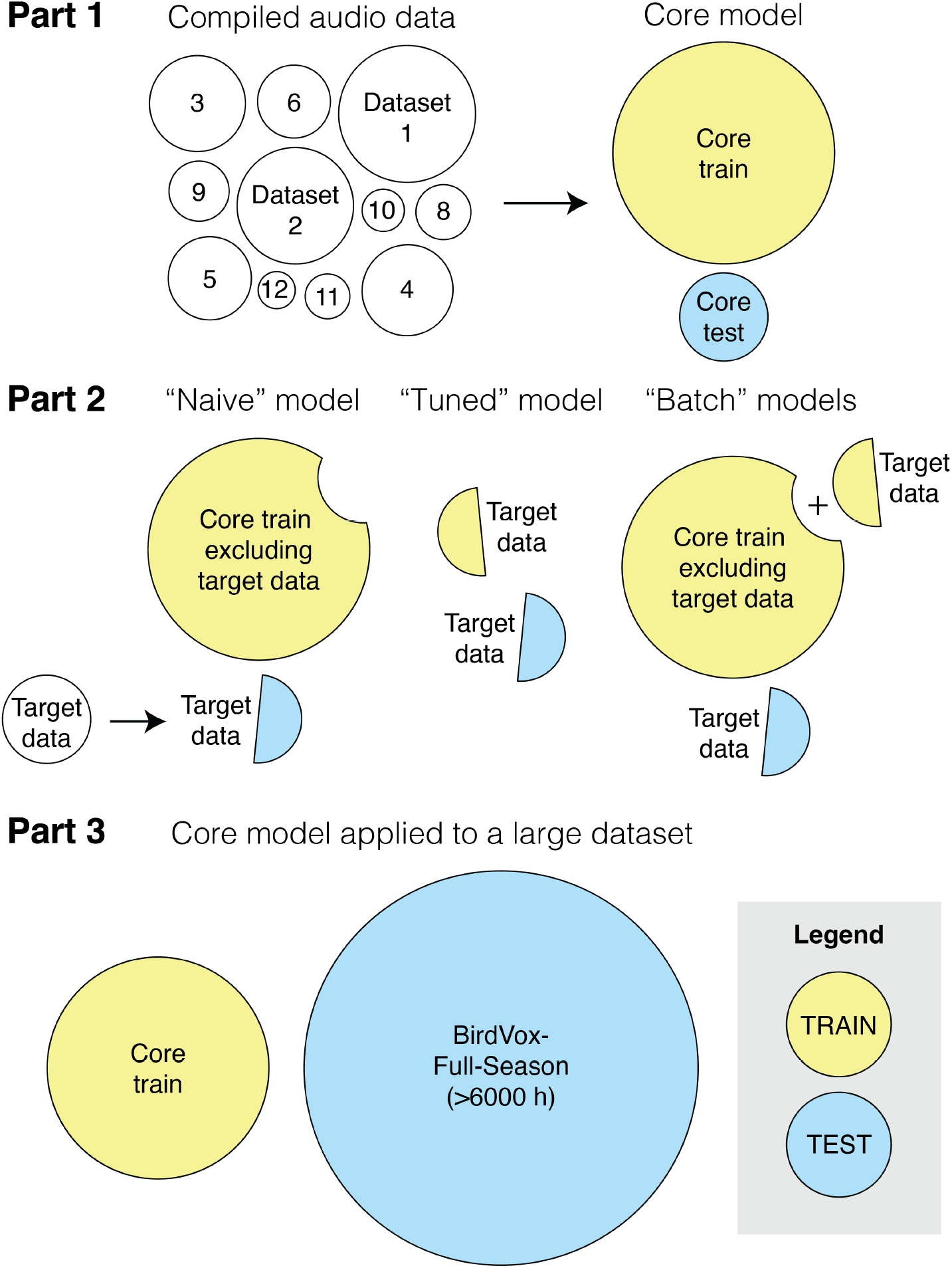
Analysis overview. In Part 1, we split annotated acoustic data into nonoverlapping “Core” test and train sets and trained the Core model. In Part 2, we applied Nighthawk to three target datasets (from Pennsylvania, New Mexico, and Colombia) using three different model construction strategies (Naive, Tuned, and Batch). In Part 3, we applied the Core model to an entire season of continuous audio from a migration monitoring array in central New York State, USA.

### Acoustic data

We obtained annotated recordings from 25 data sources, comprising published and unpublished datasets from across the Americas. Geographically, most data came from eastern North America, with some from western North America and a small sample from northern South America. A summary of data sources is in Table S1. These data primarily comprised passive nocturnal audio recordings (∼72%); the remainder were general sound datasets (∼18%), recordings of captive birds (∼2%), or not easily placed in one category (∼8%). These datasets were generated using varied approaches, and we anticipated variation in their error rates. We checked representative samples of each dataset to verify error rates were less than 5%, and we corrected errors where encountered.

### Data annotation

To assign recorded vocalizations to specific avian taxa, we visualized audio as spectrograms and annotated the onset and offset of each vocalization event. When a bird made repeated vocalizations, we considered sounds occurring more than 0.5 s apart as separate events. We used multiple software platforms that support temporal annotation, including Audacity, Raven Pro, and a custom web interface (https://merlinvision.macaulaylibrary.org).

Identification uncertainty is a fundamental part of acoustic monitoring; many flight calls can be classified to species, but others cannot (e.g., due to recording quality or overlap with similar-sounding species) (Evans and O’Brien 2002, Landsborough et al. 2019). We addressed this challenge by explicitly modeling taxonomic uncertainty. Annotators labeled flight calls at the finest taxonomic resolution possible. We used species, families, and orders as defined in the Clements Checklist, the avian taxonomy used by the eBird database (Sullivan et al. 2014). We recorded species-level classifications using the abbreviated codes defined in this taxonomy (e.g., “amered” is the code for American Redstart *Setophaga ruticilla*). For species that have broadly similar flight calls, we created an additional “group” classification level. We defined a group as two or more similar-sounding species in the same family. Each species can belong to at most one group. For example, Swamp Sparrow (*Melospiza georgiana*) and Lincoln’s Sparrow (*M. lincolnii*) give highly similar flight calls, and we treat them together here as the “SWLI” group. Thus, we used a four-level taxonomic hierarchy: species, group, family, and order. Table S2 lists all defined groups and their member species.

### Taxa included

We focused on the following taxa: (1) nocturnally migrating landbirds in the orders Passeriformes and Cu-culiformes, (2) nocturnally migrating shorebirds in the order Charadriiformes, and (3) nocturnally migrating waterbirds in the order Pelecaniformes. We included taxa known to migrate at night and vocalize during flight (Evans and O’Brien 2002). For example, we did not include the Passeriformes families of Fringillidae and Hirundinidae because these taxa primarily migrate diurnally. We included taxa for which we were able to compile at least 50 training examples and 20 testing examples, totaling 82 species, 17 groups, 18 families, and 4 orders.

### Part 1: Assembling the Core dataset

We focused annotation efforts on building a balanced and accurate test dataset. For each taxonomic class, we sampled calls from a diversity of datasets while limiting total dataset size to keep manual review tractable. We used three steps: first, we constructed a Core test split that minimized spatiotemporal overlap with the Core train split; second, we subsampled test data to obtain a taxonomically balanced and representative sample; and third, we reviewed all annotations for accuracy.

To ensure separation between Core test and train splits, we randomly assigned audio to train or test splits based on recording location wherever possible, such that each recording site was included in *either* train or test sets, but not both, thus preventing the model from using idiosyncratic information about recording location or microphone setup. We were unable to subdivide 33% of the data solely by location, either because data came from a single recording location (e.g., captive recordings from a single research station) or because location was not consistently reported. For these remaining data, we randomly assigned each audio file to either the train or test split based on recording session, such that no data from the same session was included in both train and test splits.

Next, we subsampled test data to obtain a Core test set tractable for manual review. For each taxonomic class, we randomly selected an annotated call from each of our source datasets in sequence until we either reached 500 calls or ran out of annotations for that class. We then randomly sampled an additional 10,000 negative examples (e.g., ambient noise or vocalizations that were not flight calls) following the same procedure. Our dataset includes many audio files containing more than one vocalization, and we annotated all flight call vocalizations of non-target taxa, so in practice our test set includes more than 500 examples from frequently-recorded taxa. See Table S3.

Finally, two authors (BMVD and AF) reviewed all Core test set annotations to correct any errors. If the two reviewers disagreed about the identity of a vocalization, we used a more general taxonomic categorization for that vocalization. For example, if there was agreement that a call was made by a thrush (family Turdidae), but not on its species identity, we used “Turdidae” as the classification. After review, the Core train and test sets consisted of 428009 and 47305 annotations, respectively (Table S3).

For Core model training and evaluation, we extracted all audio segments containing flight calls, as well as an approximately equal number of audio segments containing no flight calls. We refer to this approach as a “Balanced Cropped Vocalization” analysis.

### Part 1: Building the Core Nighthawk model

To build the Core *Nighthawk* model, we adapted the existing *Merlin Sound ID* deep learning framework. Merlin is trained to identify bird vocalizations but is not optimized for the short-duration calls prevalent in recordings of nocturnal bird migration. Merlin uses a deep neural network to predict the bird species present in short audio segments. Merlin takes as input a 3-second audio clip and outputs a vector that represents learned features relevant to the classification task at hand. These features are then used as inputs to a logistic regression model to predict the presence of each species. The neural network backbone of Merlin is a MobileNet (Howard et al. 2019), a compact network designed to perform well on mobile devices.

We modified the Merlin Sound ID architecture in three important ways (Figure 2). First, we decreased the input duration from 3 seconds to 1 second because flight calls are shorter vocalizations (generally 50-250 ms) than those typically evaluated by Merlin (Evans and O’Brien 2002, Farnsworth 2005). Second, because we were not constrained to smartphone deployments, we replaced the MobileNet backbone with a ResNet-34 backbone (He et al. 2016). ResNet-34 is a larger architecture that can achieve higher performance than MobileNet but requires more computational resources. Third, to facilitate classification at our four taxonomic levels (species, group, family, order), we replaced Merlin’s single, species-level output layer with four output layers, each of which received a shared feature vector as input (Figure 2). With this multi-head output, Nighthawk can simultaneously make predictions of the species, group, family, and order classes of every input.

**Figure 2:**
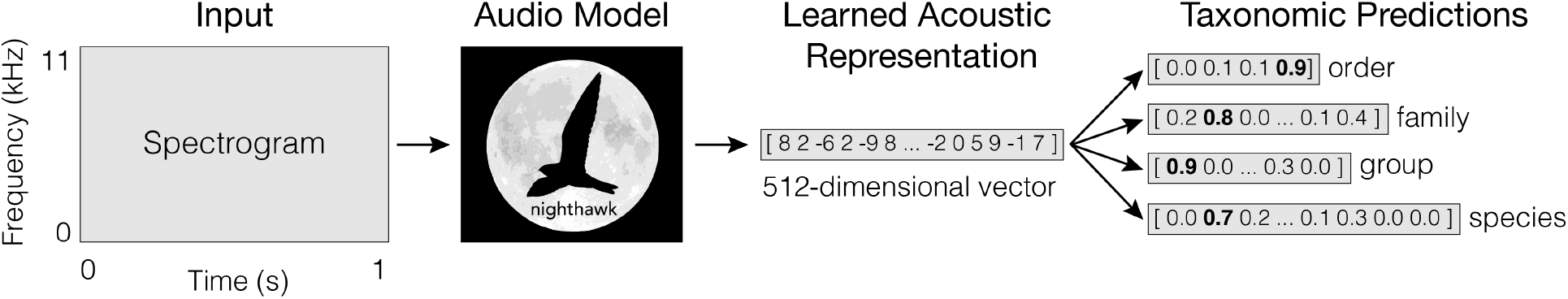
Nighthawk takes as input a 1-s spectrogram and outputs a high-dimensional feature vector that is a learned representation of the input. The final layers of the model apply logistic regression to this feature vector to make predictions across multiple taxonomic levels.

We trained Nighthawk on the Core training set of 428009 annotations. Hyperparameter sweeps revealed consistent performance after training for 100 epochs (batch size = 64) using stochastic gradient descent optimization with cosine learning rate decay, starting at a learning rate of 0.1.

### Part 1: Evaluating the Core model

We evaluated Core model performance on the Core test dataset of Balanced Cropped Vocalizations with *average precision* (AP) scores. Average precision is a metric favored by the machine learning community because AP captures overall performance without setting an arbitrary score threshold. AP summarizes the precision-recall curve for a single class and takes a value between 0 and 1, where 1 represents a system with perfect precision and perfect recall for that class. The *mean average precision* (mAP) is the mean of AP scores across multiple classes and summarizes overall model performance. Here, we evaluate models by calculating mAP scores separately for species, group, family, and order levels.

### Part 2.1: Constructing target datasets

After evaluating the Core model (Part 1), we investigated Nighthawk’s performance on data from locations not included in its training dataset (“target datasets”). We constructed and evaluated models with three target datasets, respectively from (1) Pennsylvania, USA, (2) New Mexico, USA, and (3) Colombia. In each case, we split target datasets into train and test halves by recording session.

### Part 2.2: Evaluating target models

We evaluated all target models on the holdout (test) portion of target data. We conducted two types of evaluations. In the first, we constructed Balanced Cropped Vocalization datasets for model training, in which we extracted all target audio segments containing flight calls and roughly 2x as many audio segments without flight calls. We evaluated these data using average precision (AP), as for the Core evaluation.

In field applications, we expect to deal with very unbalanced datasets, in which there are many more sections of audio without flight calls than those with flight calls. Therefore, in a second evaluation, we ran Nighthawk on the full-length, continuous recordings comprising the target dataset. All target audio files were fully annotated by an author or collaborator. For this continuous listening application, we evaluated 1-s windows incremented by 0.2 s and retained predictions that exceeded a score threshold as “detections.” We set score thresholds using the precision-recall curve from the corresponding Balanced Cropped Vocalizations evaluation, selecting a threshold that achieved high precision (0.99) on the target data.

We postprocessed detections from continuous listening in three steps: first, we enforced taxonomic consistency by only retaining detections that are consistent across species, group (if applicable), family, and order at the given score threshold. Second, to reduce false positives, we required at least two detections in a row (both exceeding the score threshold) to trigger a detection. Third, we simplified output by merging overlapping detections up to a maximum duration of 5 s per detection.

To quantify performance for continuous listening, we calculated precision and recall across all files using existing annotations. In addition, we manually reviewed predictions in case some detected calls had been missed by the original annotator. Here, we focused on evaluations of the Passeriformes class because it broadly captures the model’s ability to separate flight calls from background noise.

### Part 2.3: Well-represented target dataset

The first target dataset came from central Pennsylvania, USA (PA) (dataset 18 in Table S1, hereafter “PA dataset”). These recordings were made in the northeast US (a well-represented region in our training dataset), but >200 km from our other large datasets (e.g. datasets 5, 6, 7, 8, 9, 10). The PA dataset consists of 119 hours of annotated audio, from which we extracted 2069 annotated calls and randomly sampled 5064 audio clips without flight calls. We used 3532 of these annotations for additional model training and set aside 3600 for evaluation as described below.

We then constructed target models using three different strategies (Part 2 in Figure 1). First, we trained a new model on non-PA Core training data using the same hyperparameters as the Core model. We refer to this as the “Naive” model. It is similar to the Core model, but its training data includes no PA data. We then experimented with two approaches to fine-tuning a target model to perform well on PA data. In approach 1, we trained a Nighthawk model specifically on the PA data not used for evaluation (3532 annotations) (hereafter the “Tuned” model). In approach 2, we trained two Nighthawk models in which we took the data used to train the Naive model and augmented it with PA data (3532 annotations). To emphasize performance on PA data, we employed a custom batch construction strategy where we filled half of each batch with PA data. For the first model (hereafter the “Batch–More” model), we filled half of each batch from the PA data used in the Tuned model; in the second (hereafter the “Batch–Less” model), we filled half of each batch from a smaller pool of 1000 examples of PA data.

For Tuned, Batch–More, and Batch–Less models, we started training by initializing model weights to those of the Naive model. For the Tuned model, we performed 30 epochs of training using stochastic gradient descent optimization with cosine learning rate decay starting at a learning rate of 0.001. For Batch–More and Batch–Less, we used the same procedure with one adjustment: we trained for 15 epochs because each epoch consisted of approximately twice as many examples due to the batch construction strategy. Based on our results, we also introduced a version of the Batch–More and Batch–Less models trained for only 1 epoch instead of 15 because our results suggested that 1 epoch might be sufficient.

### Part 2.4: Under-represented target dataset

We next used a target dataset from a region poorly represented in the Core data: White Sands Missile Range, New Mexico, USA (dataset 17 in Table S1, hereafter “NM dataset”). Recording locations at White Sands Missile Range are >1000 km from the recording locations of our other large audio datasets. The NM dataset consists of 45 hours of audio annotated primarily at the order level, from which we extracted 2174 annotated calls and randomly generated 5061 clips of audio without flight calls. We used 3412 of these annotations for additional fine-tuning and set aside 3823 for evaluation.

To construct the NM dataset, we followed the same procedure as for PA. The NM dataset is annotated primarily at the order level (virtually all Passeriformes), so for this analysis we only evaluated performance for the Passeriformes class.

### Part 2.5: Out-of-domain target dataset

Nearly all our Core training data come from North America, but many of the migratory species we record in North America have migration routes that traverse Central and South America. Therefore, Nighthawk could be useful in these out-of-domain areas, although it would likely encounter soundscapes that are very different from those in the Core dataset (e.g., different ambient noises and bird species).

We used a target dataset from six recording stations in Colombia, located in or near the cities of Bogotá, Barrancabermeja, and San José del Guaviare (hereafter “CO dataset”). Some data from these locations are included in datasets 12 & 19 (Table S1), so we excluded these datasets from training and trained a new model on the remaining data (“Naive” model). The CO dataset consists of 51 hours of annotated audio, annotated primarily at the order level, from which we extracted 5650 annotated calls and randomly generated 10132 clips of audio without flight calls. We used 9990 of these annotations for additional fine-tuning and set aside 5791 for evaluation.

To construct the CO dataset, we followed the same procedure as for PA and NM.

### Part 3: Evaluating continuous listening for migration monitoring

Lastly, we applied Nighthawk to a large dataset of continuous recordings from an entire autumn migration season; this scenario represents our envisioned use case of Nighthawk for continuous nocturnal listening. This dataset comprises >6000 h of audio from autumn 2015 in Tompkins County, New York, USA (the BirdVox-full-season dataset (Farnsworth et al. 2022)). Previous work used this dataset to relate acoustic measures of bird migration to independent radar and citizen science measures (Van Doren et al. 2023). Their model, called BirdVoxDetect (Salamon et al. 2016, Lostanlen and Salamon 2022, Van Doren et al. 2023), was trained to detect and classify the flight calls of 14 bird species. As a comparison, we ran the Core Nighthawk model on BirdVox-full-season dataset. We calibrated model scores with logistic regression using the Core test dataset (effectively treating it as a validation split for this analysis), and then extracted outputs for the >6000 h of audio. We kept all detections that exceeded a score 0.8.

We then replicated the analysis from Van Doren et al. (2023) using the original analysis code. Briefly, we (a) related acoustic measures of nightly migration intensity to concurrent measures of migration intensity from nearby Doppler radar; (b) compared how well data generated with Nighthawk and BirdVoxDetect explained radar measures of migration; and (c) generated species-specific migration timing estimates from acoustic data and compared them to migration timing estimates derived from citizen science observations. See Van Doren et al. (2023) for details. For analysis (c), we followed that study in including all species with average daily eBird reporting frequencies of at least 1% of checklists and average nightly call rates of at least 0.25 calls per hour.

## Results

### Core model performance (Part 1)

We trained Nighthawk on the Core training dataset and evaluated performance on the Core test dataset using mean average precision (mAP) (Table 1). Nighthawk achieved performance of AP > 0.80 on 50 (61%) species classes, 17 (100%) group classes, 13 (72%) family classes, and 4 (100%) order classes (Table 1). Full performance metrics for all taxa are in Table S3.

**Table 1:**
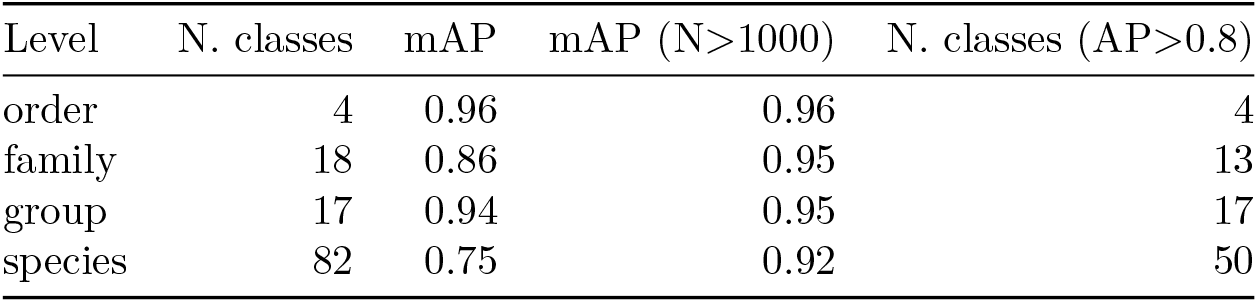
Summary of model performance across taxonomic levels on the Core test dataset. Mean average precision (mAP) scores are given for all evaluated classes and for the subset of classes with at least 1000 training examples.

| Level | N. classes | mAP | mAP (N>1000) | N. classes (AP>0.8) |
| --- | --- | --- | --- | --- |
| order | 4 | 0.96 | 0.96 | 4 |
| family | 18 | 0.86 | 0.95 | 13 |
| group | 17 | 0.94 | 0.95 | 17 |
| species | 82 | 0.75 | 0.92 | 50 |

There was a strong association between training sample size and model performance (Figure 3, Figure S1, Figure S2, Figure S3). Species classes with at least 1000 training examples performed very well (mAP=0.92; n=26). For example, see results for White-throated Sparrow in Figure 4, and for all other classes in Figure S6 through Figure S125. In contrast, species classes with fewer than 200 training examples often performed poorly (mAP=0.42; n=17).

**Figure 3:**
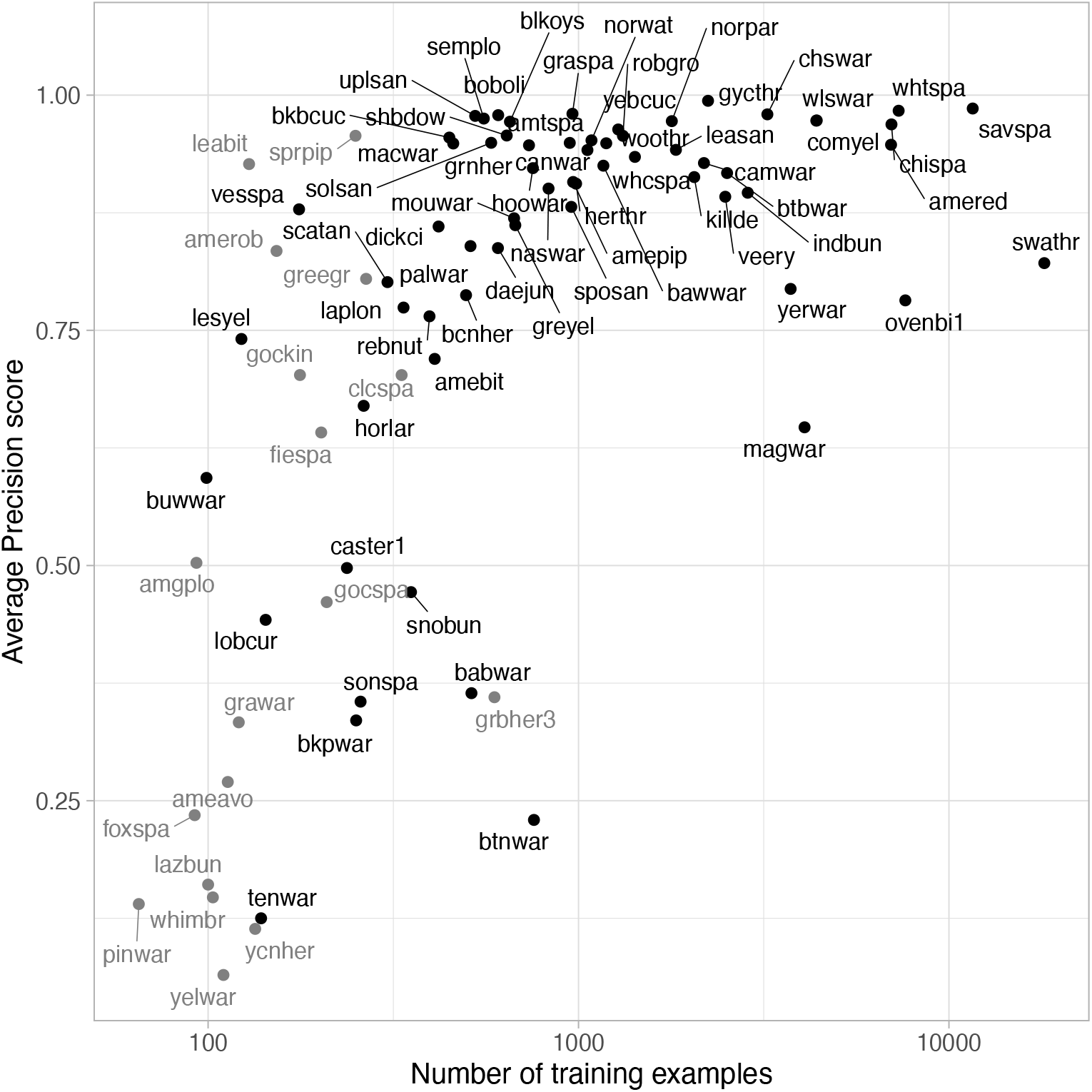
Model performance for species classes on Core test data. Performance is measured by average precision (AP) and plotted against the number of training examples. Species with more training examples generally performed better. Classes plotted in gray have less than 20 testing examples, so their reported performance is subject to increased uncertainty.

**Figure 4:**
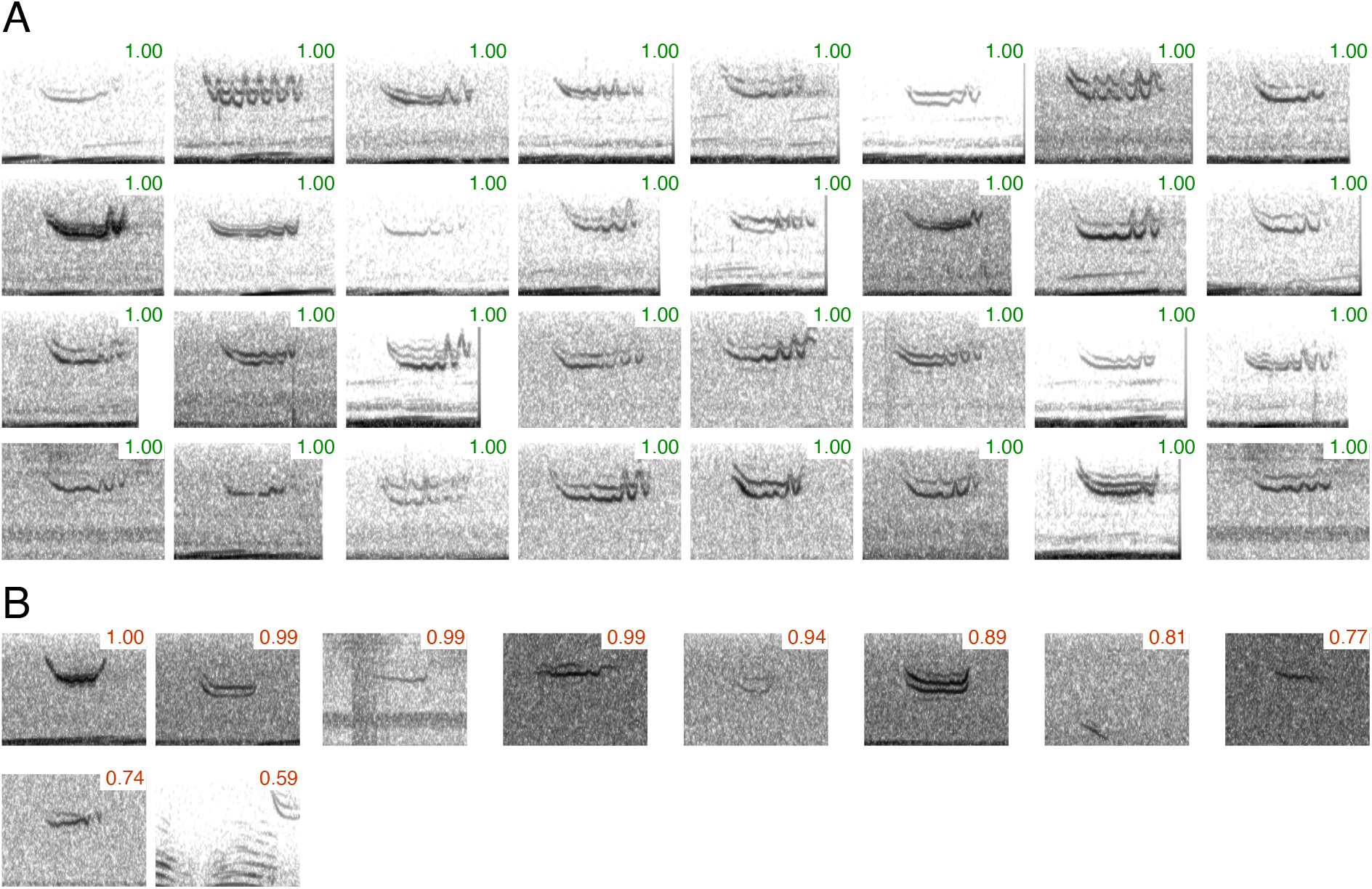
Classification examples for White-throated Sparrow drawn from the test dataset, shown as cropped spectrograms. Numbers in upper-right corners are calibrated probability values returned by the model. (A) Test examples scoring highest for this species. Any incorrect classifications are outlined in red. (B) Incorrectly-classified examples scoring highest for this species (i.e., the most confusing cases for the model). Spectrogram parameters: x-axis: 0-0.3 s; y-axis: 3-11 kHz; window type: Hanning; window length: 200 samples; hop size: 20 samples; dynamic range floor: -60 dB).

### Performance on well-represented target dataset (Part 2.3)

We evaluated Nighthawk on target audio from Pennsylvania, USA (dataset 18 in Table S1) (Figure 5). All Nighthawk models performed well, but the Tuned and Batch–More models performed best on PA data. However, there was a difference in how these two top models performed on the Core test data: Batch–More maintained good performance (−0.008 species mAP compared to Naive), while Tuned lost performance (−0.063 species mAP compared to Naive). This indicates that fine-tuning only on target data came at a cost of lower performance on the Core dataset. See Table S4 and Figure 5.

**Figure 5:**
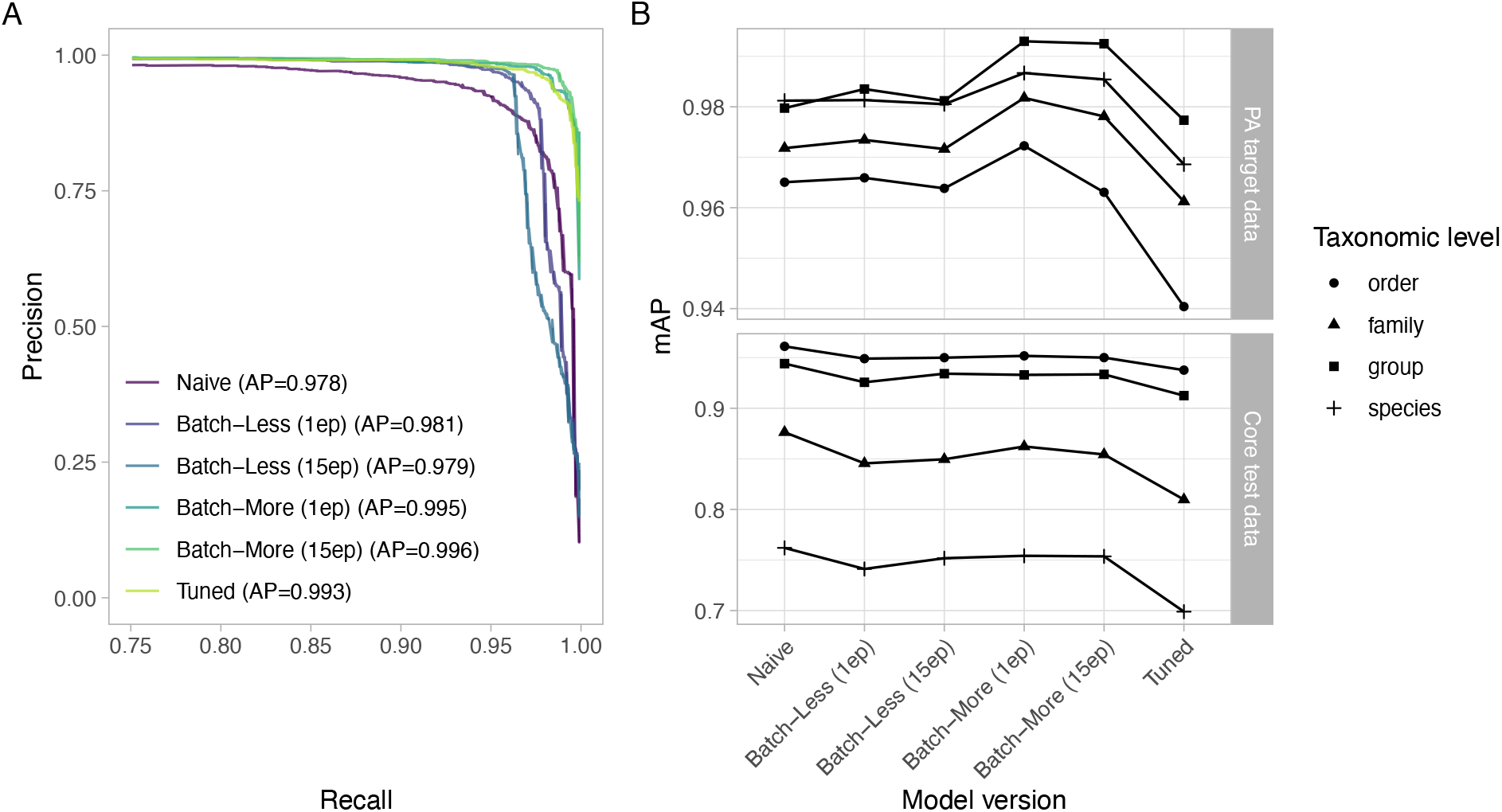
Performance on central Pennsylvania holdout dataset. (A) Precision-recall curves for order Passeriformes. Recall values are displayed only from 0.75-1.00 to emphasize the subtle differences in performance between models. (B) Performance across taxonomic levels. Shown are mean average precision (mAP) scores for taxa with at least 20 examples in the holdout test set.

For the continuous listening evaluation, tuning with Batch–More decreased the false positive rate from 33.4% (N=315) to 0.7% (N=6) while *increasing* the percentage of successfully detected annotator-marked calls from 59% to 78%. Manual review of detections revealed that some calls Nighthawk found had been missed by the annotator. See Table S4.

### Performance on under-represented target dataset (Part 2.4)

We next evaluated Nighthawk on target audio from White Sands Missile Range in New Mexico (NM), USA (dataset 17 in Table S1) (Figure 6). Tuned and Batch–More models performed best on NM data, and substantially better than the Naive model. However, we again saw a difference in how these two top models performed on our original Core test dataset: Batch–More maintained good performance (−0.003 species mAP compared to Naive), while Tuned lost performance (−0.06 species mAP compared to Naive), indicating that fine-tuning only on target data came at a cost of lower performance on the Core dataset. See Table S5 and Figure 6.

**Figure 6:**
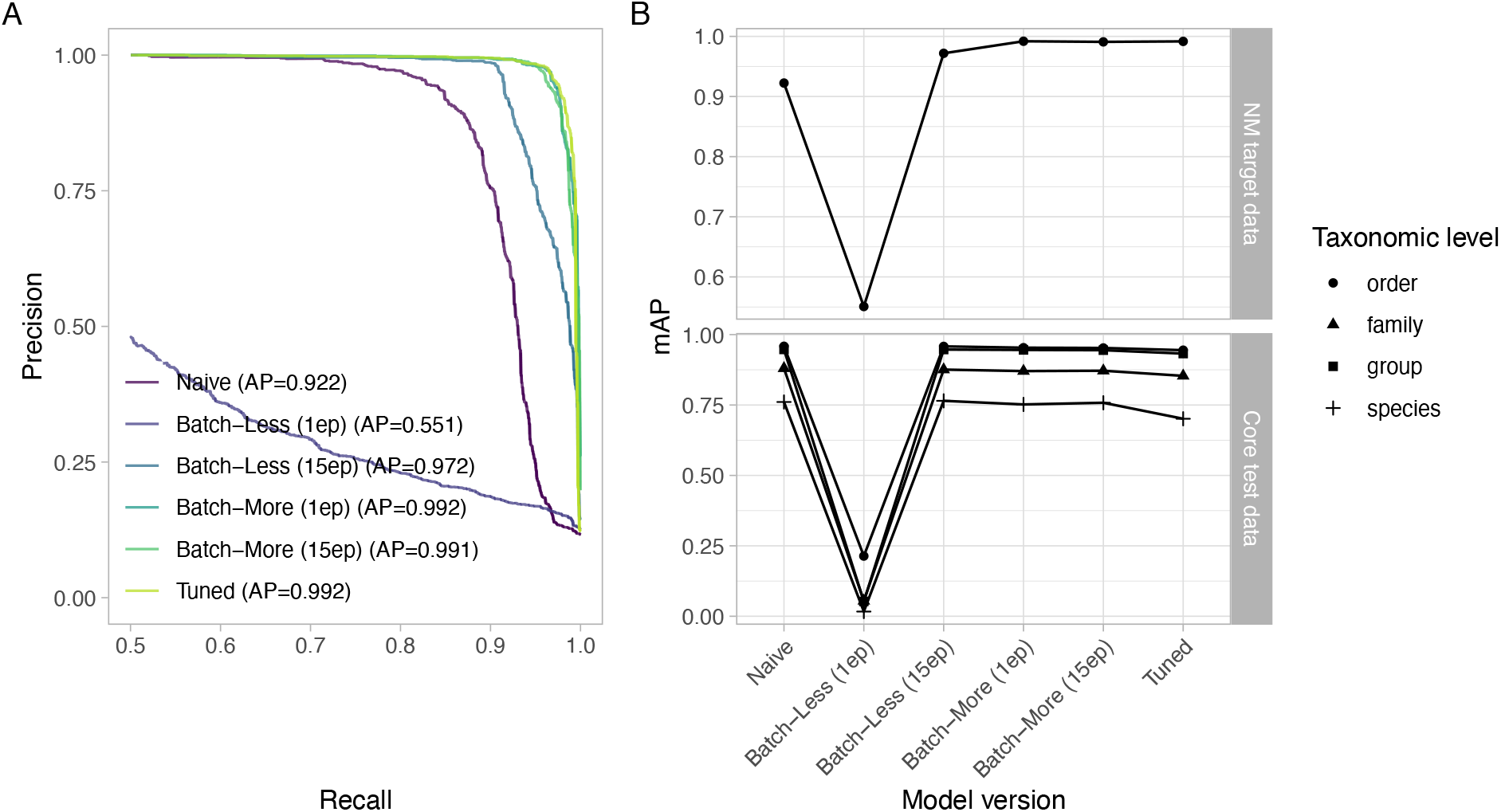
Precision-recall curves for order Passeriformes on dataset from White Sands Missile Range, NM, USA. Note that recall values are displayed only from 0.5-1.00 to emphasize the subtle differences in performance between models.

For the continuous listening evaluation, tuning with Batch–More decreased the false positive rate from 18.8% (N=228) to 3.5% (N=33) while keeping constant the percentage of successfully detected annotator-marked calls (69% and 69%). Manual review of detections again revealed that some calls Nighthawk found had been missed by the annotator. See Table S5.

### Performance on out-of-domain target dataset (Part 2.5)

Our final target evaluation was on target audio from Colombia. Although the Naive model performed poorly, fine-tuning with CO data yielded a high-performing model for both Tuned and Batch–More strategies (Figure 7). Again, there was a difference in how these two top models performed on our Core test dataset: Batch–More lost some performance (−0.037 species mAP compared to Naive), while Tuned substantially lost performance (−0.108 species mAP compared to Naive). See Table S6 and Figure 7.

**Figure 7:**
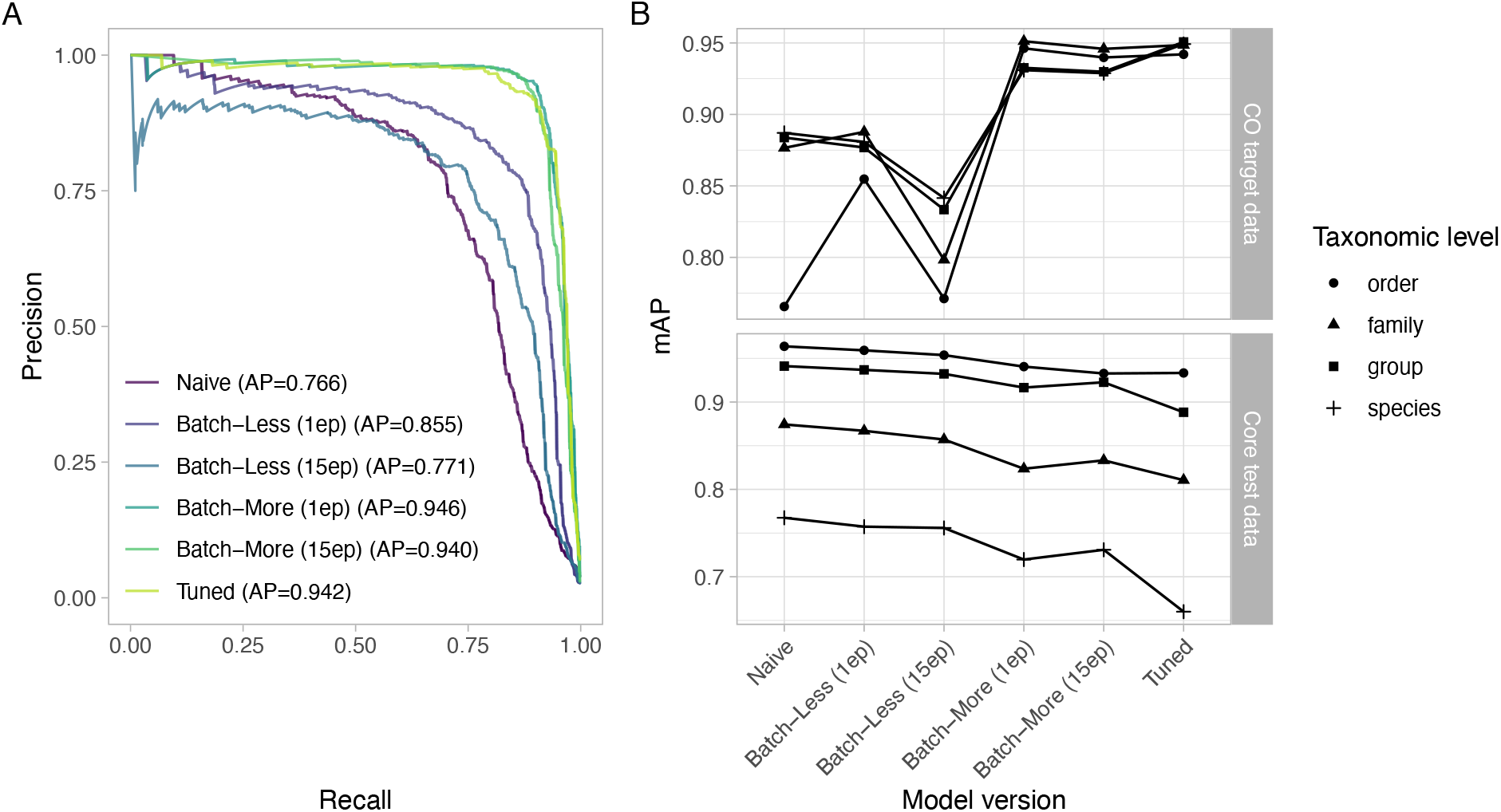
Performance on Colombia holdout dataset. (A) Precision-recall curves for order Passeriformes. (B) Performance across taxonomic levels. Shown are mean average precision (mAP) scores for taxa with at least 20 examples in the holdout test set.

For the continuous listening evaluation, tuning with Batch–More decreased the false positive rate from 24.7% (N=21) to 9.3% (N=25) while *increasing* the percentage of successfully detected annotator-marked calls from 14% to 51%. See Table S6.

### Continuous listening in New York State, USA (Part 3)

We used Core Nighthawk output to model nightly nocturnal migration intensity in central New York State. Acoustic detections alone explained 69% of variation in radar-based migration intensity (Figure 8 A; methods in Van Doren et al. (2023)). Our ability to explain migration intensity using acoustics improved after incorporating wind variables and time of season, which yielded a model explaining 80% of variation in nightly migration intensities (Figure 8 B). Models using Nighthawk detections outperformed otherwise identical models using detections generated by BirdVoxDetect (from Van Doren et al. (2023); Figure 8 C,D). Evaluation on an annotated subset of the dataset (BirdVox-296h: Farnsworth et al. (2021)) for stations in the test dataset yielded a precision of 0.51 and recall of 0.89.

**Figure 8:**
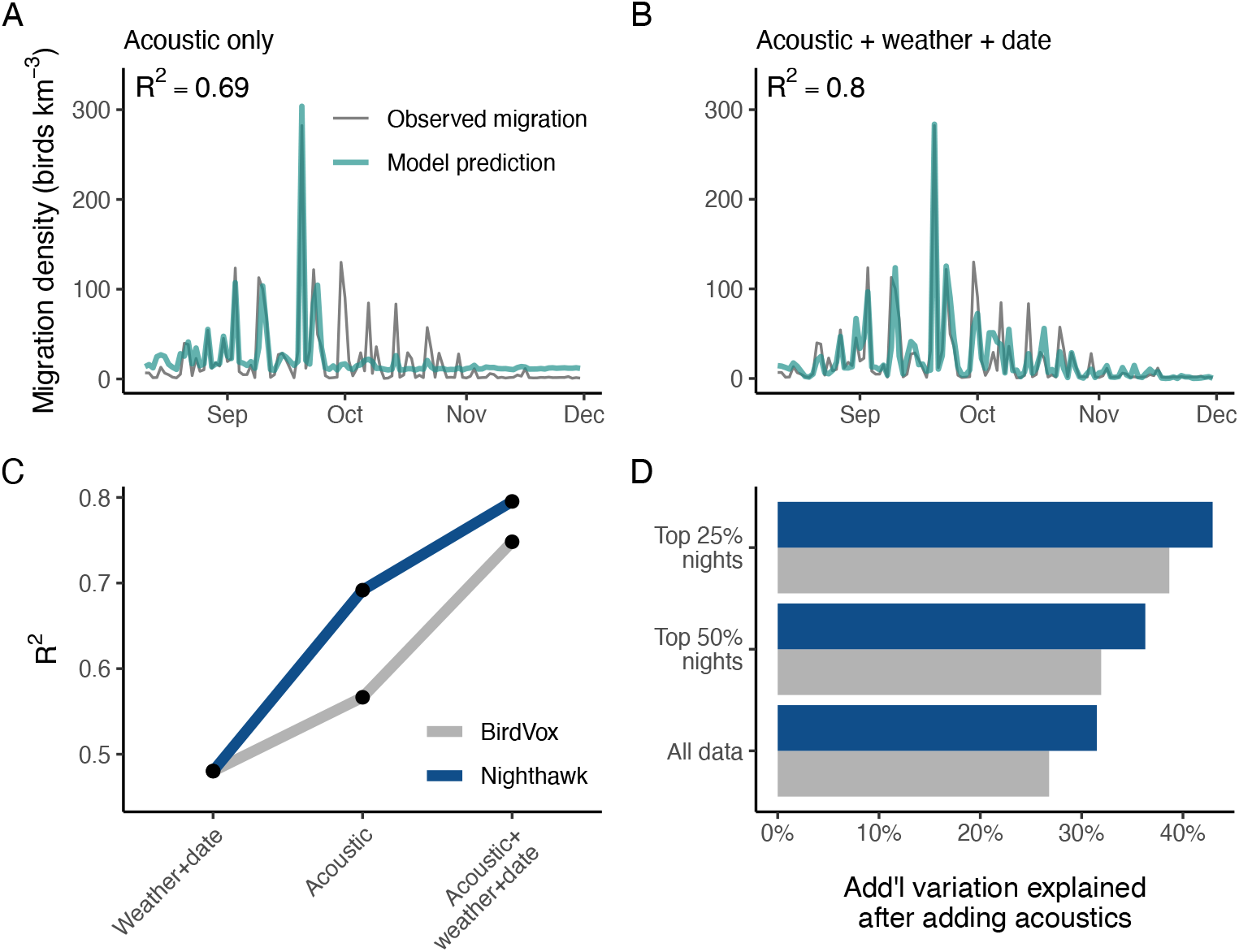
Using Nighthawk to monitor the intensity of nocturnal bird migration over Tompkins County, New York, USA in autumn 2015. (A) Explaining migration intensity with acoustics. Gray line shows migration densities observed by radar; teal lines show cross-validated predictions made with a generalized linear model comprising only flight call counts. (B) Explaining migration intensity with acoustics, weather, and date information. Same as (A), with weather and date added to the generalized linear model. (C) R-squared values for three generalized linear models of migration intensity, comparing acoustic data generated with BirdVoxDetect and Nighthawk. (D) Observed increase in model fit (R-squared) after adding acoustic data to a model including weather and date variables, for acoustic data generated with BirdVoxDetect (gray) and Nighthawk (blue).

Acoustic detections generated with Nighthawk captured fine temporal variation in migration behavior among species (Figure 9 and Figure S4). Nightly time series of acoustic detections showed that many species used airspace over central New York during only a handful of nights in the fall season. See Figure S4 for plots of all included species.

**Figure 9:**
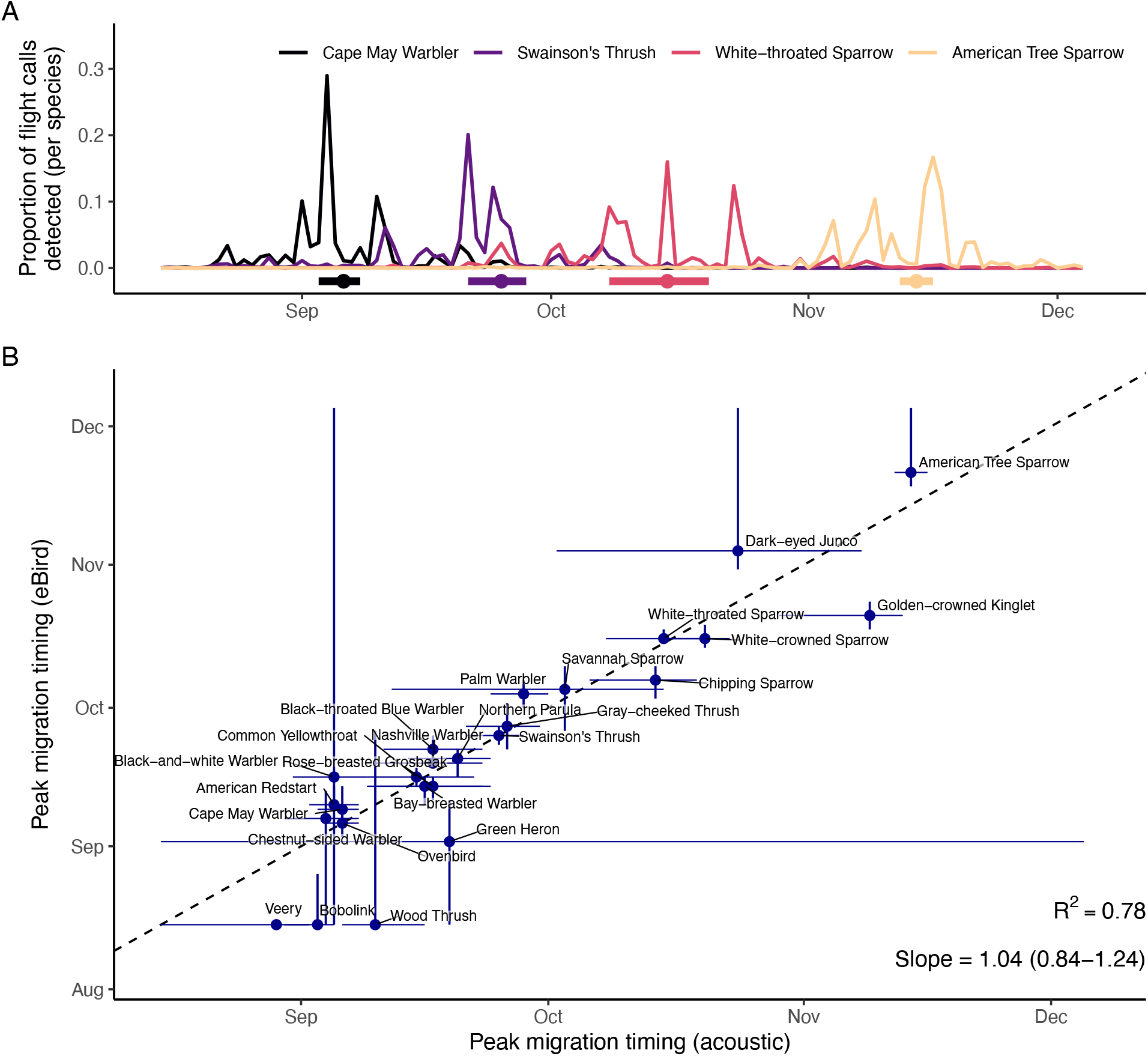
Acoustic monitoring of migration timing at the species level, estimated with acoustic and eBird data, from Tompkins County, New York, USA in fall 2015. (A) Nightly flight call detections made by Nighthawk for four representative migrant species. The y-axis shows the proportion of the season’s calls detected for that species on a given night. For example, 29% of all Cape May Warbler calls recorded during this season were detected on 4 September. Colored lines and points beneath the scatterplot show estimated peak migration timing and 95% confidence interval. (B) Comparison of migration timing at the species level, estimated with acoustic and eBird data. Each point represents one species, with confidence bars showing 95% credible intervals for acoustic and eBird estimates. The dashed line is the identity line; when the acoustic confidence interval (x-axis) intersects the identity line, the 95% CI in acoustic timing overlaps the eBird estimate.

Species-specific migration timing estimates derived from Nighthawk detections closely matched timing estimates from the independent eBird database (Figure 9 B). This close correspondence was quantified by the estimated slope of the relationship between these two measures, which did not significantly differ from 1 (slope = 1.04; 95% CI [0.84-1.24]). Acoustic timing estimates explained 78% of variation in eBird-derived timing estimates. The species included in the timing analysis had a mean AP score on the Core test set of 0.90 (SD 0.13).

## Discussion

Our results demonstrate that Nighthawk performs very well as a nocturnal flight call detector and classifier for dozens of avian taxa. The taxonomic breadth, geographic coverage, and quantity of training data exceed previously published tools (e.g. compare BirdVoxDetect, trained on 14 species and focused on one migration season in one US county; Lostanlen and Salamon (2022); Lostanlen et al. (2019); Van Doren et al. (2023)). Taxa with at least 1000 training examples consistently performed well, though performance was variable for classes with less training data. Total Nighthawk detections accurately captured nightly nocturnal migration intensity and species phenology in our migration monitoring evaluation. The target dataset analyses show that the Core Nighthawk model performed well on data from North American locations not included in model training, and that incorporating modest amounts of additional annotated audio (50-120 h) can yield high performance on custom datasets within North America and beyond.

Our analyses also show that gathering meaningful data about bird migration with acoustics does not require a perfect system. In our migration monitoring evaluation (Part 3), the combination of acoustics, weather, and date explained 80% of nightly variation in migration intensity—and acoustics alone explained 69%. These associations were drawn from a Nighthawk run that achieved a precision of “only” 0.51 (recall: 0.89) on a subset of these data. The results outperform those obtained with BirdVoxDetect and analyzed using identical methods. This result is encouraging because it suggests that even moderately skilled flight call classifiers may yield high quality data if the objective is to measure overall migration passage rates.

Our migration monitoring evaluation (Part 3) also highlights that the bulk of many migratory species used airspace over a given location during only a small number of nights. For example, 29% of all Cape May Warbler detections occurred on 4 September, and 38% of all White-throated Sparrow detections occurred on three dates (8, 15, and 23 October) (Figure 9 A). Previous studies using radar observations have shown that the bulk of avian migrants pass a given location on only a handful of nights in a given season (Horton et al. 2021). Our results suggest that this pattern may also hold true at the species level. This finding is particularly relevant for species of concern, as it suggests that most migratory species utilize a given aerial habitat during only a small number of nights each season—nights that may represent particularly important targets of conservation action (e.g., reducing light pollution or pausing wind turbines).

Species-specific migration timing estimates derived from acoustics (Part 3) closely matched those derived from independent citizen science observations, validating the accuracy of acoustic measures. The slope of the relationship between acoustic and eBird-derived timing measures was not distinguishable from 1, but there were a handful of species that showed larger deviations. Of particular interest are Veery (*Catharus fuscescens*) and Wood Thrush (*Hylocichla mustelina*), species that are difficult to detect visually during migration periods. eBird observations peaked at the very start of August and declined smoothly thereafter, likely reflecting dwindling detection of breeding birds and negligible detection of migrants. However, with acoustics, we were able to identify the subsequent passage periods of these secretive species. Another advantage of acoustic monitoring over citizen science data is a standardization of sampling effort. Autonomous recording units collect standardized samples, while citizen scientists submit varying numbers of checklists from varying locations from day to day. Even in Tompkins County, NY, one of the most heavily birded counties in the world, the number of checklists submitted to eBird in autumn 2015 varied greatly from day to day and declined at the beginning and end of the season (Figure S5). For long-term analyses over many years (e.g. Fink et al. (2020)), such variation can be overcome, but daily variation in observer effort poses a much greater challenge to daily analyses of migrating birds. For applications that require standardized estimates of nocturnal migration at fine taxonomic levels, acoustic monitoring may have great utility.

The Core Nighthawk model achieved AP > 0.8 on 50 species, highlighting its potential for monitoring entire communities of birds on the move. However, there were some species where the model distinctly underperformed, even with robust training sample sizes (Figure 3). These species included Magnolia Warbler (*Setophaga magnolia*), Black-throated Green Warbler (*Setophaga virens*), Bay-breasted Warbler (*Setophaga castanea*), Blackpoll Warbler (*Setophaga striata*), and Song Sparrow (*Melospiza melodia*). These species share an important characteristic: their calls are very similar to those of other species and are readily confused by the model. Fortunately, our modeling approach explicitly accounted for species-level identification uncertainty by joining similar-sounding species in “groups.” For example, the “ZEEP” group includes Magnolia Warbler, Bay-breasted Warbler, and Blackpoll Warbler, and it is one of the highest-scoring classes in the Core evaluation (AP = 0.97) (Figure S1, Table S3). Similarly, Black-throated Green Warblers belong to the “DBUP” group (AP = 0.91), and Song Sparrows belong to the “SFHS” group (AP = 0.97). All groups achieved AP scores > 0.80 on Core test data, further highlighting the usefulness of the “group” construction: even if Nighthawk is not able to accurately classify some vocalizations at the species level, it can still confidently classify these signals to a small group of closely-related species, maximizing information gain.

We tested Nighthawk on recordings from three target locations outside the Core training dataset: Pennsylvania, New Mexico. and Colombia. Performance of “Naive” models on target data varied; Naive models performed best on in-domain data from central Pennsylvania and worst on out-of-domain data from Colombia. However, in all cases, we successfully fine-tuned a high-performing model using a modest amount of annotated audio data from target locations. Performance on target data after fine-tuning was comparable to (or better than) performance on locations included in the Core training set. We observed dramatic performance improvements when using a custom batch construction strategy in which half of each batch came from the target dataset. Remarkably, Batch–More models appeared to need only one epoch of fine-tuning to achieve excellent performance on target data. Although we could achieve similar, if not better, performance by training *only* on target data (“Tuned” models), the Batch–More models retained better performance on the Core test set, and sometimes on the target dataset. This behavior could be advantageous in cases where a researcher tunes Nighthawk for a particular target dataset but may not have a fully representative sample of annotated audio. In these cases, custom batch construction could boost performance by maintaining skill on species not included in the target training data. For all three target datasets, Nighthawk detected a meaningful number of calls that had been overlooked during the original annotation process, thus exceeding the abilities of skilled humans. Annotating hours of continuous audio is a tedious process, and humans may miss, for example, faint vocalizations that Nighthawk is able to detect. These results highlight both the difficulty of constructing a high quality evaluation split and the advantages of systems that integrate both humans and machines in iterative data processing pipelines (Branson et al. 2014). Although it may be tempting to just “set the computer loose” on a dataset, human input and quality control is still essential to producing a high performing system. In the case of evaluating performance on target datasets, having a human review the final Nighthawk output drastically changed our interpretation of performance. For example, on PA target data our estimate of continuous-listening precision on order Passeriformes went from 0.73 before human review to 0.99 after review. Thus, including a human in the loop changed our view of the Nighthawk system from one that performed poorly (precision = 0.73) to one that performed impressively well (precision = 0.99).

### Using Nighthawk

By monitoring the vocalizations of actively migrating birds, Nighthawk provides a detailed window onto nocturnal bird migration that is not presently attainable by other means (e.g., radar or citizen science). Important advantages of this method include its ability to collect data at simultaneously fine spatial, temporal, and taxonomic resolutions. Scientists, managers, and practitioners could use acoustic monitoring for a number of applications, including: monitoring migration passage at wind farms at the species level; studying how bird species use different parts of the landscape during migratory flights; monitoring the changing arrival, departure, and passage times of species susceptible to climate change; and revealing previously unknown migration routes and behaviors. These applications could be most important in areas lacking adequate radar or citizen science coverage.

Our results suggest that Nighthawk’s Core model will function well for monitoring nocturnal bird migration in much of North America, especially in the eastern half of the continent. Researchers with their own target datasets may experience meaningful benefits from fine-tuning Nighthawk with a modest amount of additional training data drawn from the target dataset. In our analyses of target data, annotators reviewed approximately 50-120 hours of audio. To achieve the greatest benefit from fine-tuning, and in order to function properly as validation data, researchers should incorporate target training data that is as representative as possible of the broader target dataset, across locations, dates, seasons, and species. In our target datasets, we split annotated data into two approximately equally-sized sets for training and validation. Validation data allow researchers to select score thresholds that fit their needs on a larger dataset. Detection results are sensitive to the choice of threshold; some applications may call for higher precision (e.g., fully autonomous monitoring) and others for higher recall (e.g., locating rare species with some manual review). We intend to continue building our library of annotated recordings and expanding the capabilities of Nighthawk to more species and locations.

## Acknowledgements

Special thanks to David Benvent, William R. Evans, Mike Farmer, Jenn Foote, Joe Gyekis, Jacob Job, Blaine Landsborough, Michael Lanzone, Debbie Leick, Nicole Liao, Jay McGowan, Sara R. Morris, Dan Men-nill, Drew Weber, Ryan Zucker, and countless contributors to the Macaulay Library (macaulaylibrary.org) for providing audio recordings and annotations. Thanks to Steve Kelling, Juan Pablo Bello, Aurora Cramer, Vincent Lostanlen, and Justin Salamon for their work on the BirdVox effort and for helpful feedback on this project. Thanks to Benjamin Hoffman and Harold Mills for invaluable technical contributions, expertise, and assistance. This work was funded by the Cornell Lab of Ornithology, Actions at EBMF, and the Cornell Presidential Postdoctoral Fellowship.

## Author Contributions

Benjamin M. Van Doren conceived of the study, acquired funding and data, performed analyses, and wrote the first manuscript draft. Grant Van Horn created the original Merlin Sound ID model and oversaw this study. Andrew Farnsworth shaped the study and contributed audio data and annotations. Kate Stone, Dylan Osterhaus, and Jacob Drucker also provided audio data and annotations. All authors contributed to the final written draft of the manuscript. The authors declare no conflicts of interest.

## Data Availability

The Nighthawk model, trained on the Core dataset presented here, is available for download on GitHub (https://github.com/bmvandoren/Nighthawk/), along with Python code demonstrating its proper use. We welcome questions, feedback, and collaboration inquiries by email to the corresponding author.

Users can also use Nighthawk by installing the program Vesper (https://github.com/HaroldMills/Vesper) and using a plugin (https://github.com/HaroldMills/vesper-nighthawk). Vesper is designed for the management and processing of audio recordings for nocturnal bird migration monitoring and is maintained by Harold Mills (https://github.com/HaroldMills).

## References

Bauer, S., J. Shamoun-Baranes, C. Nilsson, A. Farnsworth, J. Kelly, D. R. Reynolds, A. M. Dokter, J. Krauel, L. B. Petterson, K. G. Horton, and J. W. Chapman. 2019. The grand challenges of migration ecology that radar aeroecology can help answer. Ecography 42:861–875. (doi:10.1111/ecog.04083).

Bota, G., J. Traba, F. Sardà-Palomera, D. Giralt, and C. Pérez-Granados. 2020. Acoustic Monitoring of Diurnally Migrating European Bee-Eaters Agrees with Data Derived from Citizen Science. Ardea 108. (doi:10.5253/arde.v108i2.a3).

Branson, S., G. Van Horn, C. Wah, P. Perona, and S. Belongie. 2014. The ignorant led by the blind: A hybrid human–machine vision system for fine-grained categorization. International Journal of Computer Vision 108:329.

Cramer, J., V. Lostanlen, A. Farnsworth, J. Salamon, and J. P. Bello. 2020. ICASSP 2020 - 2020 IEEE International Conference on Acoustics, Speech and Signal Processing (ICASSP). Pages 901–905. IEEE, Barcelona, Spain. (doi:10.1109/ICASSP40776.2020.9052908).

Davy, C. M., A. T. Ford, and K. C. Fraser. 2017. Aeroconservation for the fragmented skies. Conservation Letters 10:773–780. (doi:10.1111/conl.12347).

Diehl, R. H. 2013. The airspace is habitat. Trends in Ecology & Evolution 28:377–379. (doi:10.1016/j.tree.2013.02.015).

Dokter, A. M., F. Liechti, H. Stark, L. Delobbe, P. Tabary, and I. Holleman. 2011. Bird migration flight altitudes studied by a network of operational weather radars. Journal of The Royal Society Interface:rsif20100116. (doi:10.1098/rsif.2010.0116).

Evans, W. R., and M. O’Brien. 2002. Flight calls of migratory birds: Eastern North American landbirds [CD-ROM]. Old Bird Inc., Ithaca, New York, USA.

Evans, W. R., and K. V. Rosenberg. 2000. Acoustic monitoring of night-migrating birds: A progress report. Page 15 in R. Bonney, D. N. Pashley, R. J. Cooper, and L. Niles, editors. U.S. Department of Agriculture, Forest Service, Rocky Mountain Research Station, Ogden, UT.

Farnsworth, A. 2005. Flight calls and their value for future ornithological studies and conservation research. The Auk 122:733–746. (doi:10.1093/auk/122.3.733).

Farnsworth, A., S. Kelling, V. Lostanlen, J. Salamon, A. Cramer, and J. P. Bello. 2021. BirdVox-296h: A large-scale dataset for detection and classification of flight calls. (doi:10.5281/zenodo.4603643).

Farnsworth, A., B. M. Van Doren, S. Kelling, V. Lostanlen, J. Salamon, A. Cramer, and J. P. Bello. 2022. BirdVox-full-season: 6672 hours of audio from migratory birds. (doi:10.5281/zenodo.5791744).

Fink, D., T. Auer, A. Johnston, V. Ruiz-Gutierrez, W. M. Hochachka, and S. Kelling. 2020. Modeling avian full annual cycle distribution and population trends with citizen science data. Ecological Applications 30:e02056. (doi:https://doi.org/10.1002/eap.2056).

Fonseca, E., X. Favory, J. Pons, F. Font, and X. Serra. 2020. FSD50K. Zenodo. (doi:10.5281/zenodo.4060432).

Fraser, K. C., K. T. A. Davies, C. M. Davy, A. T. Ford, D. T. T. Flockhart, and E. G. Martins. 2018. Tracking the conservation promise of movement ecology. Frontiers in Ecology and Evolution 6.

Gauthreaux, S. A., C. G. Belser, and D. van Blaricom. 2003. Using a network of WSR-88D weather surveillance radars to define patterns of bird migration at large spatial scales. Pages 335–346 in P. Berthold, E. Gwinner, and E. Sonnenschein, editors. Springer Berlin Heidelberg.

Gayk, Z. G., and D. J. Mennill. 2023. Acoustic similarity of flight calls corresponds with the composition and structure of mixed-species flocks of migrating birds: evidence from a three-dimensional microphone array. Philosophical Transactions of the Royal Society B 378:20220114. (doi:10.1098/rstb.2022.0114).

He, K., X. Zhang, S. Ren, and J. Sun. 2016. Deep residual learning for image recognition. Page 770778.

Horton, K. G., B. M. Van Doren, H. J. Albers, A. Farnsworth, and D. Sheldon. 2021. Near-term ecological forecasting for dynamic aeroconservation of migratory birds. Conservation Biology 35:1777–1786. (doi:10.1111/cobi.13740).

Howard, A., M. Sandler, G. Chu, L.-C. Chen, B. Chen, M. Tan, W. Wang, Y. Zhu, R. Pang, V. Vasudevan, and others. 2019. Searching for MobileNetV3. Page 13141324.

Kahl, S., C. M. Wood, M. Eibl, and H. Klinck. 2021. BirdNET: A deep learning solution for avian diversity monitoring. Ecological Informatics 61:101236. (doi:10.1016/j.ecoinf.2021.101236).

Kays, R., M. C. Crofoot, W. Jetz, and M. Wikelski. 2015. Terrestrial animal tracking as an eye on life and planet. Science 348:aaa2478. (doi:10.1126/science.aaa2478).

Landsborough, B. J., J. R. Foote, and D. J. Mennill. 2019. Decoding the ‘zeep’ complex: Quantitative analysis of interspecific variation in the nocturnal flight calls of nine wood warbler species (Parulidae spp.). Bioacoustics 28:555–574. (doi:10.1080/09524622.2018.1509373).

Lanzone, M., E. Deleon, L. Grove, and A. Farnsworth. 2009. Revealing undocumented or poorly known flight calls of warblers (Parulidae) using a novel method of recording birds in captivity. The Auk 126:511–519. (doi:10.1525/auk.2009.08187).

Liechti, F., W. Witvliet, R. Weber, and E. Bächler. 2013. First evidence of a 200-day non-stop flight in a bird. Nature Communications 4:2554. (doi:10.1038/ncomms3554).

Lostanlen, V., and J. Salamon. 2022. BirdVox/birdvoxdetect: 1.0. Zenodo. (doi:10.5281/zenodo.7414934).

Lostanlen, V., J. Salamon, A. Farnsworth, S. Kelling, and J. P. Bello. 2017. BirdVox-full-night: A dataset for avian flight call detection in continuous recordings. Zenodo. (doi:10.5281/zenodo.1205569).

Lostanlen, V., J. Salamon, A. Farnsworth, S. Kelling, and J. P. Bello. 2018. 2018 IEEE international conference on acoustics, speech and signal processing (ICASSP). Pages 266–270. (doi:10.1109/ICASSP.2018.8461410).

Lostanlen, V., J. Salamon, A. Farnsworth, S. Kelling, and J. P. Bello. 2019. Robust sound event detection in bioacoustic sensor networks. PLOS ONE 14:e0214168. (doi:10.1371/journal.pone.0214168).

Morris, S. R., K. G. Horton, A. K. Tegeler, and M. Lanzone. 2016. Individual flight-calling behaviour in wood warblers. Animal Behaviour 114:241–247. (doi:10.1016/j.anbehav.2016.01.027).

Nussbaumer, R., S. Bauer, L. Benoit, G. Mariethoz, F. Liechti, and B. Schmid. 2021. Quantifying year-round nocturnal bird migration with a fluid dynamics model. bioRxiv:2020.10.13.321844. (doi:10.1101/2020.10.13.321844).

Piczak, K. J. 2015. ESC: Dataset for environmental sound classification. Page 10151018. ACM Press. (doi:10.1145/2733373.2806390).

Reynolds, M. D., B. L. Sullivan, E. Hallstein, S. Matsumoto, S. Kelling, M. Merrifield, D. Fink, A. Johnston,W. M. Hochachka, N. E. Bruns, M. E. Reiter, S. Veloz, C. Hickey, N. Elliott, L. Martin, J. W. Fitzpatrick, P. Spraycar, G. H. Golet, C. McColl, C. Low, and S. A. Morrison. 2017. Dynamic conservation for migratory species. Science Advances 3:e1700707. (doi:10.1126/sciadv.1700707).

Rosenberg, K. V., A. M. Dokter, P. J. Blancher, J. R. Sauer, A. C. Smith, P. A. Smith, J. C. Stanton, A. Panjabi, L. Helft, M. Parr, and P. P. Marra. 2019. Decline of the North American avifauna. Science 366:120–124. (doi:10.1126/science.aaw1313).

Salamon, J., J. P. Bello, A. Farnsworth, M. Robbins, S. Keen, H. Klinck, and S. Kelling. 2016. Towards the automatic classification of avian flight calls for bioacoustic monitoring. PLOS ONE 11:e0166866. (doi:10.1371/journal.pone.0166866).

Salamon, J., C. Jacoby, and J. P. Bello. 2014. A dataset and taxonomy for urban sound research. Page 10411044. Association for Computing Machinery, New York, NY, USA. (doi:10.1145/2647868.2655045).

Shamoun-Baranes, J., S. Bauer, J. W. Chapman, P. Desmet, A. M. Dokter, A. Farnsworth, H. van Gasteren, B. Haest, J. Koistinen, B. Kranstauber, F. Liechti, T. H. E. Mason, C. Nilsson, R. Nussbaumer, B. Schmid, N. Weisshaupt, and H. Leijnse. 2022. Meteorological Data Policies Needed to Support Biodiversity Monitoring with Weather Radar. Bulletin of the American Meteorological Society 103:E1234–E1242. (doi:10.1175/BAMS-D-21-0196.1).

Shipley, J. R., J. F. Kelly, and W. F. Frick. 2018. Toward integrating citizen science and radar data for migrant bird conservation. Remote Sensing in Ecology and Conservation 4:127–136. (doi:10.1002/rse2.62).

Sugai, L. S. M., T. S. F. Silva, J. W. J. Ribeiro, and D. Llusia. 2019. Terrestrial passive acoustic monitoring: Review and perspectives. BioScience 69:15–25. (doi:10.1093/biosci/biy147).

Sullivan, B. L., J. L. Aycrigg, J. H. Barry, R. E. Bonney, N. Bruns, C. B. Cooper, T. Damoulas, A. A. Dhondt, T. Dietterich, A. Farnsworth, D. Fink, J. W. Fitzpatrick, T. Fredericks, J. Gerbracht, C. Gomes, W. M. Hochachka, M. J. Iliff, C. Lagoze, F. A. La Sorte, M. Merrifield, W. Morris, T. B. Phillips, M. Reynolds, A. D. Rodewald, K. V. Rosenberg, N. M. Trautmann, A. Wiggins, D. W. Winkler, W.-K. Wong, C. L. Wood, J. Yu, and S. Kelling. 2014. The eBird enterprise: An integrated approach to development and application of citizen science. Biological Conservation 169:31–40. (doi:10.1016/j.biocon.2013.11.003).

Van Doren, B. M., G. J. Conway, R. J. Phillips, G. C. Evans, G. C. M. Roberts, M. Liedvogel, and B. C. Sheldon. 2021. Human activity shapes the wintering ecology of a migratory bird. Global Change Biology 27:2715–2727. (doi:https://doi.org/10.1111/gcb.15597).

Van Doren, B. M., and K. G. Horton. 2018. A continental system for forecasting bird migration. Science 361:1115–1118. (doi:10.1126/science.aat7526).

Van Doren, B. M., V. Lostanlen, A. Cramer, J. Salamon, A. Dokter, S. Kelling, J. P. Bello, and A. Farnsworth. 2023. Automated acoustic monitoring captures timing and intensity of bird migration. Journal of Applied Ecology 60:433444. (doi:10.1111/1365-2664.14342).

Van Doren, B. M., D. Sheldon, J. Geevarghese, W. M. Hochachka, and A. Farnsworth. 2015. Autumn morning flights of migrant songbirds in the northeastern United States are linked to nocturnal migration and winds aloft. The Auk 132:105–118. (doi:10.1642/AUK-13-260.1).

