## Supplementary Material for "*Nighthawk*: acoustic monitoring of nocturnal bird migration in the Americas"

Table S1: Data summary by dataset. Train and test columns give the number of annotated audio segments included in each dataset. Annotations may be either flight calls or other non-flight call sounds. Pecentages give the proportion of the total train or test splits. The splits shown were used for training and evaluating the Core model; we modified some of these train and test splits for target analyses (Part 2) as described in the Methods.

| ID | Train | Test | Description | References |
|---|---|---|---|---|
| 1 | 68938 (16%) | 5389 (11%) | Field recordings from Montana. | - |
| 2 | 56990 (13%) | 2300 (4.9%) | Field recordings from eastern North America. | [a] |
| 3 | 36803 (8.6%) | 1123 (2.4%) | FSD-50K: a public sound dataset. | [b] |
| 4 | 30341 (7.1%) | 8215 (17%) | Assets from Macaulay Library. | [c] |
| 5 | 23814 (5.6%) | 6739 (14%) | BirdVox-Full-Night dataset: extracted calls. | [d] |
| 6 | 21838 (5.1%) | 1112 (2.4%) | Field recordings from New York (extracted background noise). | [e] |
| 7 | 20863 (4.9%) | 1120 (2.4%) | BirdVox-Full-Night dataset: extracted background noise. | [d] |
| 8 | 19576 (4.6%) | 2579 (5.5%) | Field recordings from New York. | [a] |
| 9 | 19452 (4.5%) | 7181 (15%) | Field recordings from New York. | [e] |
| 10 | 18177 (4.2%) | 4711 (10%) | BirdVox-296h dataset: 296 h from Ithaca, NY in fall 2015. | [f] |
| 11 | 17722 (4.1%) | 1123 (2.4%) | AudioSet: a public sound dataset. | [g] |
| 12 | 16456 (3.8%) | - | Annotated model results from Colombia. | - |
| 13 | 12678 (3%) | - | Annotated model results from a selection of datasets. | - |
| 14 | 11693 (2.7%) | 1123 (2.4%) | Non-bird assets from the Macaulay Library. | [c] |
| 15 | 9803 (2.3%) | - | Field recordings from Colorado. | - |
| 16 | 7861 (1.8%) | 870 (1.8%) | UrbanSound8K: a public sound dataset. | [h] |
| 17 | 7235 (1.7%) | - | Field recordings from New Mexico. | - |
| 18 | 7133 (1.7%) | - | Field recordings from Pennsylvania. | - |
| 19 | 5601 (1.3%) | - | Annotated model results from Colombia. | - |
| 20 | 4230 (0.99%) | 915 (1.9%) | Captive recordings from PA, NY, and ME. | [i] |
| 21 | 3745 (0.87%) | 852 (1.8%) | Captive and field recordings from AZ, NY, PA, and TX. | - |
| 22 | 3119 (0.73%) | 929 (2%) | Captive recordings from Pennsylvania. | [j] |
| 23 | 1597 (0.37%) | 400 (0.85%) | ESC-50: a public sound dataset. | [k] |
| 24 | 1350 (0.32%) | 329 (0.7%) | Captive recordings from Ontario. | [l] |
| 25 | 994 (0.23%) | 295 (0.62%) | Field recordings of North American sparrows. | - |

[a] http://OldBird.org
[b] Fonseca et al. (2020)
[c] http://macaulaylibrary.org
[d] Lostanlen et al. (2018); Lostanlen et al. (2017)
[e] Van Doren et al. (2015)
[f] Farnsworth et al. (2021)
[g] https://research.google.com/audioset
[h] Salamon et al. (2014)
[i] Morris et al. (2016)
[j] Lanzone et al. (2009)
[k] Piczak (2015)
[l] Landsborough et al. (2019)

Table S2: Species comprising each group. Groups are defined as two or more similar-sounding species that belong to the same family (see Evans and O'Brien 2002). In our modeling framework, each species can belong to at most one group.

| group | species | name | sci_name | family |
|---|---|---|---|---|
| BUNT | blugrb1 | Blue Grosbeak | Passerina caerulea | Cardinalidae |
| BUNT | indbun | Indigo Bunting | Passerina cyanea | Cardinalidae |
| BUNT | lazbun | Lazuli Bunting | Passerina amoena | Cardinalidae |
| BUNT | paibun | Painted Bunting | Passerina ciris | Cardinalidae |
| BUNT | varbun | Varied Bunting | Passerina versicolor | Cardinalidae |
| GROS | bkhgro | Black-headed Grosbeak | Pheucticus melanocephalus | Cardinalidae |
| GROS | robgro | Rose-breasted Grosbeak | Pheucticus ludovicianus | Cardinalidae |
| TANA | scatan | Scarlet Tanager | Piranga olivacea | Cardinalidae |
| TANA | sumtan | Summer Tanager | Piranga rubra | Cardinalidae |
| TANA | westan | Western Tanager | Piranga ludoviciana | Cardinalidae |
| BZWA | chswar | Chestnut-sided Warbler | Setophaga pensylvanica | Parulidae |
| BZWA | comyel | Common Yellowthroat | Geothlypis trichas | Parulidae |
| BZWA | hoowar | Hooded Warbler | Setophaga citrina | Parulidae |
| BZWA | kirwar | Kirtland's Warbler | Setophaga kirtlandii | Parulidae |
| DBUP | btnwar | Black-throated Green Warbler | Setophaga virens | Parulidae |
| DBUP | btywar | Black-throated Gray Warbler | Setophaga nigrescens | Parulidae |
| DBUP | colwar | Colima Warbler | Leiothlypis crissalis | Parulidae |
| DBUP | gchwar | Golden-cheeked Warbler | Setophaga chrysoparia | Parulidae |
| DBUP | grawar | Grace's Warbler | Setophaga graciae | Parulidae |
| DBUP | herwar | Hermit Warbler | Setophaga occidentalis | Parulidae |
| DBUP | lucwar | Lucy's Warbler | Leiothlypis luciae | Parulidae |
| DBUP | naswar | Nashville Warbler | Leiothlypis ruficapilla | Parulidae |
| DBUP | orcwar | Orange-crowned Warbler | Leiothlypis celata | Parulidae |
| DBUP | tenwar | Tennessee Warbler | Leiothlypis peregrina | Parulidae |
| DBUP | towwar | Townsend's Warbler | Setophaga townsendi | Parulidae |
| DBUP | virwar | Virginia's Warbler | Leiothlypis virginiae | Parulidae |
| DEWA | norpar | Northern Parula | Setophaga americana | Parulidae |
| DEWA | pinwar | Pine Warbler | Setophaga pinus | Parulidae |
| DEWA | yetwar | Yellow-throated Warbler | Setophaga dominica | Parulidae |
| MWAR | macwar | MacGillivray's Warbler | Geothlypis tolmiei | Parulidae |
| MWAR | mouwar | Mourning Warbler | Geothlypis philadelphia | Parulidae |
| SBUF | btbwar | Black-throated Blue Warbler | Setophaga caerulescens | Parulidae |
| SBUF | buwwar | Blue-winged Warbler | Vermivora cyanoptera | Parulidae |
| SBUF | gowwar | Golden-winged Warbler | Vermivora chrysoptera | Parulidae |
| SBUF | ovenbi1 | Ovenbird | Seiurus aurocapilla | Parulidae |
| SBUF | palwar | Palm Warbler | Setophaga palmarum | Parulidae |
| SBUF | prowar | Prothonotary Warbler | Protonotaria citrea | Parulidae |
| SBUF | swawar | Swainson's Warbler | Limnothlypis swainsonii | Parulidae |
| SBUF | yerwar | Yellow-rumped Warbler | Setophaga coronata | Parulidae |
| ZEEP | babwar | Bay-breasted Warbler | Setophaga castanea | Parulidae |
| ZEEP | bkbwar | Blackburnian Warbler | Setophaga fusca | Parulidae |
| ZEEP | bkpwar | Blackpoll Warbler | Setophaga striata | Parulidae |
| ZEEP | camwar | Cape May Warbler | Setophaga tigrina | Parulidae |
| ZEEP | cerwar | Cerulean Warbler | Setophaga cerulea | Parulidae |
| ZEEP | conwar | Connecticut Warbler | Oporornis agilis | Parulidae |
| ZEEP | kenwar | Kentucky Warbler | Geothlypis formosa | Parulidae |

| | | | | |
|---|---|---|---|---|
| ZEEP | louwat | Louisiana Waterthrush | Parkesia motacilla | Parulidae |
| ZEEP | magwar | Magnolia Warbler | Setophaga magnolia | Parulidae |
| ZEEP | norwat | Northern Waterthrush | Parkesia noveboracensis | Parulidae |
| ZEEP | refwar | Red-faced Warbler | Cardellina rubrifrons | Parulidae |
| ZEEP | woewar1 | Worm-eating Warbler | Helmitheros vermivorum | Parulidae |
| ZEEP | yelwar | Yellow Warbler | Setophaga petechia | Parulidae |
| CCBRS | brespa | Brewer's Sparrow | Spizella breweri | Passerellidae |
| CCBRS | clcspa | Clay-colored Sparrow | Spizella pallida | Passerellidae |
| CUPS | amtspa | American Tree Sparrow | Spizelloides arborea | Passerellidae |
| CUPS | chispa | Chipping Sparrow | Spizella passerina | Passerellidae |
| DESP | bacspa | Bachman's Sparrow | Peucaea aestivalis | Passerellidae |
| DESP | fiespa | Field Sparrow | Spizella pusilla | Passerellidae |
| DESP | henspa | Henslow's Sparrow | Centronyx henslowii | Passerellidae |
| DESP | lecspa | LeConte's Sparrow | Ammospiza leconteii | Passerellidae |
| DESP | nstspa | Nelson's Sparrow | Ammospiza nelsoni | Passerellidae |
| DESP | savspa | Savannah Sparrow | Passerculus sandwichensis | Passerellidae |
| DESP | sstspa | Saltmarsh Sparrow | Ammospiza caudacuta | Passerellidae |
| HSSP | graspa | Grasshopper Sparrow | Ammodramus savannarum | Passerellidae |
| HSSP | seaspa | Seaside Sparrow | Ammospiza maritima | Passerellidae |
| HSSP | vesspa | Vesper Sparrow | Pooecetes gramineus | Passerellidae |
| HSSP | whcspa | White-crowned Sparrow | Zonotrichia leucophrys | Passerellidae |
| SFHS | foxspa | Fox Sparrow | Passerella iliaca | Passerellidae |
| SFHS | gocspa | Golden-crowned Sparrow | Zonotrichia atricapilla | Passerellidae |
| SFHS | harspa | Harris's Sparrow | Zonotrichia querula | Passerellidae |
| SFHS | sonspa | Song Sparrow | Melospiza melodia | Passerellidae |
| SFHS | whtspa | White-throated Sparrow | Zonotrichia albicollis | Passerellidae |
| SWLI | linspa | Lincoln's Sparrow | Melospiza lincolnii | Passerellidae |
| SWLI | swaspa | Swamp Sparrow | Melospiza georgiana | Passerellidae |
| GCBI | bicthr | Bicknell's Thrush | Catharus bicknelli | Turdidae |
| GCBI | gycthr | Gray-cheeked Thrush | Catharus minimus | Turdidae |
| THSH | herthr | Hermit Thrush | Catharus guttatus | Turdidae |
| THSH | swathr | Swainson's Thrush | Catharus ustulatus | Turdidae |
| THSH | veery | Veery | Catharus fuscescens | Turdidae |
| THSH | woothr | Wood Thrush | Hylocichla mustelina | Turdidae |

Table S3: Performance by class, with number of train and test examples. Performance is measured by average precision (AP). Percentages are calculated separately for each taxonomic level. "Negative" examples contained no nocturnal flight calls (e.g. ambient noise or other bird vocalizations).

| Level | Class | AP | Train | Test |
|---|---|---|---|---|
| order | Passeriformes | 0.99 | 197558 (46%) | 30794 (65%) |
| order | Cuculiformes | 0.98 | 845 (0.2%) | 235 (0.5%) |
| order | Charadriiformes | 0.94 | 9606 (2.2%) | 2403 (5.1%) |
| order | Pelecaniformes | 0.94 | 2770 (0.65%) | 671 (1.4%) |
| order | (negative) | - | 204671 (48%) | 11689 (25%) |
| order | (other) | - | 6025 (1.4%) | 1295 (2.7%) |
| order | (unknown) | - | 6534 (1.5%) | 247 (0.52%) |
| family | Parulidae | 0.98 | 84817 (20%) | 10740 (23%) |
| family | Passerellidae | 0.98 | 40466 (9.5%) | 5789 (12%) |
| family | Icteridae | 0.98 | 989 (0.23%) | 784 (1.7%) |

| | | | | |
|---|---|---|---|---|
| family | Cuculidae | 0.98 | 845 (0.2%) | 235 (0.5%) |
| family | Haematopodidae | 0.96 | 176 (0.041%) | 103 (0.22%) |
| family | Turdidae | 0.94 | 23946 (5.6%) | 7183 (15%) |
| family | Ardeidae | 0.94 | 2733 (0.64%) | 663 (1.4%) |
| family | Scolopacidae | 0.94 | 5573 (1.3%) | 1544 (3.3%) |
| family | Bombycillidae | 0.94 | 190 (0.044%) | 34 (0.072%) |
| family | Charadriidae | 0.93 | 2000 (0.47%) | 272 (0.57%) |
| family | Calcariidae | 0.93 | 506 (0.12%) | 333 (0.7%) |
| family | Cardinalidae | 0.92 | 5183 (1.2%) | 910 (1.9%) |
| family | Motacillidae | 0.92 | 1081 (0.25%) | 241 (0.51%) |
| family | Sittidae | 0.78 | 333 (0.078%) | 52 (0.11%) |
| family | Laridae | 0.72 | 1180 (0.28%) | 323 (0.68%) |
| family | Regulidae | 0.70 | 170 (0.04%) | 38 (0.08%) |
| family | Alaudidae | 0.68 | 148 (0.035%) | 238 (0.5%) |
| family | Recurvirostridae | 0.30 | 105 (0.025%) | 35 (0.074%) |
| family | (negative) | - | 204671 (48%) | 11689 (25%) |
| family | (other) | - | 8188 (1.9%) | 1774 (3.7%) |
| family | (unknown) | - | 44709 (10%) | 4354 (9.2%) |
| group | CUPS | 0.99 | 9552 (2.2%) | 1116 (2.4%) |
| group | SWLI | 0.99 | 1746 (0.41%) | 541 (1.1%) |
| group | GCBI | 0.99 | 1455 (0.34%) | 735 (1.6%) |
| group | DESP | 0.98 | 11802 (2.8%) | 686 (1.4%) |
| group | SFHS | 0.97 | 8026 (1.9%) | 702 (1.5%) |
| group | HSSP | 0.97 | 3215 (0.75%) | 1292 (2.7%) |
| group | ZEEP | 0.97 | 28380 (6.6%) | 2681 (5.7%) |
| group | DEWA | 0.96 | 1739 (0.41%) | 573 (1.2%) |
| group | BZWA | 0.96 | 8364 (2%) | 1778 (3.8%) |
| group | GROS | 0.95 | 1061 (0.25%) | 464 (0.98%) |
| group | BUNT | 0.94 | 3160 (0.74%) | 204 (0.43%) |
| group | MWAR | 0.92 | 1106 (0.26%) | 340 (0.72%) |
| group | THSH | 0.91 | 19284 (4.5%) | 5398 (11%) |
| group | DBUP | 0.91 | 13194 (3.1%) | 1378 (2.9%) |
| group | SBUF | 0.90 | 13254 (3.1%) | 1897 (4%) |
| group | CCBRS | 0.84 | 927 (0.22%) | 60 (0.13%) |
| group | TANA | 0.81 | 322 (0.075%) | 98 (0.21%) |
| group | (N/A) | - | 37747 (8.8%) | 8709 (18%) |
| group | (negative) | - | 204671 (48%) | 11689 (25%) |
| group | (unknown) | - | 59004 (14%) | 6993 (15%) |
| species | gycthr | 0.99 | 1367 (0.32%) | 648 (1.4%) |
| species | savspa | 0.99 | 11327 (2.6%) | 595 (1.3%) |
| species | chswar | 0.98 | 3185 (0.74%) | 550 (1.2%) |
| species | whtspa | 0.98 | 7118 (1.7%) | 516 (1.1%) |
| species | graspa | 0.98 | 1038 (0.24%) | 505 (1.1%) |
| species | boboli | 0.98 | 511 (0.12%) | 671 (1.4%) |
| species | uplsan | 0.98 | 465 (0.11%) | 197 (0.42%) |
| species | semplo | 0.98 | 451 (0.11%) | 78 (0.16%) |
| species | chispa | 0.97 | 8510 (2%) | 527 (1.1%) |
| species | comyel | 0.97 | 4209 (0.98%) | 690 (1.5%) |
| species | norpar | 0.97 | 1667 (0.39%) | 542 (1.1%) |
| species | wlswar | 0.97 | 4901 (1.1%) | 521 (1.1%) |
| species | blkoys | 0.97 | 67 (0.016%) | 94 (0.2%) |
| species | robgro | 0.96 | 1053 (0.25%) | 462 (0.98%) |

| | | | | |
|---|---|---|---|---|
| species | yebcuc | 0.96 | 441 (0.1%) | 135 (0.29%) |
| species | shbdow | 0.96 | 469 (0.11%) | 358 (0.76%) |
| species | sprpip | 0.96 | 213 (0.05%) | 24 (0.051%) |
| species | amered | 0.95 | 7041 (1.6%) | 615 (1.3%) |
| species | amtspa | 0.95 | 902 (0.21%) | 531 (1.1%) |
| species | woothr | 0.95 | 883 (0.21%) | 378 (0.8%) |
| species | norwat | 0.95 | 1053 (0.25%) | 169 (0.36%) |
| species | bkbcuc | 0.95 | 404 (0.094%) | 100 (0.21%) |
| species | grnher | 0.95 | 632 (0.15%) | 310 (0.65%) |
| species | macwar | 0.95 | 464 (0.11%) | 145 (0.31%) |
| species | solsan | 0.95 | 510 (0.12%) | 66 (0.14%) |
| species | canwar | 0.94 | 1028 (0.24%) | 317 (0.67%) |
| species | leasan | 0.94 | 1432 (0.33%) | 312 (0.66%) |
| species | bawwar | 0.93 | 1085 (0.25%) | 227 (0.48%) |
| species | camwar | 0.93 | 1613 (0.38%) | 383 (0.81%) |
| species | whcspa | 0.93 | 1678 (0.39%) | 364 (0.77%) |
| species | leabit | 0.93 | 121 (0.028%) | 29 (0.061%) |
| species | btbwar | 0.92 | 2264 (0.53%) | 498 (1.1%) |
| species | hoowar | 0.92 | 753 (0.18%) | 127 (0.27%) |
| species | herthr | 0.91 | 901 (0.21%) | 290 (0.61%) |
| species | amepip | 0.91 | 868 (0.2%) | 217 (0.46%) |
| species | killde | 0.91 | 1400 (0.33%) | 171 (0.36%) |
| species | indbun | 0.90 | 2787 (0.65%) | 150 (0.32%) |
| species | naswar | 0.90 | 836 (0.2%) | 333 (0.7%) |
| species | veery | 0.89 | 1883 (0.44%) | 658 (1.4%) |
| species | vesspa | 0.88 | 476 (0.11%) | 423 (0.89%) |
| species | sposan | 0.88 | 640 (0.15%) | 165 (0.35%) |
| species | mouwar | 0.87 | 641 (0.15%) | 195 (0.41%) |
| species | dickci | 0.86 | 386 (0.09%) | 110 (0.23%) |
| species | greyel | 0.86 | 497 (0.12%) | 128 (0.27%) |
| species | daejun | 0.84 | 523 (0.12%) | 325 (0.69%) |
| species | palwar | 0.84 | 500 (0.12%) | 298 (0.63%) |
| species | amerob | 0.83 | 97 (0.023%) | 41 (0.087%) |
| species | swathr | 0.82 | 15582 (3.6%) | 4072 (8.6%) |
| species | scatan | 0.80 | 249 (0.058%) | 98 (0.21%) |
| species | greegr | 0.80 | 228 (0.053%) | 21 (0.044%) |
| species | yerwar | 0.79 | 3777 (0.88%) | 494 (1%) |
| species | bcnher | 0.79 | 459 (0.11%) | 111 (0.23%) |
| species | ovenbi1 | 0.78 | 6500 (1.5%) | 525 (1.1%) |
| species | laplon | 0.77 | 285 (0.067%) | 197 (0.42%) |
| species | rebnut | 0.76 | 320 (0.075%) | 52 (0.11%) |
| species | lesyel | 0.74 | 91 (0.021%) | 43 (0.091%) |
| species | amebit | 0.72 | 318 (0.074%) | 57 (0.12%) |
| species | clcspa | 0.70 | 506 (0.12%) | 49 (0.1%) |
| species | gockin | 0.70 | 168 (0.039%) | 38 (0.08%) |
| species | horlar | 0.67 | 148 (0.035%) | 238 (0.5%) |
| species | magwar | 0.65 | 4057 (0.95%) | 498 (1.1%) |
| species | fiespa | 0.64 | 198 (0.046%) | 43 (0.091%) |
| species | buwwar | 0.59 | 97 (0.023%) | 75 (0.16%) |
| species | caster1 | 0.50 | 154 (0.036%) | 45 (0.095%) |
| species | amgplo | 0.50 | 49 (0.011%) | 8 (0.017%) |
| species | snobun | 0.47 | 186 (0.043%) | 99 (0.21%) |

| | | | | |
|---|---|---|---|---|
| species | gocspa | 0.46 | 182 (0.043%) | 40 (0.085%) |
| species | lobcur | 0.44 | 72 (0.017%) | 22 (0.046%) |
| species | babwar | 0.36 | 359 (0.084%) | 176 (0.37%) |
| species | grbher3 | 0.36 | 303 (0.071%) | 46 (0.097%) |
| species | sonspa | 0.36 | 550 (0.13%) | 105 (0.22%) |
| species | bkpwar | 0.34 | 249 (0.058%) | 195 (0.41%) |
| species | grawar | 0.33 | 115 (0.027%) | 45 (0.095%) |
| species | ameavo | 0.27 | 59 (0.014%) | 14 (0.03%) |
| species | btnwar | 0.23 | 767 (0.18%) | 171 (0.36%) |
| species | foxspa | 0.23 | 94 (0.022%) | 23 (0.049%) |
| species | lazbun | 0.16 | 123 (0.029%) | 32 (0.068%) |
| species | whimbr | 0.15 | 59 (0.014%) | 26 (0.055%) |
| species | pinwar | 0.14 | 58 (0.014%) | 31 (0.065%) |
| species | tenwar | 0.13 | 145 (0.034%) | 93 (0.2%) |
| species | ycnher | 0.11 | 131 (0.031%) | 16 (0.034%) |
| species | yelwar | 0.06 | 314 (0.073%) | 20 (0.042%) |
| species | (negative) | - | 204671 (48%) | 11689 (25%) |
| species | (other) | - | 12223 (2.9%) | 2616 (5.5%) |
| species | (unknown) | - | 92873 (22%) | 10023 (21%) |

Table S4: Summary of model performance on the PA target dataset.

| Data | Metric | Naive | Batch-Less (1ep) | Batch-Less (15ep) | Batch-More (1ep) | Batch-More (15ep) | Tuned |
|---|---|---|---|---|---|---|---|
| PA | AP: Passeriformes | 0.978 | 0.981 | 0.979 | 0.995 | 0.996 | 0.993 |
| PA | Precision (continuous listening): Passeriformes | 0.498 | | | 0.754 | 0.731 | 0.590 |
| PA | Precision (continuous listening): Passeriformes (reviewed) | 0.666 | | | 0.995 | 0.993 | 0.932 |
| PA | Recall (continuous listening): Passeriformes | 0.591 | | | 0.763 | 0.777 | 0.892 |
| PA | mAP: family (4 taxa) | 0.972 | 0.973 | 0.972 | 0.982 | 0.978 | 0.961 |
| PA | mAP: group (6 taxa) | 0.980 | 0.984 | 0.981 | 0.993 | 0.992 | 0.977 |
| PA | mAP: order (2 taxa) | 0.965 | 0.966 | 0.964 | 0.972 | 0.963 | 0.940 |
| PA | mAP: species (6 taxa) | 0.981 | 0.981 | 0.981 | 0.987 | 0.985 | 0.969 |
| Core | AP: Passeriformes | 0.992 | 0.990 | 0.990 | 0.990 | 0.991 | 0.989 |
| Core | mAP: family (18 taxa) | 0.876 | 0.846 | 0.850 | 0.862 | 0.854 | 0.810 |
| Core | mAP: group (17 taxa) | 0.944 | 0.926 | 0.934 | 0.933 | 0.933 | 0.912 |
| Core | mAP: order (4 taxa) | 0.961 | 0.949 | 0.950 | 0.952 | 0.950 | 0.938 |
| Core | mAP: species (82 taxa) | 0.762 | 0.741 | 0.752 | 0.754 | 0.754 | 0.699 |

Table S5: Summary of model performance on the NM target dataset.

| Data | Metric | Naive | Batch-Less (1ep) | Batch-Less (15ep) | Batch-More (1ep) | Batch-More (15ep) | Tuned |
|---|---|---|---|---|---|---|---|
| NM | AP: Passeriformes | 0.922 | 0.551 | 0.972 | 0.992 | 0.991 | 0.992 |
| NM | Precision (continuous listening): Passeriformes | 0.635 | | | 0.771 | 0.815 | 0.672 |
| NM | Precision (continuous listening): Passeriformes (reviewed) | 0.812 | | | 0.965 | 0.965 | 0.935 |
| NM | Recall (continuous listening): Passeriformes | 0.689 | | | 0.777 | 0.689 | 0.915 |
| Core | AP: Passeriformes | 0.991 | 0.725 | 0.991 | 0.992 | 0.992 | 0.991 |

Table S6: Summary of model performance on the CO target dataset.

| Data | Metric | Naive | Batch-Less (1ep) | Batch-Less (15ep) | Batch-More (1ep) | Batch-More (15ep) | Tuned |
|---|---|---|---|---|---|---|---|
| CO | AP: Passeriformes | 0.766 | 0.855 | 0.771 | 0.946 | 0.940 | 0.942 |
| CO | AP: gycthr | 0.893 | 0.869 | 0.873 | 0.905 | 0.908 | 0.949 |
| CO | AP: swathr | 0.881 | 0.893 | 0.810 | 0.957 | 0.950 | 0.949 |
| CO | Precision (continuous listening): Passeriformes | 0.682 | | | 0.906 | 0.796 | 0.946 |
| CO | Precision (continuous listening): Passeriformes (reviewed) | 0.753 | | | 0.981 | 0.907 | 0.964 |
| CO | Precision (continuous listening): gycthr | 0.767 | | | 0.759 | 0.812 | 0.821 |
| CO | Precision (continuous listening): swathr | 0.925 | | | 0.867 | 0.852 | 0.918 |
| CO | Recall (continuous listening): Passeriformes | 0.142 | | | 0.119 | 0.509 | 0.153 |
| CO | Recall (continuous listening): gycthr | 0.719 | | | 0.688 | 0.812 | 0.719 |
| CO | Recall (continuous listening): swathr | 0.327 | | | 0.359 | 0.387 | 0.191 |
| CO | mAP: family (1 taxa) | 0.877 | 0.888 | 0.798 | 0.951 | 0.946 | 0.949 |
| CO | mAP: group (2 taxa) | 0.884 | 0.877 | 0.833 | 0.933 | 0.930 | 0.951 |
| CO | mAP: order (1 taxa) | 0.766 | 0.855 | 0.771 | 0.946 | 0.940 | 0.942 |
| CO | mAP: species (2 taxa) | 0.887 | 0.881 | 0.841 | 0.931 | 0.929 | 0.949 |
| Core | AP: Passeriformes | 0.992 | 0.991 | 0.990 | 0.987 | 0.986 | 0.971 |
| Core | AP: gycthr | 0.994 | 0.991 | 0.993 | 0.989 | 0.990 | 0.972 |
| Core | AP: swathr | 0.858 | 0.822 | 0.819 | 0.797 | 0.802 | 0.766 |

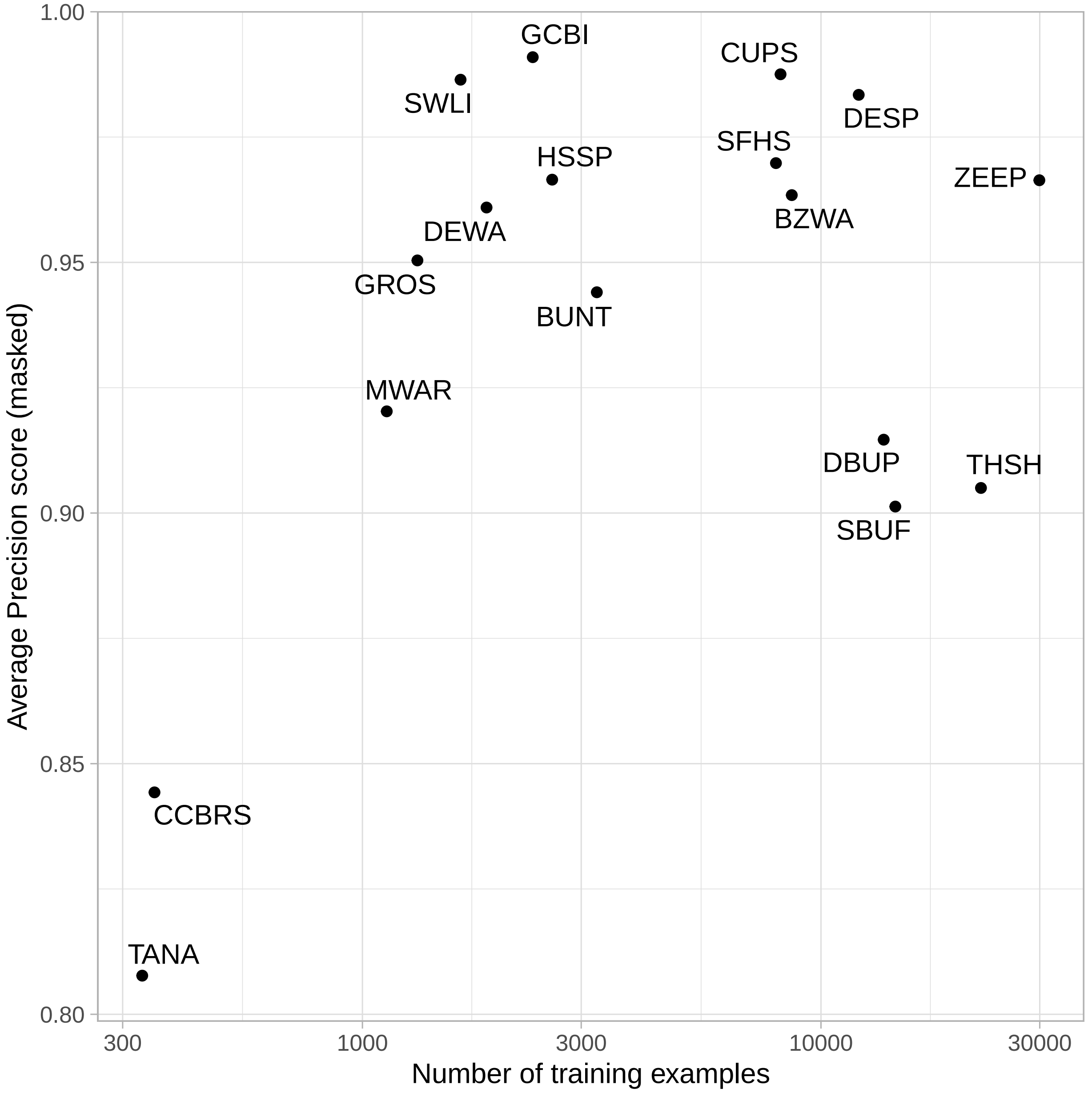


Figure S1: Model performance for group classes on Core test data. Performance is measured by average precision (AP) and plotted against the number of training examples. Groups with more training examples generally performed better.

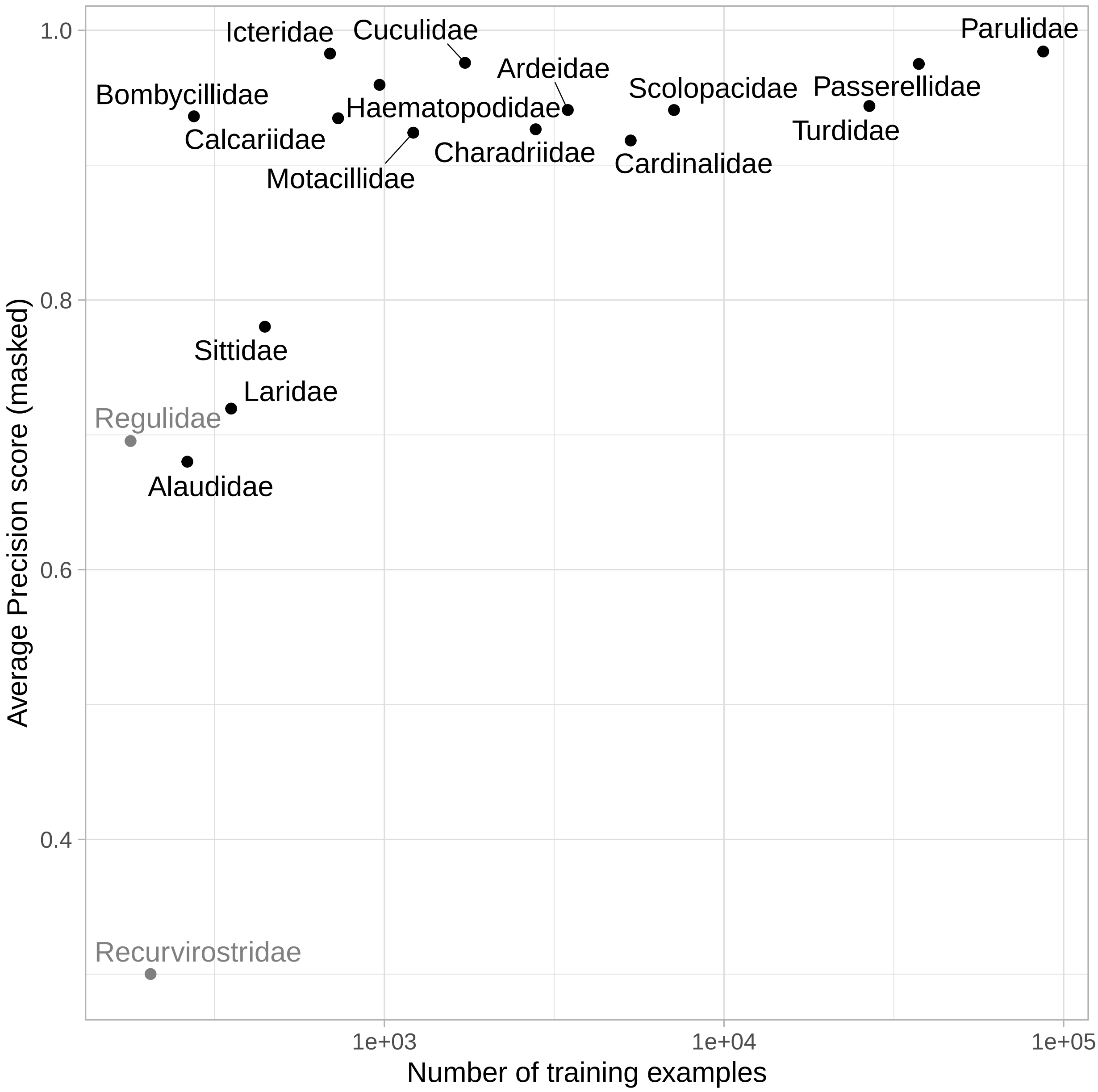


Figure S2: Model performance for family classes on Core test data. Performance is measured by average precision (AP) and plotted against the number of training examples. Families with more training examples generally performed better. Classes plotted in gray have less than 20 testing examples, so their reported performance is subject to increased uncertainty.

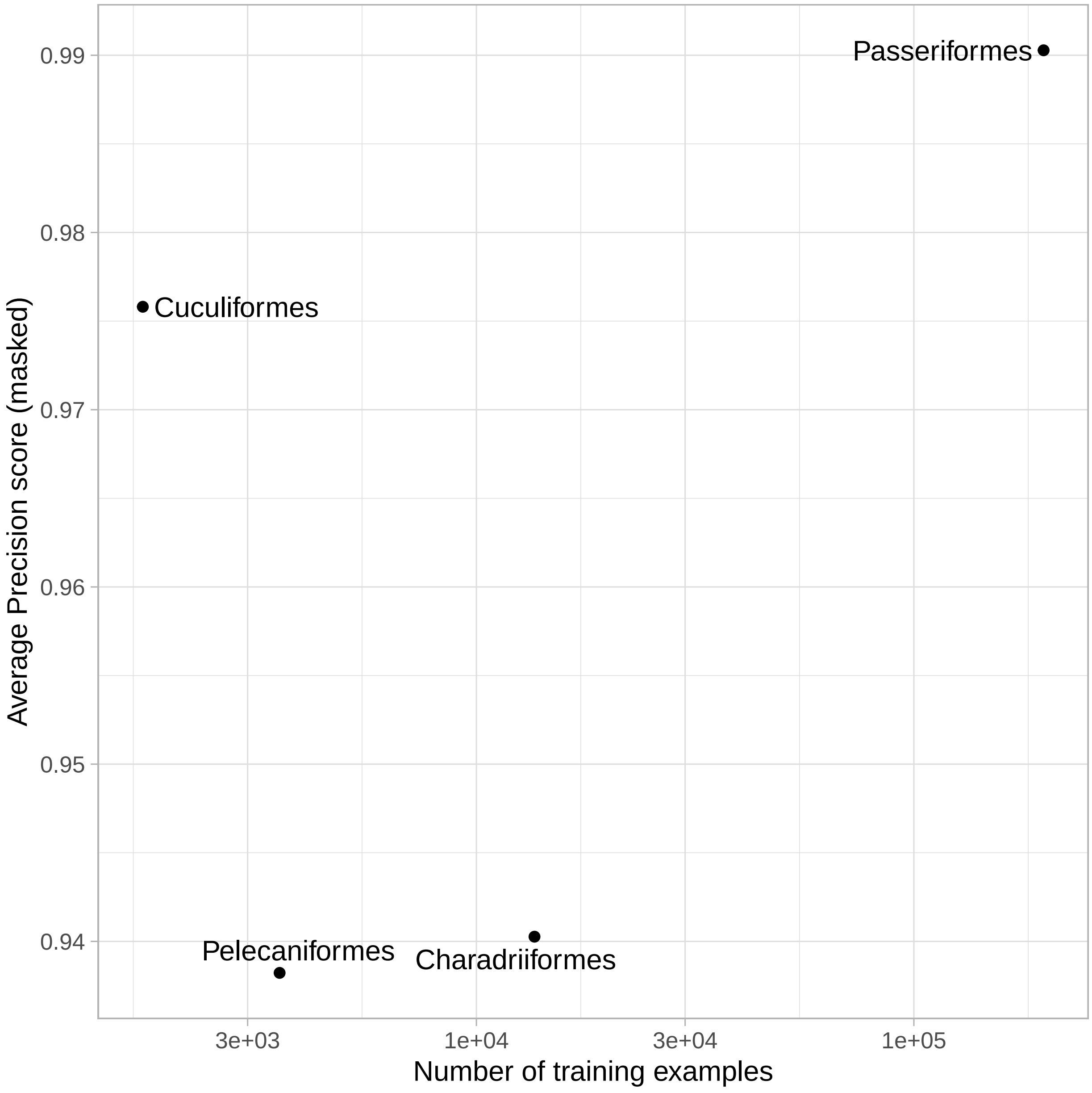


Figure S3: Model performance for order classes on the Core test data. Performance is measured by average precision (AP) and plotted against the number of training examples.

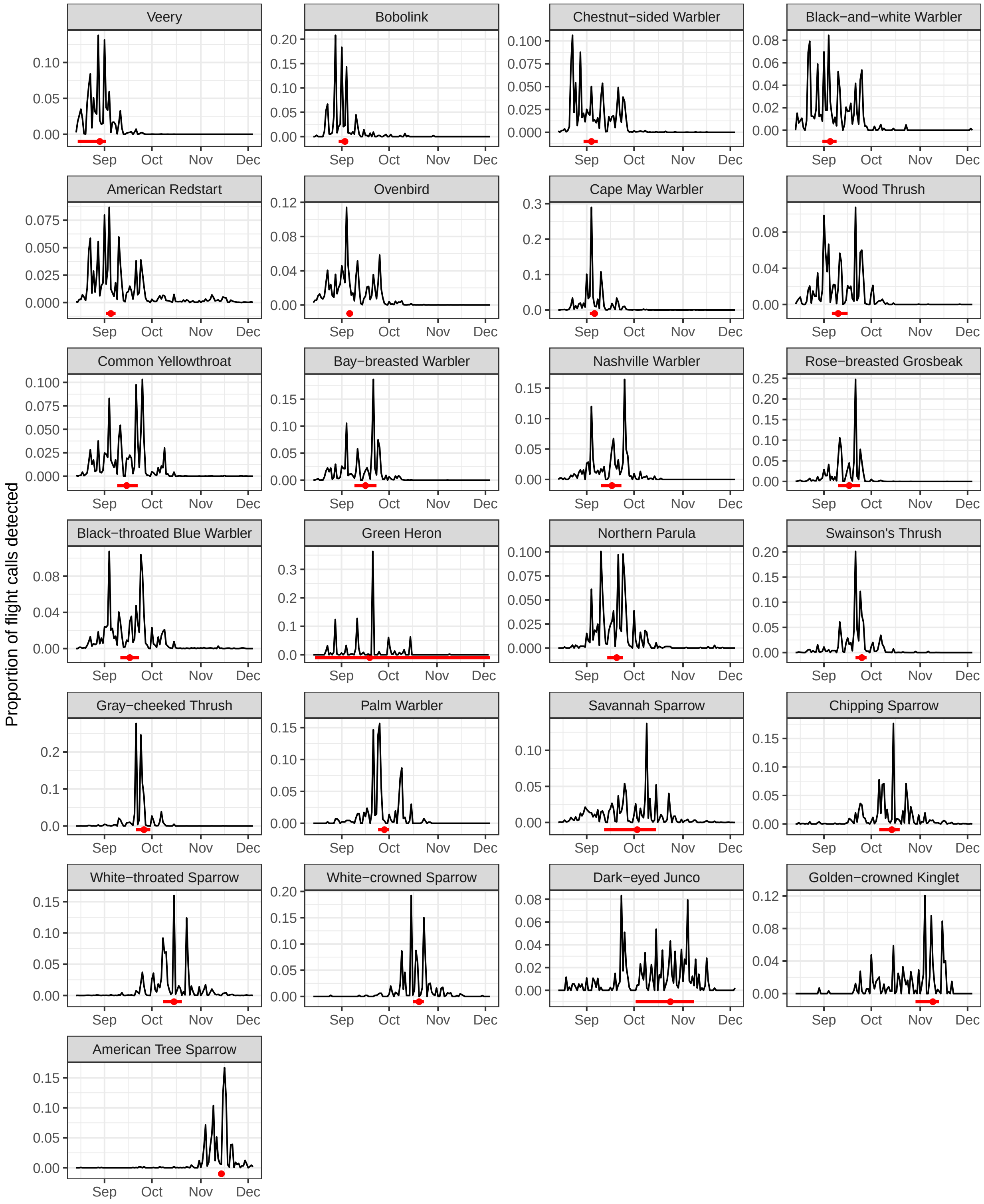


Figure S4: Nightly flight call detections made by Nighthawk in Ithaca, New York, USA during autumn 2015 for all migrant species. The y-axis shows the proportion of the season's calls detected for that species on a given night. For example, 29% of all Cape May Warbler calls recorded during this season were detected on 4 September. Red line and point beneath each plot shows estimated peak migration timing and 95% confidence interval.

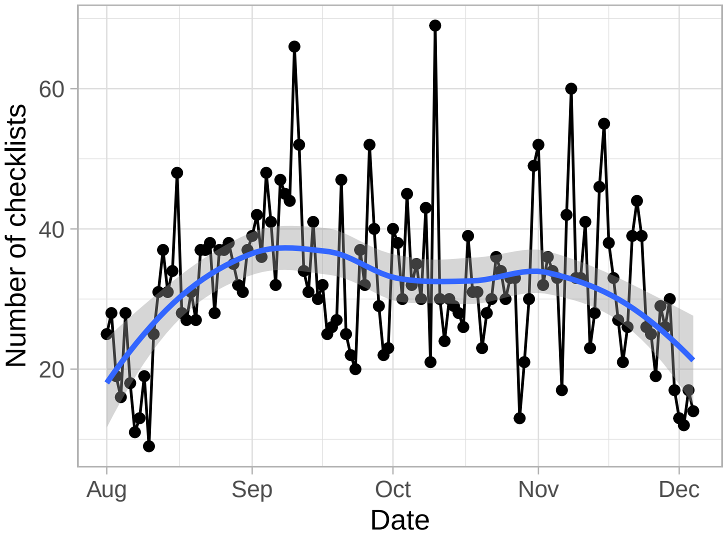


Figure S5: Daily number of complete eBird checklists submitted in Tompkins County, New York, USA in autumn 2015.

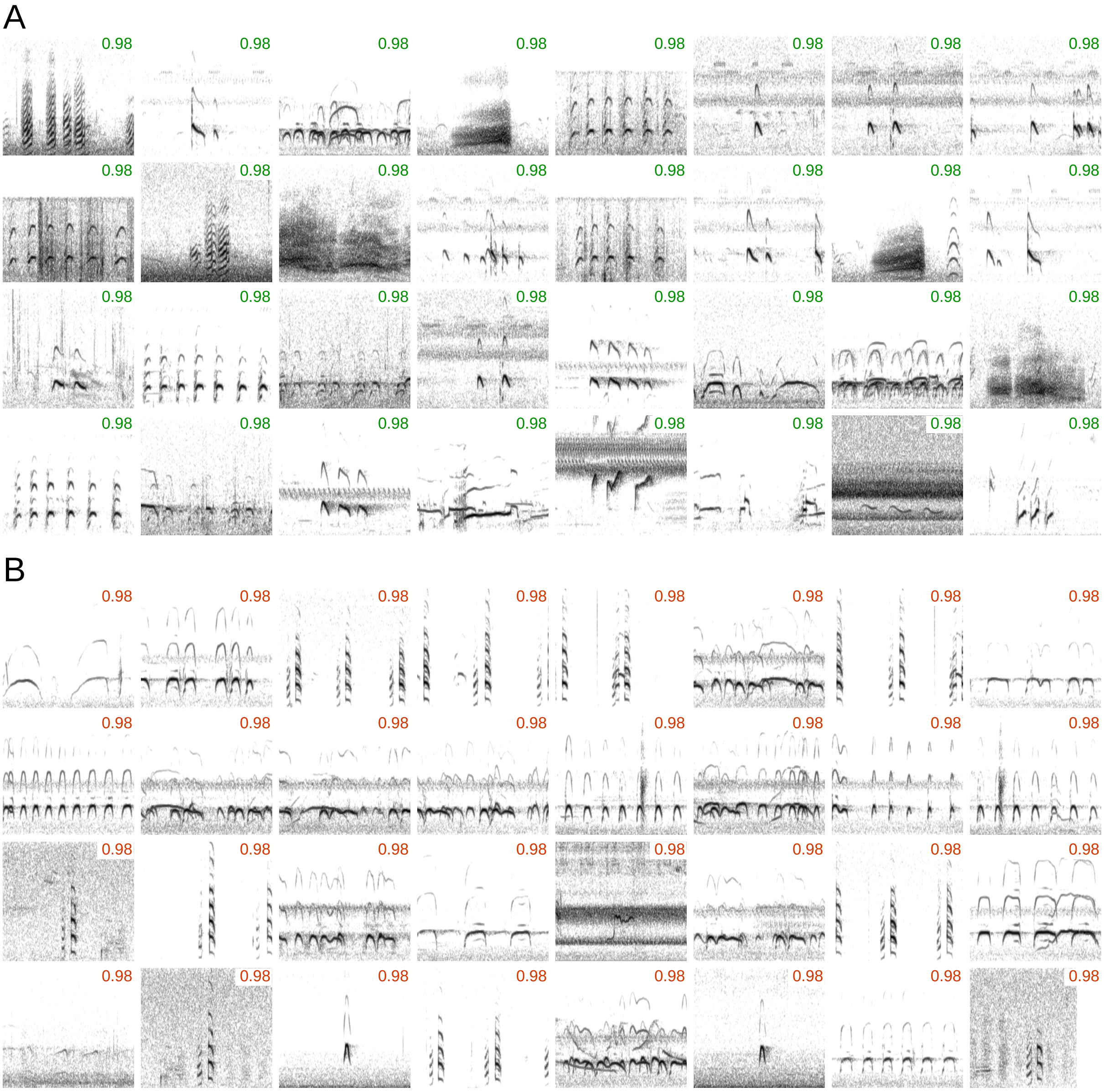


Figure S6: Classification examples for **Charadriiformes** drawn from the test dataset, shown as cropped spectrograms. Numbers in upper-right corners are calibrated probability values returned by the model. (A) Test examples scoring highest for this species. Any incorrect classifications are outlined in red. (B) Incorrectly-classified examples scoring highest for this species (i.e., the most confusing cases for the model). Spectrogram parameters: x-axis: 0-0.8 s; y-axis: 1-10 kHz; window type: Hanning; window length: 300 samples; hop size: 30 samples; dynamic range floor: -50 dB).

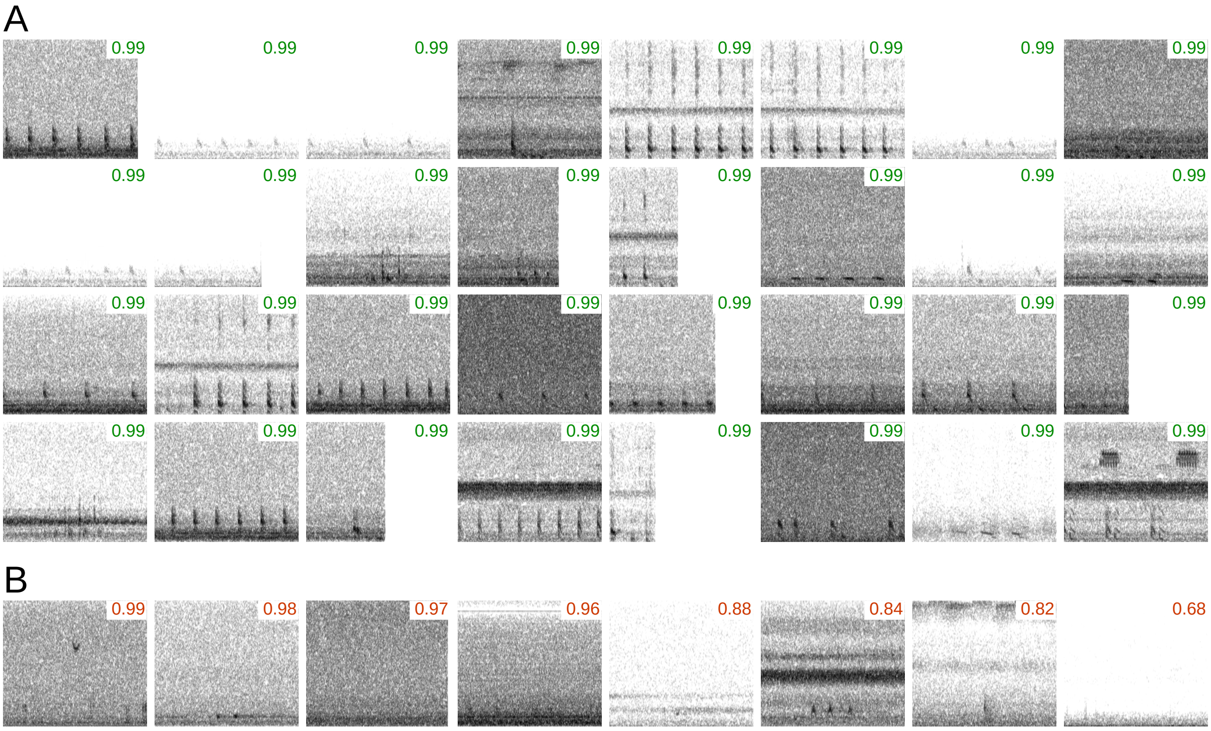


Figure S7: Classification examples for **Cuculiformes** drawn from the test dataset, shown as cropped spectrograms. Numbers in upper-right corners are calibrated probability values returned by the model. (A) Test examples scoring highest for this species. Any incorrect classifications are outlined in red. (B) Incorrectly-classified examples scoring highest for this species (i.e., the most confusing cases for the model). Spectrogram parameters: x-axis: 0-0.8 s; y-axis: 1-10 kHz; window type: Hanning; window length: 300 samples; hop size: 30 samples; dynamic range floor: -50 dB).

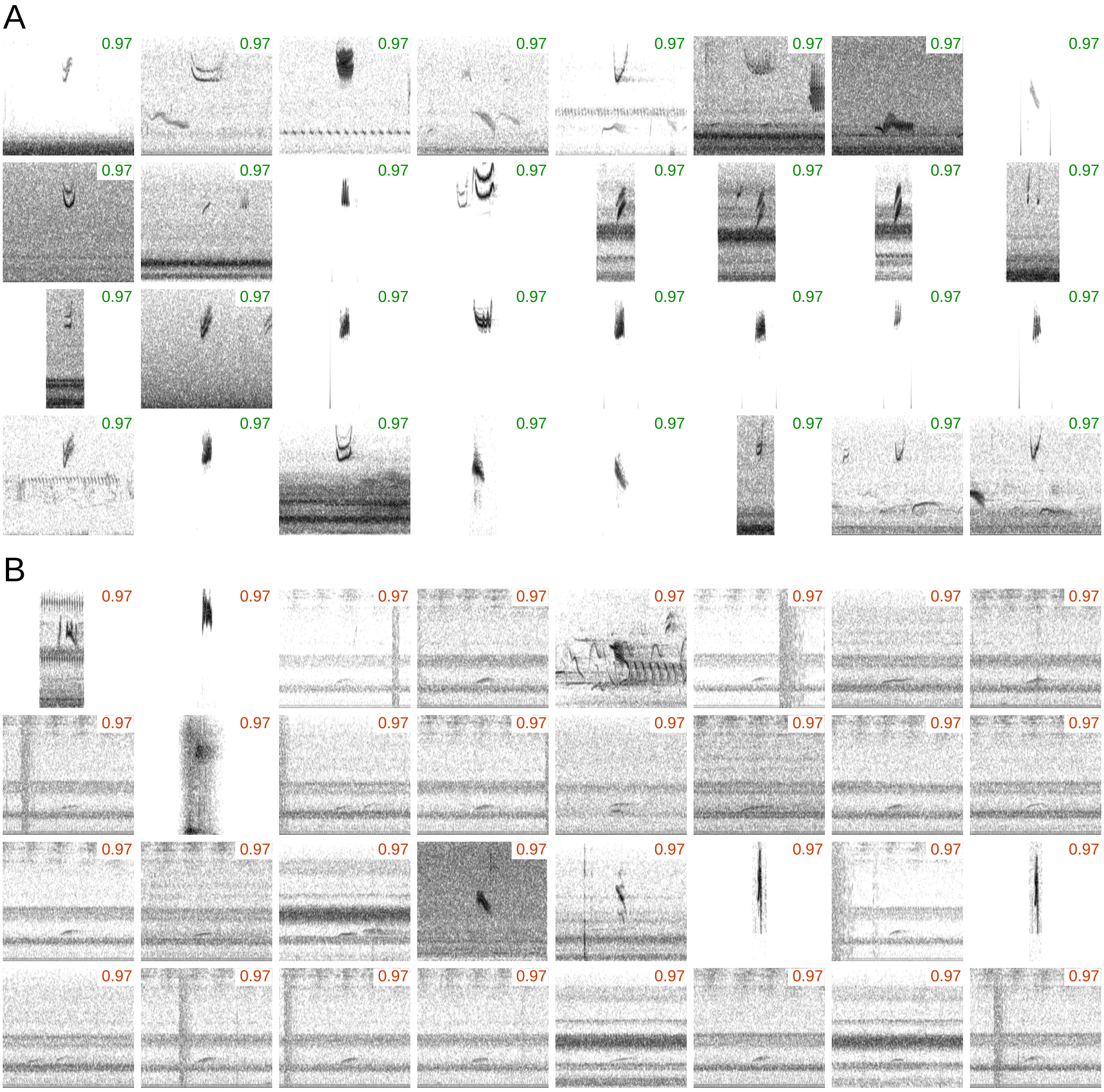


Figure S8: Classification examples for **Passeriformes** drawn from the test dataset, shown as cropped spectrograms. Numbers in upper-right corners are calibrated probability values returned by the model. (A) Test examples scoring highest for this species. Any incorrect classifications are outlined in red. (B) Incorrectly-classified examples scoring highest for this species (i.e., the most confusing cases for the model). Spectrogram parameters: x-axis: 0-0.8 s; y-axis: 1-10 kHz; window type: Hanning; window length: 300 samples; hop size: 30 samples; dynamic range floor: -50 dB).

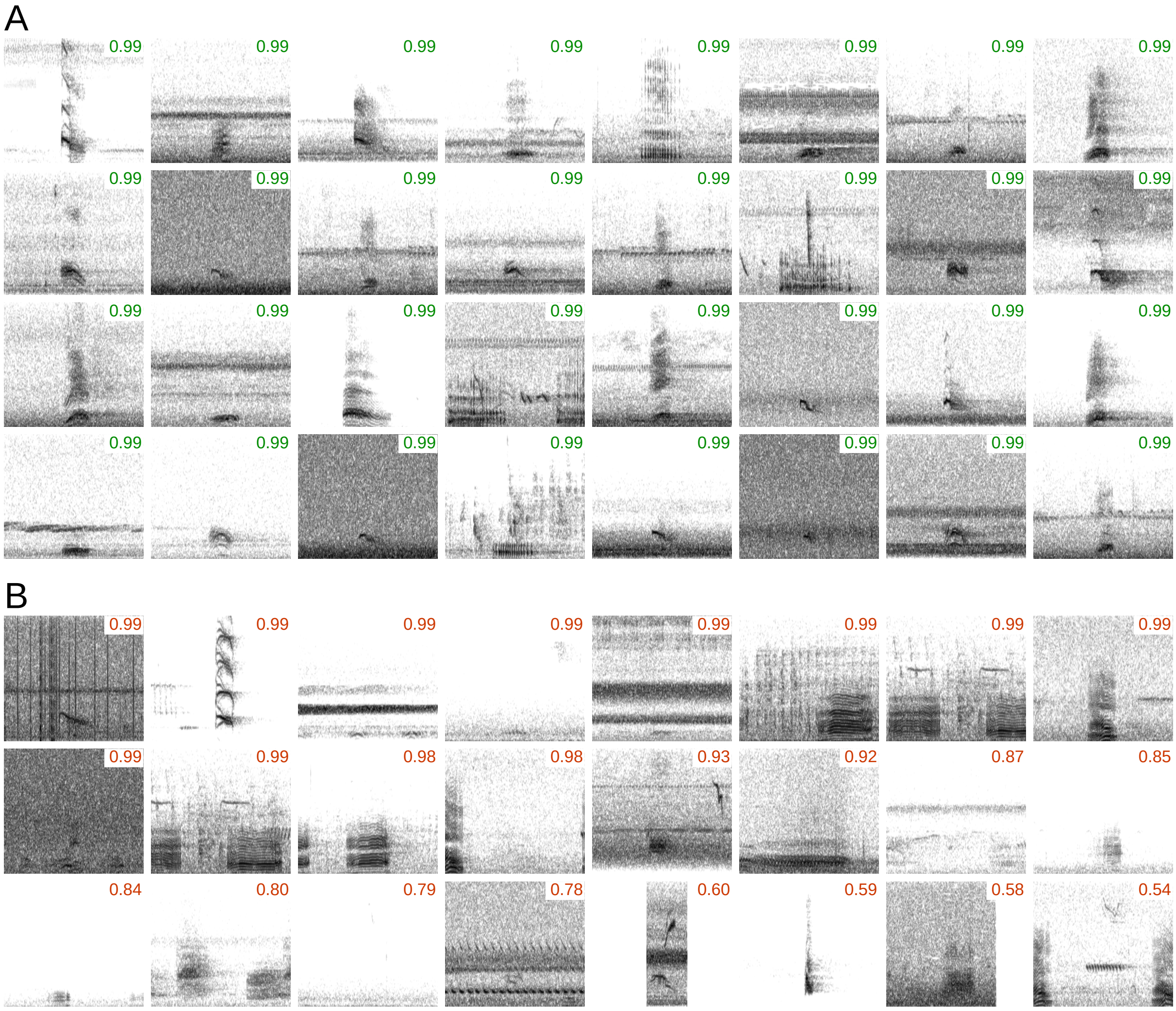


Figure S9: Classification examples for **Pelecaniformes** drawn from the test dataset, shown as cropped spectrograms. Numbers in upper-right corners are calibrated probability values returned by the model. (A) Test examples scoring highest for this species. Any incorrect classifications are outlined in red. (B) Incorrectly-classified examples scoring highest for this species (i.e., the most confusing cases for the model). Spectrogram parameters: x-axis: 0-0.8 s; y-axis: 1-10 kHz; window type: Hanning; window length: 300 samples; hop size: 30 samples; dynamic range floor: -50 dB).

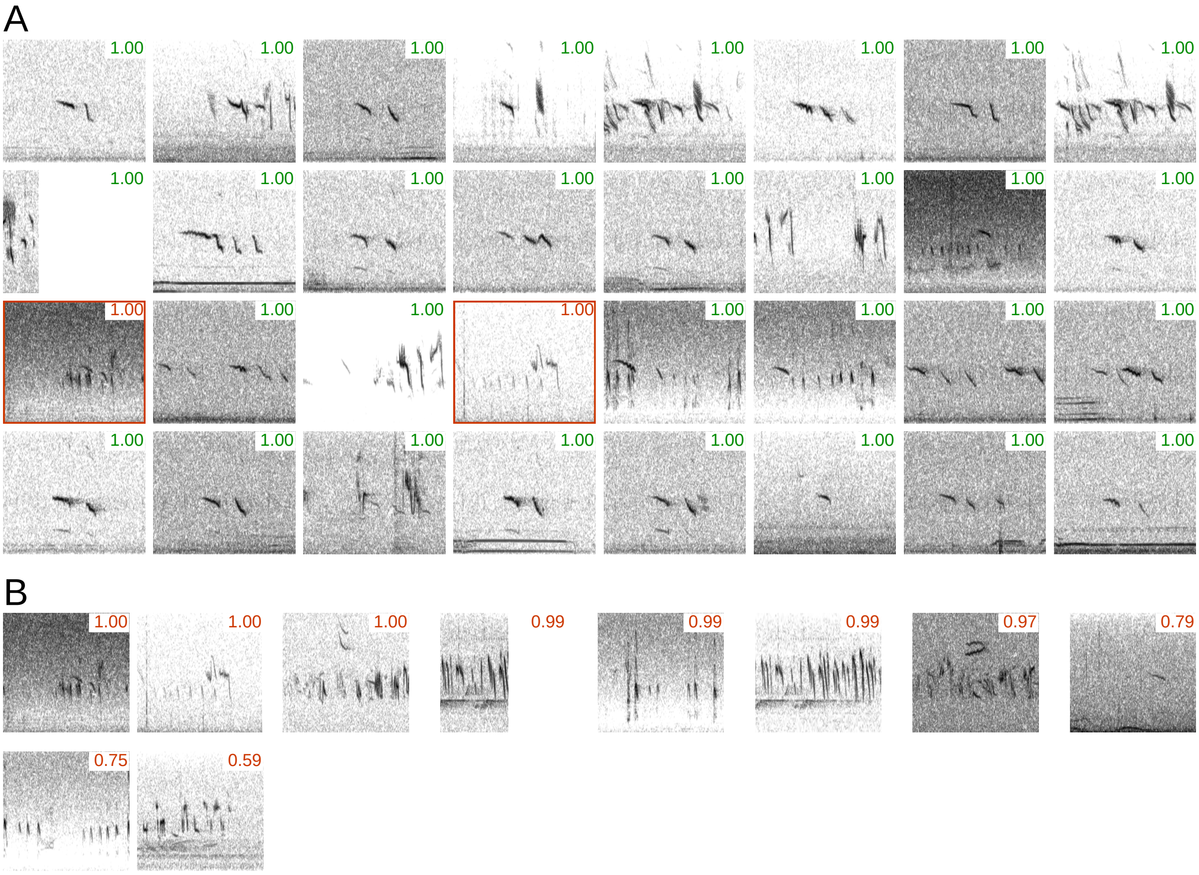


Figure S10: Classification examples for **Alaudidae** drawn from the test dataset, shown as cropped spectrograms. Numbers in upper-right corners are calibrated probability values returned by the model. (A) Test examples scoring highest for this species. Any incorrect classifications are outlined in red. (B) Incorrectly-classified examples scoring highest for this species (i.e., the most confusing cases for the model). Spectrogram parameters: x-axis: 0-0.8 s; y-axis: 1-10 kHz; window type: Hanning; window length: 300 samples; hop size: 30 samples; dynamic range floor: -50 dB).

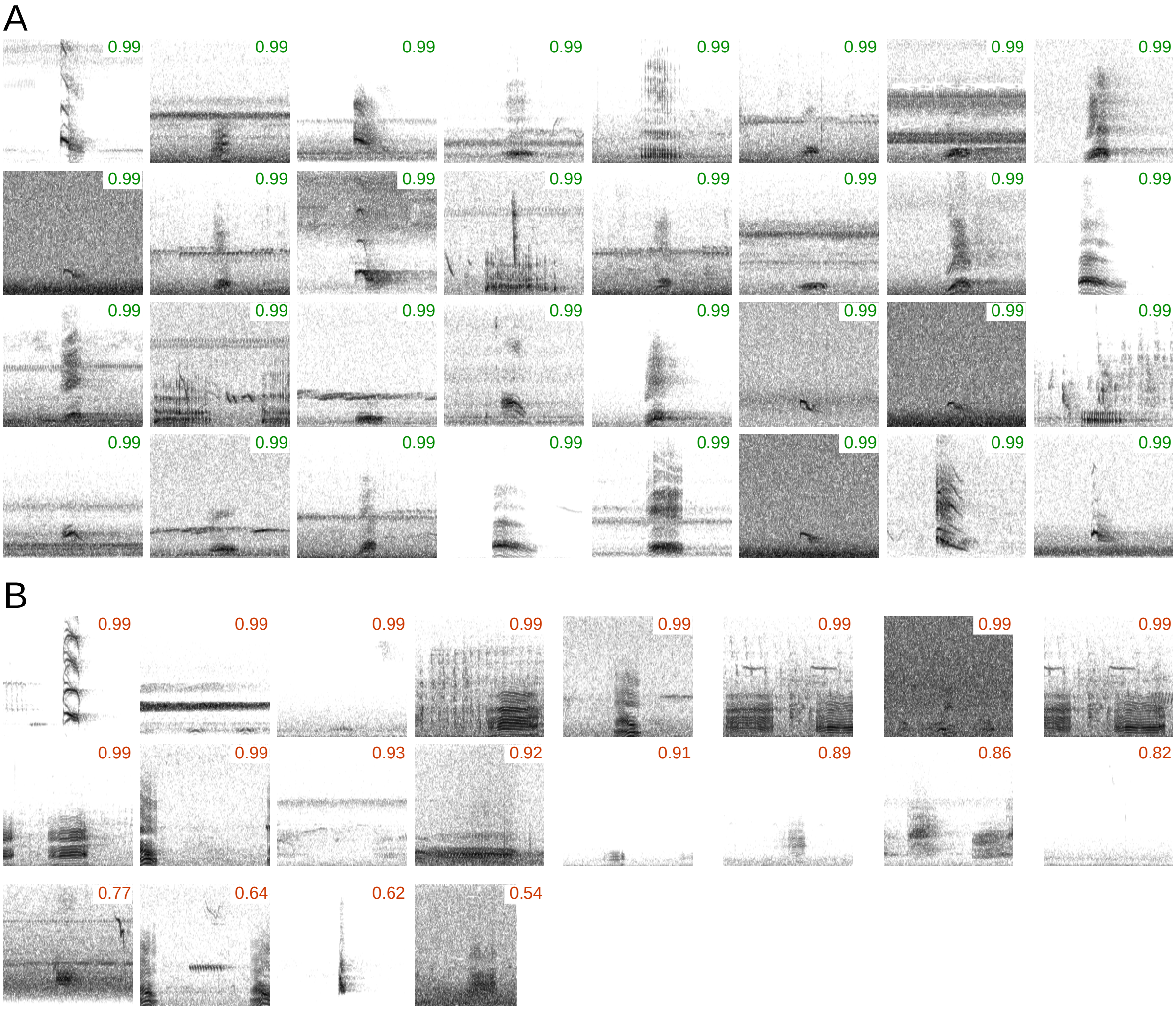


Figure S11: Classification examples for **Ardeidae** drawn from the test dataset, shown as cropped spectrograms. Numbers in upper-right corners are calibrated probability values returned by the model. (A) Test examples scoring highest for this species. Any incorrect classifications are outlined in red. (B) Incorrectly-classified examples scoring highest for this species (i.e., the most confusing cases for the model). Spectrogram parameters: x-axis: 0-0.8 s; y-axis: 1-10 kHz; window type: Hanning; window length: 300 samples; hop size: 30 samples; dynamic range floor: -50 dB).

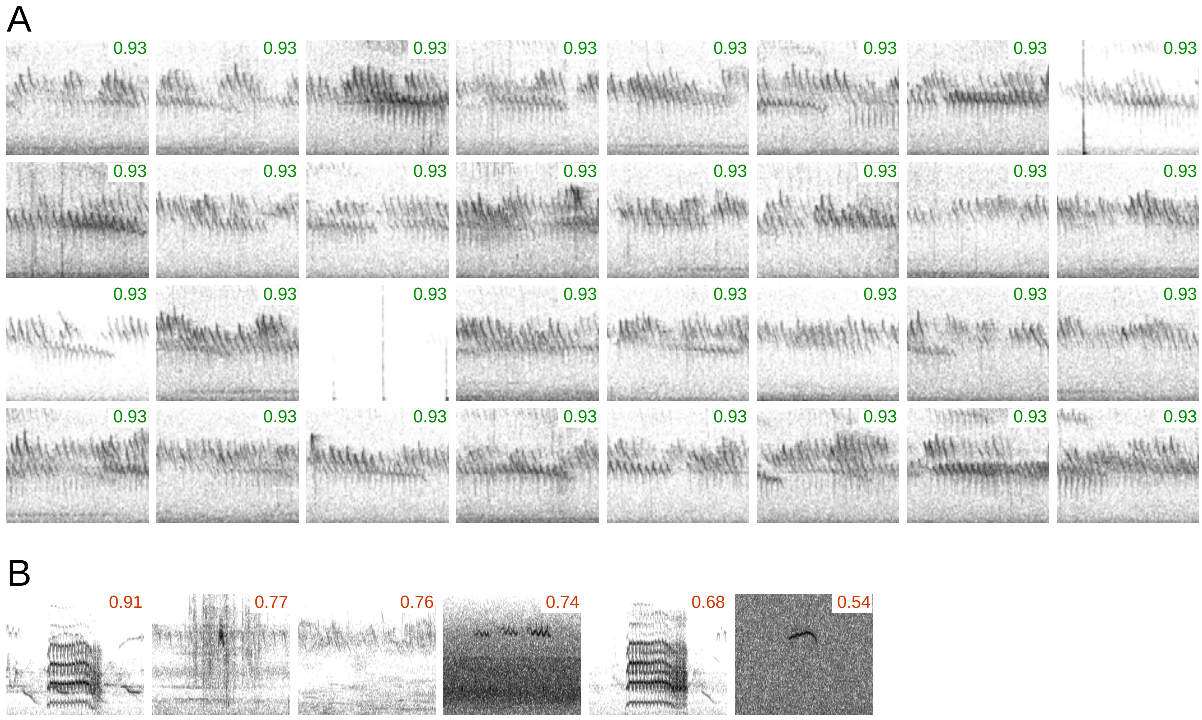


Figure S12: Classification examples for **Bombycillidae** drawn from the test dataset, shown as cropped spectrograms. Numbers in upper-right corners are calibrated probability values returned by the model. (A) Test examples scoring highest for this species. Any incorrect classifications are outlined in red. (B) Incorrectly-classified examples scoring highest for this species (i.e., the most confusing cases for the model). Spectrogram parameters: x-axis: 0-0.8 s; y-axis: 1-10 kHz; window type: Hanning; window length: 300 samples; hop size: 30 samples; dynamic range floor: -50 dB).

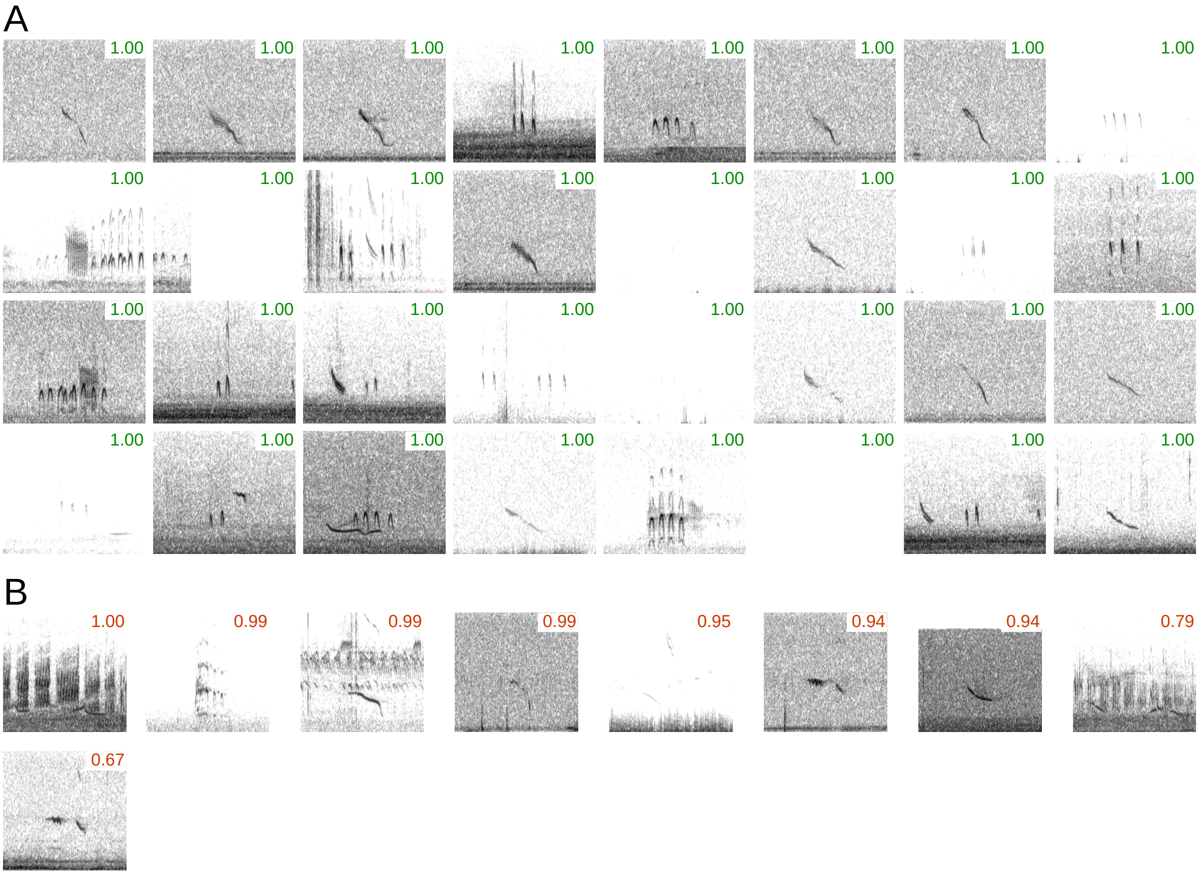


Figure S13: Classification examples for **Calcariidae** drawn from the test dataset, shown as cropped spectrograms. Numbers in upper-right corners are calibrated probability values returned by the model. (A) Test examples scoring highest for this species. Any incorrect classifications are outlined in red. (B) Incorrectly-classified examples scoring highest for this species (i.e., the most confusing cases for the model). Spectrogram parameters: x-axis: 0-0.8 s; y-axis: 1-10 kHz; window type: Hanning; window length: 300 samples; hop size: 30 samples; dynamic range floor: -50 dB).

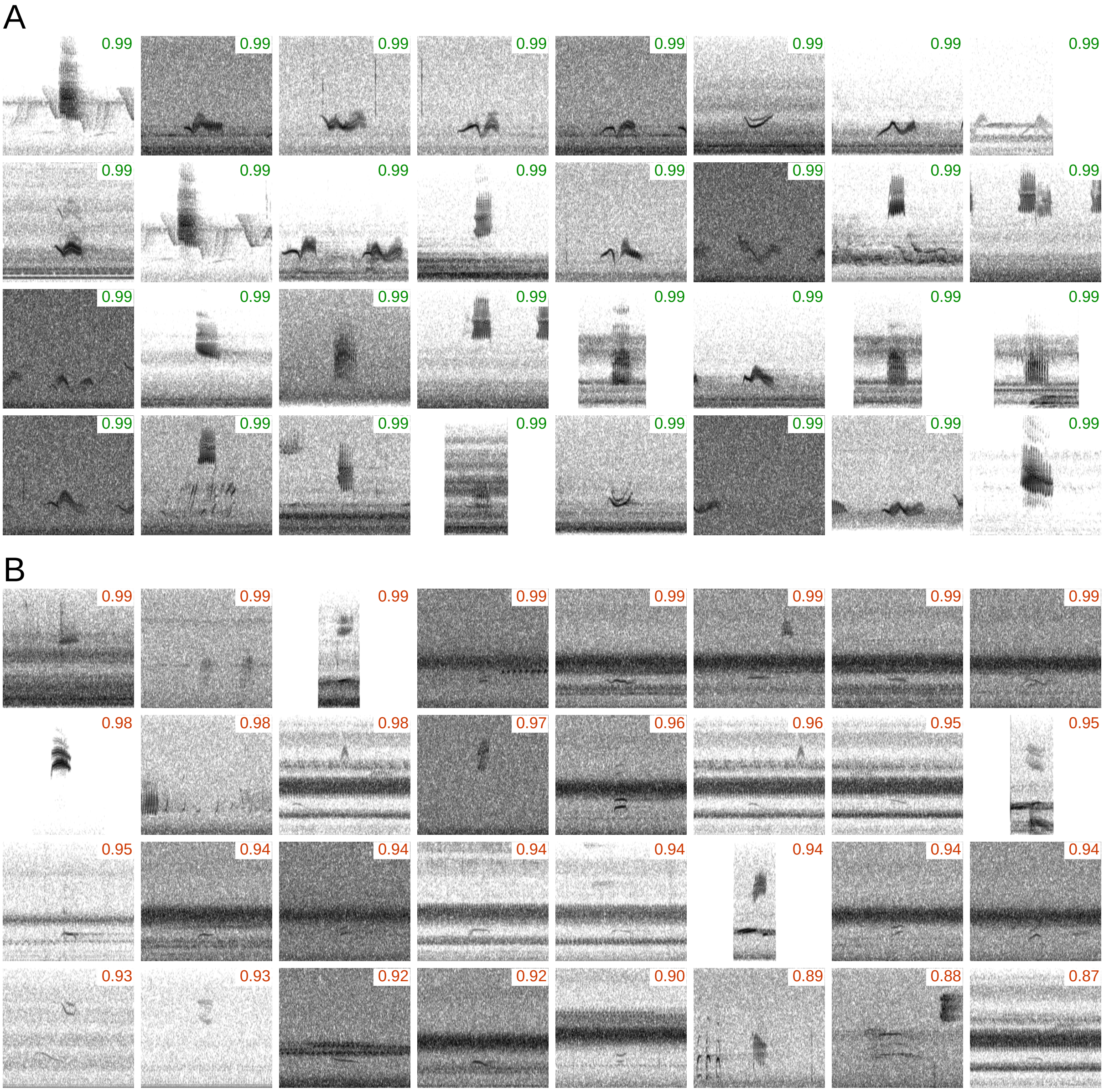


Figure S14: Classification examples for **Cardinalidae** drawn from the test dataset, shown as cropped spectrograms. Numbers in upper-right corners are calibrated probability values returned by the model. (A) Test examples scoring highest for this species. Any incorrect classifications are outlined in red. (B) Incorrectly-classified examples scoring highest for this species (i.e., the most confusing cases for the model). Spectrogram parameters: x-axis: 0-0.8 s; y-axis: 1-10 kHz; window type: Hanning; window length: 300 samples; hop size: 30 samples; dynamic range floor: -50 dB).

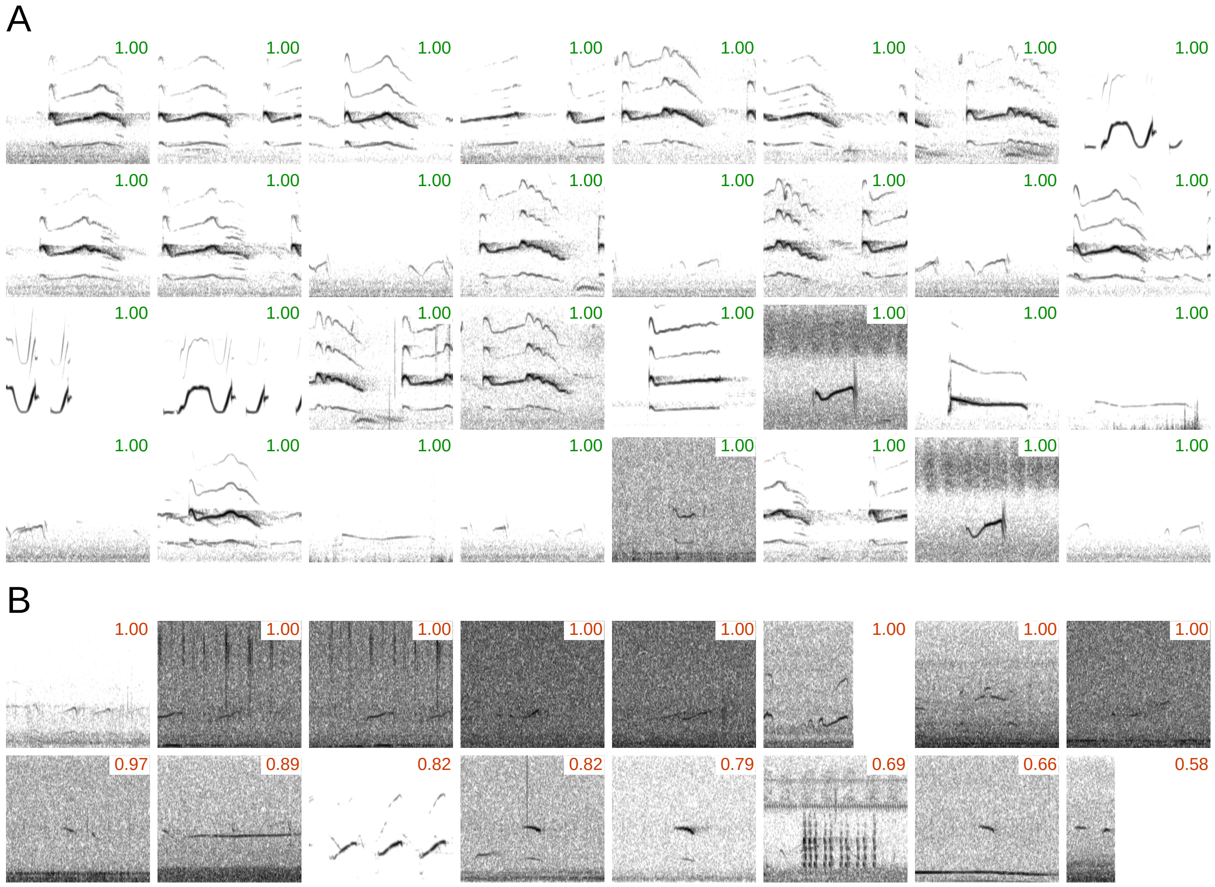


Figure S15: Classification examples for **Charadriidae** drawn from the test dataset, shown as cropped spectrograms. Numbers in upper-right corners are calibrated probability values returned by the model. (A) Test examples scoring highest for this species. Any incorrect classifications are outlined in red. (B) Incorrectly-classified examples scoring highest for this species (i.e., the most confusing cases for the model). Spectrogram parameters: x-axis: 0-0.8 s; y-axis: 1-10 kHz; window type: Hanning; window length: 300 samples; hop size: 30 samples; dynamic range floor: -50 dB).

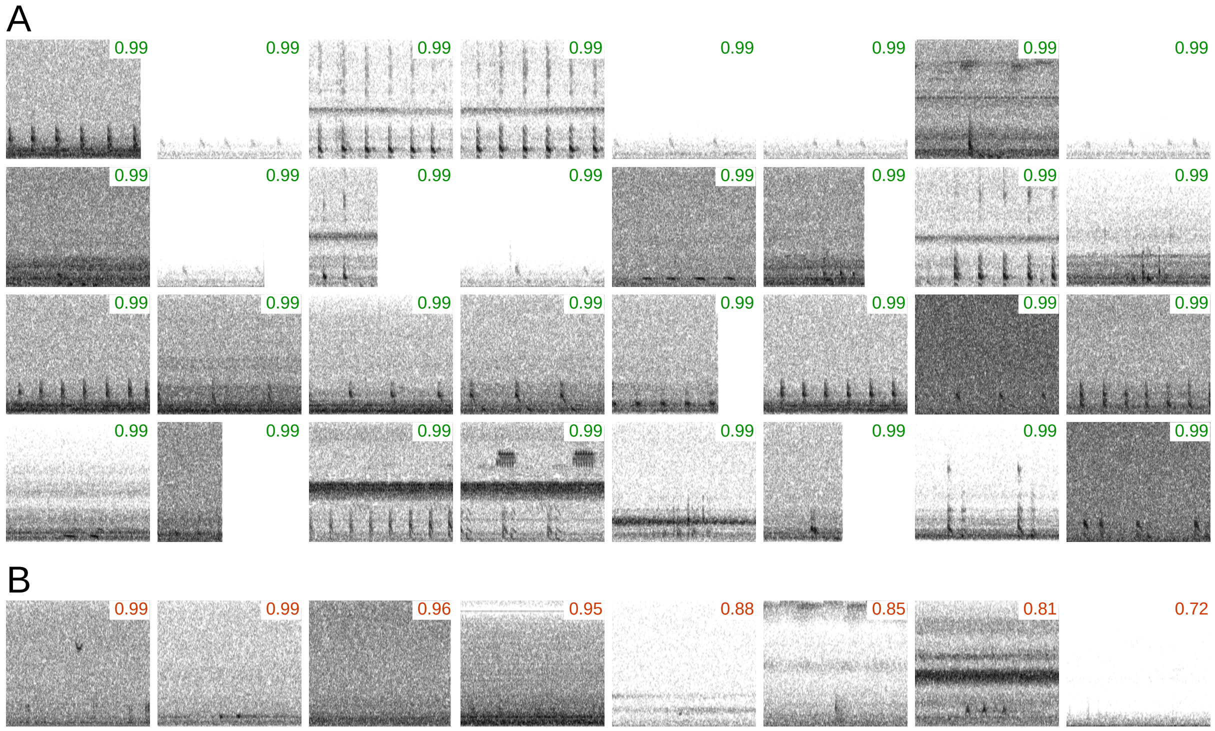


Figure S16: Classification examples for **Cuculidae** drawn from the test dataset, shown as cropped spectrograms. Numbers in upper-right corners are calibrated probability values returned by the model. (A) Test examples scoring highest for this species. Any incorrect classifications are outlined in red. (B) Incorrectly-classified examples scoring highest for this species (i.e., the most confusing cases for the model). Spectrogram parameters: x-axis: 0-0.8 s; y-axis: 1-10 kHz; window type: Hanning; window length: 300 samples; hop size: 30 samples; dynamic range floor: -50 dB).

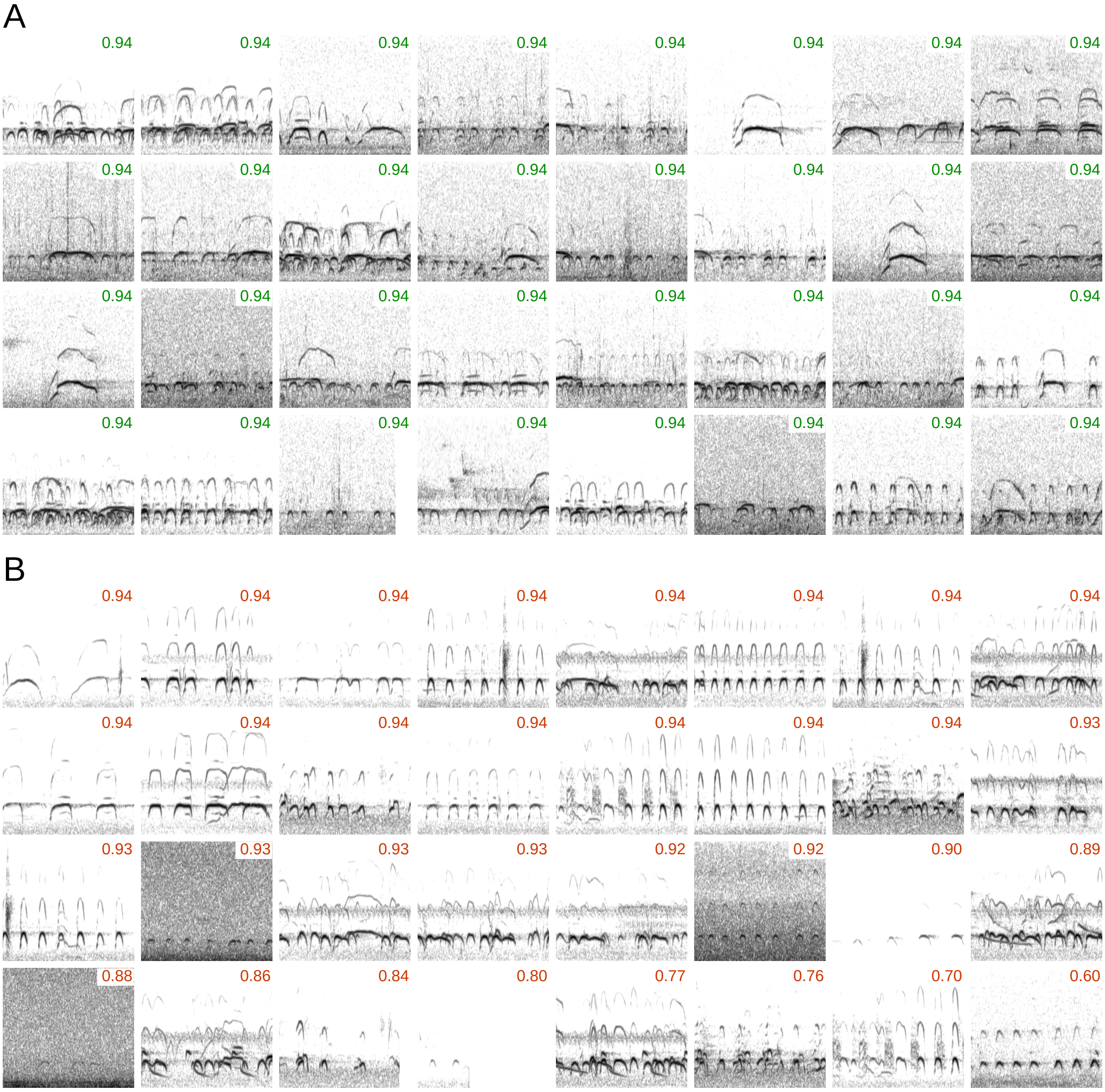


Figure S17: Classification examples for **Haematopodidae** drawn from the test dataset, shown as cropped spectrograms. Numbers in upper-right corners are calibrated probability values returned by the model. (A) Test examples scoring highest for this species. Any incorrect classifications are outlined in red. (B) Incorrectly-classified examples scoring highest for this species (i.e., the most confusing cases for the model). Spectrogram parameters: x-axis: 0-0.8 s; y-axis: 1-10 kHz; window type: Hanning; window length: 300 samples; hop size: 30 samples; dynamic range floor: -50 dB).

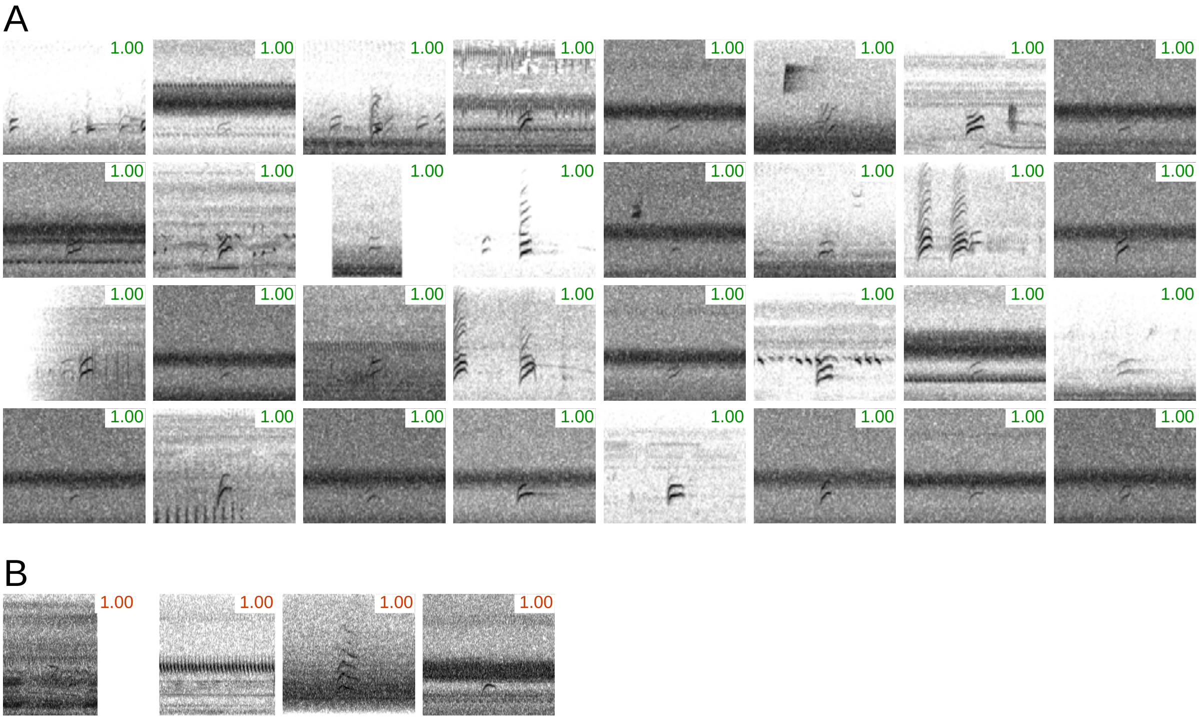


Figure S18: Classification examples for **Icteridae** drawn from the test dataset, shown as cropped spectrograms. Numbers in upper-right corners are calibrated probability values returned by the model. (A) Test examples scoring highest for this species. Any incorrect classifications are outlined in red. (B) Incorrectly-classified examples scoring highest for this species (i.e., the most confusing cases for the model). Spectrogram parameters: x-axis: 0-0.8 s; y-axis: 1-10 kHz; window type: Hanning; window length: 300 samples; hop size: 30 samples; dynamic range floor: -50 dB).

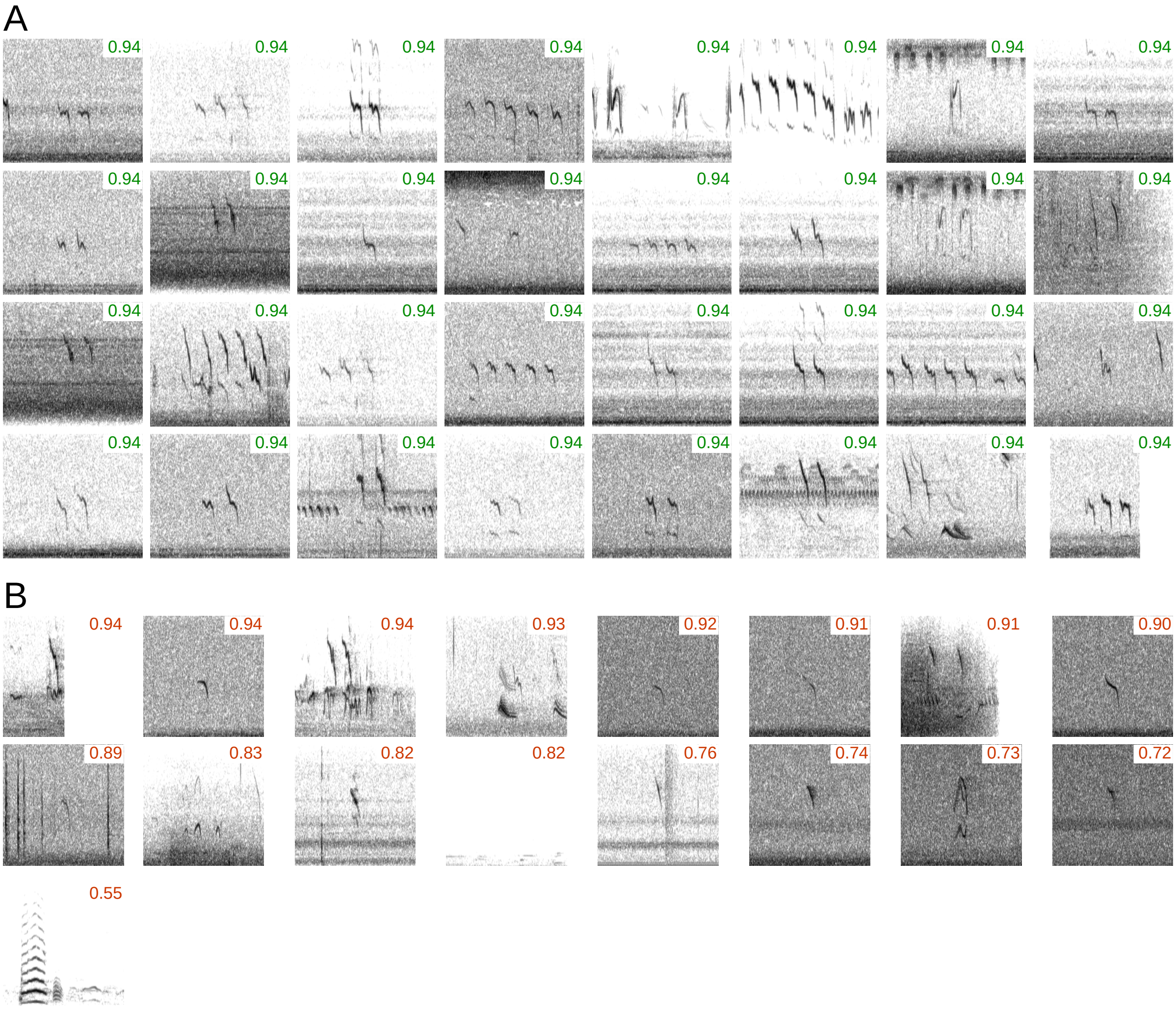


Figure S20: Classification examples for **Motacillidae** drawn from the test dataset, shown as cropped spectrograms. Numbers in upper-right corners are calibrated probability values returned by the model. (A) Test examples scoring highest for this species. Any incorrect classifications are outlined in red. (B) Incorrectly-classified examples scoring highest for this species (i.e., the most confusing cases for the model). Spectrogram parameters: x-axis: 0-0.8 s; y-axis: 1-10 kHz; window type: Hanning; window length: 300 samples; hop size: 30 samples; dynamic range floor: -50 dB).

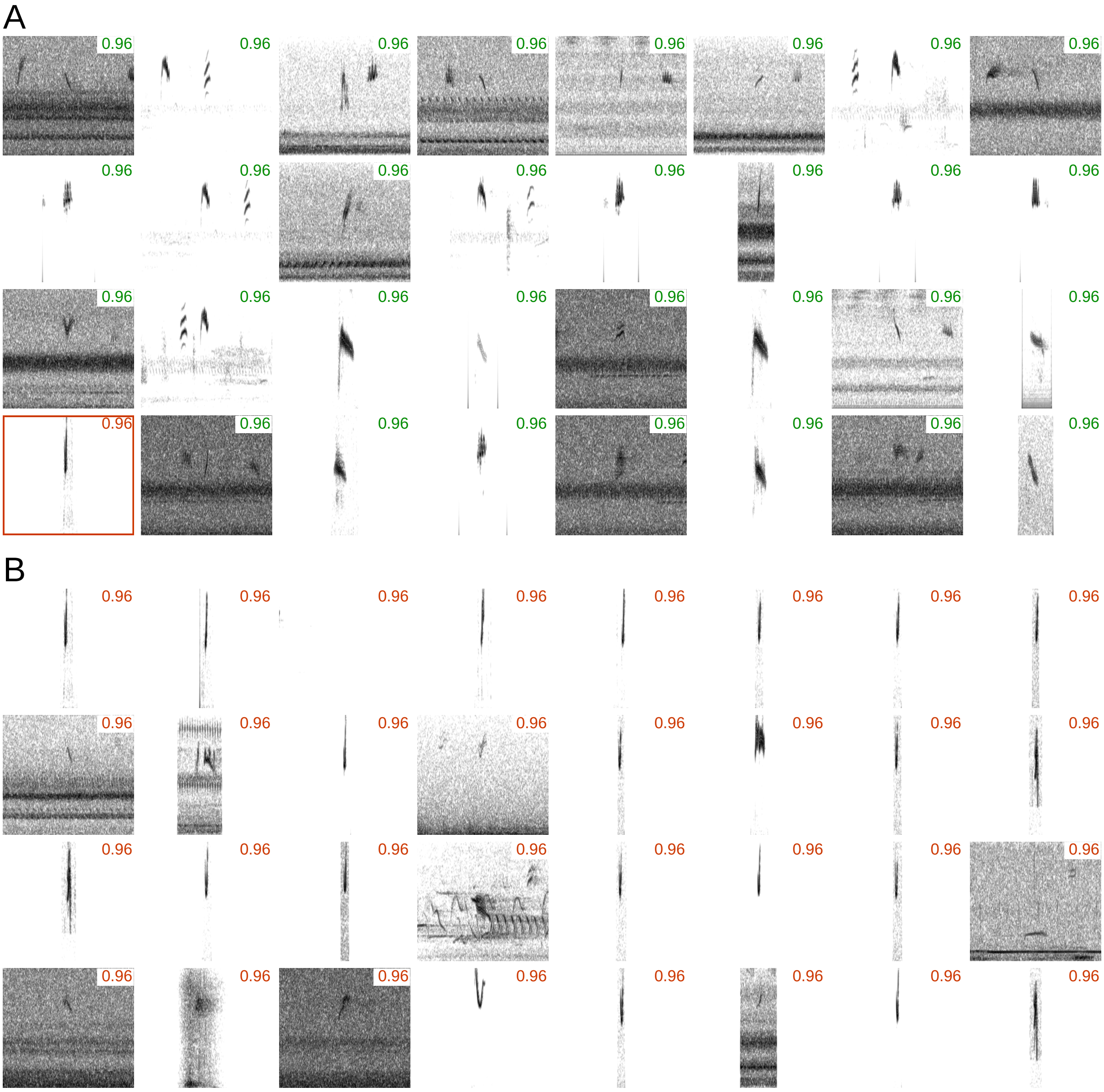


Figure S21: Classification examples for **Parulidae** drawn from the test dataset, shown as cropped spectrograms. Numbers in upper-right corners are calibrated probability values returned by the model. (A) Test examples scoring highest for this species. Any incorrect classifications are outlined in red. (B) Incorrectly-classified examples scoring highest for this species (i.e., the most confusing cases for the model). Spectrogram parameters: x-axis: 0-0.8 s; y-axis: 1-10 kHz; window type: Hanning; window length: 300 samples; hop size: 30 samples; dynamic range floor: -50 dB).

A

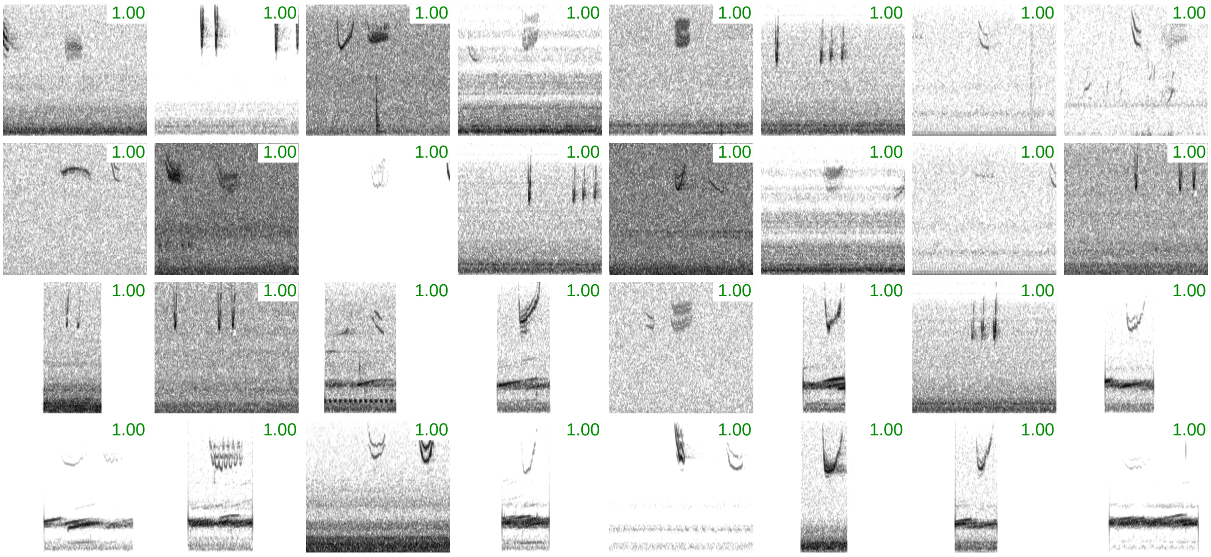

B

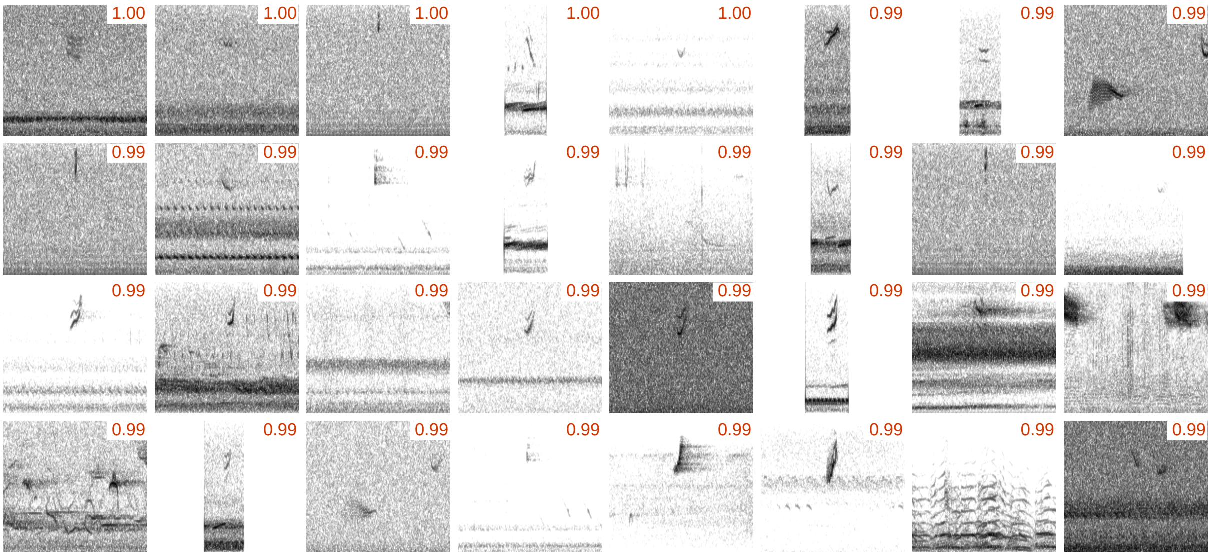

Figure S22: Classification examples for **Passerellidae** drawn from the test dataset, shown as cropped spectrograms. Numbers in upper-right corners are calibrated probability values returned by the model. (A) Test examples scoring highest for this species. Any incorrect classifications are outlined in red. (B) Incorrectly-classified examples scoring highest for this species (i.e., the most confusing cases for the model). Spectrogram parameters: x-axis: 0-0.8 s; y-axis: 1-10 kHz; window type: Hanning; window length: 300 samples; hop size: 30 samples; dynamic range floor: -50 dB).

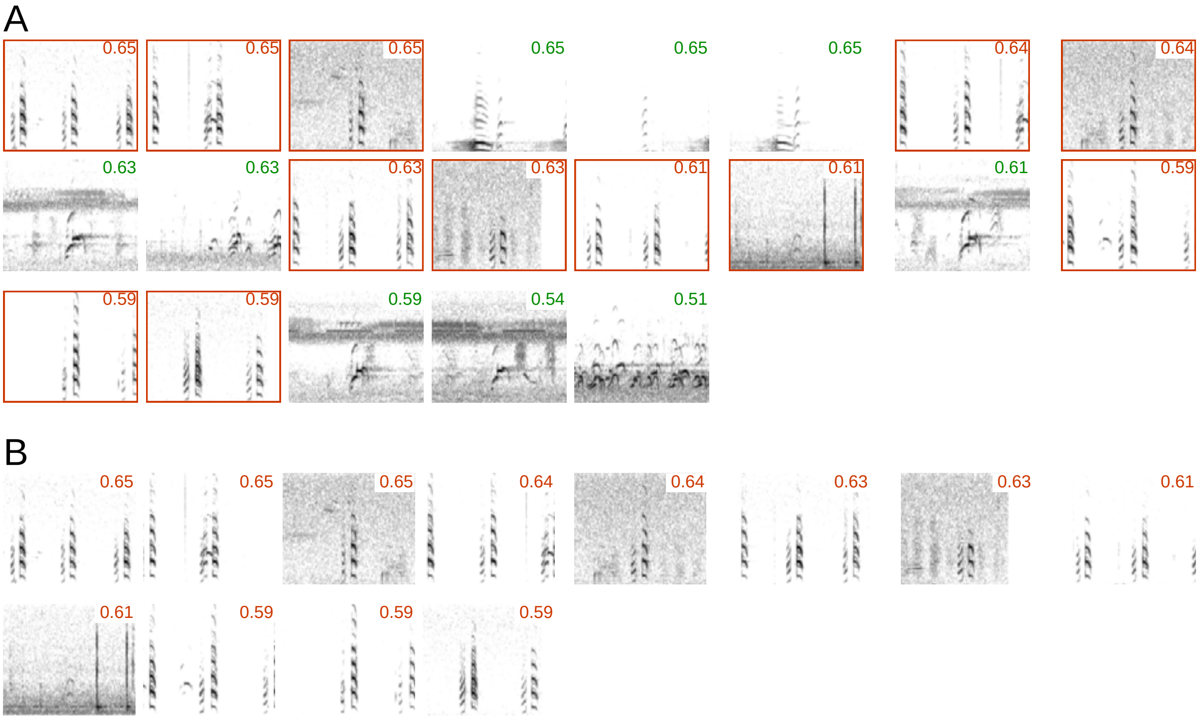


Figure S23: Classification examples for **Recurvirostridae** drawn from the test dataset, shown as cropped spectrograms. Numbers in upper-right corners are calibrated probability values returned by the model. (A) Test examples scoring highest for this species. Any incorrect classifications are outlined in red. (B) Incorrectly-classified examples scoring highest for this species (i.e., the most confusing cases for the model). Spectrogram parameters: x-axis: 0-0.8 s; y-axis: 1-10 kHz; window type: Hanning; window length: 300 samples; hop size: 30 samples; dynamic range floor: -50 dB).

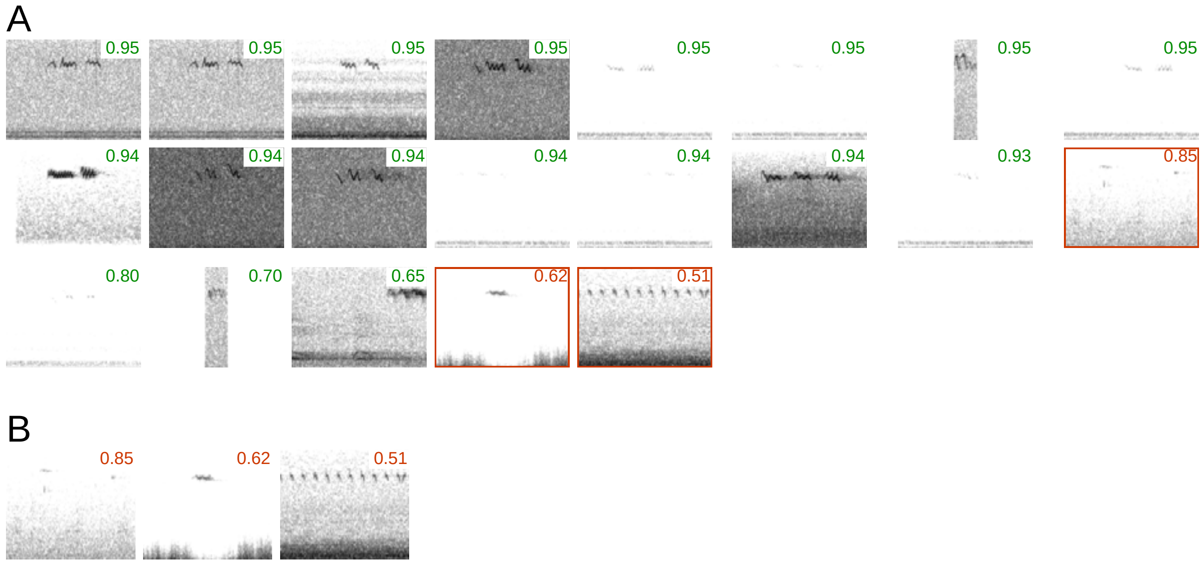


Figure S24: Classification examples for **Regulidae** drawn from the test dataset, shown as cropped spectrograms. Numbers in upper-right corners are calibrated probability values returned by the model. (A) Test examples scoring highest for this species. Any incorrect classifications are outlined in red. (B) Incorrectly-classified examples scoring highest for this species (i.e., the most confusing cases for the model). Spectrogram parameters: x-axis: 0-0.8 s; y-axis: 1-10 kHz; window type: Hanning; window length: 300 samples; hop size: 30 samples; dynamic range floor: -50 dB).

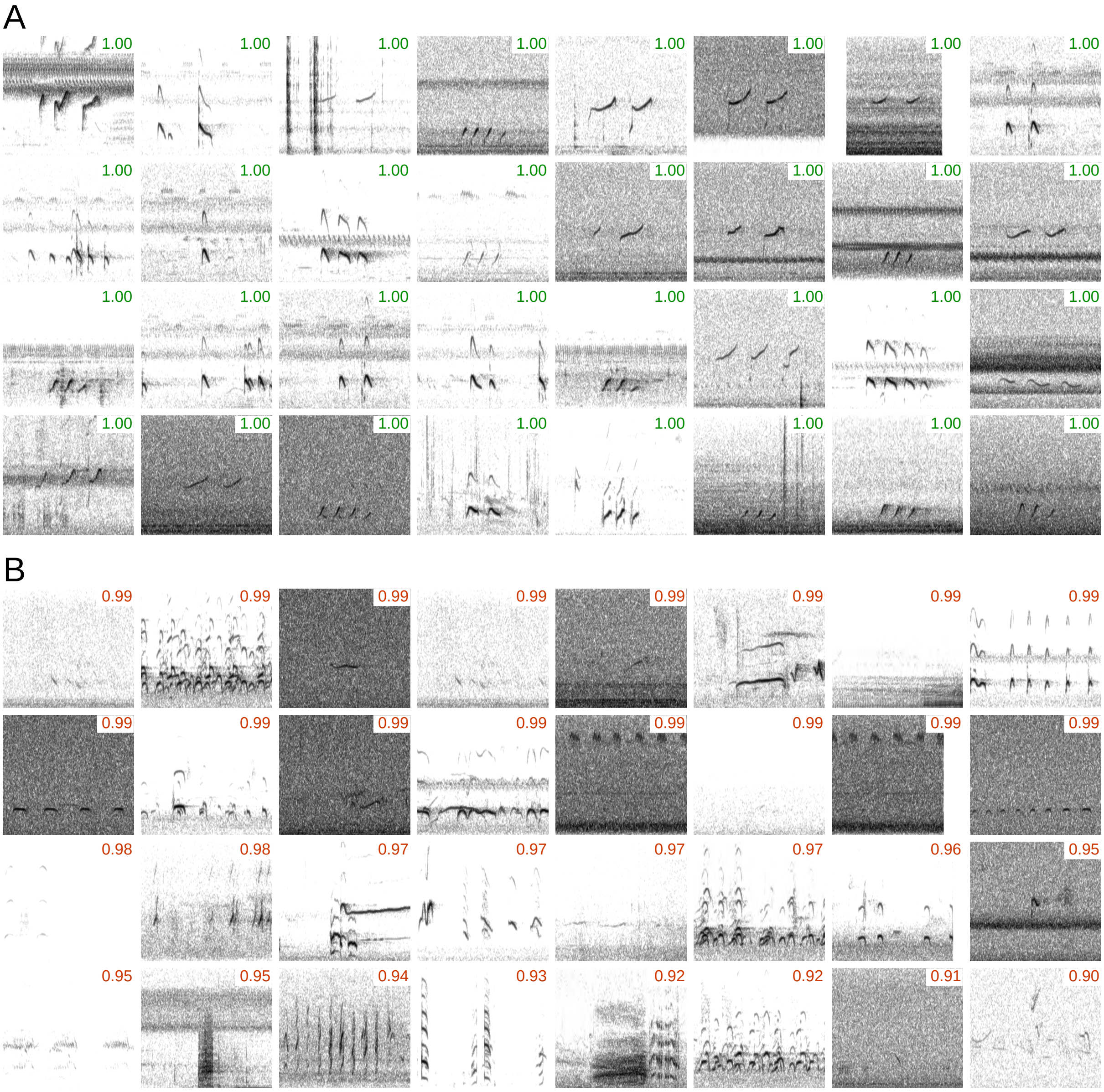


Figure S25: Classification examples for **Scolopacidae** drawn from the test dataset, shown as cropped spectrograms. Numbers in upper-right corners are calibrated probability values returned by the model. (A) Test examples scoring highest for this species. Any incorrect classifications are outlined in red. (B) Incorrectly-classified examples scoring highest for this species (i.e., the most confusing cases for the model). Spectrogram parameters: x-axis: 0-0.8 s; y-axis: 1-10 kHz; window type: Hanning; window length: 300 samples; hop size: 30 samples; dynamic range floor: -50 dB).

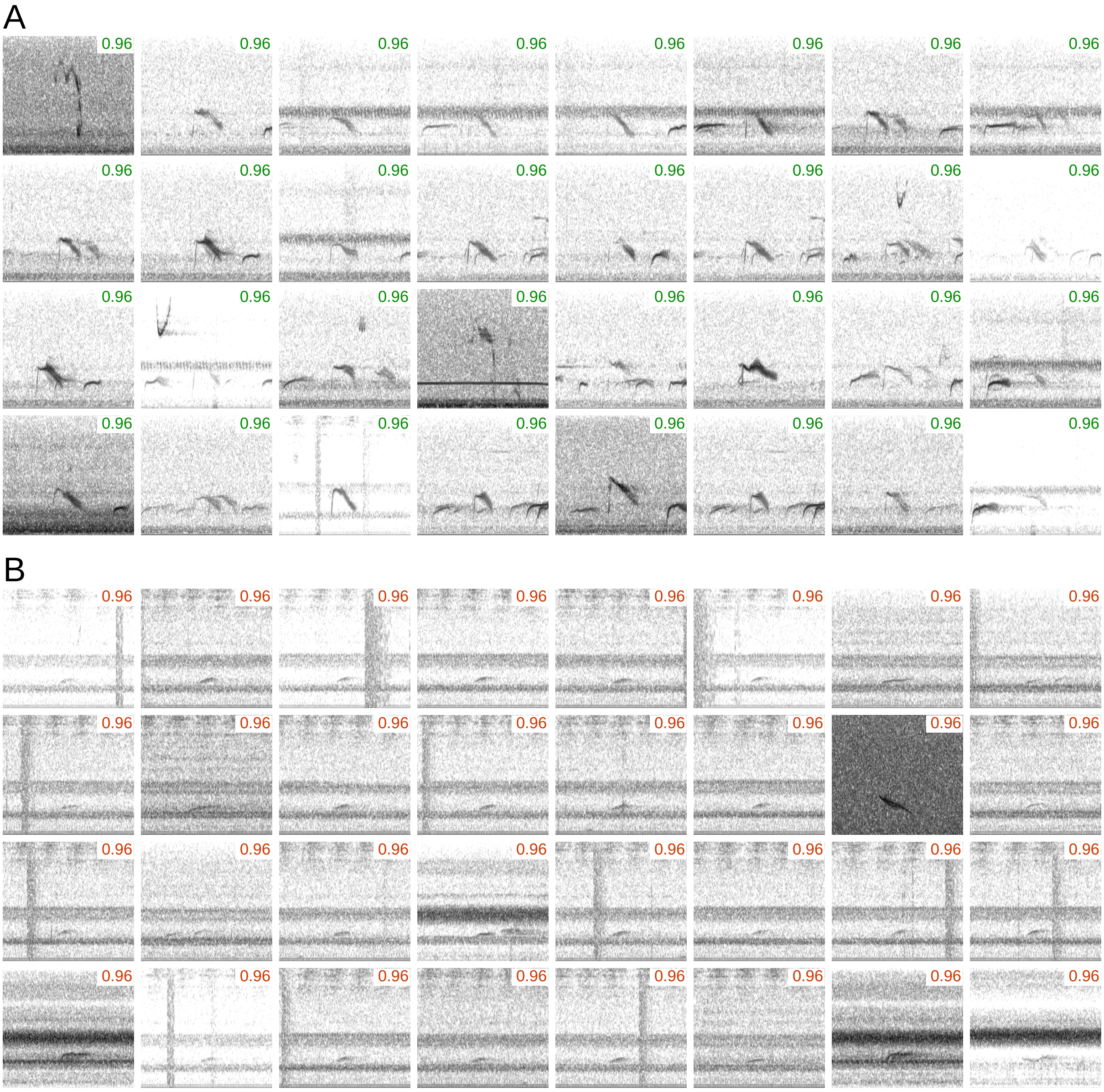


Figure S27: Classification examples for **Turdidae** drawn from the test dataset, shown as cropped spectrograms. Numbers in upper-right corners are calibrated probability values returned by the model. (A) Test examples scoring highest for this species. Any incorrect classifications are outlined in red. (B) Incorrectly-classified examples scoring highest for this species (i.e., the most confusing cases for the model). Spectrogram parameters: x-axis: 0-0.8 s; y-axis: 1-10 kHz; window type: Hanning; window length: 300 samples; hop size: 30 samples; dynamic range floor: -50 dB).

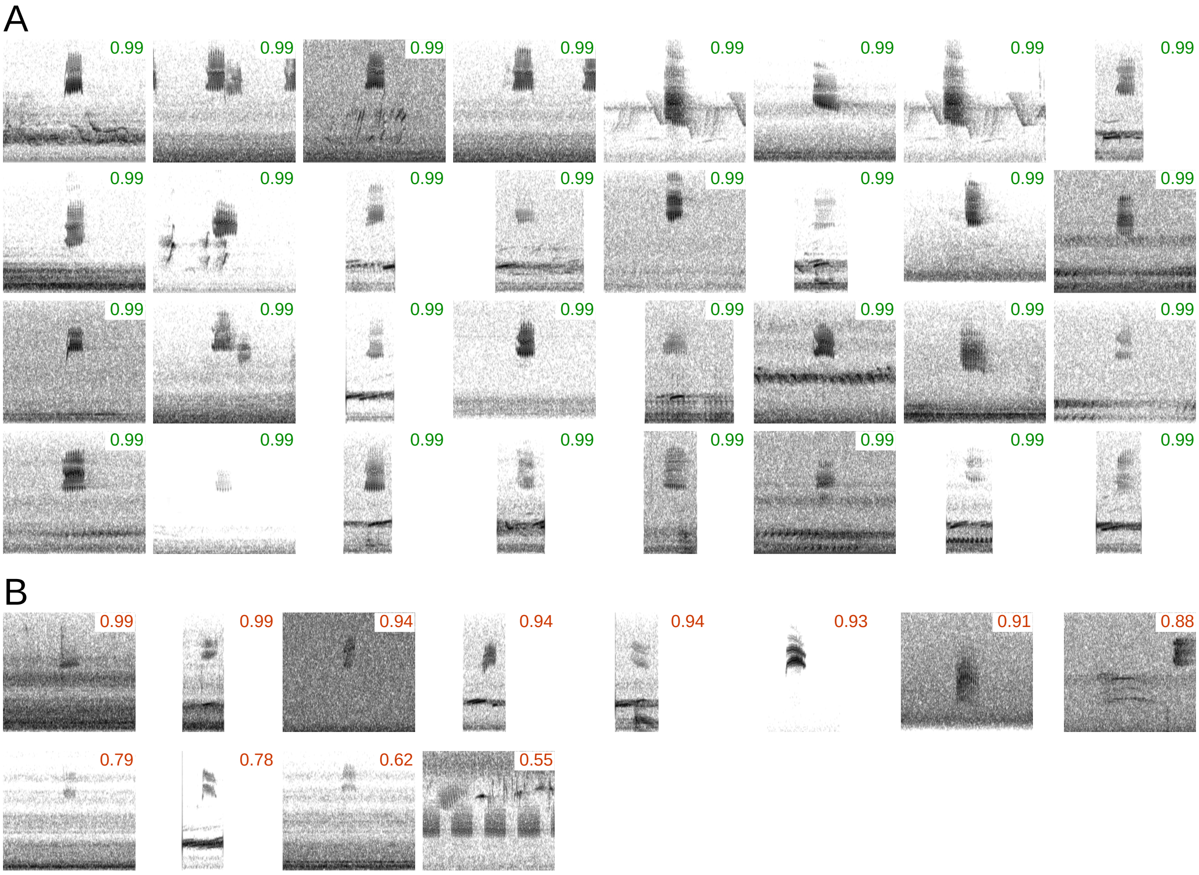


Figure S28: Classification examples for **BUNT** drawn from the test dataset, shown as cropped spectrograms. Numbers in upper-right corners are calibrated probability values returned by the model. (A) Test examples scoring highest for this species. Any incorrect classifications are outlined in red. (B) Incorrectly-classified examples scoring highest for this species (i.e., the most confusing cases for the model). Spectrogram parameters: x-axis: 0-0.8 s; y-axis: 1-10 kHz; window type: Hanning; window length: 300 samples; hop size: 30 samples; dynamic range floor: -50 dB).

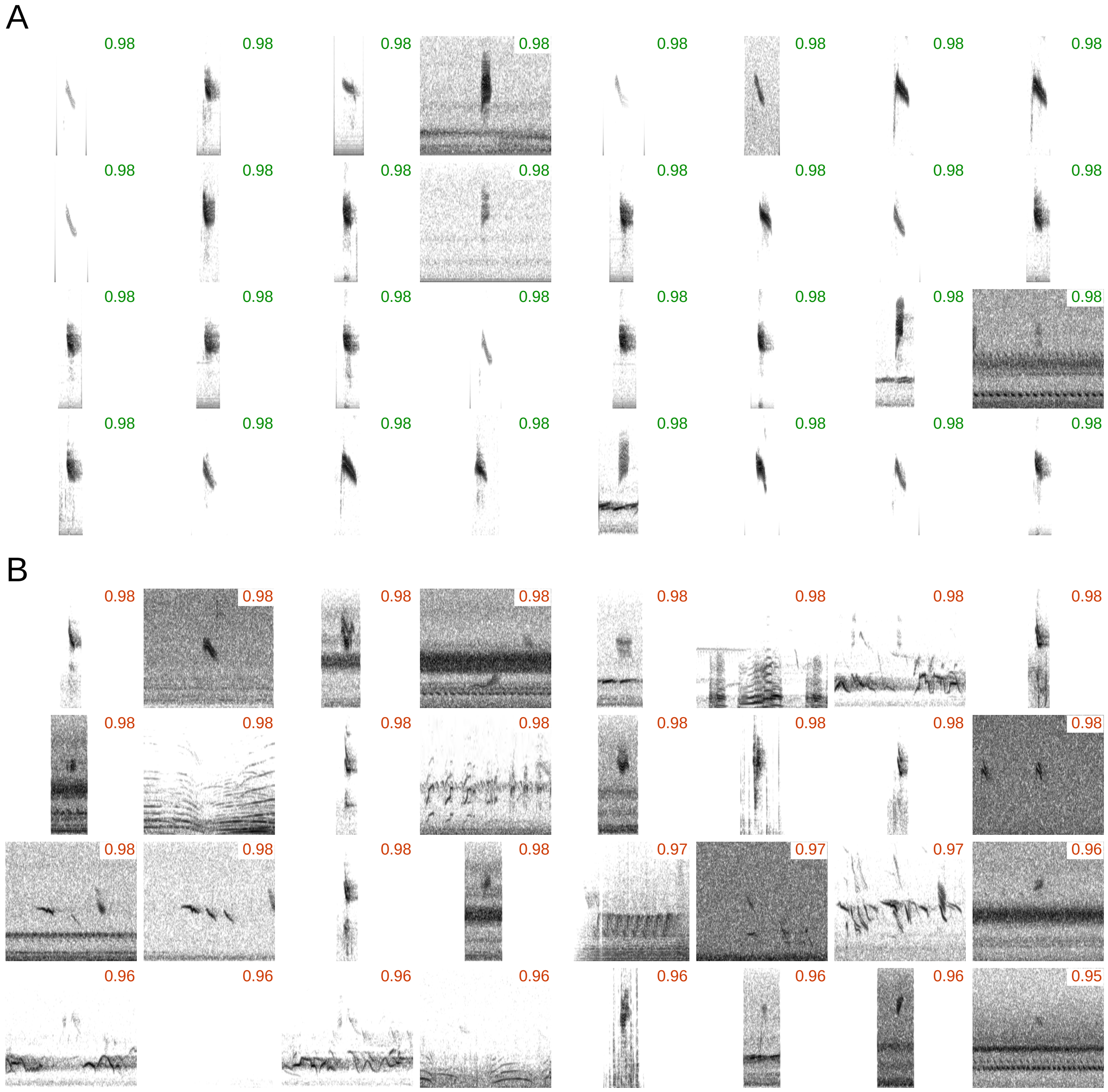


Figure S29: Classification examples for **BZWA** drawn from the test dataset, shown as cropped spectrograms. Numbers in upper-right corners are calibrated probability values returned by the model. (A) Test examples scoring highest for this species. Any incorrect classifications are outlined in red. (B) Incorrectly-classified examples scoring highest for this species (i.e., the most confusing cases for the model). Spectrogram parameters: x-axis: 0-0.8 s; y-axis: 1-10 kHz; window type: Hanning; window length: 300 samples; hop size: 30 samples; dynamic range floor: -50 dB).

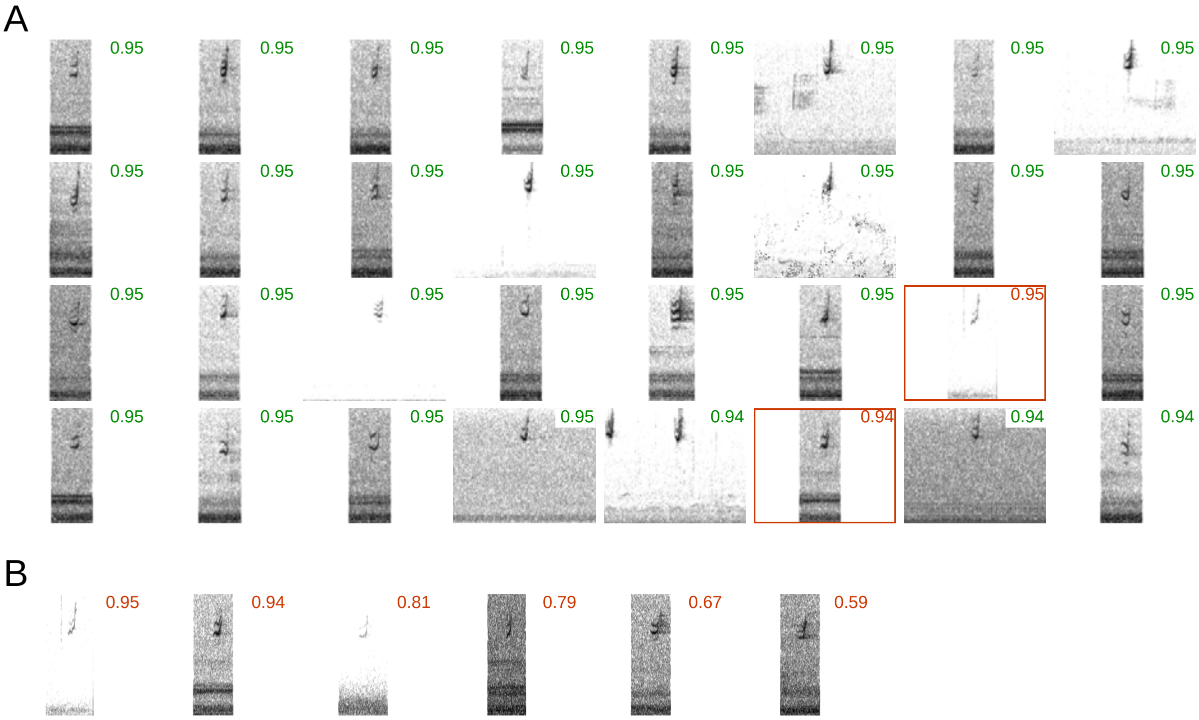


Figure S30: Classification examples for **CCBRS** drawn from the test dataset, shown as cropped spectrograms. Numbers in upper-right corners are calibrated probability values returned by the model. (A) Test examples scoring highest for this species. Any incorrect classifications are outlined in red. (B) Incorrectly-classified examples scoring highest for this species (i.e., the most confusing cases for the model). Spectrogram parameters: x-axis: 0-0.8 s; y-axis: 1-10 kHz; window type: Hanning; window length: 300 samples; hop size: 30 samples; dynamic range floor: -50 dB).

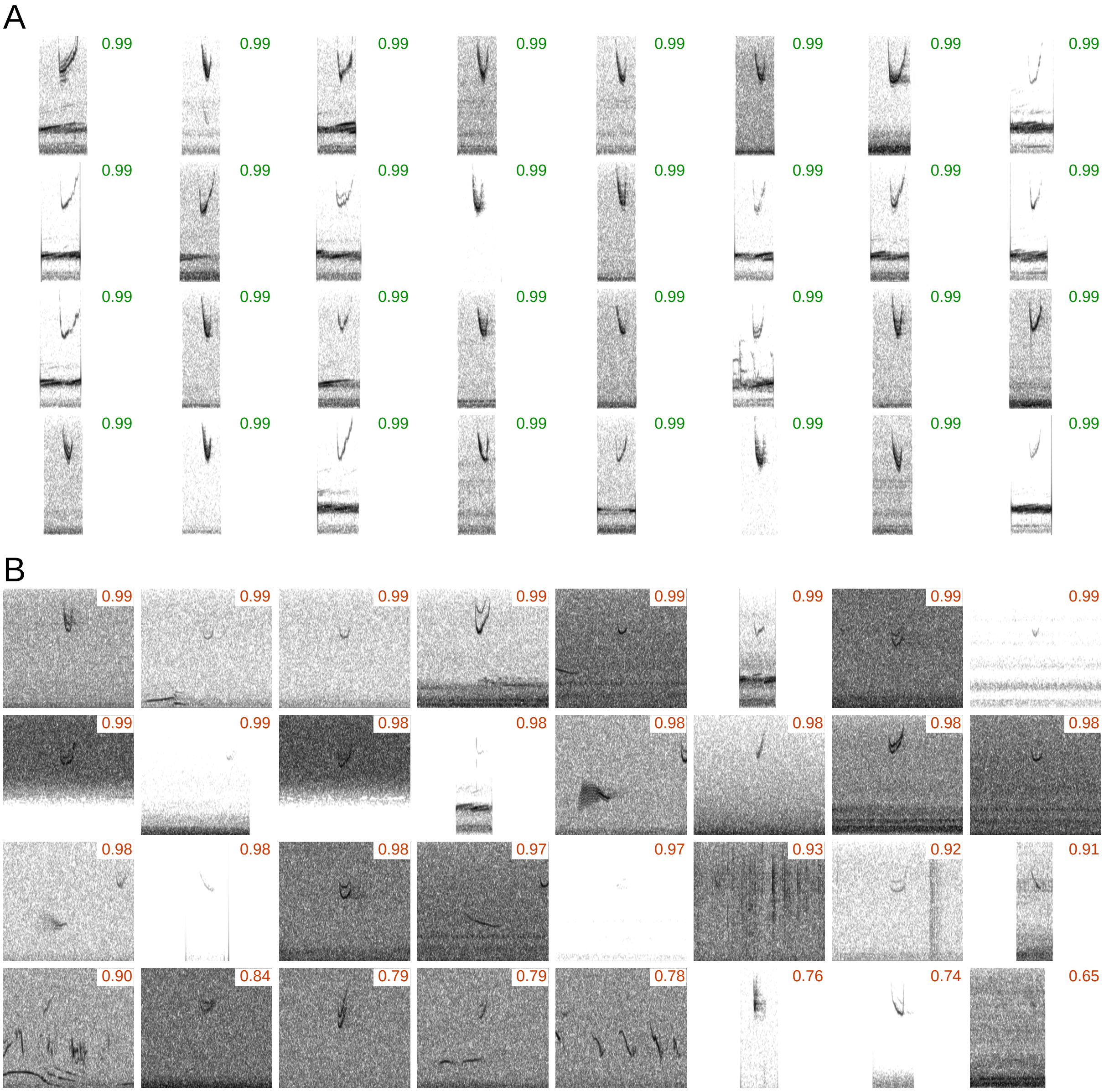


Figure S31: Classification examples for **CUPS** drawn from the test dataset, shown as cropped spectrograms. Numbers in upper-right corners are calibrated probability values returned by the model. (A) Test examples scoring highest for this species. Any incorrect classifications are outlined in red. (B) Incorrectly-classified examples scoring highest for this species (i.e., the most confusing cases for the model). Spectrogram parameters: x-axis: 0-0.8 s; y-axis: 1-10 kHz; window type: Hanning; window length: 300 samples; hop size: 30 samples; dynamic range floor: -50 dB).

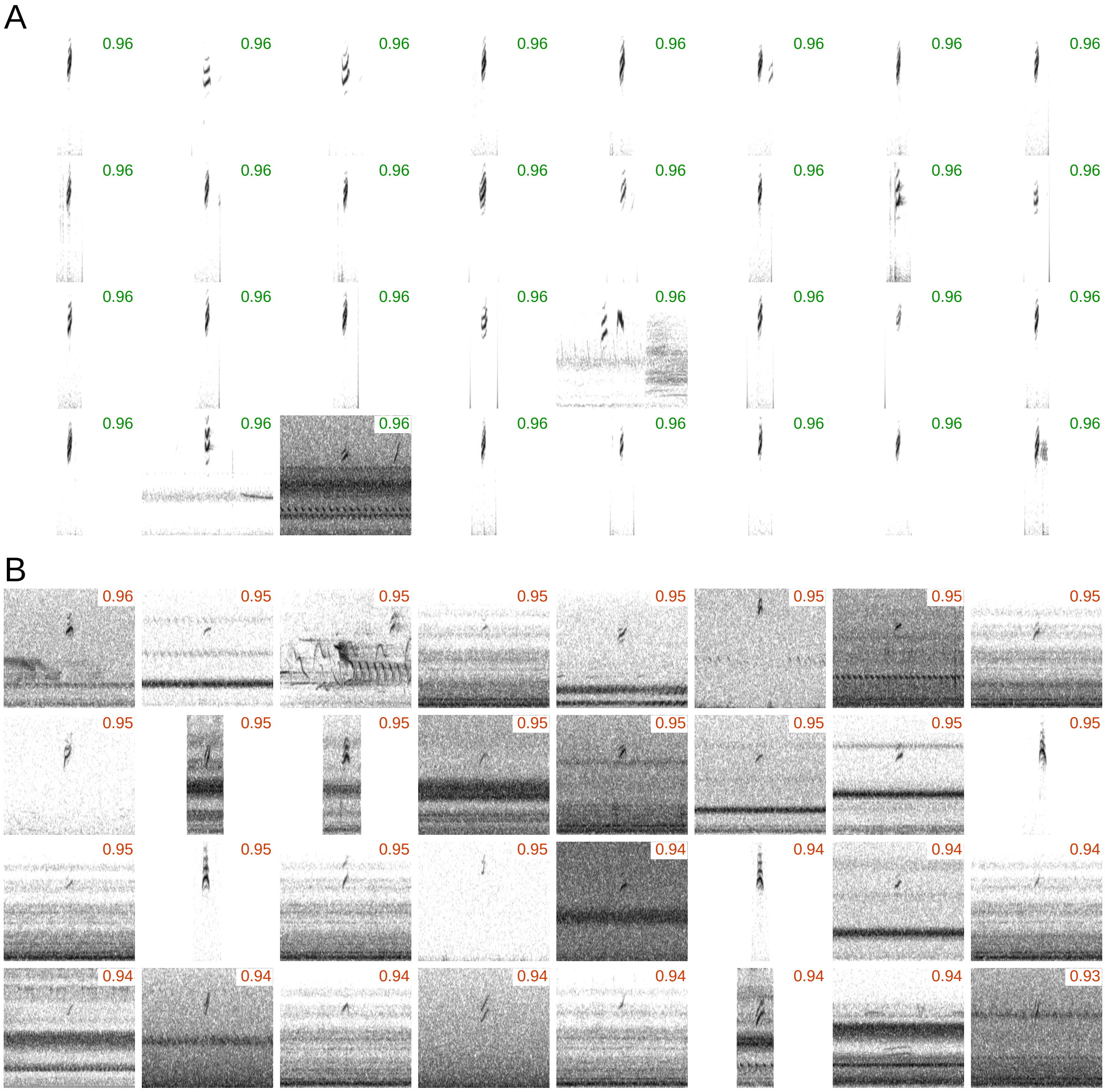


Figure S32: Classification examples for **DBUP** drawn from the test dataset, shown as cropped spectrograms. Numbers in upper-right corners are calibrated probability values returned by the model. (A) Test examples scoring highest for this species. Any incorrect classifications are outlined in red. (B) Incorrectly-classified examples scoring highest for this species (i.e., the most confusing cases for the model). Spectrogram parameters: x-axis: 0-0.8 s; y-axis: 1-10 kHz; window type: Hanning; window length: 300 samples; hop size: 30 samples; dynamic range floor: -50 dB).

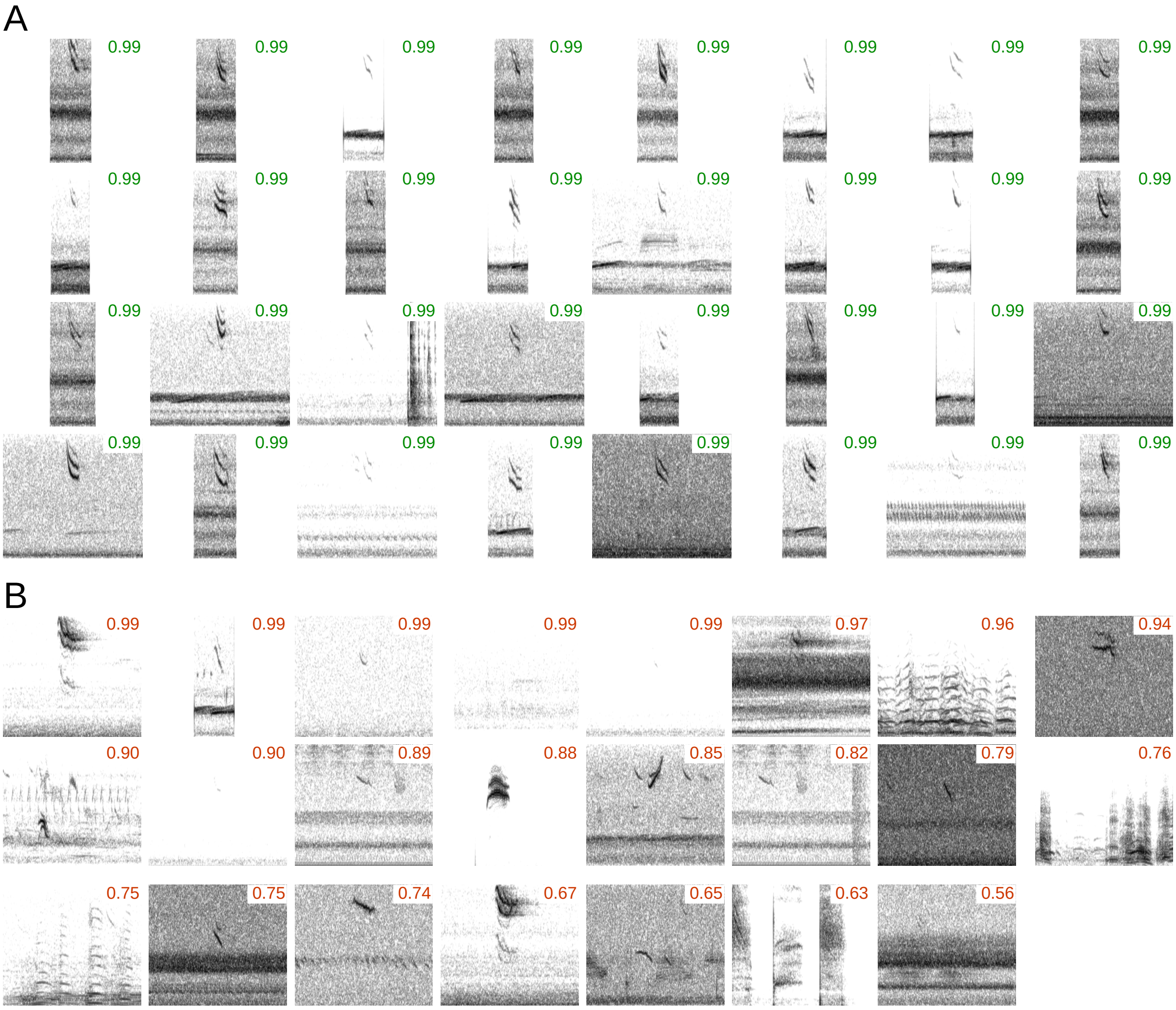


Figure S33: Classification examples for **DESP** drawn from the test dataset, shown as cropped spectrograms. Numbers in upper-right corners are calibrated probability values returned by the model. (A) Test examples scoring highest for this species. Any incorrect classifications are outlined in red. (B) Incorrectly-classified examples scoring highest for this species (i.e., the most confusing cases for the model). Spectrogram parameters: x-axis: 0-0.8 s; y-axis: 1-10 kHz; window type: Hanning; window length: 300 samples; hop size: 30 samples; dynamic range floor: -50 dB).

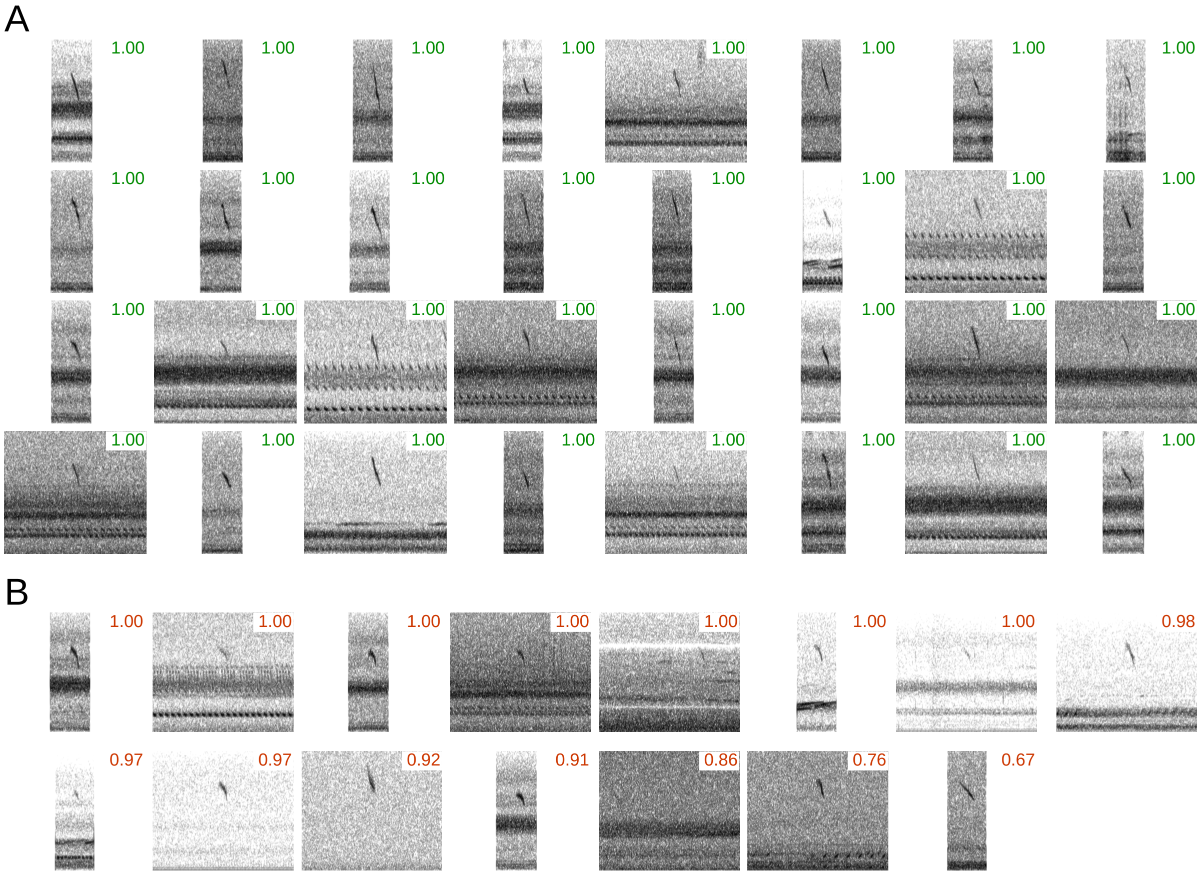


Figure S34: Classification examples for **DEWA** drawn from the test dataset, shown as cropped spectrograms. Numbers in upper-right corners are calibrated probability values returned by the model. (A) Test examples scoring highest for this species. Any incorrect classifications are outlined in red. (B) Incorrectly-classified examples scoring highest for this species (i.e., the most confusing cases for the model). Spectrogram parameters: x-axis: 0-0.8 s; y-axis: 1-10 kHz; window type: Hanning; window length: 300 samples; hop size: 30 samples; dynamic range floor: -50 dB).

A

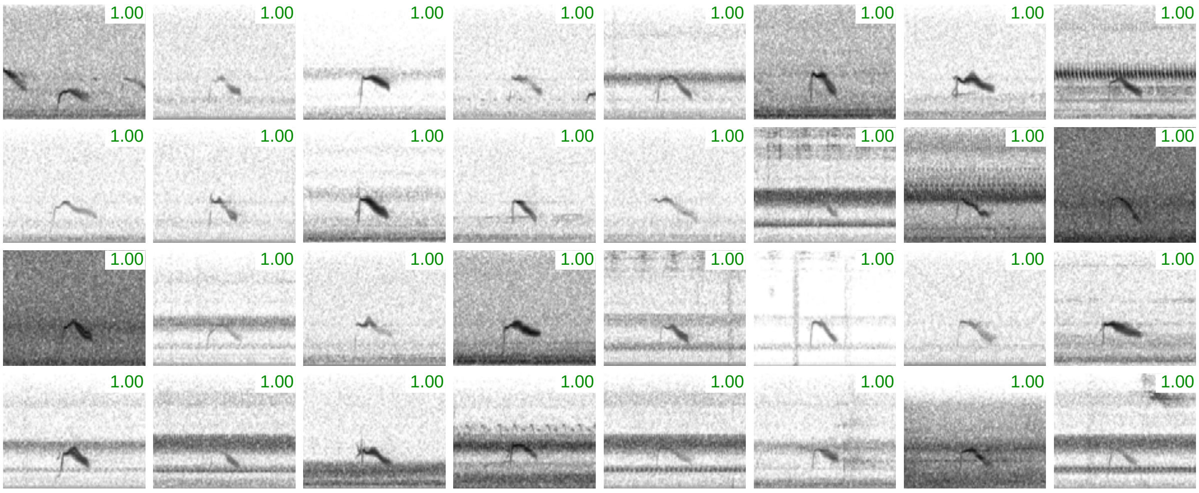


B

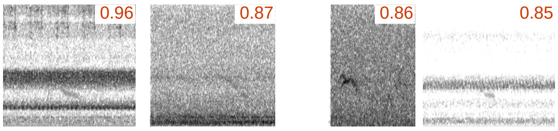


Figure S35: Classification examples for **GCBI** drawn from the test dataset, shown as cropped spectrograms. Numbers in upper-right corners are calibrated probability values returned by the model. (A) Test examples scoring highest for this species. Any incorrect classifications are outlined in red. (B) Incorrectly-classified examples scoring highest for this species (i.e., the most confusing cases for the model). Spectrogram parameters: x-axis: 0-0.8 s; y-axis: 1-10 kHz; window type: Hanning; window length: 300 samples; hop size: 30 samples; dynamic range floor: -50 dB).

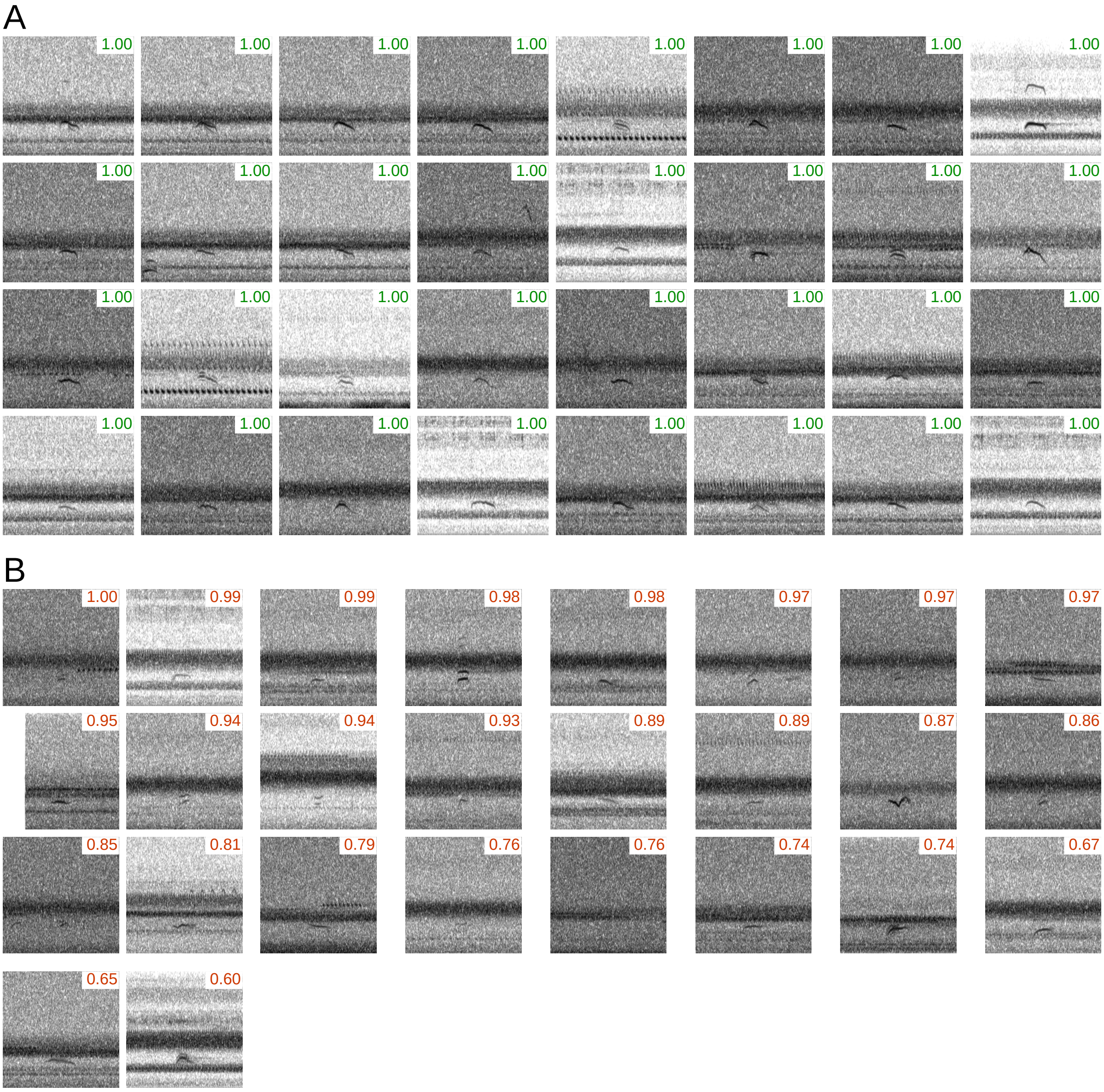


Figure S36: Classification examples for **GROS** drawn from the test dataset, shown as cropped spectrograms. Numbers in upper-right corners are calibrated probability values returned by the model. (A) Test examples scoring highest for this species. Any incorrect classifications are outlined in red. (B) Incorrectly-classified examples scoring highest for this species (i.e., the most confusing cases for the model). Spectrogram parameters: x-axis: 0-0.8 s; y-axis: 1-10 kHz; window type: Hanning; window length: 300 samples; hop size: 30 samples; dynamic range floor: -50 dB).

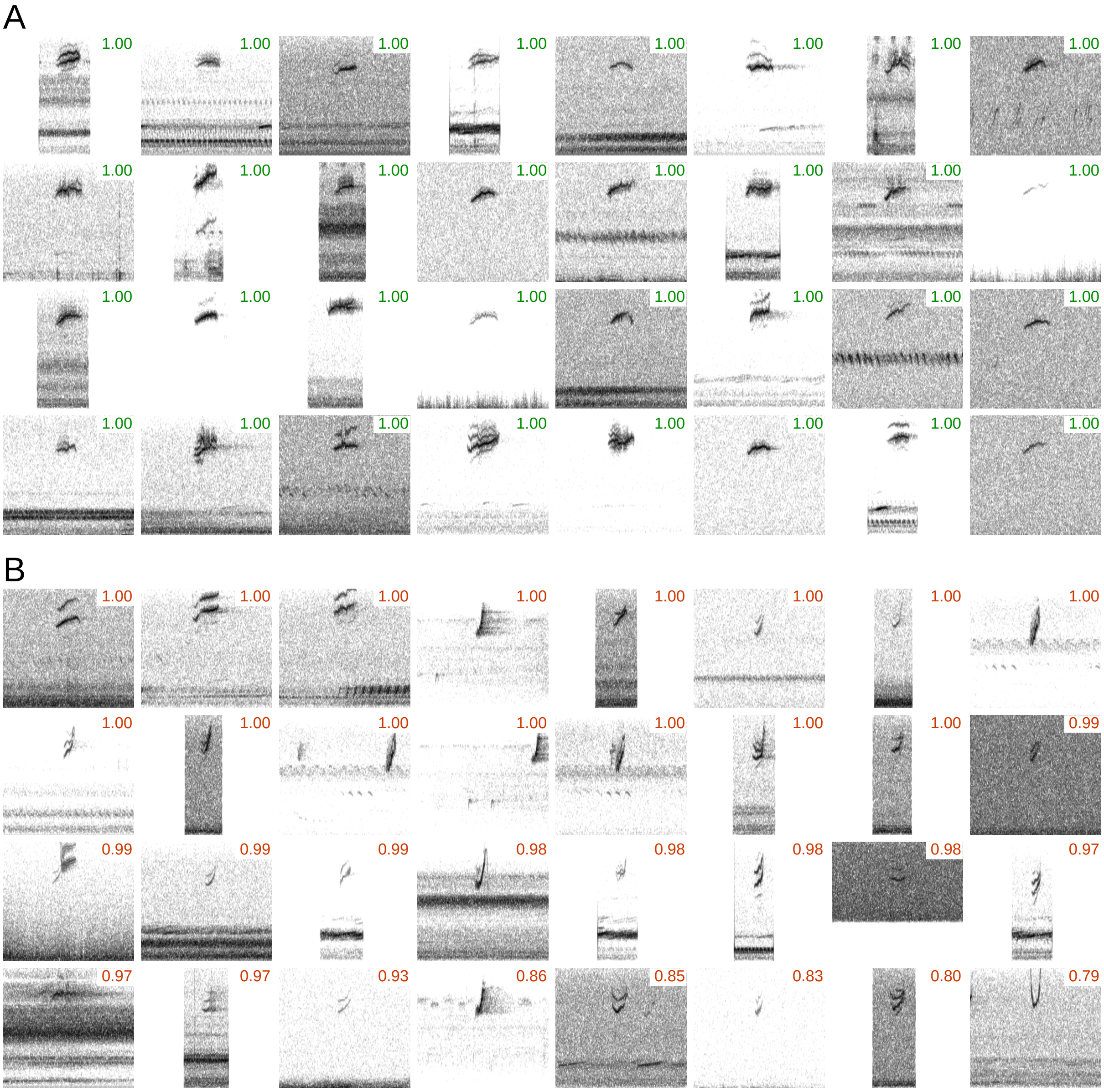


Figure S37: Classification examples for **HSSP** drawn from the test dataset, shown as cropped spectrograms. Numbers in upper-right corners are calibrated probability values returned by the model. (A) Test examples scoring highest for this species. Any incorrect classifications are outlined in red. (B) Incorrectly-classified examples scoring highest for this species (i.e., the most confusing cases for the model). Spectrogram parameters: x-axis: 0-0.8 s; y-axis: 1-10 kHz; window type: Hanning; window length: 300 samples; hop size: 30 samples; dynamic range floor: -50 dB).

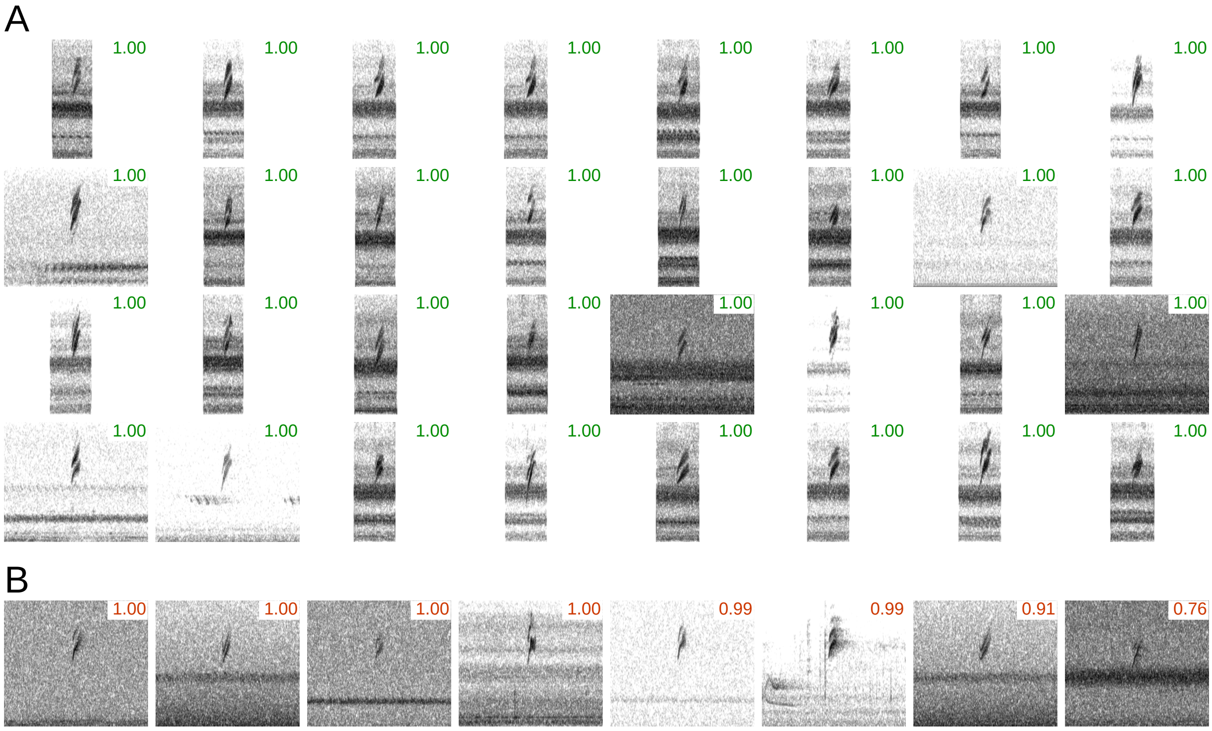


Figure S38: Classification examples for **MWAR** drawn from the test dataset, shown as cropped spectrograms. Numbers in upper-right corners are calibrated probability values returned by the model. (A) Test examples scoring highest for this species. Any incorrect classifications are outlined in red. (B) Incorrectly-classified examples scoring highest for this species (i.e., the most confusing cases for the model). Spectrogram parameters: x-axis: 0-0.8 s; y-axis: 1-10 kHz; window type: Hanning; window length: 300 samples; hop size: 30 samples; dynamic range floor: -50 dB).

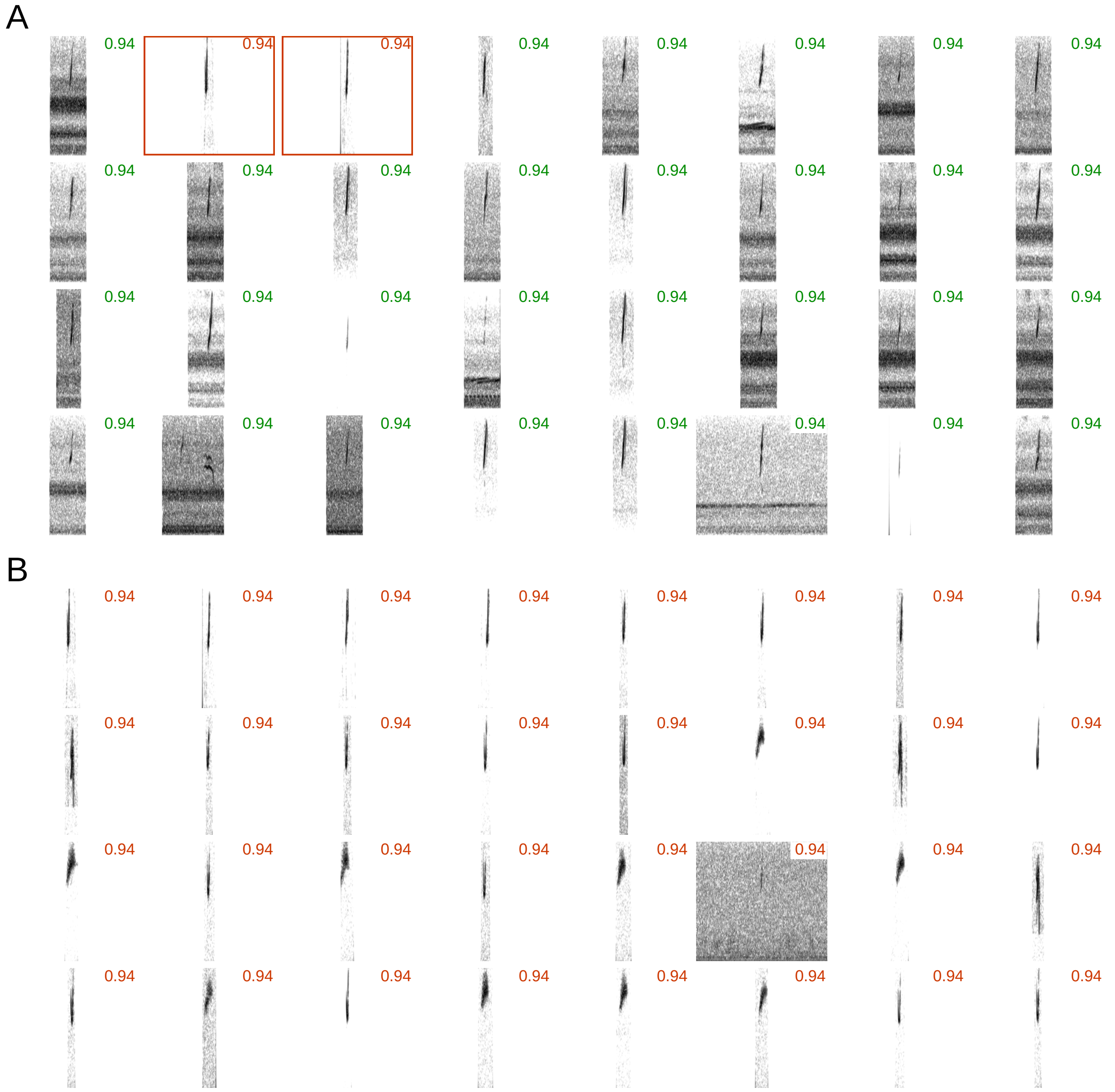


Figure S39: Classification examples for **SBUF** drawn from the test dataset, shown as cropped spectrograms. Numbers in upper-right corners are calibrated probability values returned by the model. (A) Test examples scoring highest for this species. Any incorrect classifications are outlined in red. (B) Incorrectly-classified examples scoring highest for this species (i.e., the most confusing cases for the model). Spectrogram parameters: x-axis: 0-0.8 s; y-axis: 1-10 kHz; window type: Hanning; window length: 300 samples; hop size: 30 samples; dynamic range floor: -50 dB).

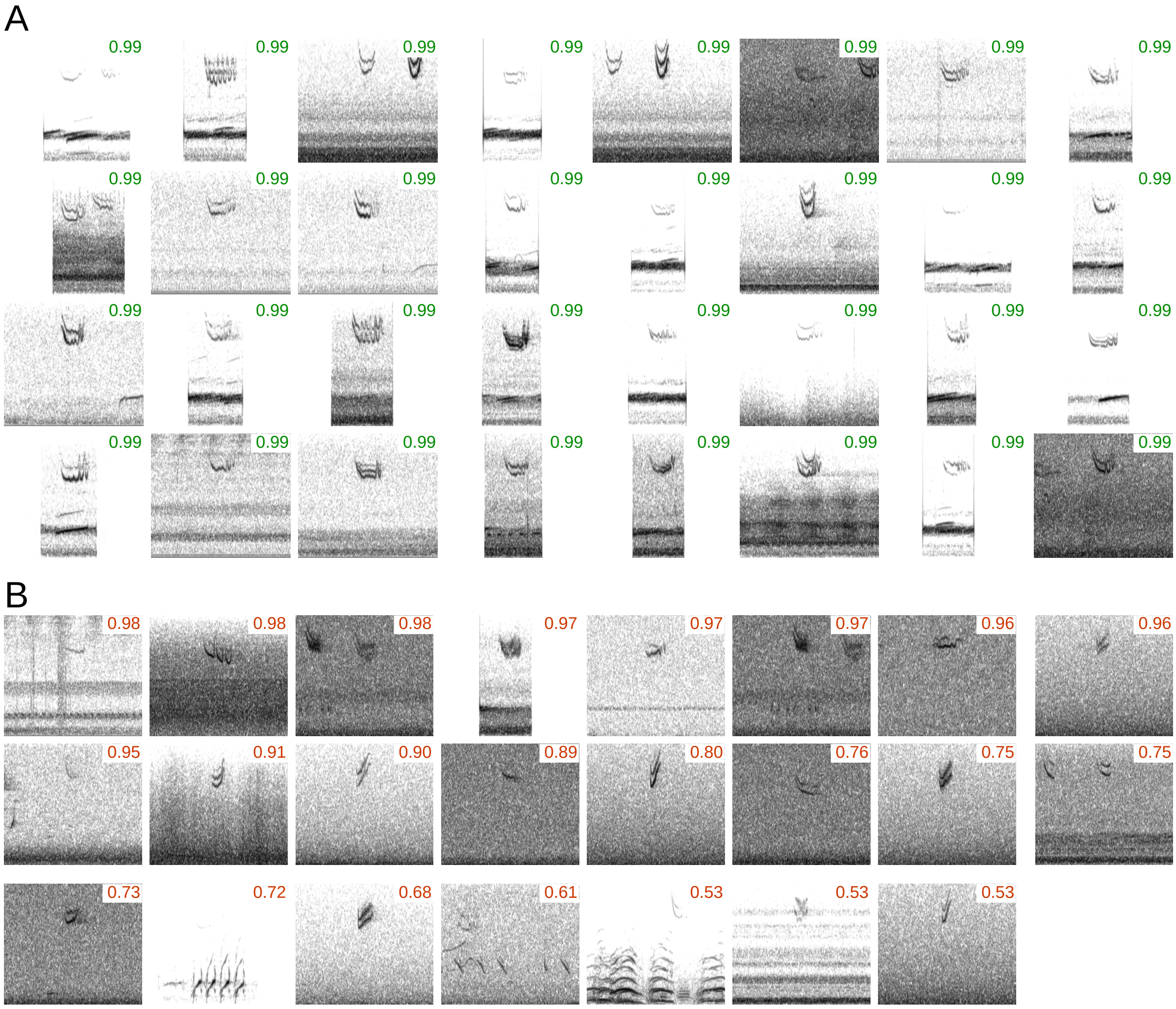


Figure S40: Classification examples for **SFHS** drawn from the test dataset, shown as cropped spectrograms. Numbers in upper-right corners are calibrated probability values returned by the model. (A) Test examples scoring highest for this species. Any incorrect classifications are outlined in red. (B) Incorrectly-classified examples scoring highest for this species (i.e., the most confusing cases for the model). Spectrogram parameters: x-axis: 0-0.8 s; y-axis: 1-10 kHz; window type: Hanning; window length: 300 samples; hop size: 30 samples; dynamic range floor: -50 dB).

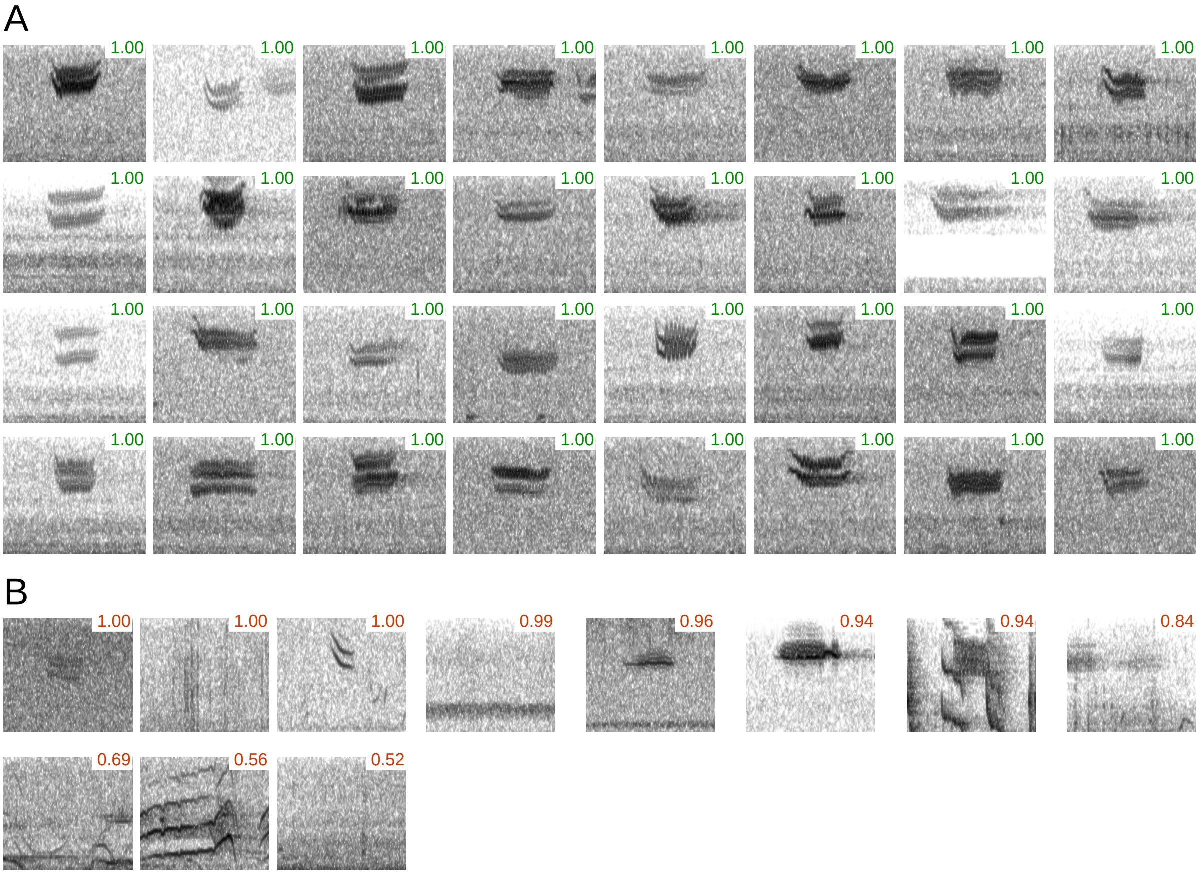


Figure S41: Classification examples for **SWLI** drawn from the test dataset, shown as cropped spectrograms. Numbers in upper-right corners are calibrated probability values returned by the model. (A) Test examples scoring highest for this species. Any incorrect classifications are outlined in red. (B) Incorrectly-classified examples scoring highest for this species (i.e., the most confusing cases for the model). Spectrogram parameters: x-axis: 0-0.3 s; y-axis: 3-11 kHz; window type: Hanning; window length: 200 samples; hop size: 20 samples; dynamic range floor: -60 dB).

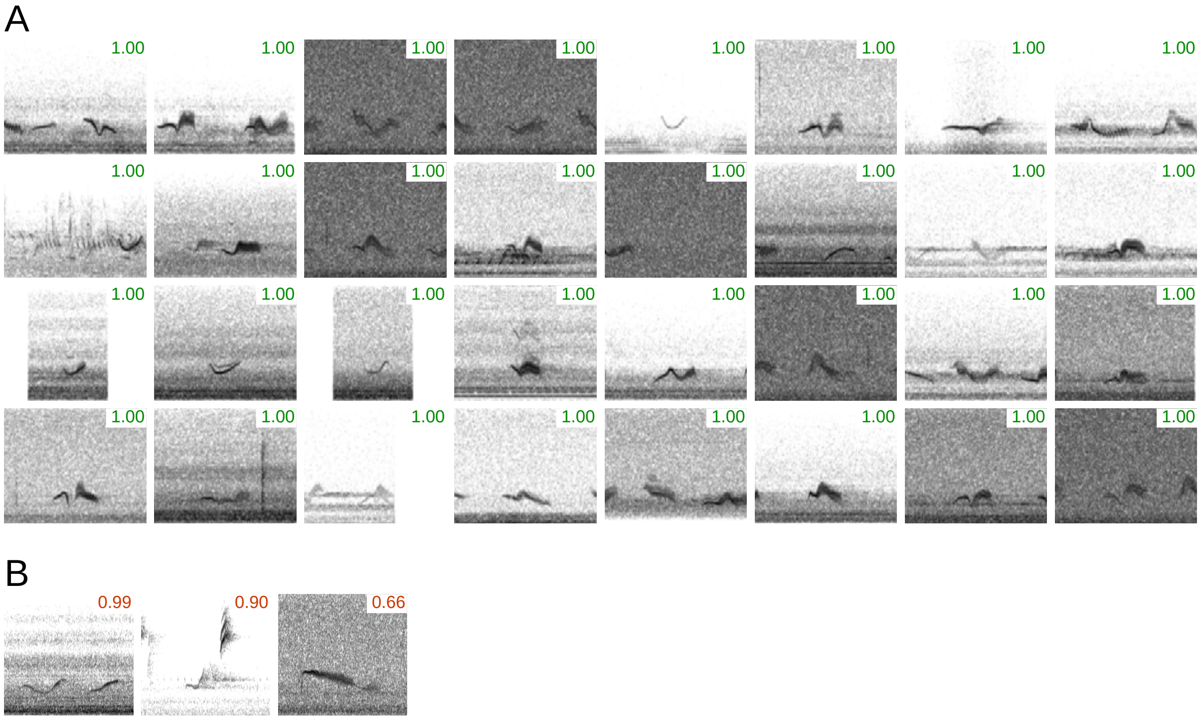


Figure S42: Classification examples for **TANA** drawn from the test dataset, shown as cropped spectrograms. Numbers in upper-right corners are calibrated probability values returned by the model. (A) Test examples scoring highest for this species. Any incorrect classifications are outlined in red. (B) Incorrectly-classified examples scoring highest for this species (i.e., the most confusing cases for the model). Spectrogram parameters: x-axis: 0-0.8 s; y-axis: 1-10 kHz; window type: Hanning; window length: 300 samples; hop size: 30 samples; dynamic range floor: -50 dB).

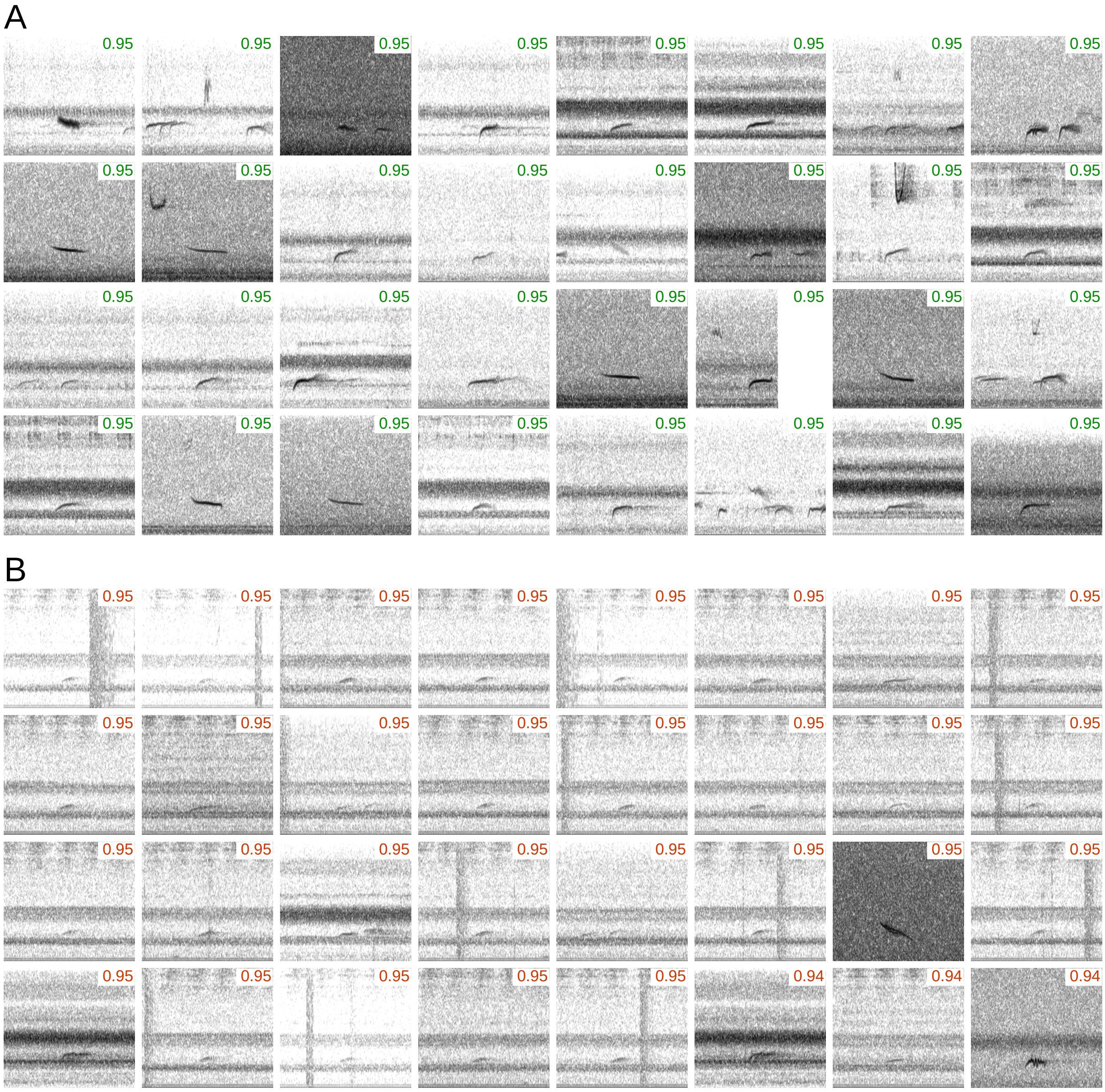


Figure S43: Classification examples for **THSH** drawn from the test dataset, shown as cropped spectrograms. Numbers in upper-right corners are calibrated probability values returned by the model. (A) Test examples scoring highest for this species. Any incorrect classifications are outlined in red. (B) Incorrectly-classified examples scoring highest for this species (i.e., the most confusing cases for the model). Spectrogram parameters: x-axis: 0-0.8 s; y-axis: 1-10 kHz; window type: Hanning; window length: 300 samples; hop size: 30 samples; dynamic range floor: -50 dB).

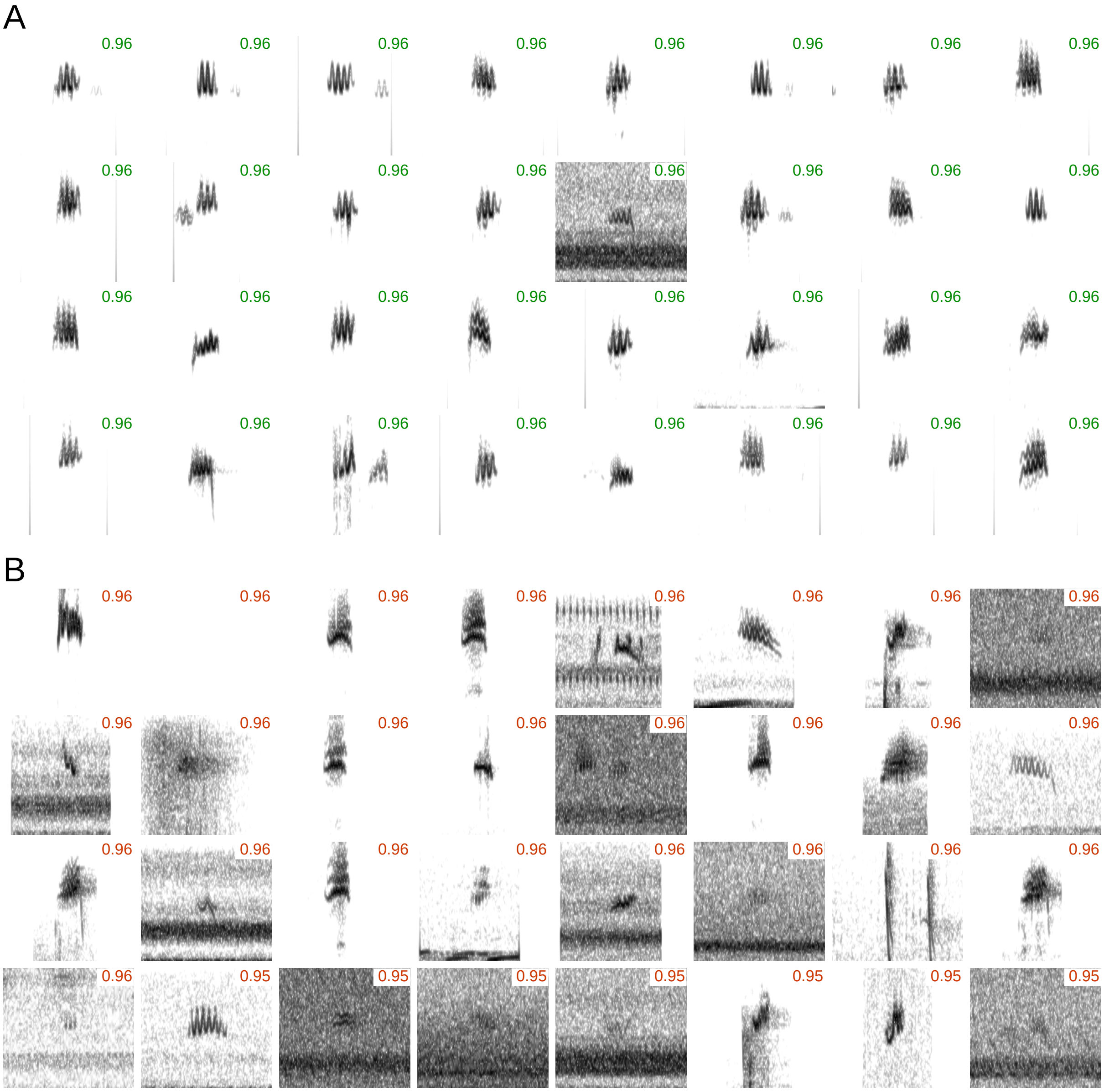


Figure S44: Classification examples for **ZEEP** drawn from the test dataset, shown as cropped spectrograms. Numbers in upper-right corners are calibrated probability values returned by the model. (A) Test examples scoring highest for this species. Any incorrect classifications are outlined in red. (B) Incorrectly-classified examples scoring highest for this species (i.e., the most confusing cases for the model). Spectrogram parameters: x-axis: 0-0.3 s; y-axis: 3-10.5 kHz; window type: Hanning; window length: 200 samples; hop size: 20 samples; dynamic range floor: -50 dB).

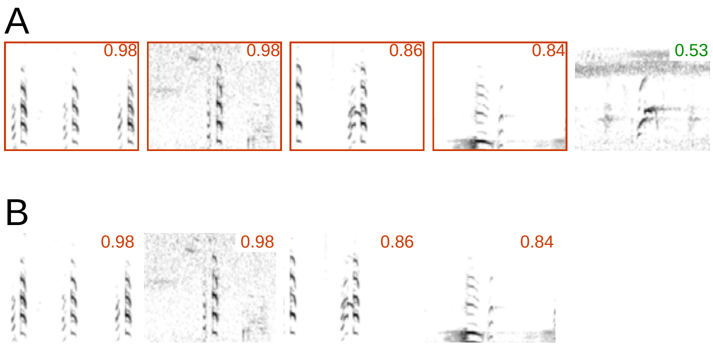


Figure S45: Classification examples for **ameavo** drawn from the test dataset, shown as cropped spectrograms. Numbers in upper-right corners are calibrated probability values returned by the model. (A) Test examples scoring highest for this species. Any incorrect classifications are outlined in red. (B) Incorrectly-classified examples scoring highest for this species (i.e., the most confusing cases for the model). Spectrogram parameters: x-axis: 0-0.8 s; y-axis: 1-8 kHz; window type: Hanning; window length: 400 samples; hop size: 30 samples; dynamic range floor: -45 dB).

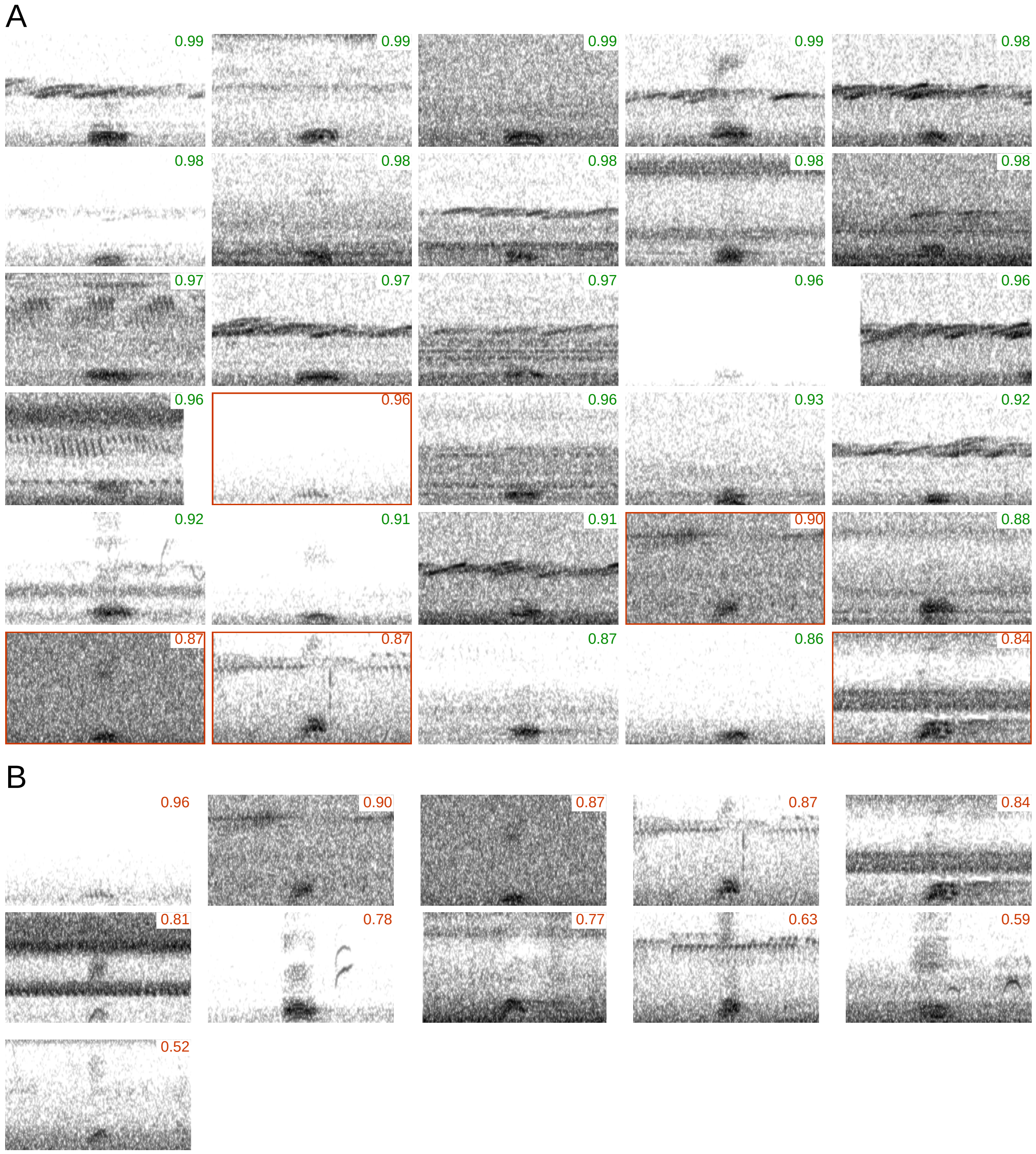


Figure S46: Classification examples for **amebit** drawn from the test dataset, shown as cropped spectrograms. Numbers in upper-right corners are calibrated probability values returned by the model. (A) Test examples scoring highest for this species. Any incorrect classifications are outlined in red. (B) Incorrectly-classified examples scoring highest for this species (i.e., the most confusing cases for the model). Spectrogram parameters: x-axis: 0-0.8 s; y-axis: 1-5 kHz; window type: Hanning; window length: 400 samples; hop size: 30 samples; dynamic range floor: -45 dB).

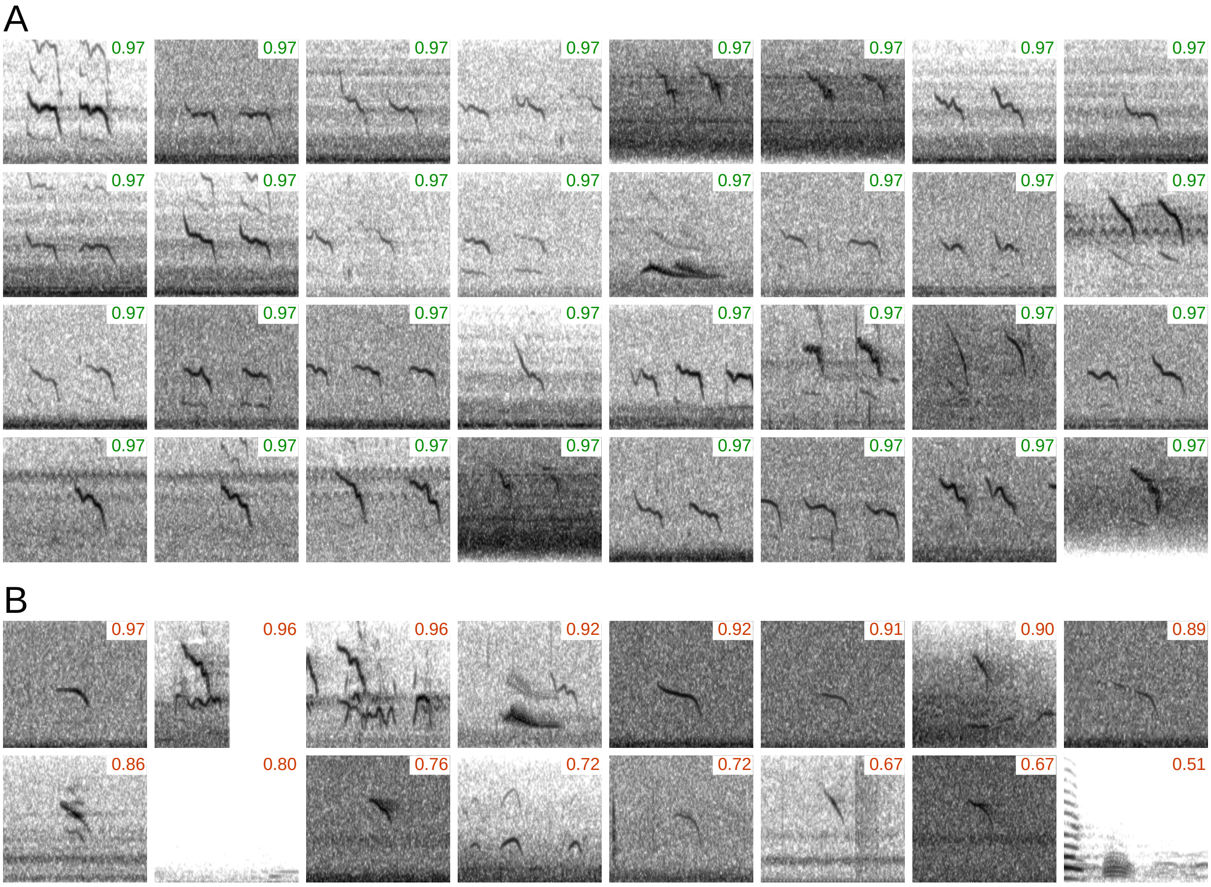


Figure S47: Classification examples for **amepip** drawn from the test dataset, shown as cropped spectrograms. Numbers in upper-right corners are calibrated probability values returned by the model. (A) Test examples scoring highest for this species. Any incorrect classifications are outlined in red. (B) Incorrectly-classified examples scoring highest for this species (i.e., the most confusing cases for the model). Spectrogram parameters: x-axis: 0-0.3 s; y-axis: 1-10 kHz; window type: Hanning; window length: 200 samples; hop size: 20 samples; dynamic range floor: -60 dB).

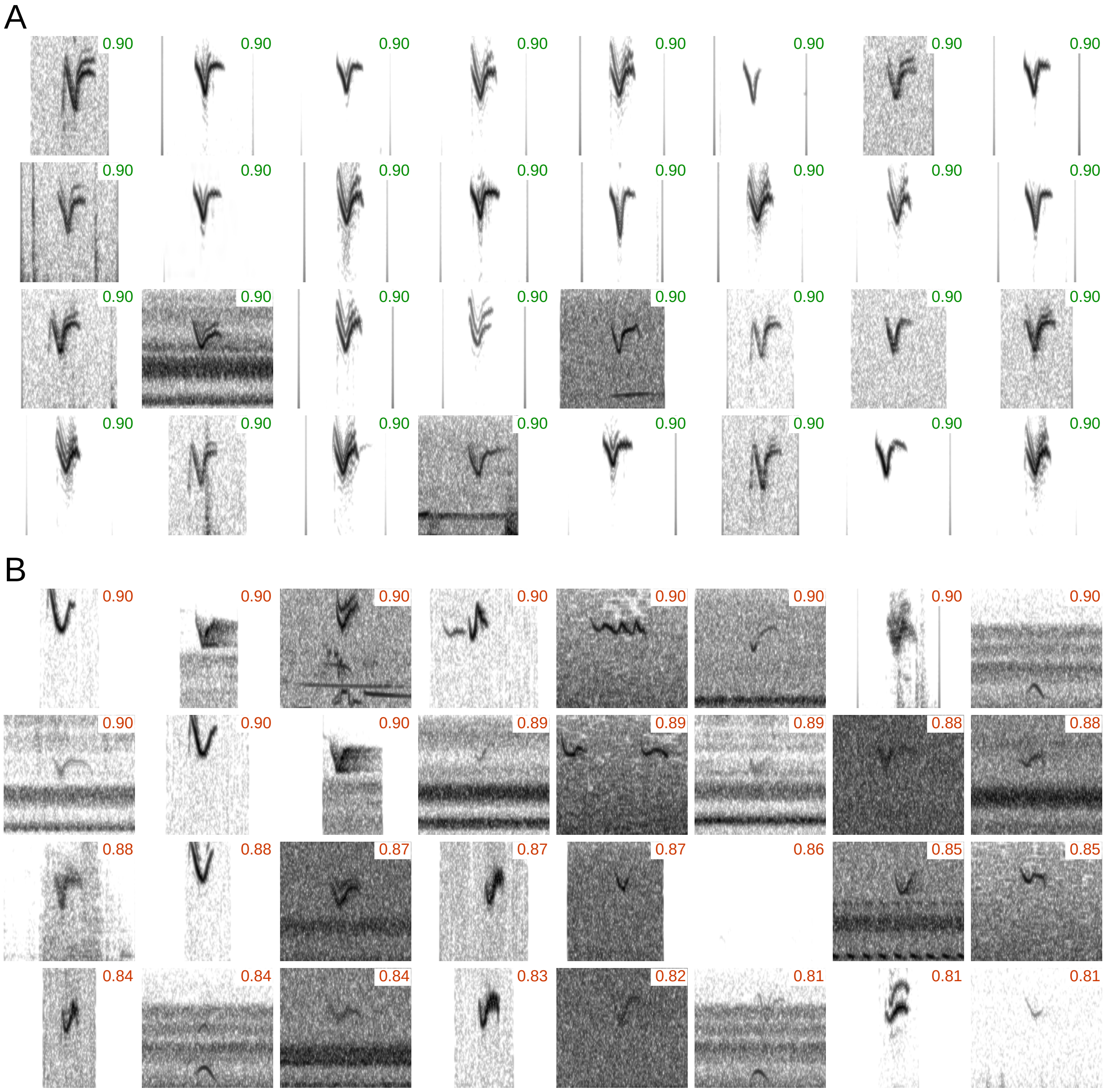


Figure S48: Classification examples for **amered** drawn from the test dataset, shown as cropped spectrograms. Numbers in upper-right corners are calibrated probability values returned by the model. (A) Test examples scoring highest for this species. Any incorrect classifications are outlined in red. (B) Incorrectly-classified examples scoring highest for this species (i.e., the most confusing cases for the model). Spectrogram parameters: x-axis: 0-0.3 s; y-axis: 2-10 kHz; window type: Hanning; window length: 200 samples; hop size: 20 samples; dynamic range floor: -60 dB).

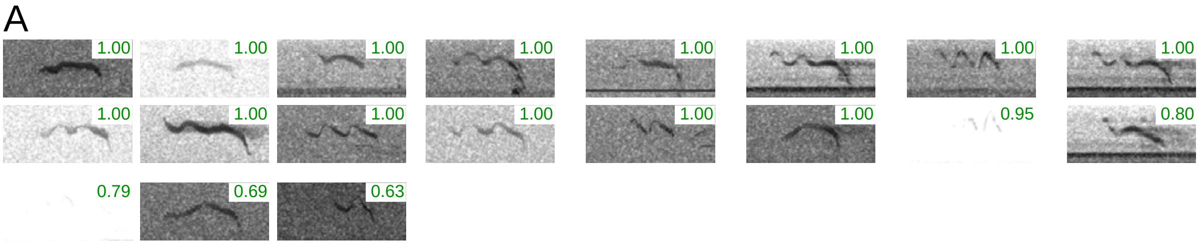


B

No highly confident incorrect clips.

Figure S49: Classification examples for **amerob** drawn from the test dataset, shown as cropped spectrograms. Numbers in upper-right corners are calibrated probability values returned by the model. (A) Test examples scoring highest for this species. Any incorrect classifications are outlined in red. (B) Incorrectly-classified examples scoring highest for this species (i.e., the most confusing cases for the model). Spectrogram parameters: x-axis: 0-0.3 s; y-axis: 2-10 kHz; window type: Hanning; window length: 200 samples; hop size: 20 samples; dynamic range floor: -60 dB).

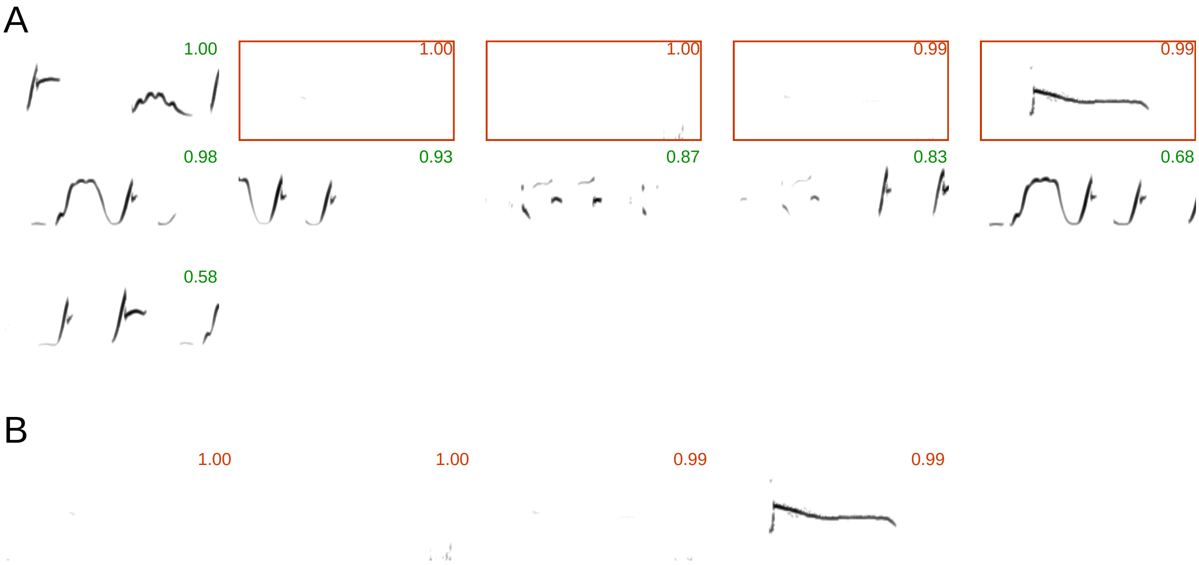


Figure S50: Classification examples for **amgplo** drawn from the test dataset, shown as cropped spectrograms. Numbers in upper-right corners are calibrated probability values returned by the model. (A) Test examples scoring highest for this species. Any incorrect classifications are outlined in red. (B) Incorrectly-classified examples scoring highest for this species (i.e., the most confusing cases for the model). Spectrogram parameters: x-axis: 0-0.8 s; y-axis: 1-5 kHz; window type: Hanning; window length: 400 samples; hop size: 30 samples; dynamic range floor: -30 dB).

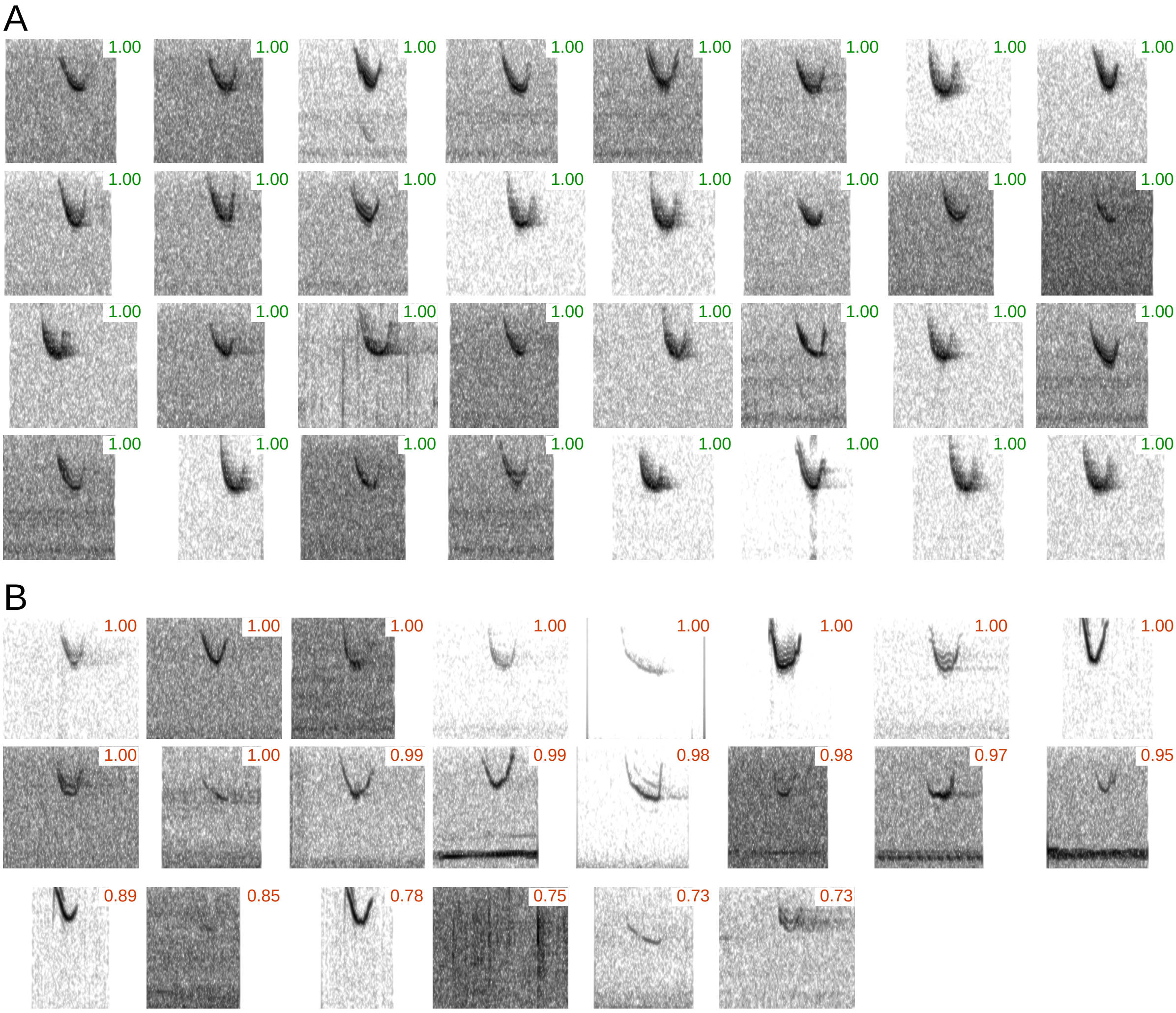


Figure S51: Classification examples for **amtspa** drawn from the test dataset, shown as cropped spectrograms. Numbers in upper-right corners are calibrated probability values returned by the model. (A) Test examples scoring highest for this species. Any incorrect classifications are outlined in red. (B) Incorrectly-classified examples scoring highest for this species (i.e., the most confusing cases for the model). Spectrogram parameters: x-axis: 0-0.3 s; y-axis: 2-10 kHz; window type: Hanning; window length: 200 samples; hop size: 20 samples; dynamic range floor: -60 dB).

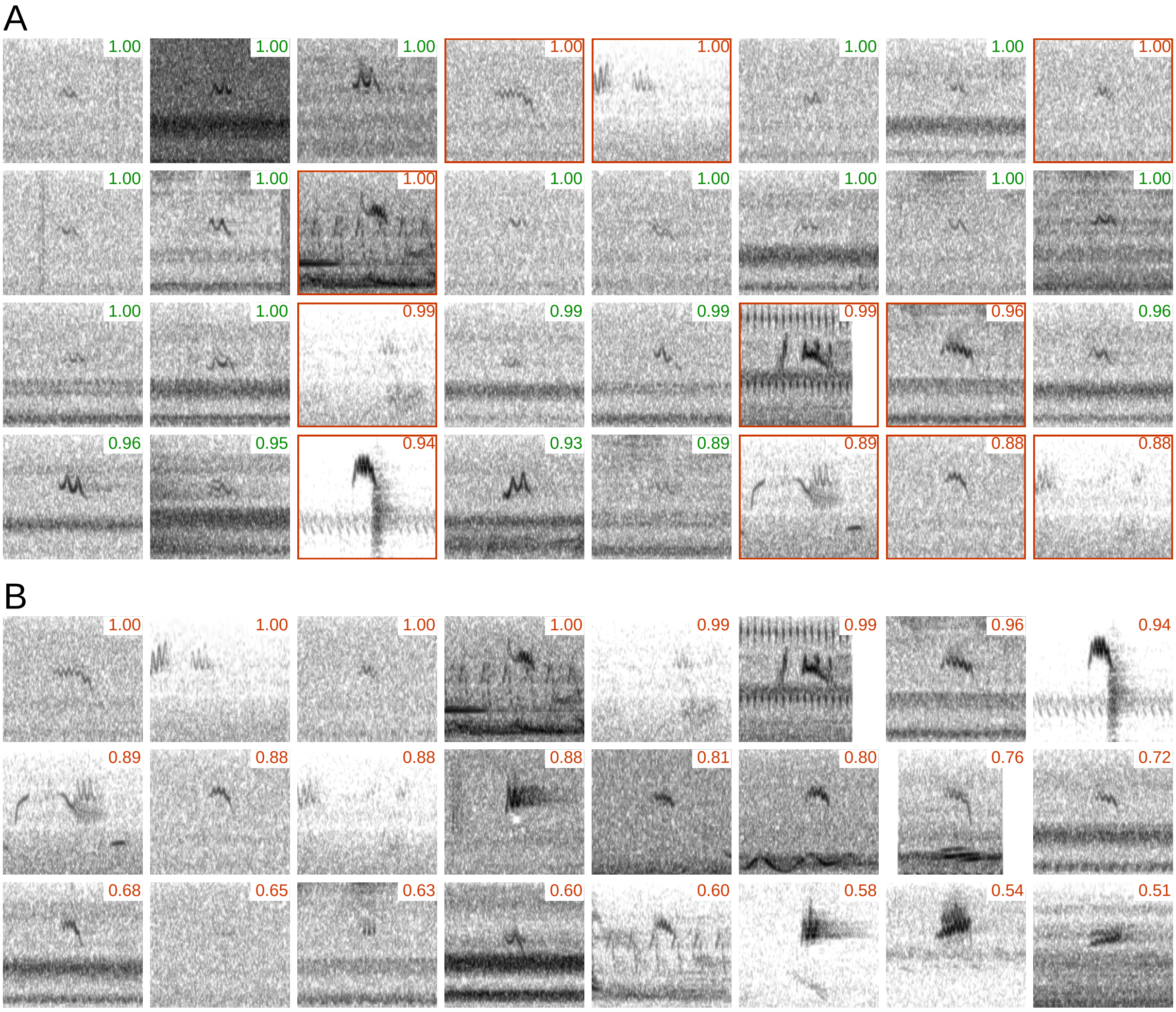

Figure S52: Classification examples for **babwar** drawn from the test dataset, shown as cropped spectrograms. Numbers in upper-right corners are calibrated probability values returned by the model. (A) Test examples scoring highest for this species. Any incorrect classifications are outlined in red. (B) Incorrectly-classified examples scoring highest for this species (i.e., the most confusing cases for the model). Spectrogram parameters: x-axis: 0-0.3 s; y-axis: 2-10 kHz; window type: Hanning; window length: 200 samples; hop size: 20 samples; dynamic range floor: -60 dB).

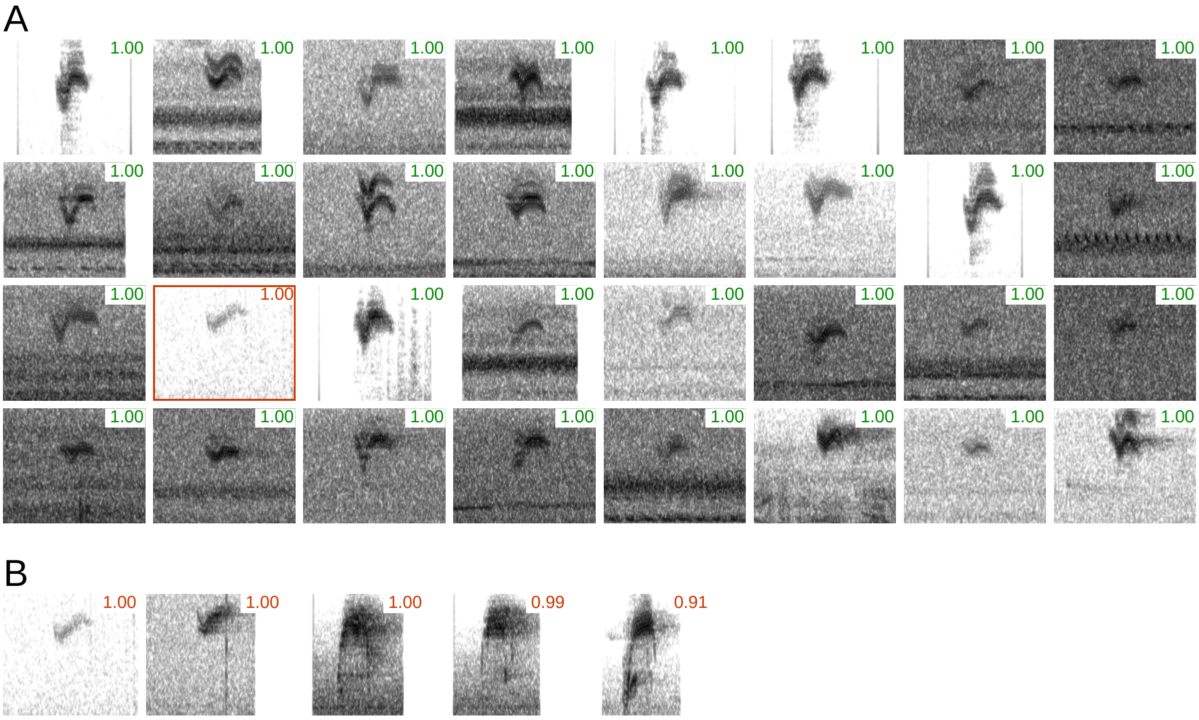


Figure S53: Classification examples for **bawwar** drawn from the test dataset, shown as cropped spectrograms. Numbers in upper-right corners are calibrated probability values returned by the model. (A) Test examples scoring highest for this species. Any incorrect classifications are outlined in red. (B) Incorrectly-classified examples scoring highest for this species (i.e., the most confusing cases for the model). Spectrogram parameters: x-axis: 0-0.3 s; y-axis: 2-10 kHz; window type: Hanning; window length: 200 samples; hop size: 20 samples; dynamic range floor: -60 dB).

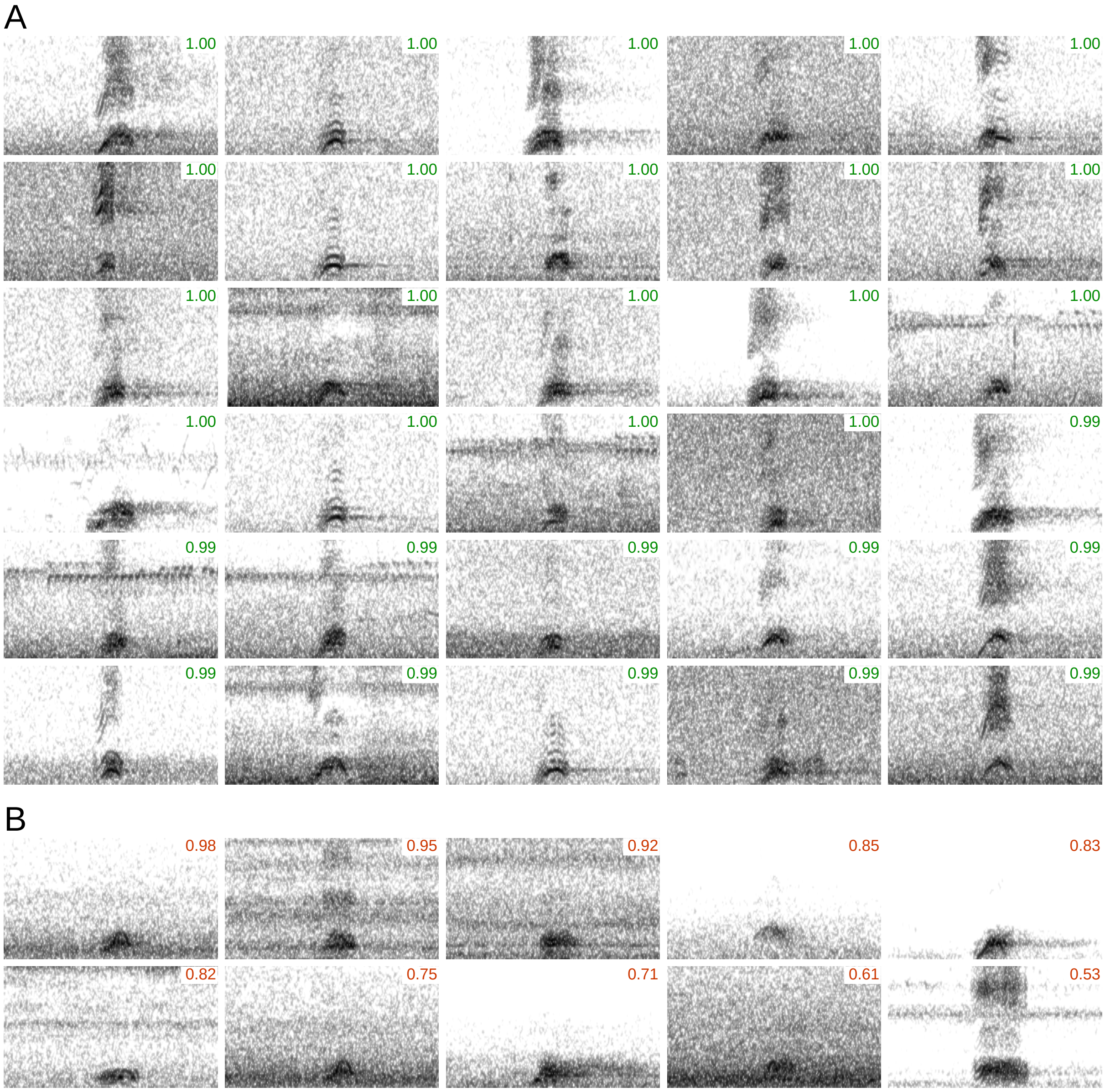


Figure S54: Classification examples for **bcnher** drawn from the test dataset, shown as cropped spectrograms. Numbers in upper-right corners are calibrated probability values returned by the model. (A) Test examples scoring highest for this species. Any incorrect classifications are outlined in red. (B) Incorrectly-classified examples scoring highest for this species (i.e., the most confusing cases for the model). Spectrogram parameters: x-axis: 0-0.8 s; y-axis: 1-5 kHz; window type: Hanning; window length: 400 samples; hop size: 30 samples; dynamic range floor: -45 dB).

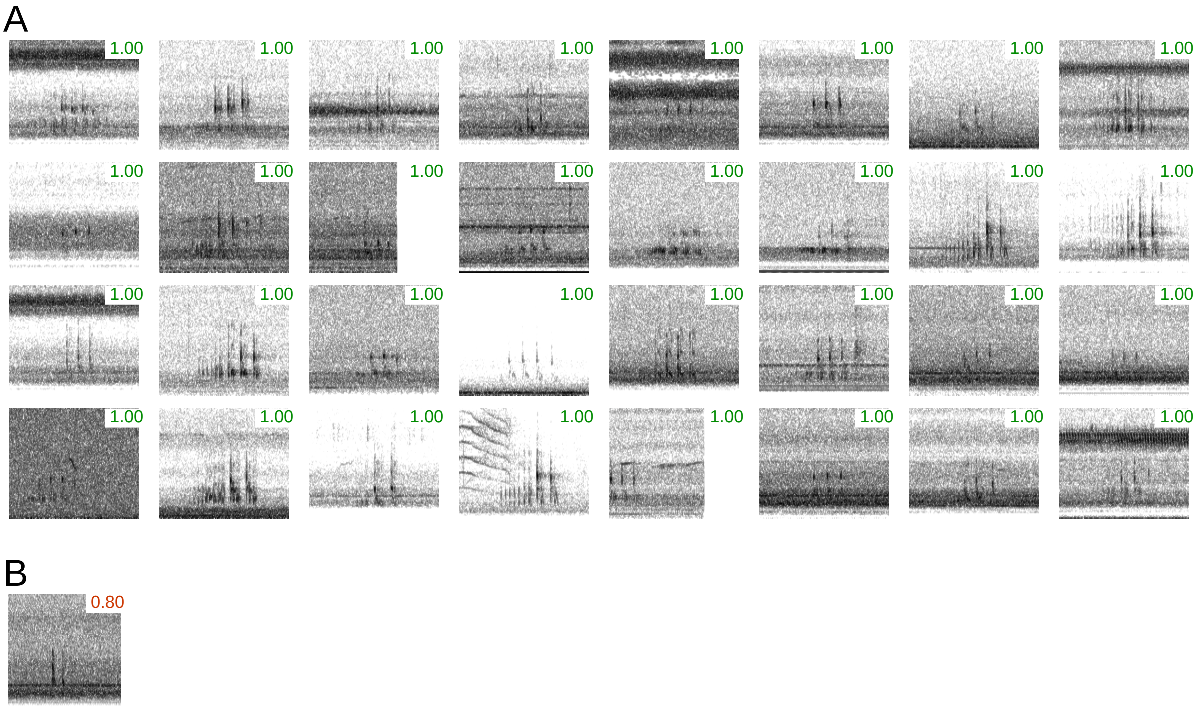


Figure S55: Classification examples for **bkbcuc** drawn from the test dataset, shown as cropped spectrograms. Numbers in upper-right corners are calibrated probability values returned by the model. (A) Test examples scoring highest for this species. Any incorrect classifications are outlined in red. (B) Incorrectly-classified examples scoring highest for this species (i.e., the most confusing cases for the model). Spectrogram parameters: x-axis: 0-1 s; y-axis: 0-6 kHz; window type: Hanning; window length: 420 samples; hop size: 35 samples; dynamic range floor: -50 dB).

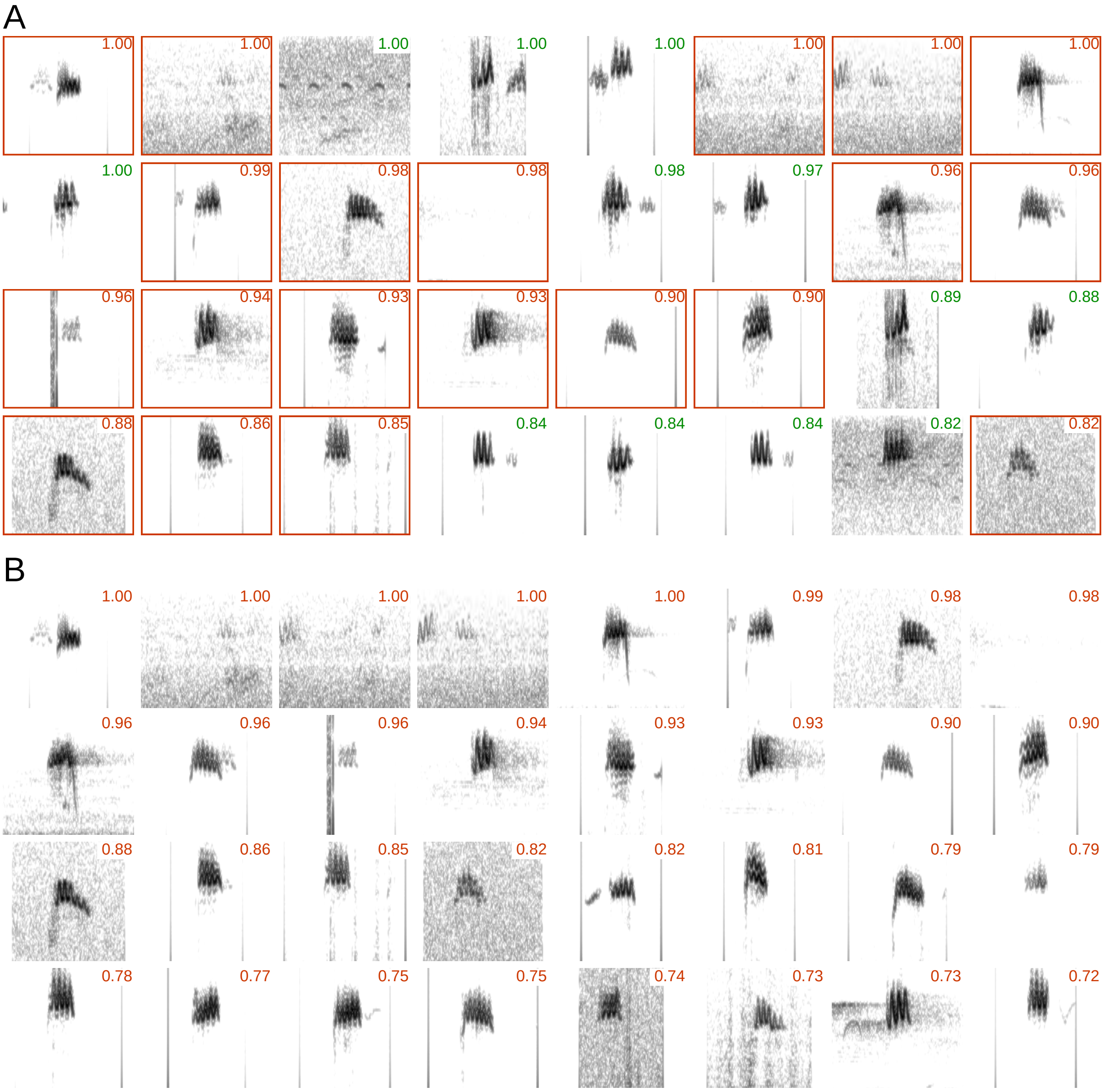


Figure S56: Classification examples for **bkpwar** drawn from the test dataset, shown as cropped spectrograms. Numbers in upper-right corners are calibrated probability values returned by the model. (A) Test examples scoring highest for this species. Any incorrect classifications are outlined in red. (B) Incorrectly-classified examples scoring highest for this species (i.e., the most confusing cases for the model). Spectrogram parameters: x-axis: 0-0.3 s; y-axis: 2-10 kHz; window type: Hanning; window length: 200 samples; hop size: 20 samples; dynamic range floor: -60 dB).

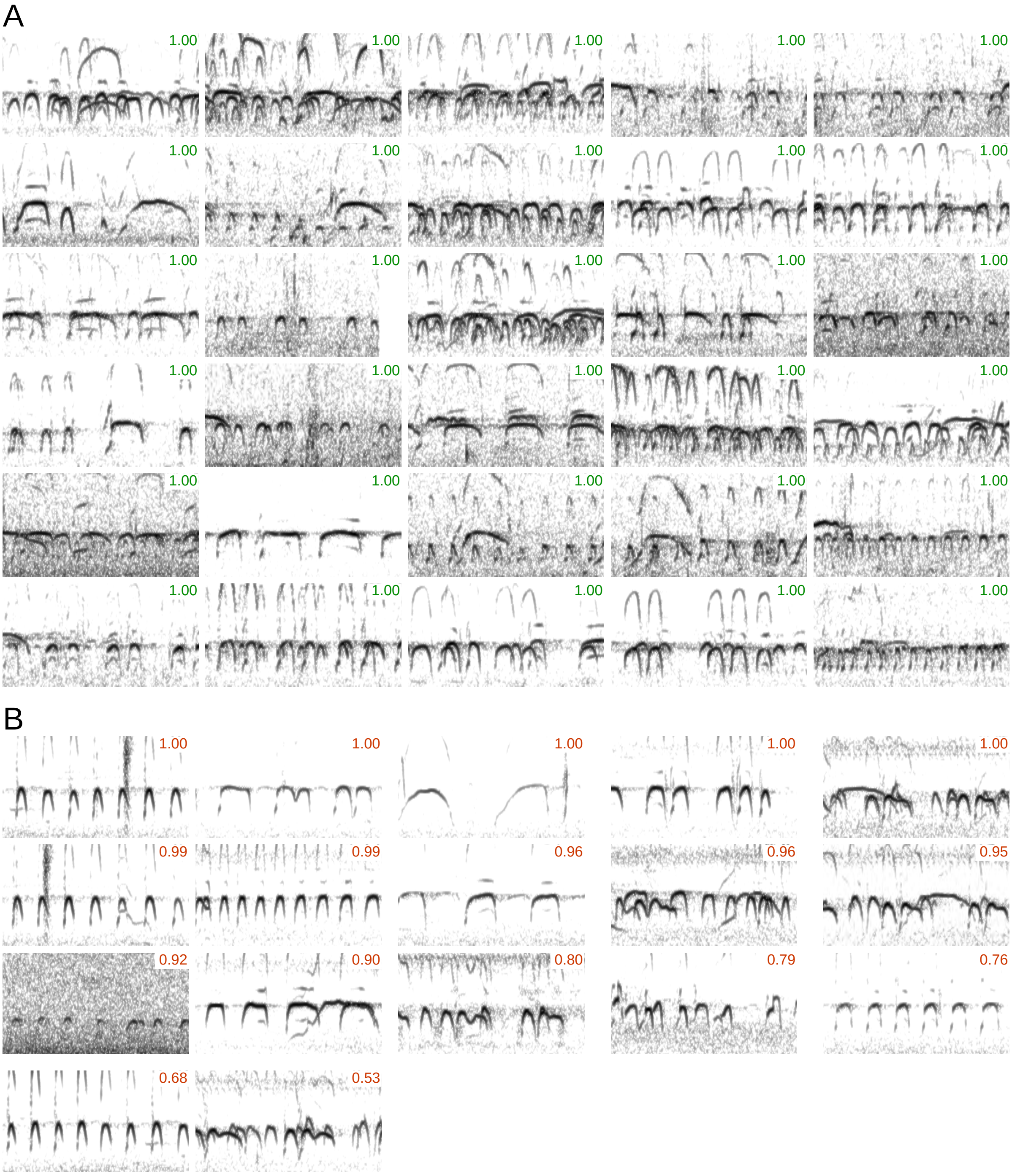


Figure S57: Classification examples for **blkoys** drawn from the test dataset, shown as cropped spectrograms. Numbers in upper-right corners are calibrated probability values returned by the model. (A) Test examples scoring highest for this species. Any incorrect classifications are outlined in red. (B) Incorrectly-classified examples scoring highest for this species (i.e., the most confusing cases for the model). Spectrogram parameters: x-axis: 0-0.8 s; y-axis: 1-5 kHz; window type: Hanning; window length: 400 samples; hop size: 30 samples; dynamic range floor: -45 dB).

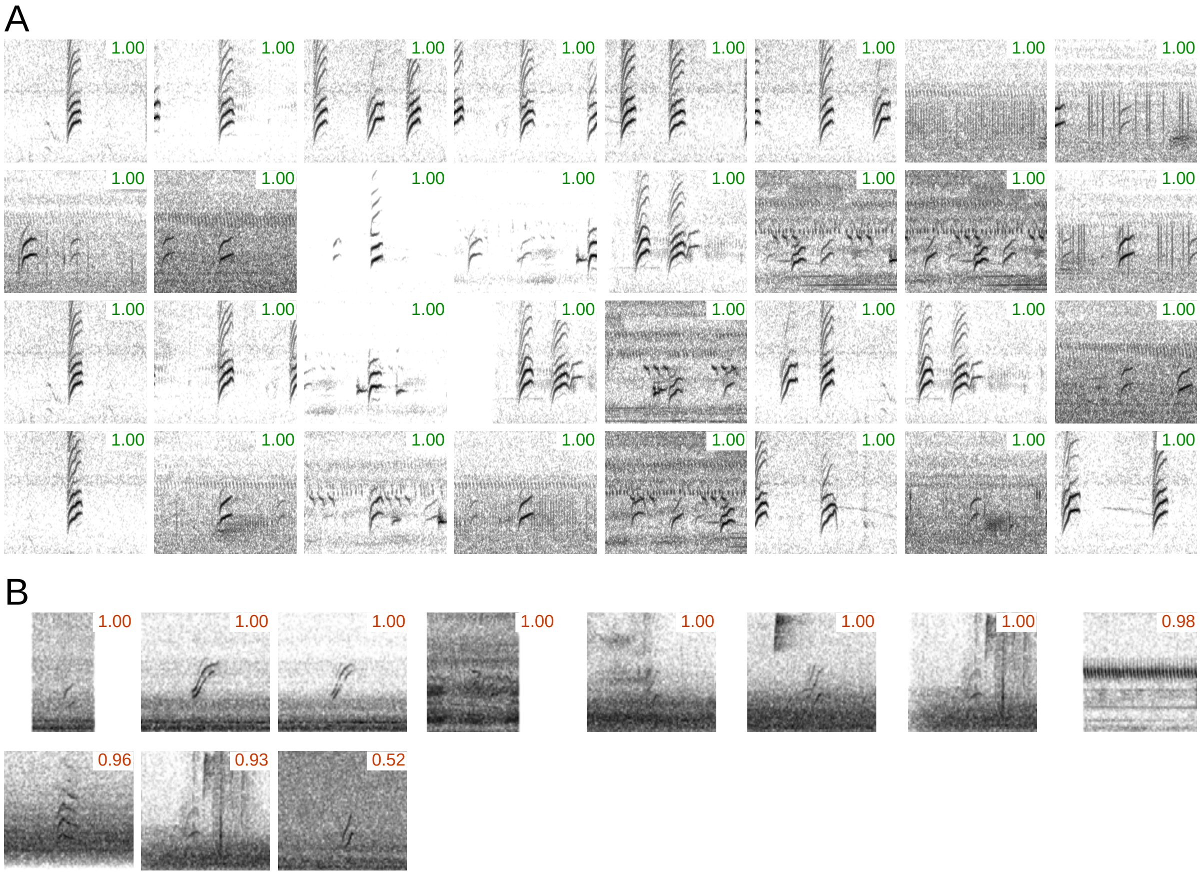


Figure S58: Classification examples for **boboli** drawn from the test dataset, shown as cropped spectrograms. Numbers in upper-right corners are calibrated probability values returned by the model. (A) Test examples scoring highest for this species. Any incorrect classifications are outlined in red. (B) Incorrectly-classified examples scoring highest for this species (i.e., the most confusing cases for the model). Spectrogram parameters: x-axis: 0-0.8 s; y-axis: 1-8 kHz; window type: Hanning; window length: 400 samples; hop size: 30 samples; dynamic range floor: -45 dB).

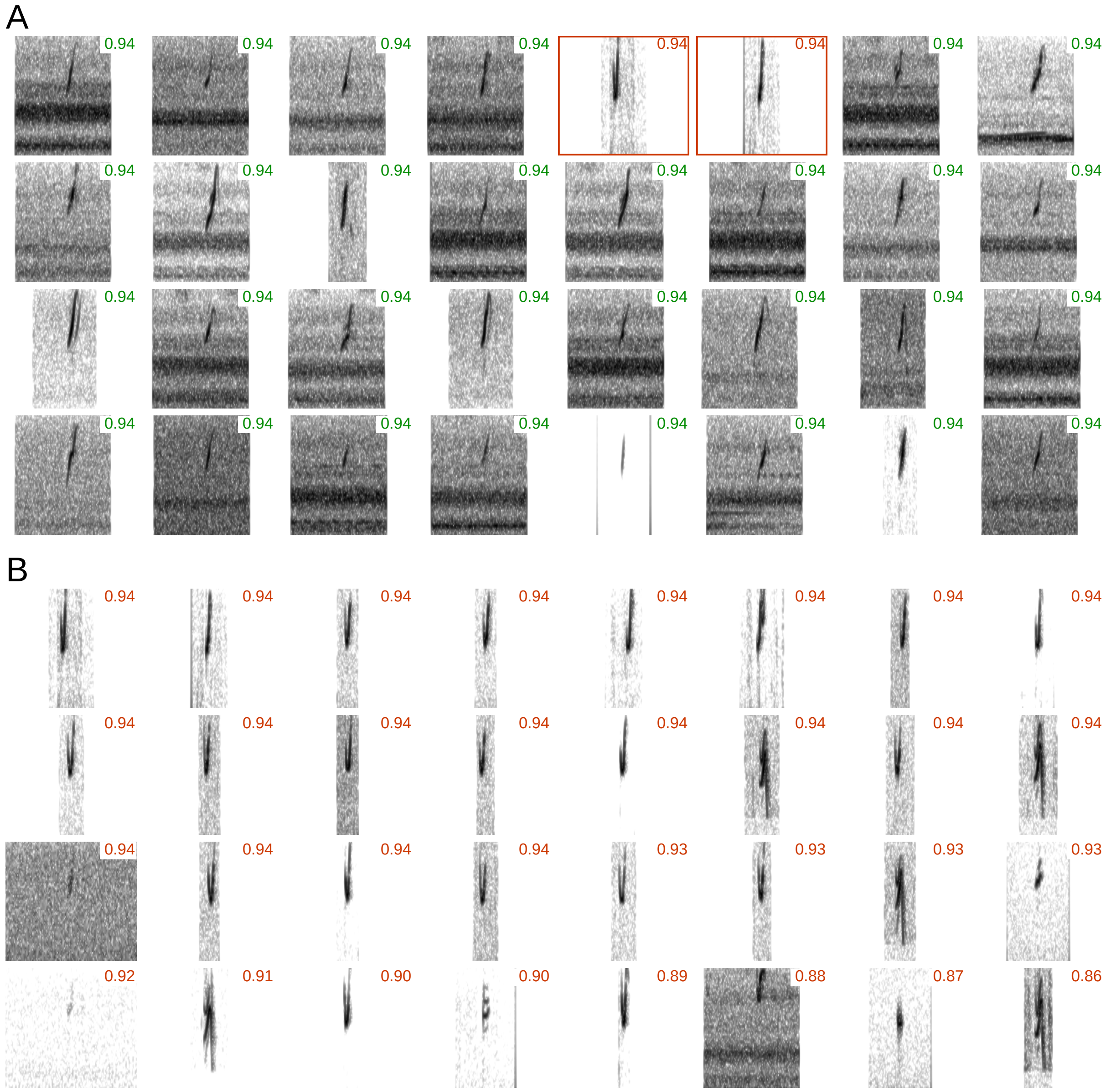


Figure S59: Classification examples for **btbwar** drawn from the test dataset, shown as cropped spectrograms. Numbers in upper-right corners are calibrated probability values returned by the model. (A) Test examples scoring highest for this species. Any incorrect classifications are outlined in red. (B) Incorrectly-classified examples scoring highest for this species (i.e., the most confusing cases for the model). Spectrogram parameters: x-axis: 0-0.3 s; y-axis: 2-10 kHz; window type: Hanning; window length: 200 samples; hop size: 20 samples; dynamic range floor: -60 dB).

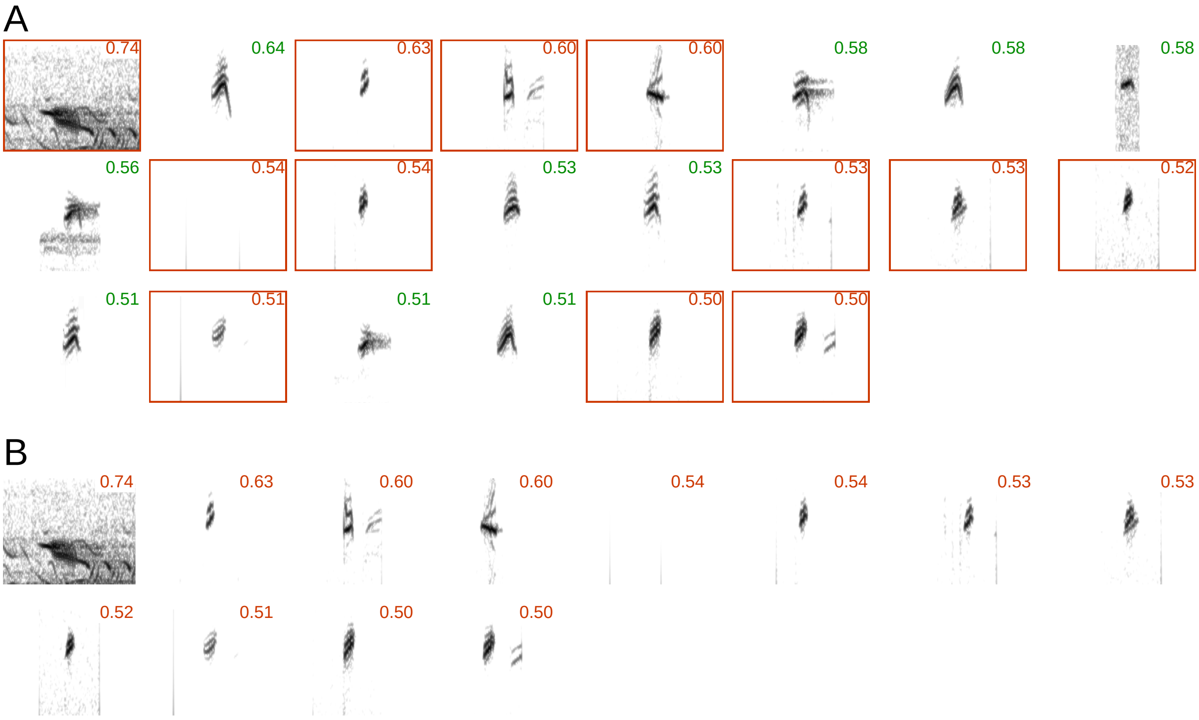


Figure S60: Classification examples for **btnwar** drawn from the test dataset, shown as cropped spectrograms. Numbers in upper-right corners are calibrated probability values returned by the model. (A) Test examples scoring highest for this species. Any incorrect classifications are outlined in red. (B) Incorrectly-classified examples scoring highest for this species (i.e., the most confusing cases for the model). Spectrogram parameters: x-axis: 0-0.3 s; y-axis: 3-11 kHz; window type: Hanning; window length: 200 samples; hop size: 20 samples; dynamic range floor: -50 dB).

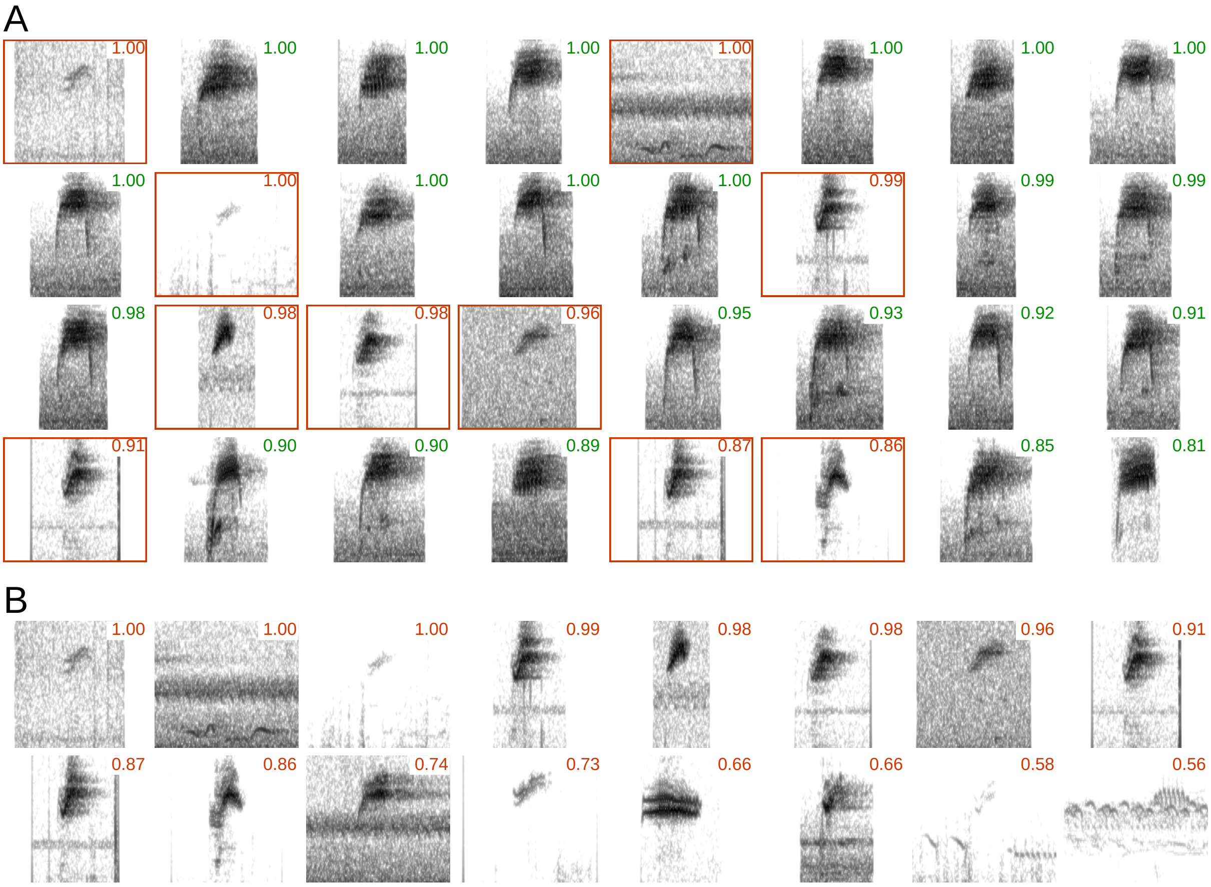


Figure S61: Classification examples for **buwwar** drawn from the test dataset, shown as cropped spectrograms. Numbers in upper-right corners are calibrated probability values returned by the model. (A) Test examples scoring highest for this species. Any incorrect classifications are outlined in red. (B) Incorrectly-classified examples scoring highest for this species (i.e., the most confusing cases for the model). Spectrogram parameters: x-axis: 0-0.3 s; y-axis: 2-10 kHz; window type: Hanning; window length: 200 samples; hop size: 20 samples; dynamic range floor: -60 dB).

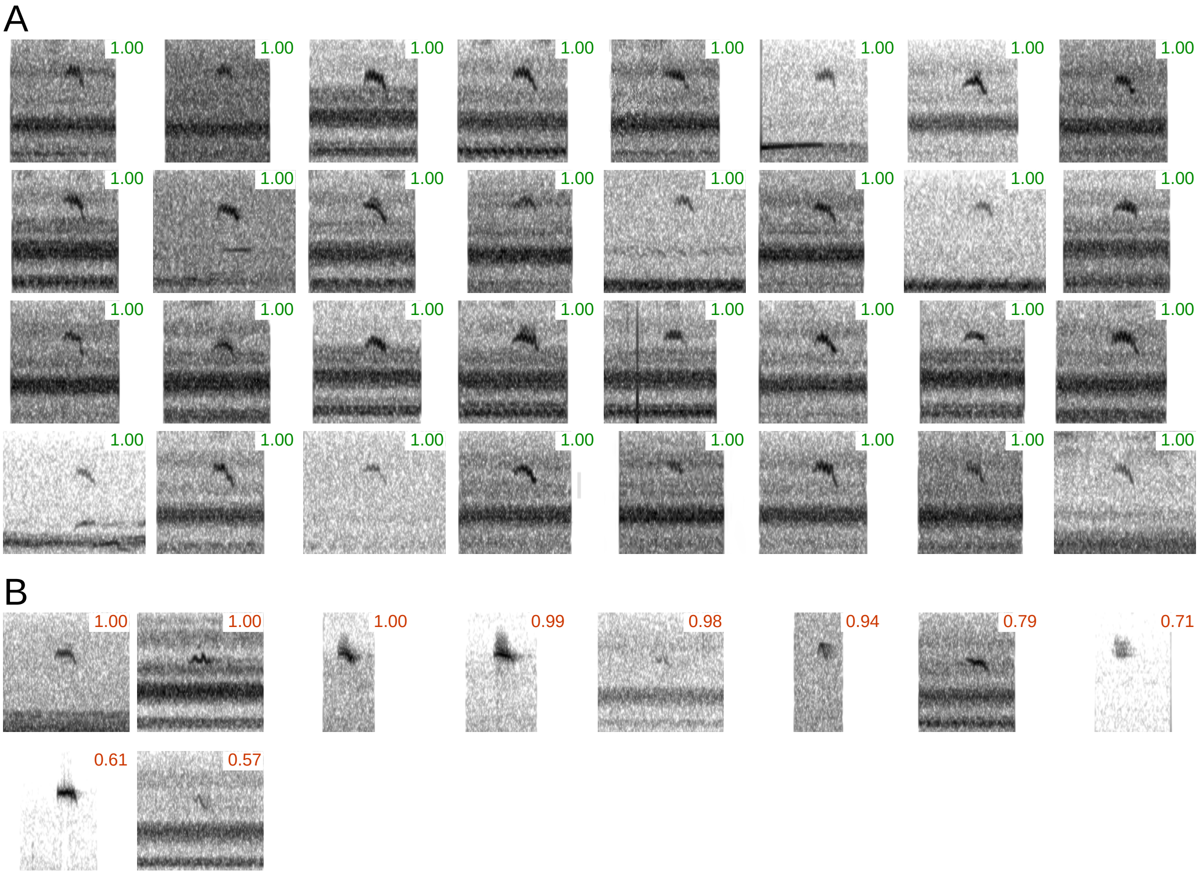


Figure S62: Classification examples for **camwar** drawn from the test dataset, shown as cropped spectrograms. Numbers in upper-right corners are calibrated probability values returned by the model. (A) Test examples scoring highest for this species. Any incorrect classifications are outlined in red. (B) Incorrectly-classified examples scoring highest for this species (i.e., the most confusing cases for the model). Spectrogram parameters: x-axis: 0-0.3 s; y-axis: 2-10 kHz; window type: Hanning; window length: 200 samples; hop size: 20 samples; dynamic range floor: -60 dB).

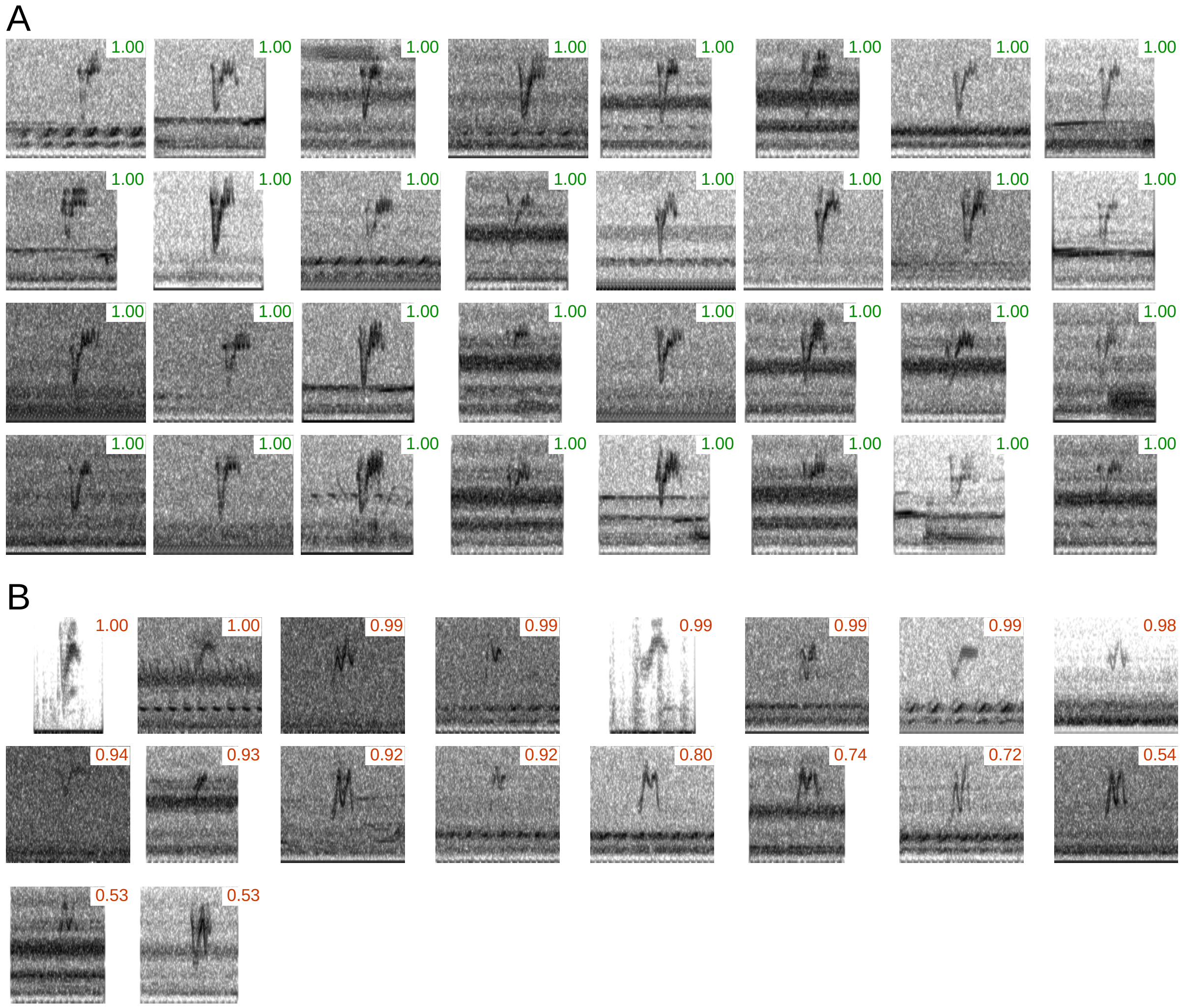


Figure S63: Classification examples for **canwar** drawn from the test dataset, shown as cropped spectrograms. Numbers in upper-right corners are calibrated probability values returned by the model. (A) Test examples scoring highest for this species. Any incorrect classifications are outlined in red. (B) Incorrectly-classified examples scoring highest for this species (i.e., the most confusing cases for the model). Spectrogram parameters: x-axis: 0-0.3 s; y-axis: 0-9 kHz; window type: Hanning; window length: 200 samples; hop size: 20 samples; dynamic range floor: -60 dB).

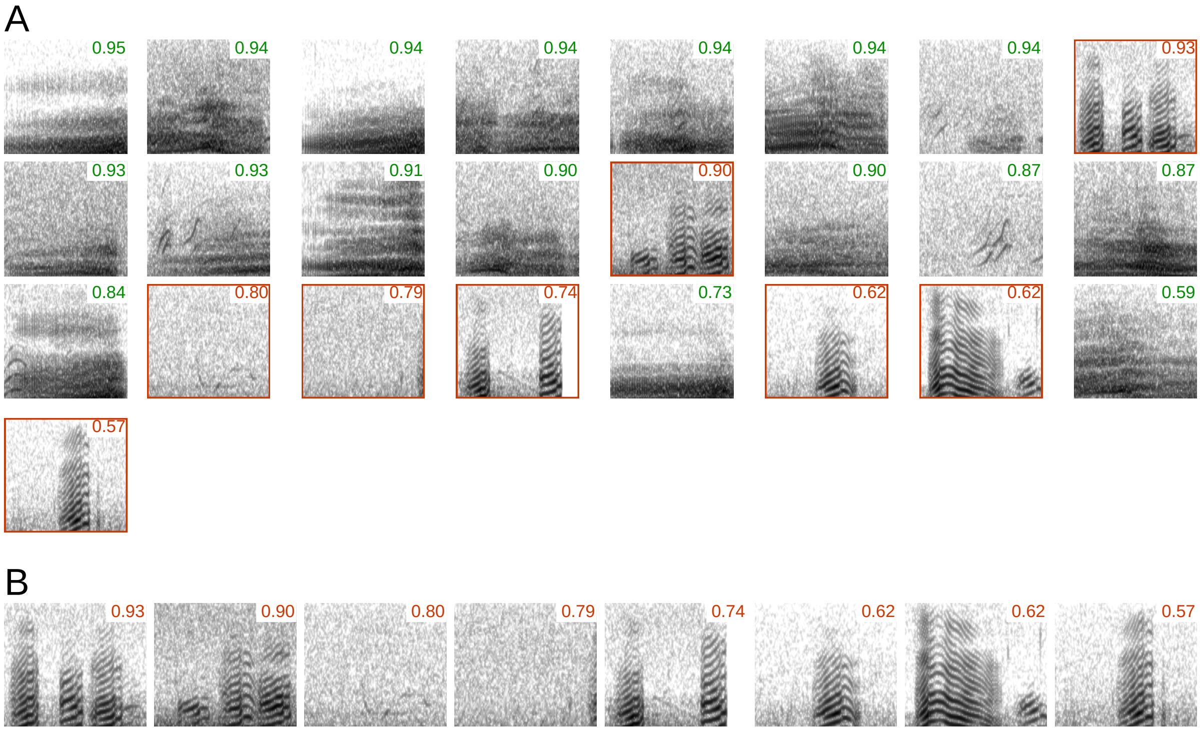


Figure S64: Classification examples for **caster1** drawn from the test dataset, shown as cropped spectrograms. Numbers in upper-right corners are calibrated probability values returned by the model. (A) Test examples scoring highest for this species. Any incorrect classifications are outlined in red. (B) Incorrectly-classified examples scoring highest for this species (i.e., the most confusing cases for the model). Spectrogram parameters: x-axis: 0-0.3 s; y-axis: 2-10 kHz; window type: Hanning; window length: 200 samples; hop size: 20 samples; dynamic range floor: -60 dB).

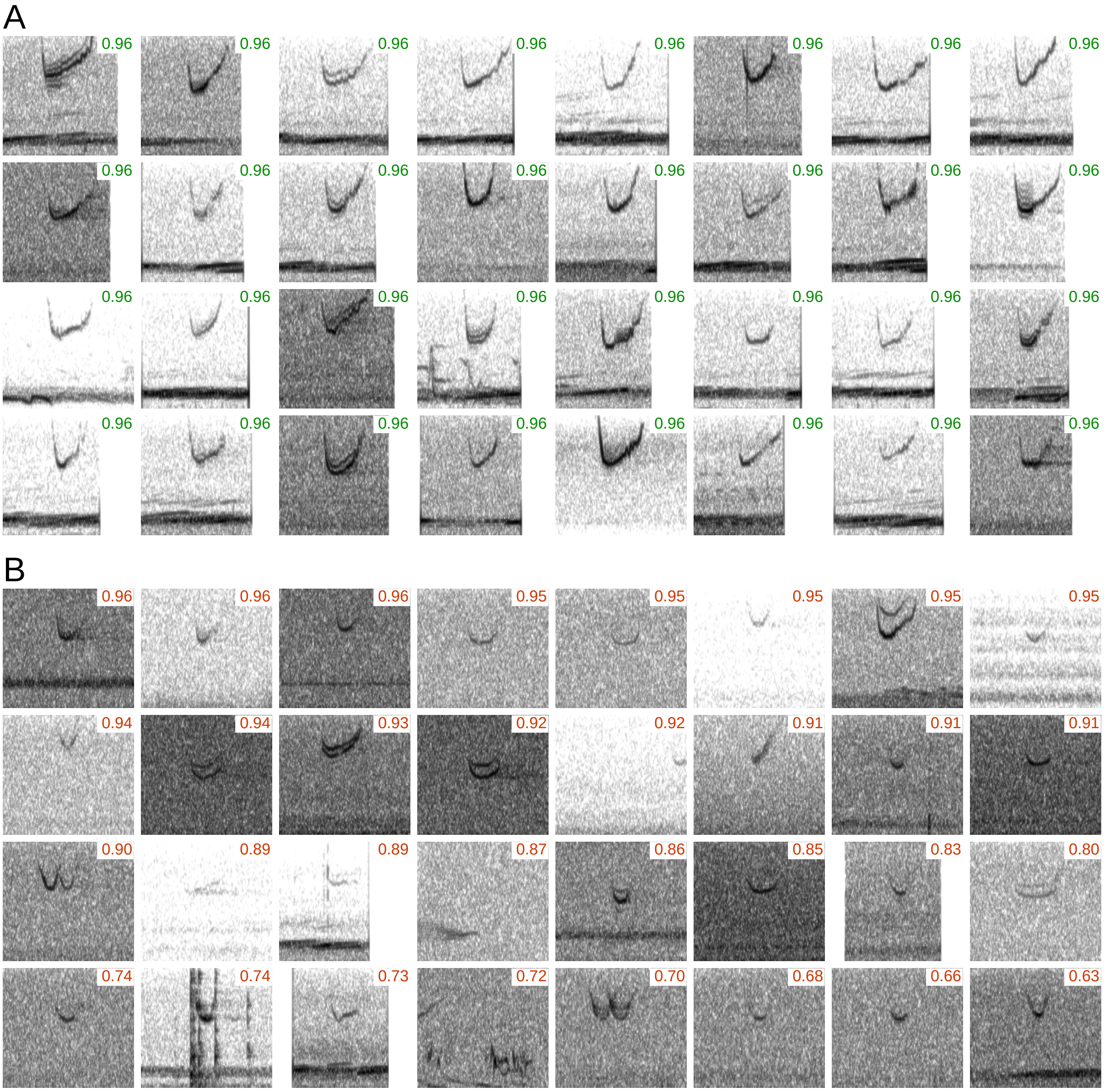


Figure S65: Classification examples for **chispa** drawn from the test dataset, shown as cropped spectrograms. Numbers in upper-right corners are calibrated probability values returned by the model. (A) Test examples scoring highest for this species. Any incorrect classifications are outlined in red. (B) Incorrectly-classified examples scoring highest for this species (i.e., the most confusing cases for the model). Spectrogram parameters: x-axis: 0-0.3 s; y-axis: 2-10 kHz; window type: Hanning; window length: 200 samples; hop size: 20 samples; dynamic range floor: -60 dB).

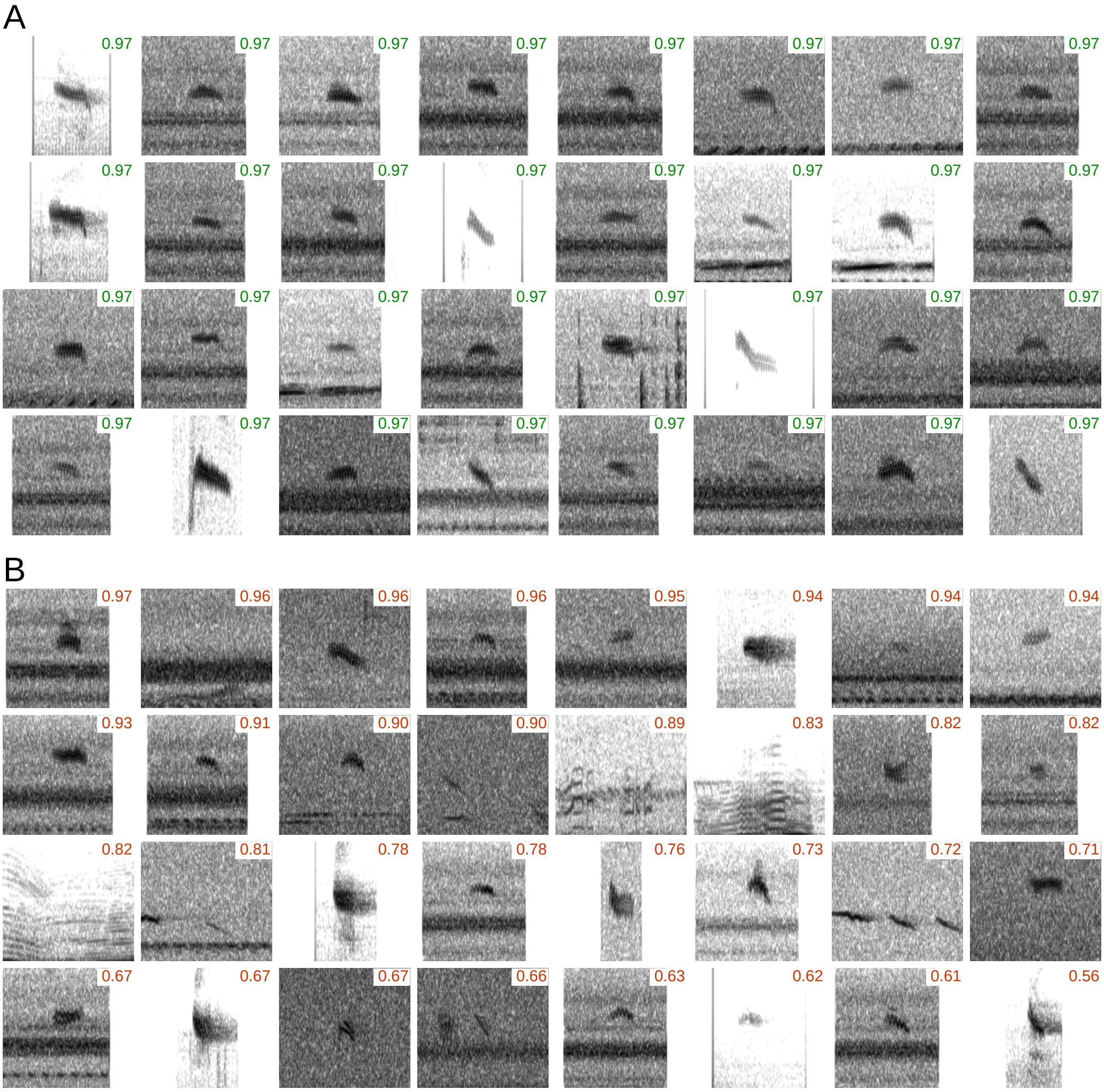


Figure S66: Classification examples for **chswar** drawn from the test dataset, shown as cropped spectrograms. Numbers in upper-right corners are calibrated probability values returned by the model. (A) Test examples scoring highest for this species. Any incorrect classifications are outlined in red. (B) Incorrectly-classified examples scoring highest for this species (i.e., the most confusing cases for the model). Spectrogram parameters: x-axis: 0-0.3 s; y-axis: 2-10 kHz; window type: Hanning; window length: 200 samples; hop size: 20 samples; dynamic range floor: -60 dB).

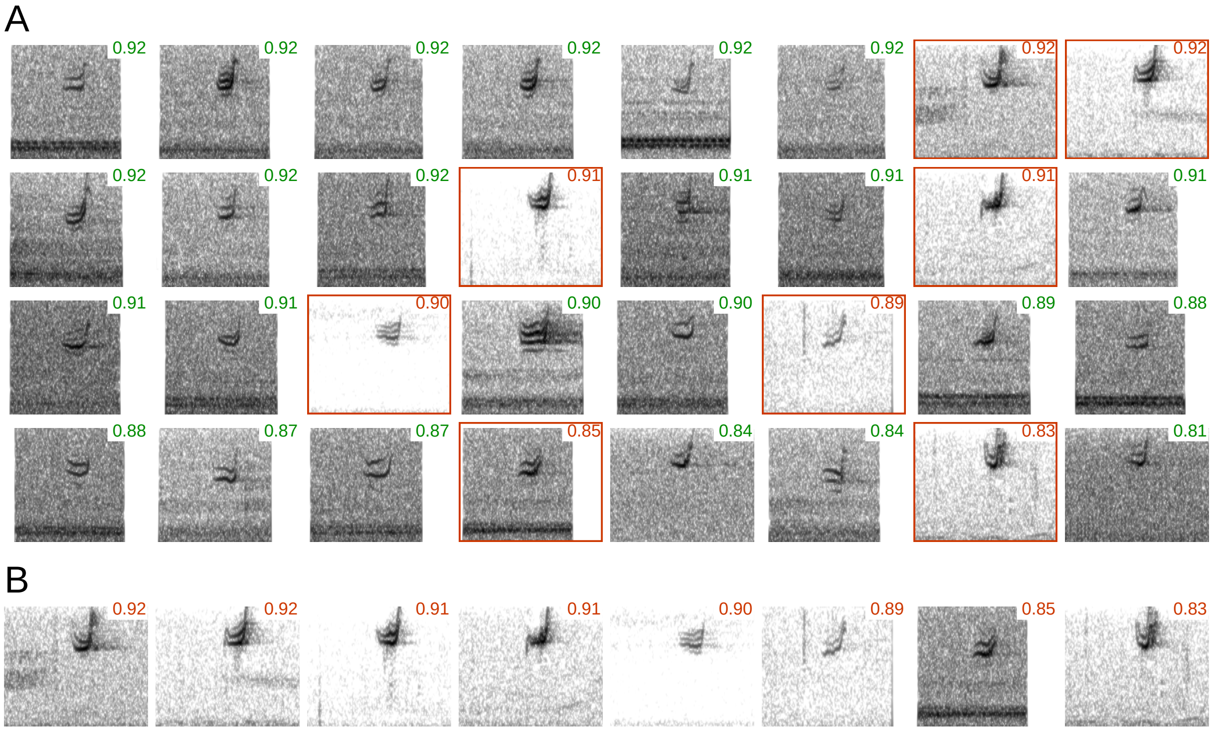


Figure S67: Classification examples for **clcspa** drawn from the test dataset, shown as cropped spectrograms. Numbers in upper-right corners are calibrated probability values returned by the model. (A) Test examples scoring highest for this species. Any incorrect classifications are outlined in red. (B) Incorrectly-classified examples scoring highest for this species (i.e., the most confusing cases for the model). Spectrogram parameters: x-axis: 0-0.3 s; y-axis: 2-11 kHz; window type: Hanning; window length: 200 samples; hop size: 20 samples; dynamic range floor: -60 dB).

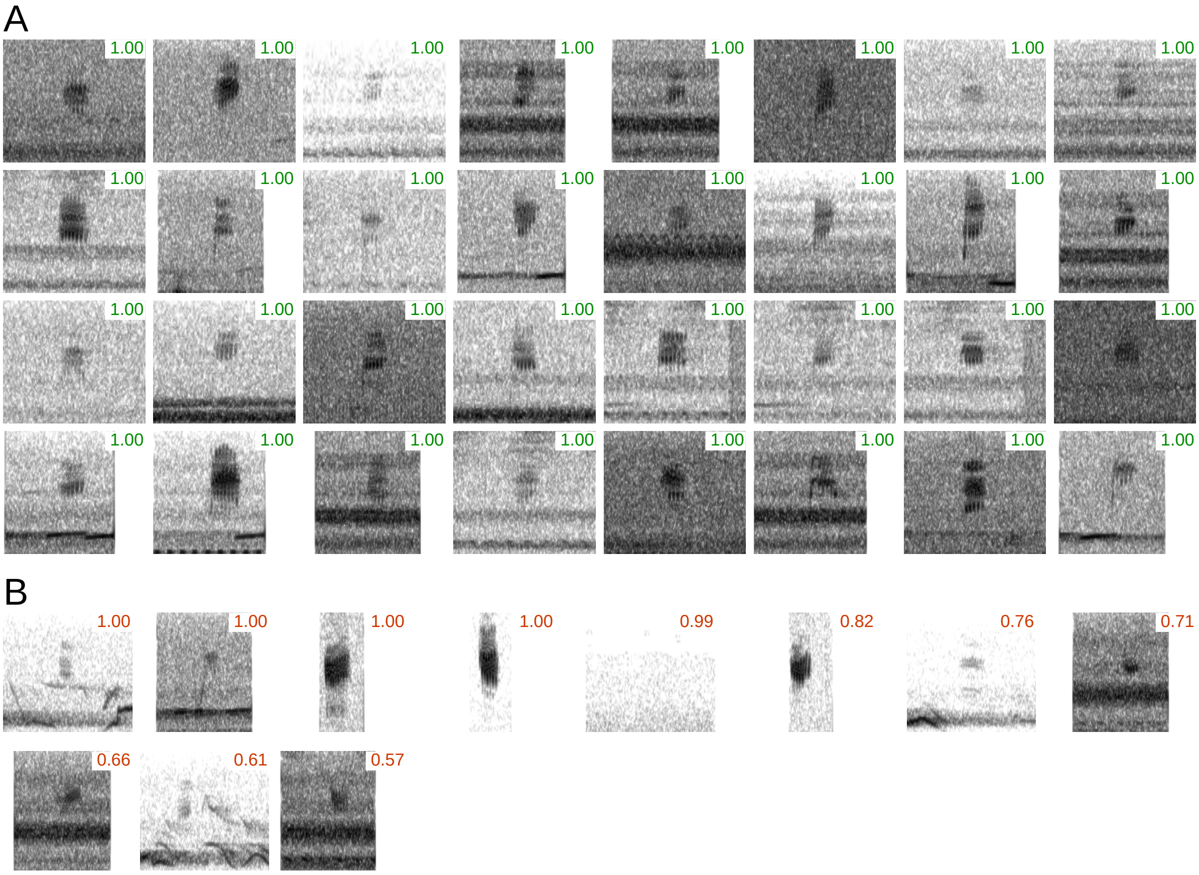


Figure S68: Classification examples for **comyel** drawn from the test dataset, shown as cropped spectrograms. Numbers in upper-right corners are calibrated probability values returned by the model. (A) Test examples scoring highest for this species. Any incorrect classifications are outlined in red. (B) Incorrectly-classified examples scoring highest for this species (i.e., the most confusing cases for the model). Spectrogram parameters: x-axis: 0-0.3 s; y-axis: 2-10 kHz; window type: Hanning; window length: 200 samples; hop size: 20 samples; dynamic range floor: -60 dB).

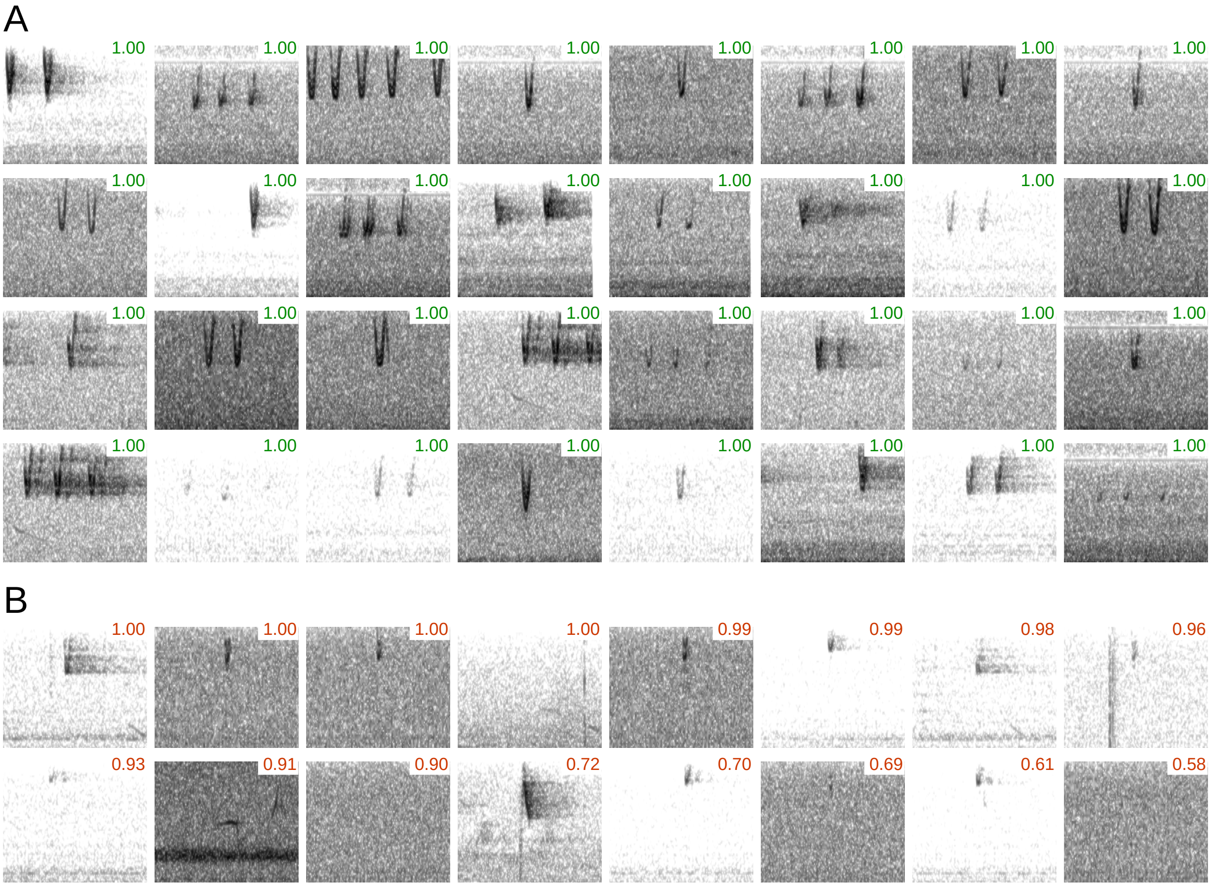


Figure S69: Classification examples for **daejun** drawn from the test dataset, shown as cropped spectrograms. Numbers in upper-right corners are calibrated probability values returned by the model. (A) Test examples scoring highest for this species. Any incorrect classifications are outlined in red. (B) Incorrectly-classified examples scoring highest for this species (i.e., the most confusing cases for the model). Spectrogram parameters: x-axis: 0-0.3 s; y-axis: 2-11 kHz; window type: Hanning; window length: 200 samples; hop size: 20 samples; dynamic range floor: -60 dB).

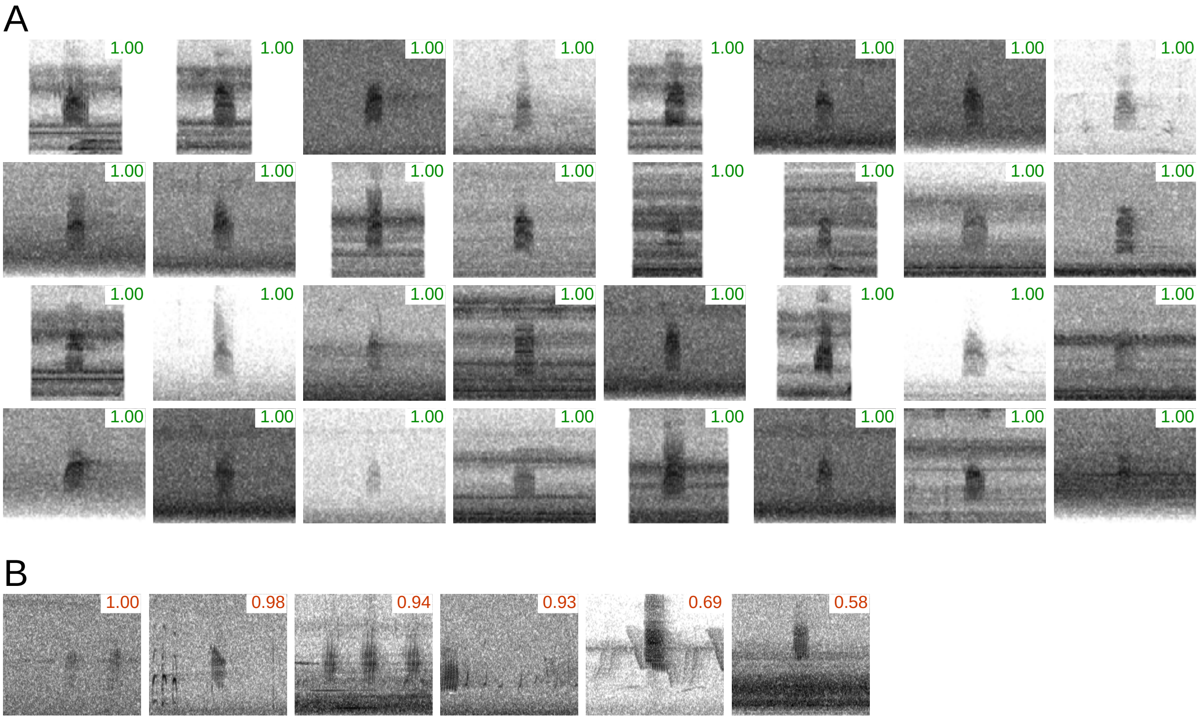


Figure S70: Classification examples for **dickci** drawn from the test dataset, shown as cropped spectrograms. Numbers in upper-right corners are calibrated probability values returned by the model. (A) Test examples scoring highest for this species. Any incorrect classifications are outlined in red. (B) Incorrectly-classified examples scoring highest for this species (i.e., the most confusing cases for the model). Spectrogram parameters: x-axis: 0-0.8 s; y-axis: 1-8 kHz; window type: Hanning; window length: 400 samples; hop size: 30 samples; dynamic range floor: -60 dB).

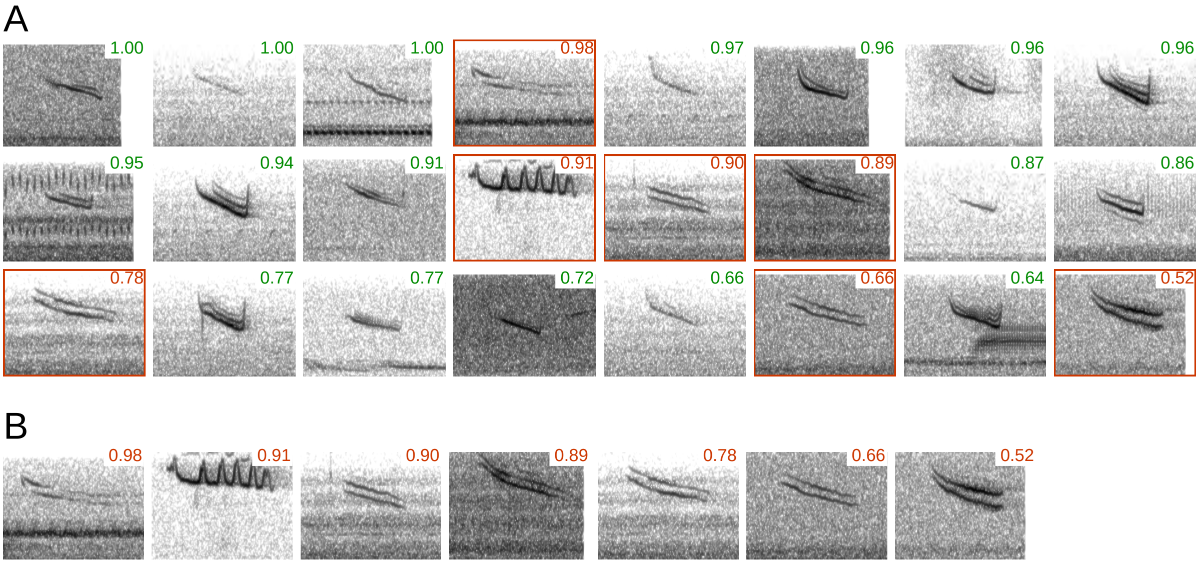


Figure S71: Classification examples for **fiespa** drawn from the test dataset, shown as cropped spectrograms. Numbers in upper-right corners are calibrated probability values returned by the model. (A) Test examples scoring highest for this species. Any incorrect classifications are outlined in red. (B) Incorrectly-classified examples scoring highest for this species (i.e., the most confusing cases for the model). Spectrogram parameters: x-axis: 0-0.3 s; y-axis: 2-11 kHz; window type: Hanning; window length: 200 samples; hop size: 20 samples; dynamic range floor: -60 dB).

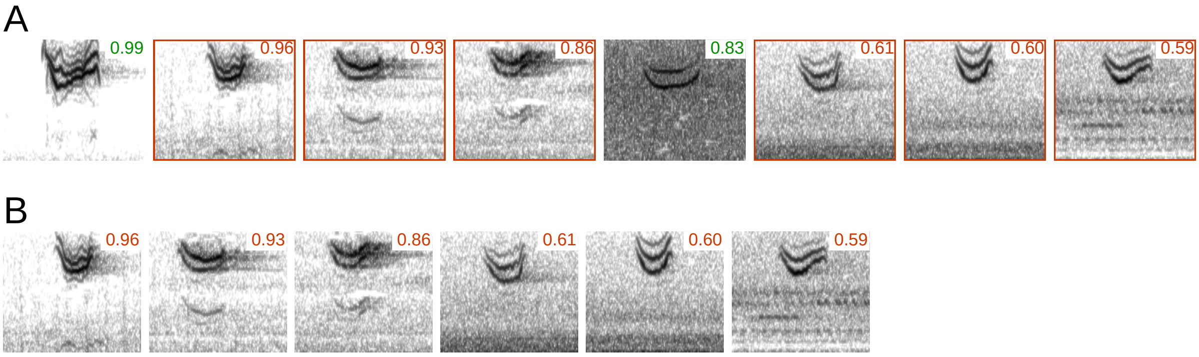


Figure S72: Classification examples for **foxspa** drawn from the test dataset, shown as cropped spectrograms. Numbers in upper-right corners are calibrated probability values returned by the model. (A) Test examples scoring highest for this species. Any incorrect classifications are outlined in red. (B) Incorrectly-classified examples scoring highest for this species (i.e., the most confusing cases for the model). Spectrogram parameters: x-axis: 0-0.3 s; y-axis: 2-10 kHz; window type: Hanning; window length: 200 samples; hop size: 20 samples; dynamic range floor: -60 dB).

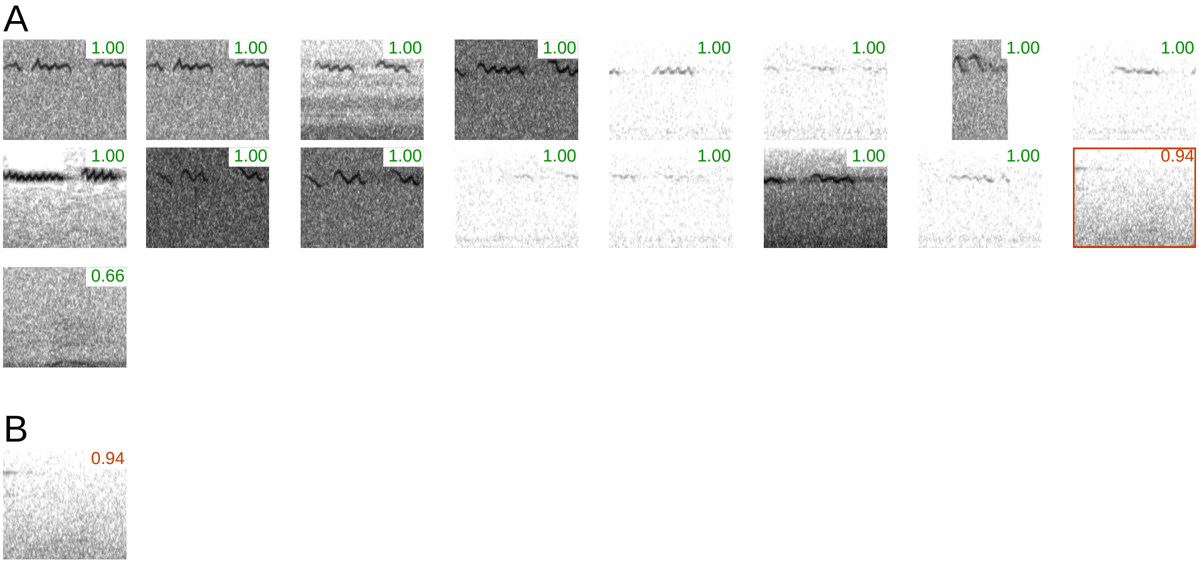


Figure S73: Classification examples for **gockin** drawn from the test dataset, shown as cropped spectrograms. Numbers in upper-right corners are calibrated probability values returned by the model. (A) Test examples scoring highest for this species. Any incorrect classifications are outlined in red. (B) Incorrectly-classified examples scoring highest for this species (i.e., the most confusing cases for the model). Spectrogram parameters: x-axis: 0-0.3 s; y-axis: 2-10 kHz; window type: Hanning; window length: 200 samples; hop size: 20 samples; dynamic range floor: -60 dB).

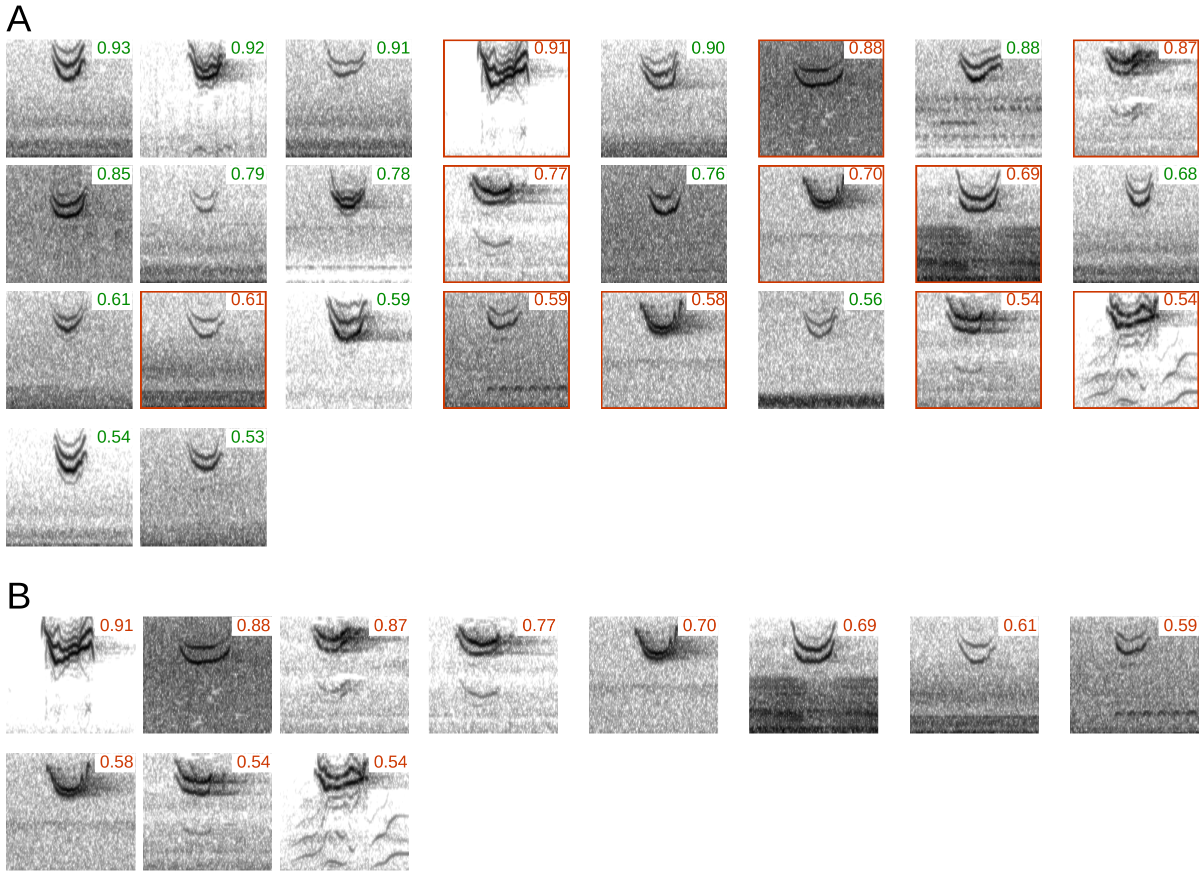


Figure S74: Classification examples for **gocspa** drawn from the test dataset, shown as cropped spectrograms. Numbers in upper-right corners are calibrated probability values returned by the model. (A) Test examples scoring highest for this species. Any incorrect classifications are outlined in red. (B) Incorrectly-classified examples scoring highest for this species (i.e., the most confusing cases for the model). Spectrogram parameters: x-axis: 0-0.3 s; y-axis: 2-10 kHz; window type: Hanning; window length: 200 samples; hop size: 20 samples; dynamic range floor: -60 dB).

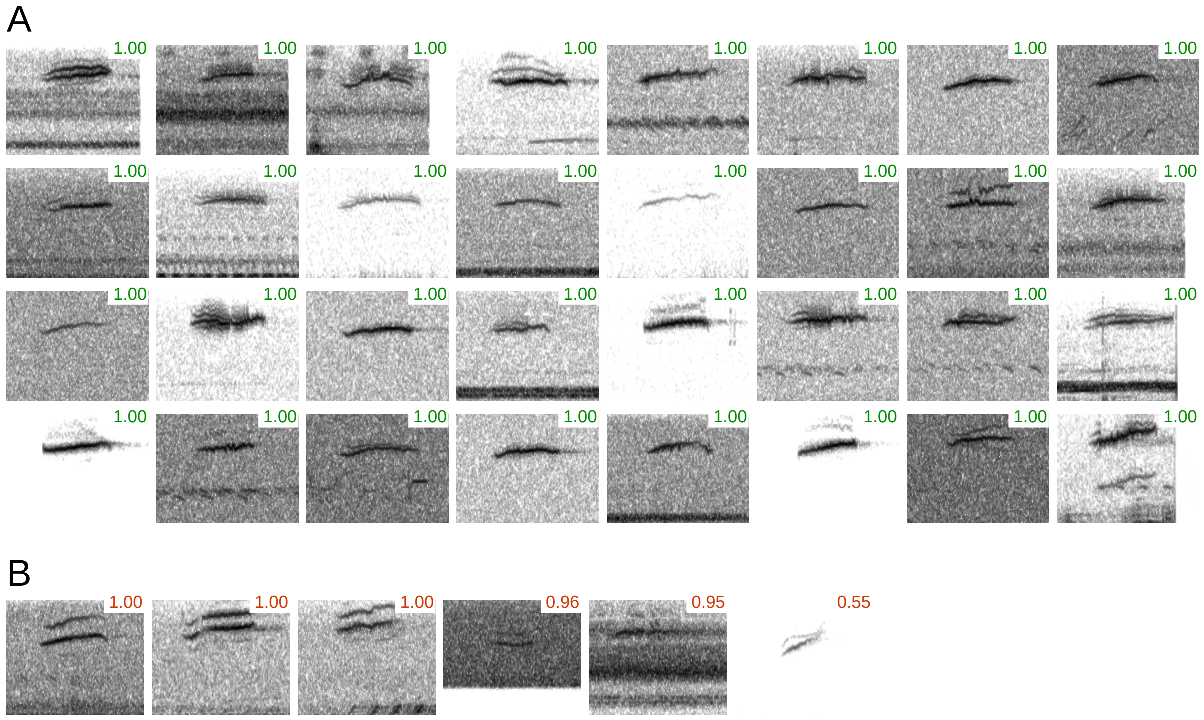


Figure S75: Classification examples for **graspa** drawn from the test dataset, shown as cropped spectrograms. Numbers in upper-right corners are calibrated probability values returned by the model. (A) Test examples scoring highest for this species. Any incorrect classifications are outlined in red. (B) Incorrectly-classified examples scoring highest for this species (i.e., the most confusing cases for the model). Spectrogram parameters: x-axis: 0-0.3 s; y-axis: 2-11 kHz; window type: Hanning; window length: 200 samples; hop size: 20 samples; dynamic range floor: -60 dB).

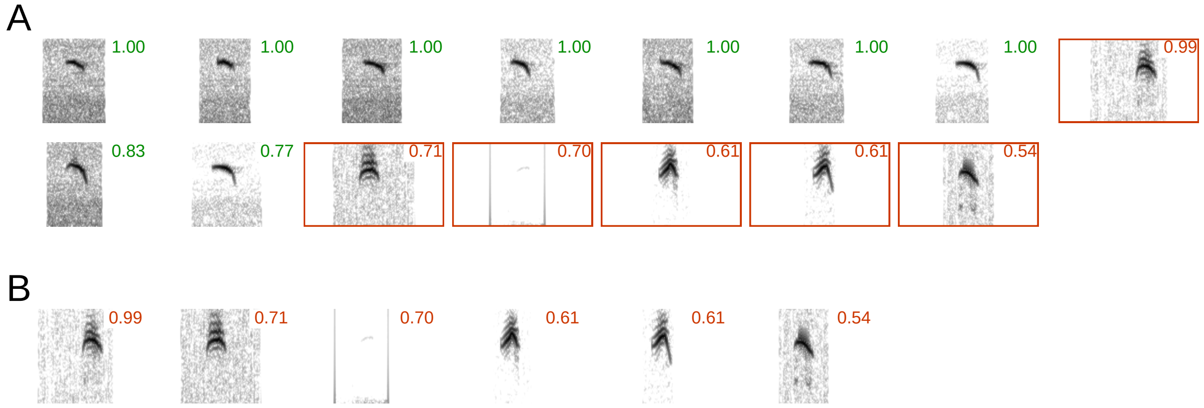


Figure S76: Classification examples for **grawar** drawn from the test dataset, shown as cropped spectrograms. Numbers in upper-right corners are calibrated probability values returned by the model. (A) Test examples scoring highest for this species. Any incorrect classifications are outlined in red. (B) Incorrectly-classified examples scoring highest for this species (i.e., the most confusing cases for the model). Spectrogram parameters: x-axis: 0-0.3 s; y-axis: 2-10 kHz; window type: Hanning; window length: 200 samples; hop size: 20 samples; dynamic range floor: -60 dB).

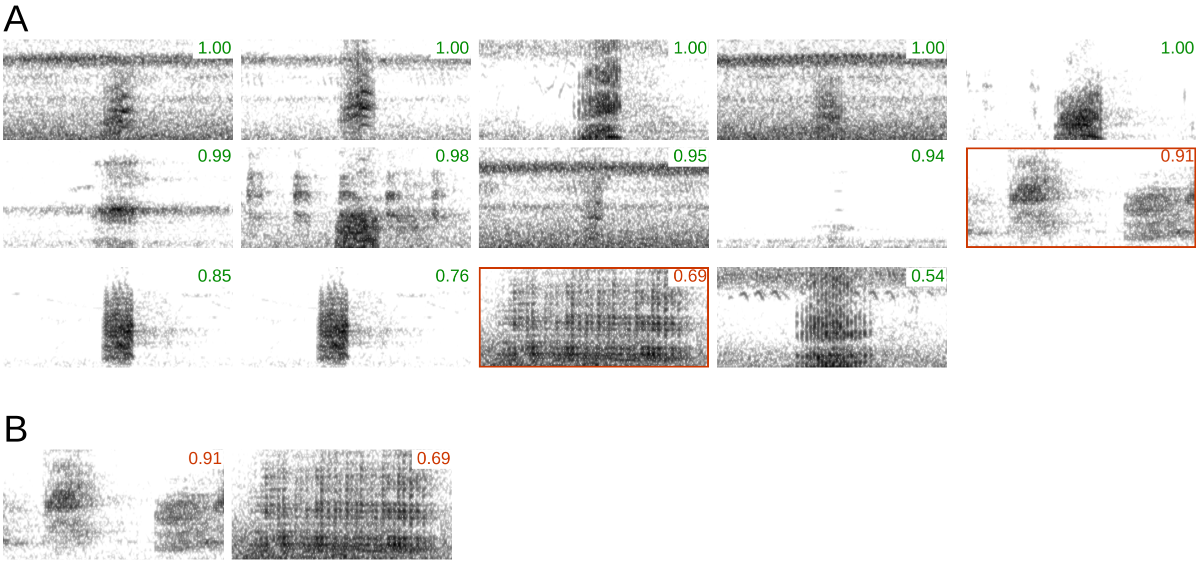


Figure S77: Classification examples for **grbher3** drawn from the test dataset, shown as cropped spectrograms. Numbers in upper-right corners are calibrated probability values returned by the model. (A) Test examples scoring highest for this species. Any incorrect classifications are outlined in red. (B) Incorrectly-classified examples scoring highest for this species (i.e., the most confusing cases for the model). Spectrogram parameters: x-axis: 0-0.8 s; y-axis: 1-5 kHz; window type: Hanning; window length: 400 samples; hop size: 30 samples; dynamic range floor: -45 dB).

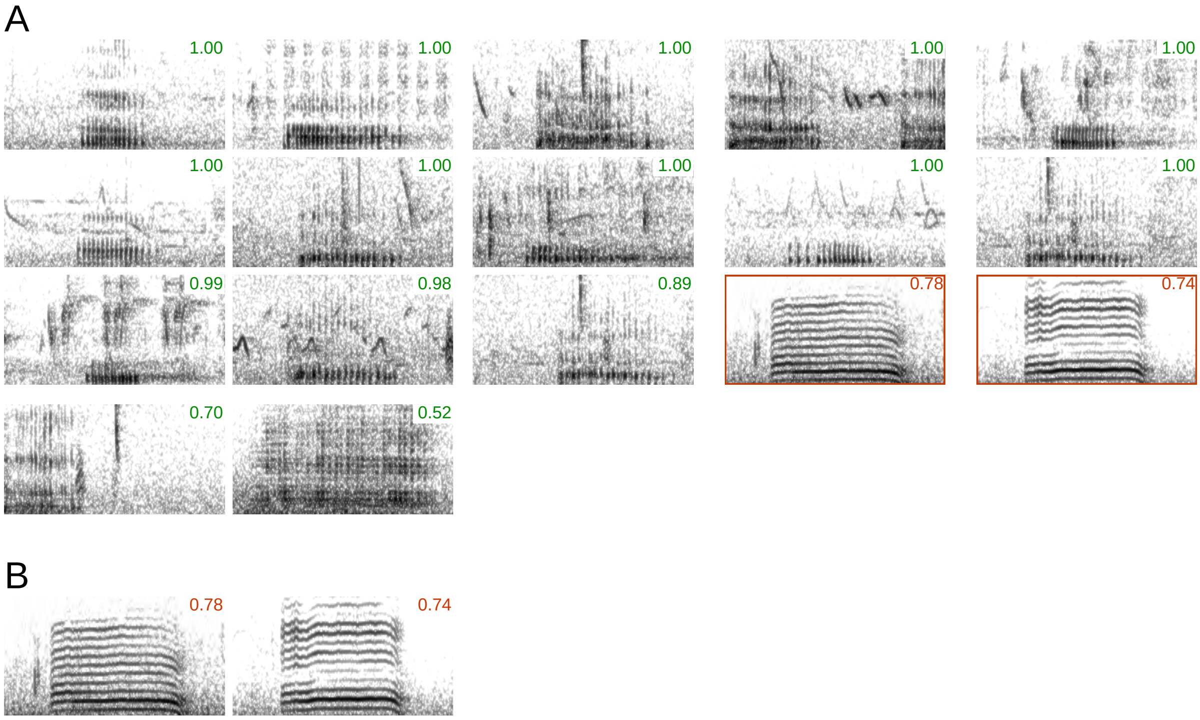


Figure S78: Classification examples for **greegr** drawn from the test dataset, shown as cropped spectrograms. Numbers in upper-right corners are calibrated probability values returned by the model. (A) Test examples scoring highest for this species. Any incorrect classifications are outlined in red. (B) Incorrectly-classified examples scoring highest for this species (i.e., the most confusing cases for the model). Spectrogram parameters: x-axis: 0-0.8 s; y-axis: 1-5 kHz; window type: Hanning; window length: 400 samples; hop size: 30 samples; dynamic range floor: -45 dB).

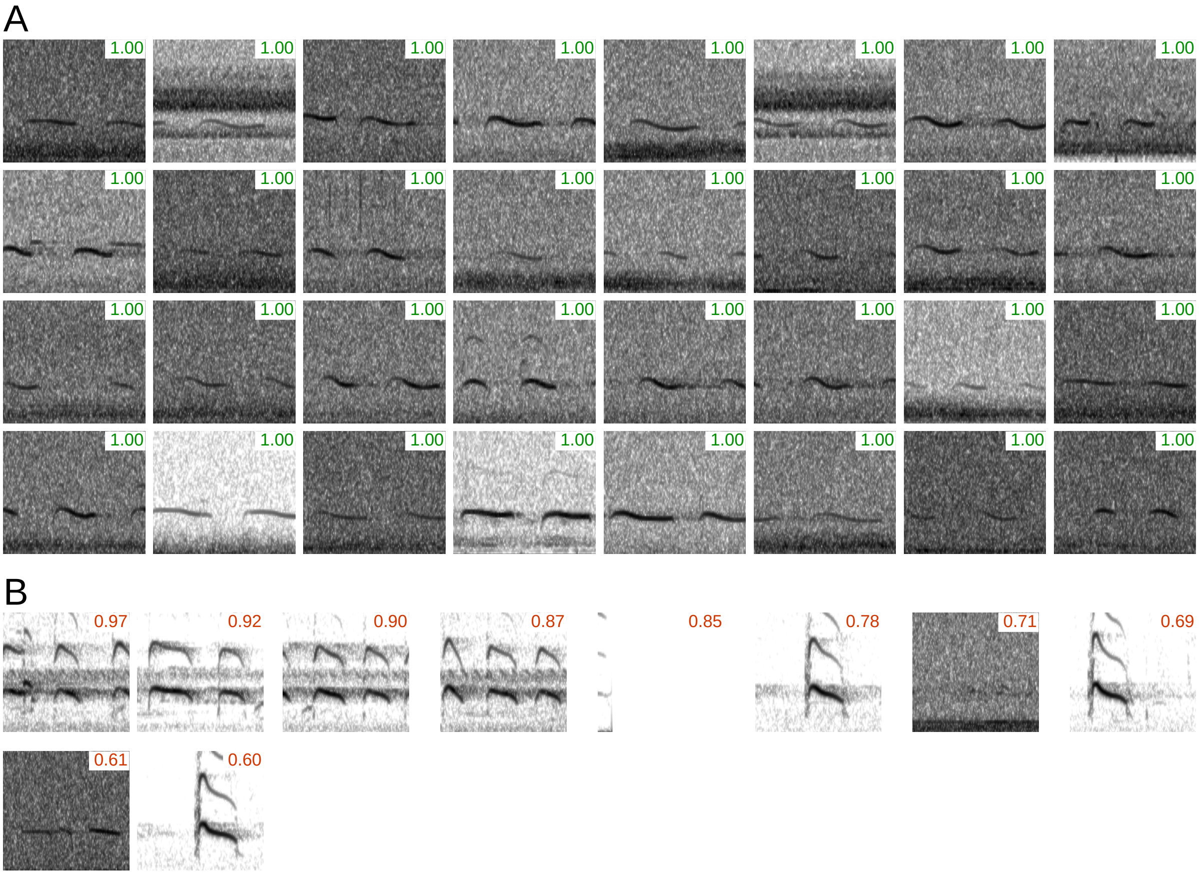


Figure S79: Classification examples for **greyel** drawn from the test dataset, shown as cropped spectrograms. Numbers in upper-right corners are calibrated probability values returned by the model. (A) Test examples scoring highest for this species. Any incorrect classifications are outlined in red. (B) Incorrectly-classified examples scoring highest for this species (i.e., the most confusing cases for the model). Spectrogram parameters: x-axis: 0-0.3 s; y-axis: 0.5-8 kHz; window type: Hanning; window length: 200 samples; hop size: 20 samples; dynamic range floor: -60 dB).

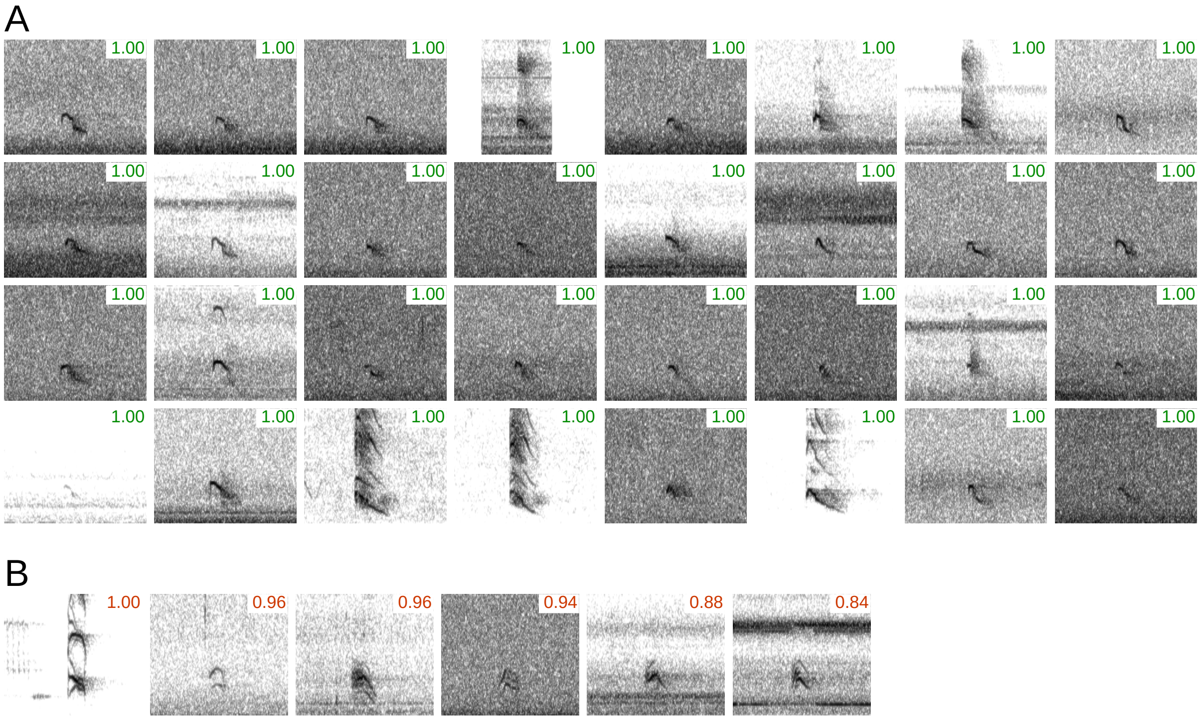


Figure S80: Classification examples for **grnher** drawn from the test dataset, shown as cropped spectrograms. Numbers in upper-right corners are calibrated probability values returned by the model. (A) Test examples scoring highest for this species. Any incorrect classifications are outlined in red. (B) Incorrectly-classified examples scoring highest for this species (i.e., the most confusing cases for the model). Spectrogram parameters: x-axis: 0-0.8 s; y-axis: 1-6 kHz; window type: Hanning; window length: 400 samples; hop size: 30 samples; dynamic range floor: -50 dB).

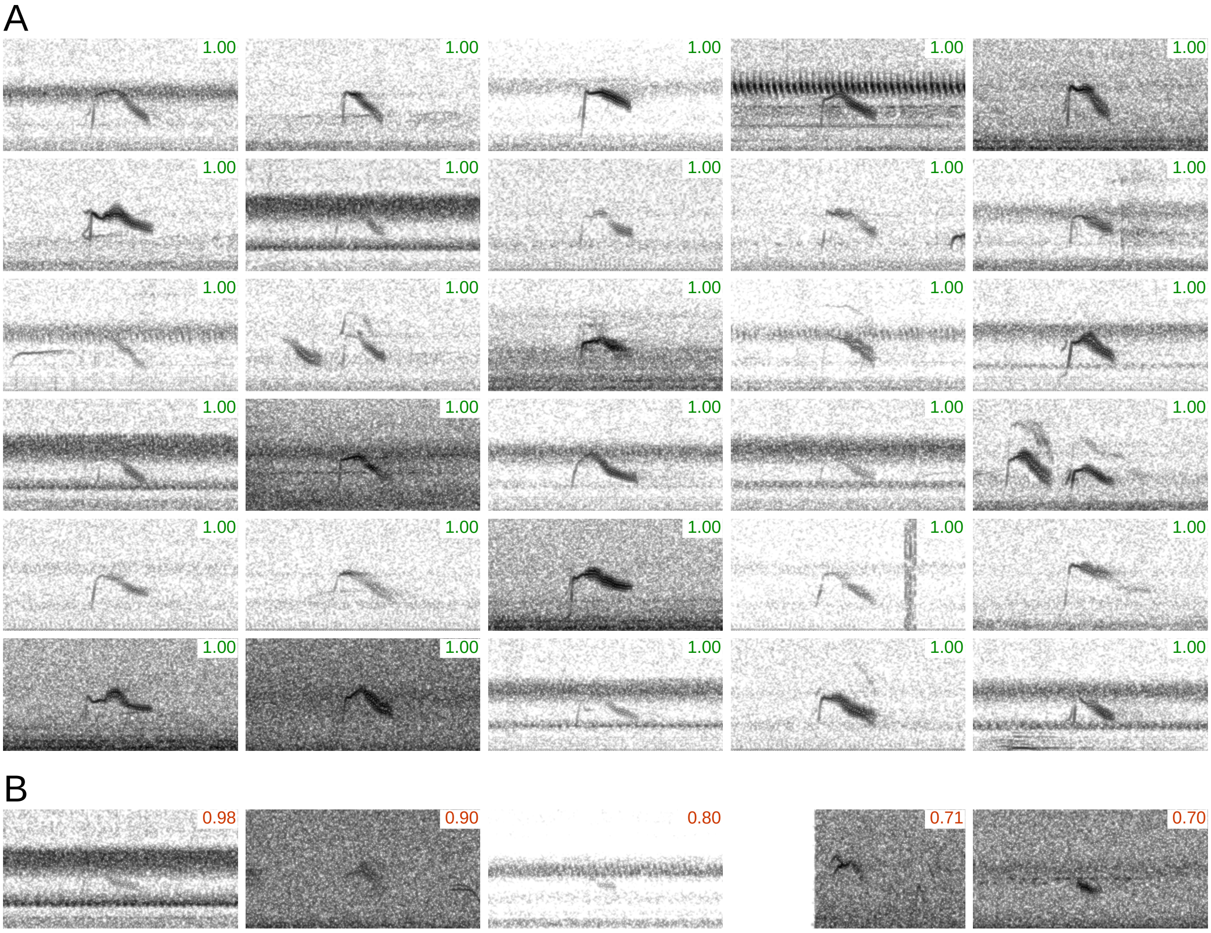


Figure S81: Classification examples for **gycthr** drawn from the test dataset, shown as cropped spectrograms. Numbers in upper-right corners are calibrated probability values returned by the model. (A) Test examples scoring highest for this species. Any incorrect classifications are outlined in red. (B) Incorrectly-classified examples scoring highest for this species (i.e., the most confusing cases for the model). Spectrogram parameters: x-axis: 0-0.8 s; y-axis: 1-7 kHz; window type: Hanning; window length: 400 samples; hop size: 30 samples; dynamic range floor: -50 dB).

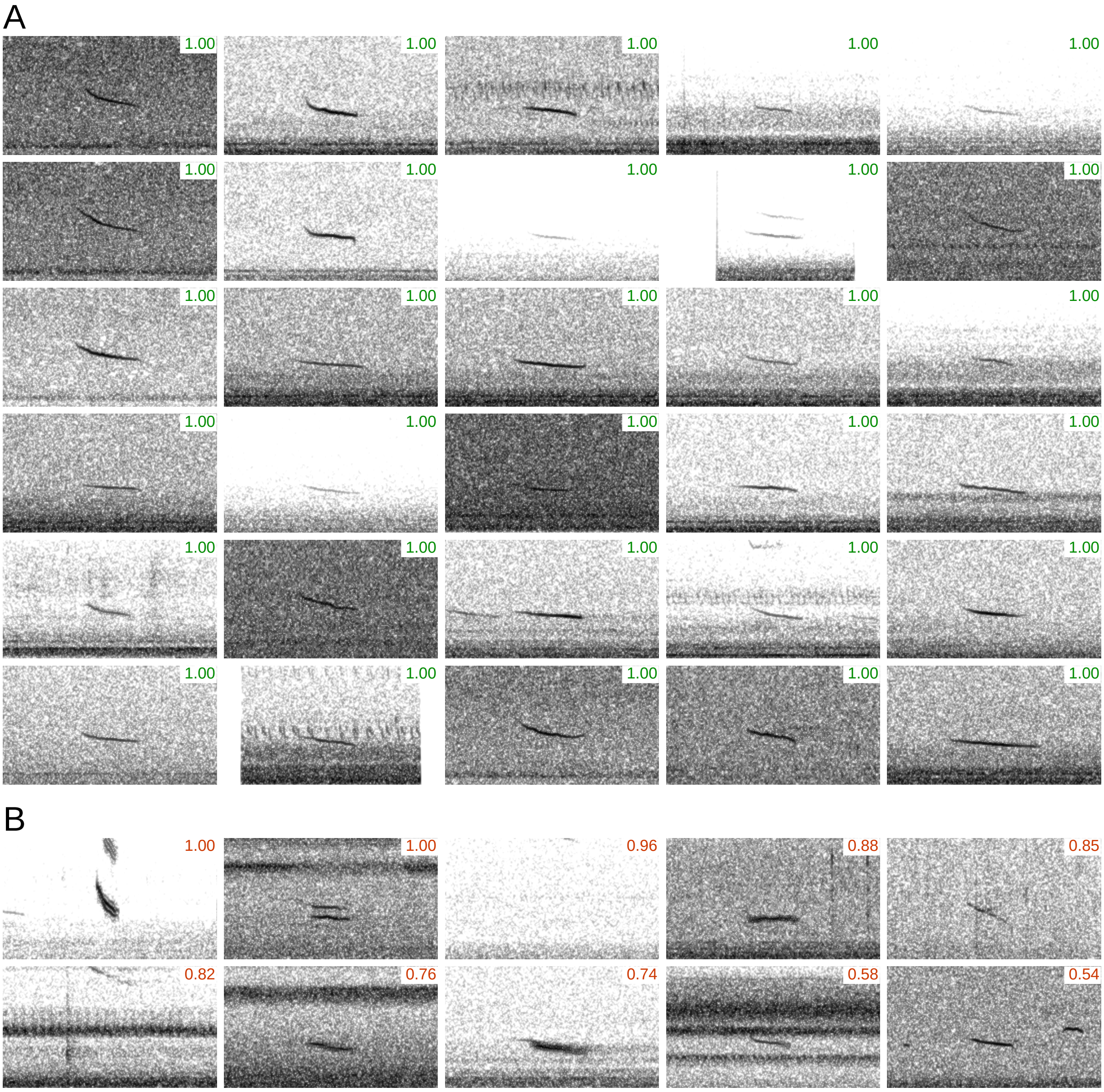


Figure S82: Classification examples for **herthr** drawn from the test dataset, shown as cropped spectrograms. Numbers in upper-right corners are calibrated probability values returned by the model. (A) Test examples scoring highest for this species. Any incorrect classifications are outlined in red. (B) Incorrectly-classified examples scoring highest for this species (i.e., the most confusing cases for the model). Spectrogram parameters: x-axis: 0-0.8 s; y-axis: 1-7 kHz; window type: Hanning; window length: 400 samples; hop size: 30 samples; dynamic range floor: -45 dB).

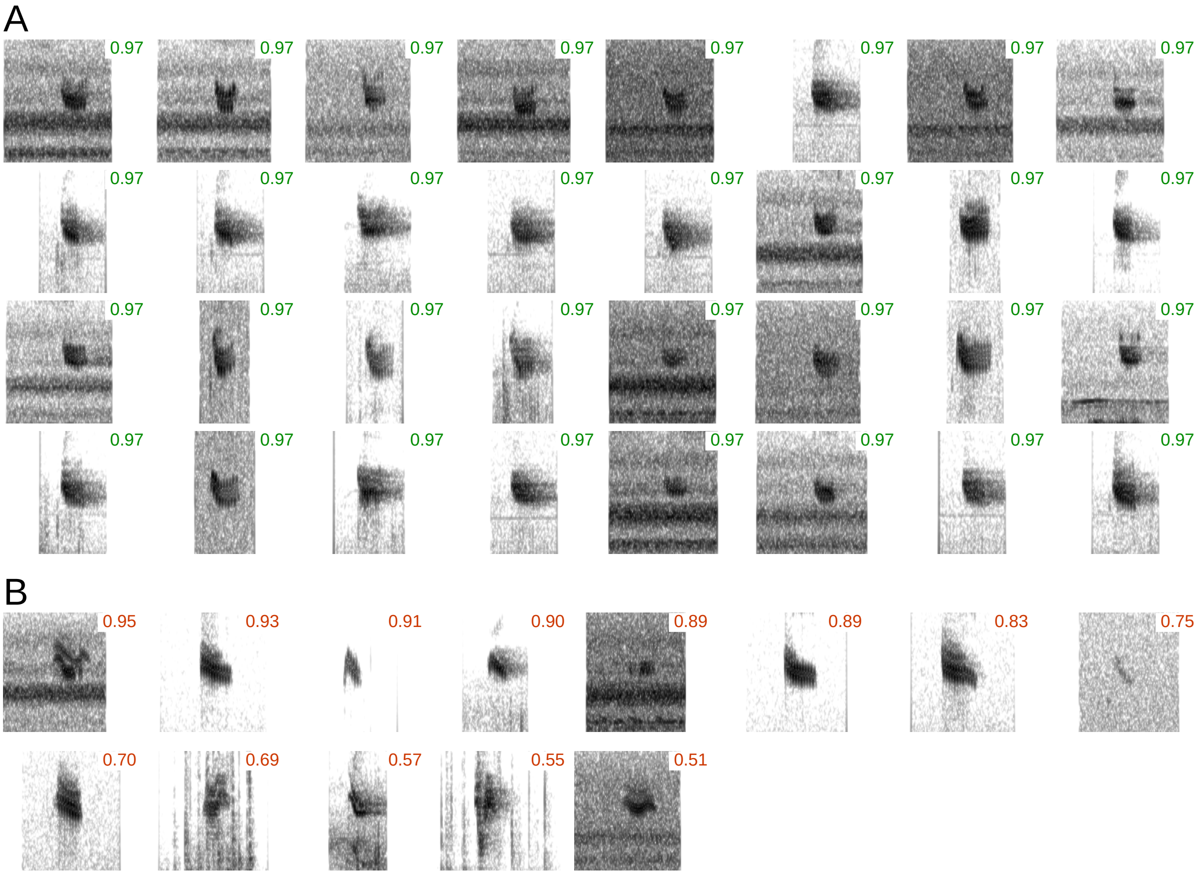


Figure S83: Classification examples for **hoowar** drawn from the test dataset, shown as cropped spectrograms. Numbers in upper-right corners are calibrated probability values returned by the model. (A) Test examples scoring highest for this species. Any incorrect classifications are outlined in red. (B) Incorrectly-classified examples scoring highest for this species (i.e., the most confusing cases for the model). Spectrogram parameters: x-axis: 0-0.3 s; y-axis: 2-10 kHz; window type: Hanning; window length: 200 samples; hop size: 20 samples; dynamic range floor: -60 dB).

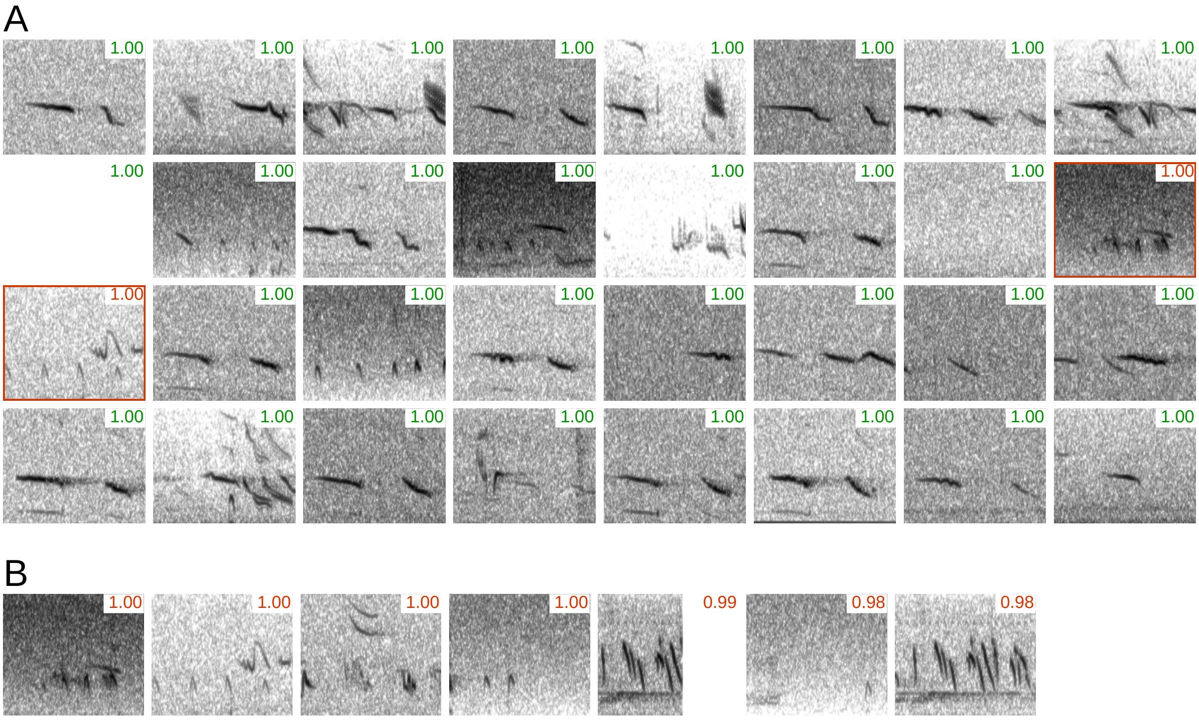


Figure S84: Classification examples for **horlar** drawn from the test dataset, shown as cropped spectrograms. Numbers in upper-right corners are calibrated probability values returned by the model. (A) Test examples scoring highest for this species. Any incorrect classifications are outlined in red. (B) Incorrectly-classified examples scoring highest for this species (i.e., the most confusing cases for the model). Spectrogram parameters: x-axis: 0-0.3 s; y-axis: 2-10 kHz; window type: Hanning; window length: 200 samples; hop size: 20 samples; dynamic range floor: -60 dB).

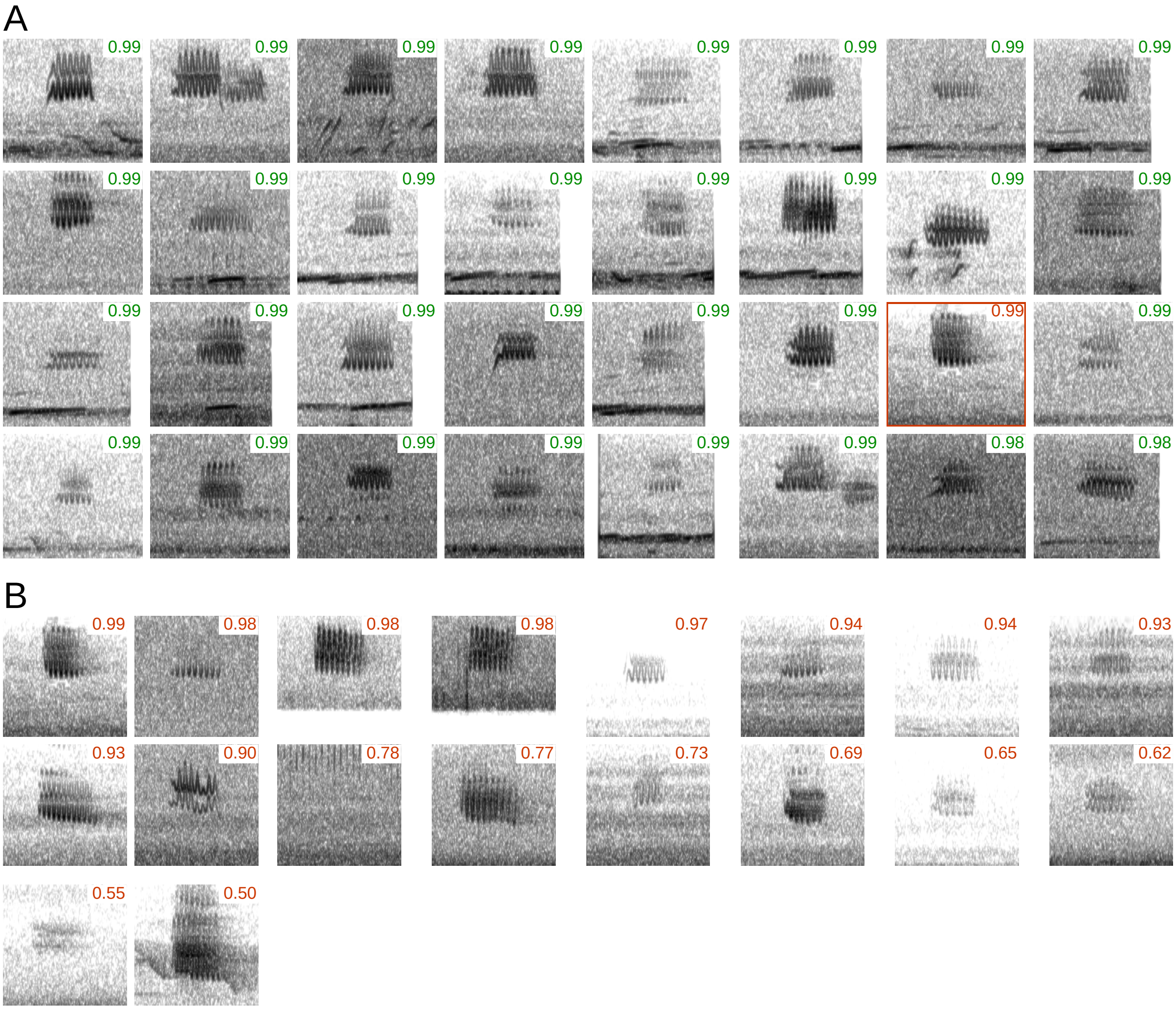


Figure S85: Classification examples for **indbun** drawn from the test dataset, shown as cropped spectrograms. Numbers in upper-right corners are calibrated probability values returned by the model. (A) Test examples scoring highest for this species. Any incorrect classifications are outlined in red. (B) Incorrectly-classified examples scoring highest for this species (i.e., the most confusing cases for the model). Spectrogram parameters: x-axis: 0-0.3 s; y-axis: 2-10 kHz; window type: Hanning; window length: 200 samples; hop size: 20 samples; dynamic range floor: -60 dB).

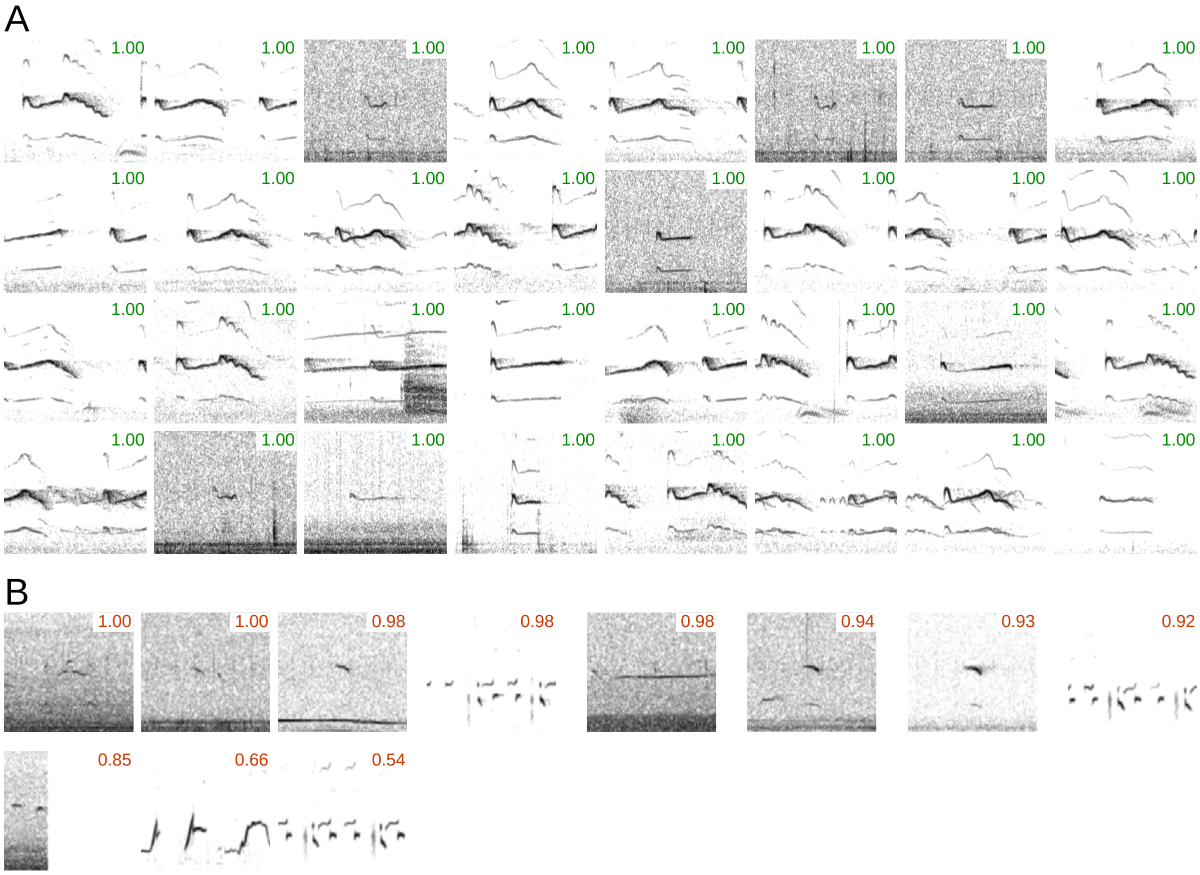


Figure S86: Classification examples for **killde** drawn from the test dataset, shown as cropped spectrograms. Numbers in upper-right corners are calibrated probability values returned by the model. (A) Test examples scoring highest for this species. Any incorrect classifications are outlined in red. (B) Incorrectly-classified examples scoring highest for this species (i.e., the most confusing cases for the model). Spectrogram parameters: x-axis: 0-0.8 s; y-axis: 1-8 kHz; window type: Hanning; window length: 400 samples; hop size: 30 samples; dynamic range floor: -45 dB).

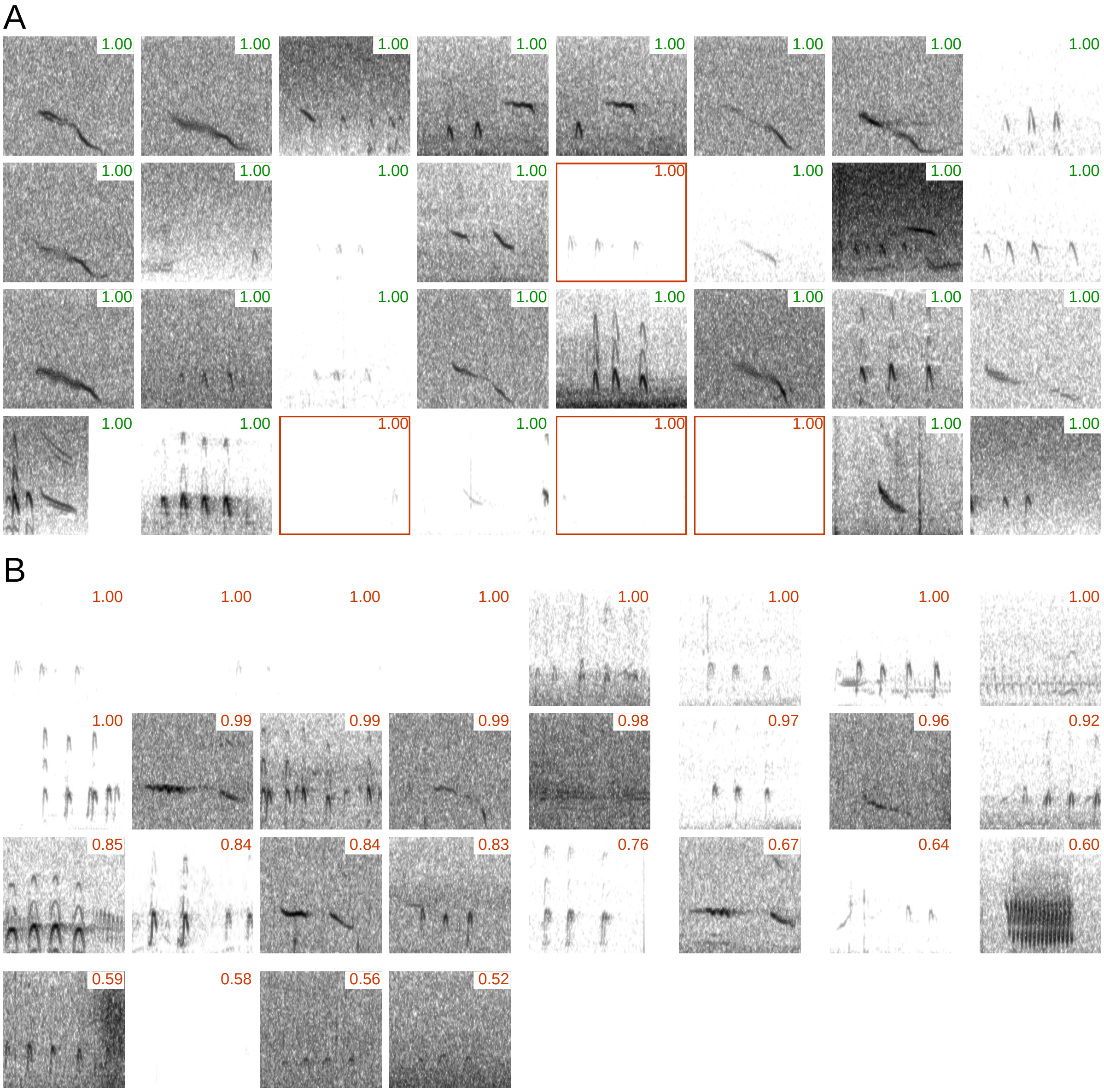


Figure S87: Classification examples for **laplon** drawn from the test dataset, shown as cropped spectrograms. Numbers in upper-right corners are calibrated probability values returned by the model. (A) Test examples scoring highest for this species. Any incorrect classifications are outlined in red. (B) Incorrectly-classified examples scoring highest for this species (i.e., the most confusing cases for the model). Spectrogram parameters: x-axis: 0-0.3 s; y-axis: 2-10 kHz; window type: Hanning; window length: 200 samples; hop size: 20 samples; dynamic range floor: -60 dB).

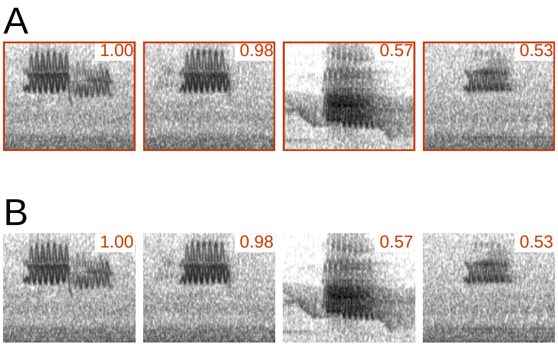


Figure S88: Classification examples for **lazbun** drawn from the test dataset, shown as cropped spectrograms. Numbers in upper-right corners are calibrated probability values returned by the model. (A) Test examples scoring highest for this species. Any incorrect classifications are outlined in red. (B) Incorrectly-classified examples scoring highest for this species (i.e., the most confusing cases for the model). Spectrogram parameters: x-axis: 0-0.3 s; y-axis: 2-10 kHz; window type: Hanning; window length: 200 samples; hop size: 20 samples; dynamic range floor: -60 dB).

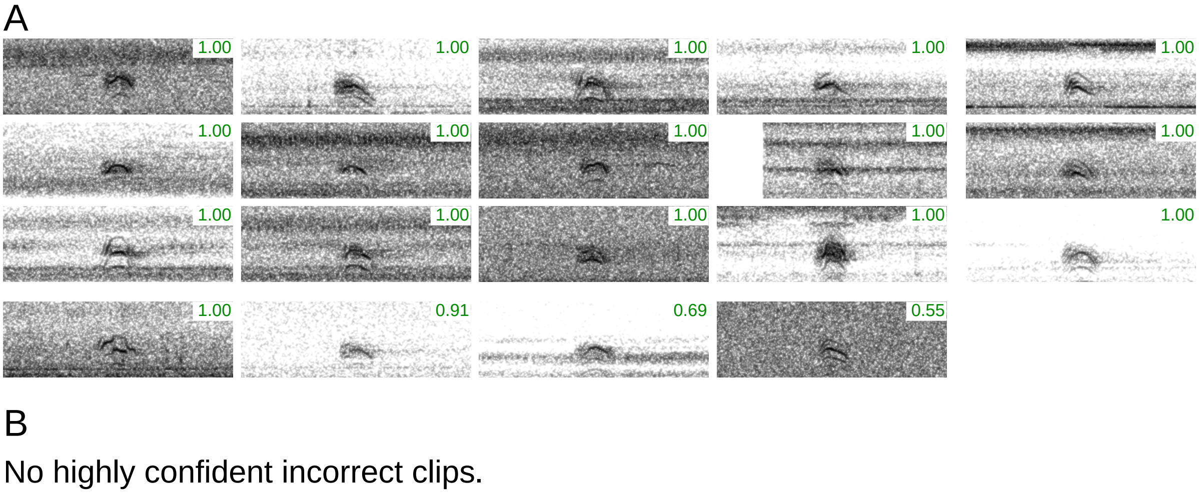


Figure S89: Classification examples for **leabit** drawn from the test dataset, shown as cropped spectrograms. Numbers in upper-right corners are calibrated probability values returned by the model. (A) Test examples scoring highest for this species. Any incorrect classifications are outlined in red. (B) Incorrectly-classified examples scoring highest for this species (i.e., the most confusing cases for the model). Spectrogram parameters: x-axis: 0-0.8 s; y-axis: 1-5 kHz; window type: Hanning; window length: 400 samples; hop size: 30 samples; dynamic range floor: -45 dB).

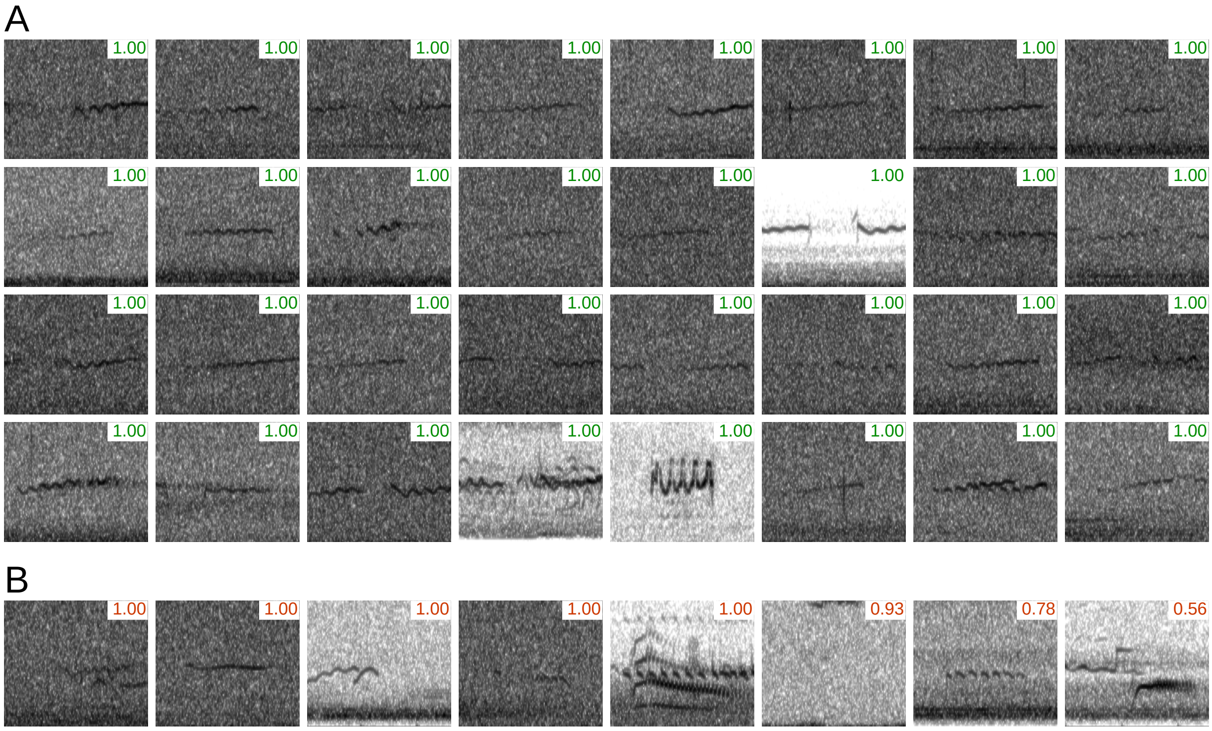


Figure S90: Classification examples for **leasan** drawn from the test dataset, shown as cropped spectrograms. Numbers in upper-right corners are calibrated probability values returned by the model. (A) Test examples scoring highest for this species. Any incorrect classifications are outlined in red. (B) Incorrectly-classified examples scoring highest for this species (i.e., the most confusing cases for the model). Spectrogram parameters: x-axis: 0-0.3 s; y-axis: 0.5-8 kHz; window type: Hanning; window length: 200 samples; hop size: 20 samples; dynamic range floor: -60 dB).

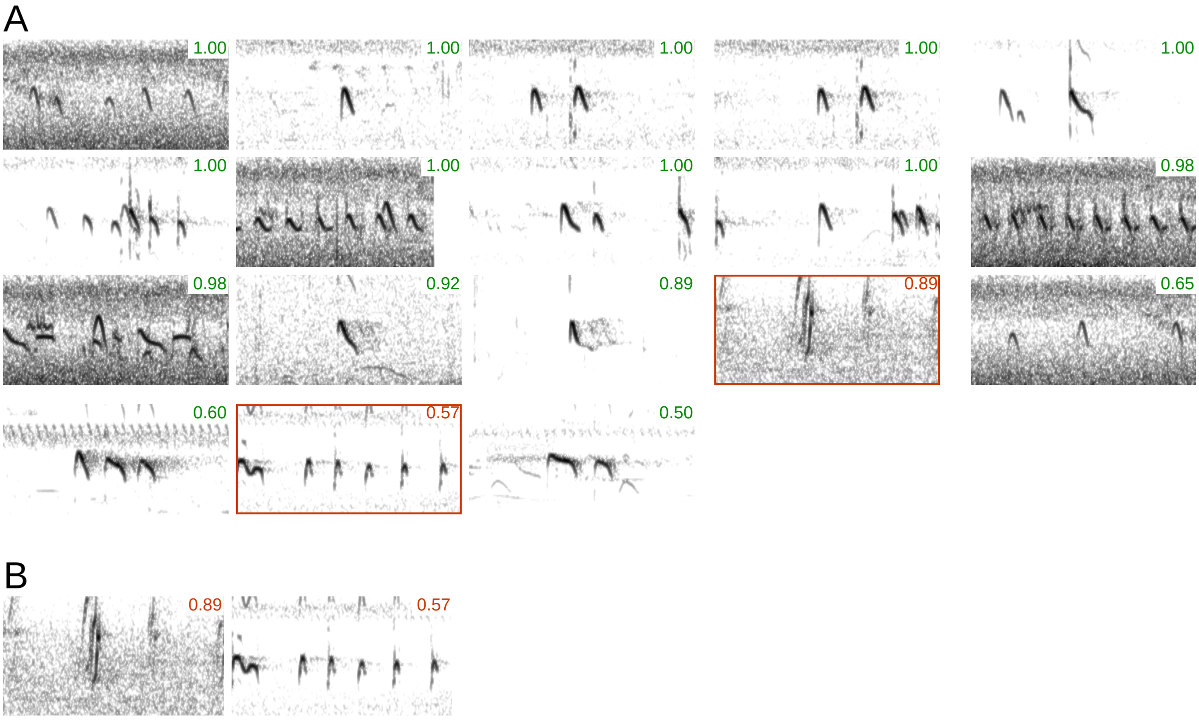


Figure S91: Classification examples for **lesyel** drawn from the test dataset, shown as cropped spectrograms. Numbers in upper-right corners are calibrated probability values returned by the model. (A) Test examples scoring highest for this species. Any incorrect classifications are outlined in red. (B) Incorrectly-classified examples scoring highest for this species (i.e., the most confusing cases for the model). Spectrogram parameters: x-axis: 0-0.8 s; y-axis: 1-5 kHz; window type: Hanning; window length: 400 samples; hop size: 30 samples; dynamic range floor: -45 dB).

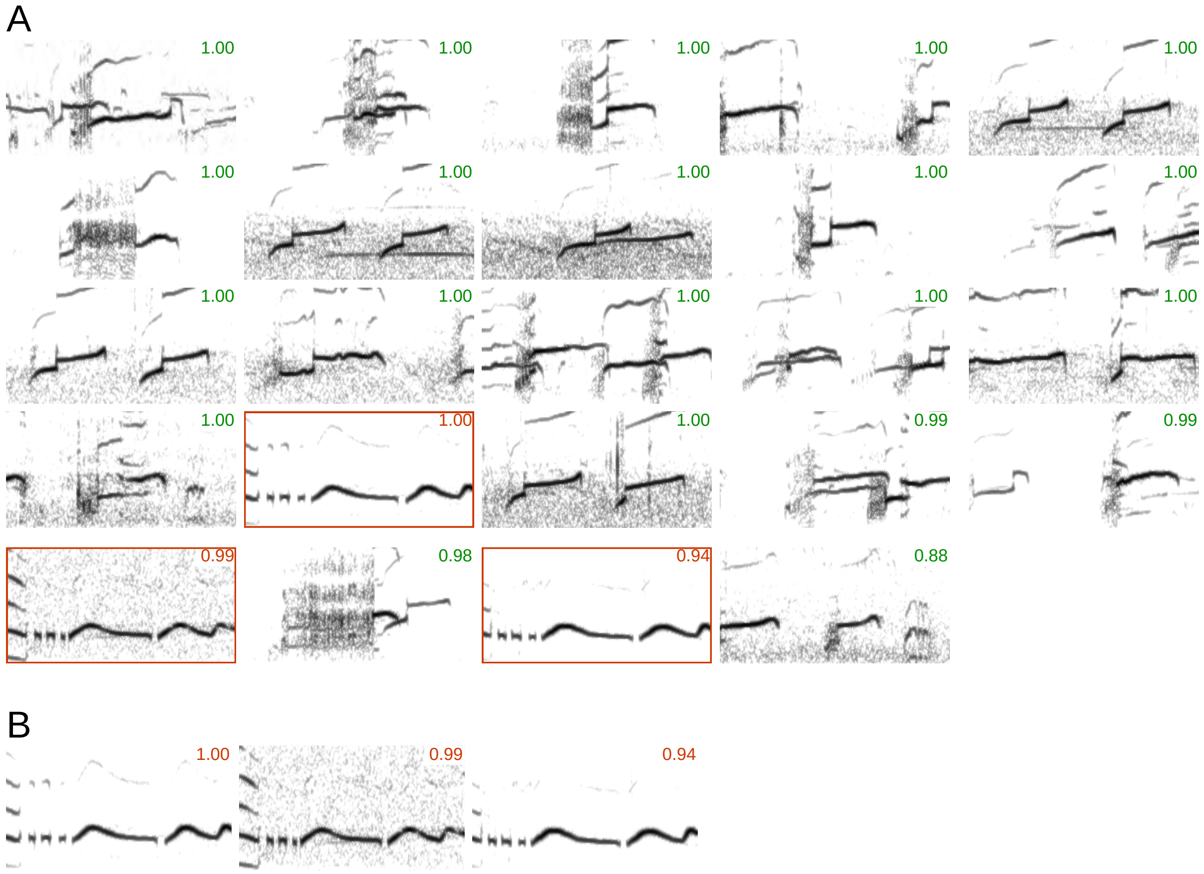


Figure S92: Classification examples for **lobcur** drawn from the test dataset, shown as cropped spectrograms. Numbers in upper-right corners are calibrated probability values returned by the model. (A) Test examples scoring highest for this species. Any incorrect classifications are outlined in red. (B) Incorrectly-classified examples scoring highest for this species (i.e., the most confusing cases for the model). Spectrogram parameters: x-axis: 0-0.8 s; y-axis: 1-5 kHz; window type: Hanning; window length: 400 samples; hop size: 30 samples; dynamic range floor: -45 dB).

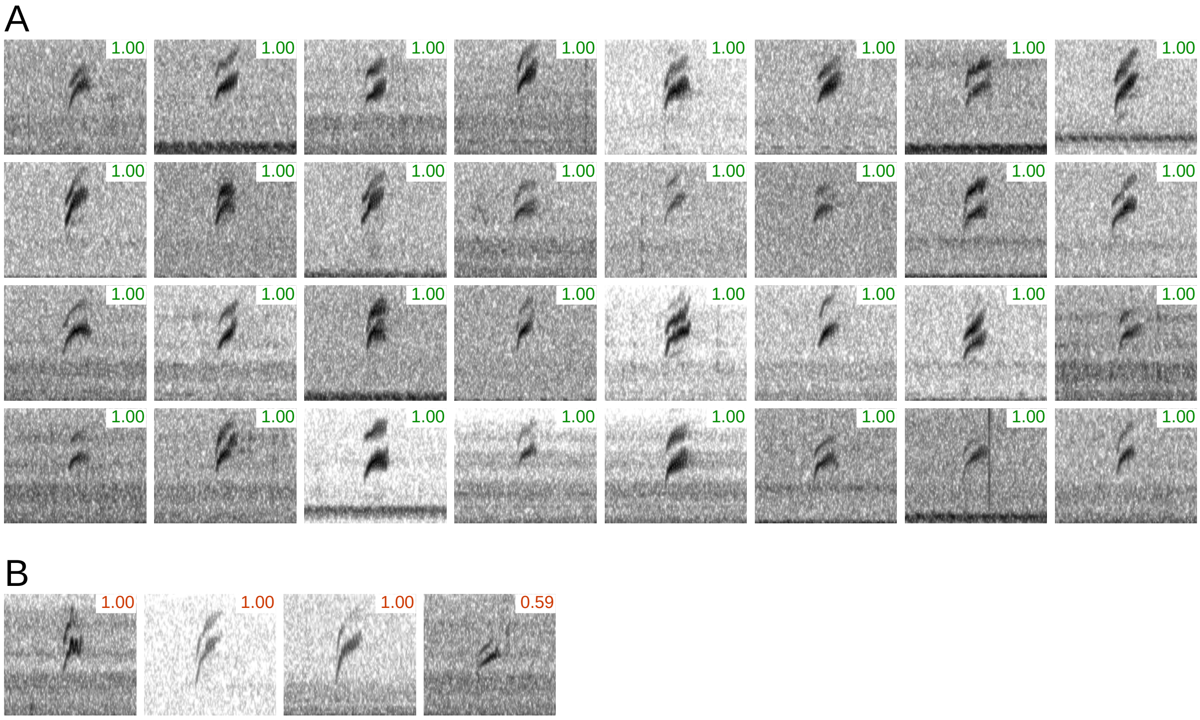


Figure S93: Classification examples for **macwar** drawn from the test dataset, shown as cropped spectrograms. Numbers in upper-right corners are calibrated probability values returned by the model. (A) Test examples scoring highest for this species. Any incorrect classifications are outlined in red. (B) Incorrectly-classified examples scoring highest for this species (i.e., the most confusing cases for the model). Spectrogram parameters: x-axis: 0-0.3 s; y-axis: 3-10 kHz; window type: Hanning; window length: 200 samples; hop size: 20 samples; dynamic range floor: -60 dB).

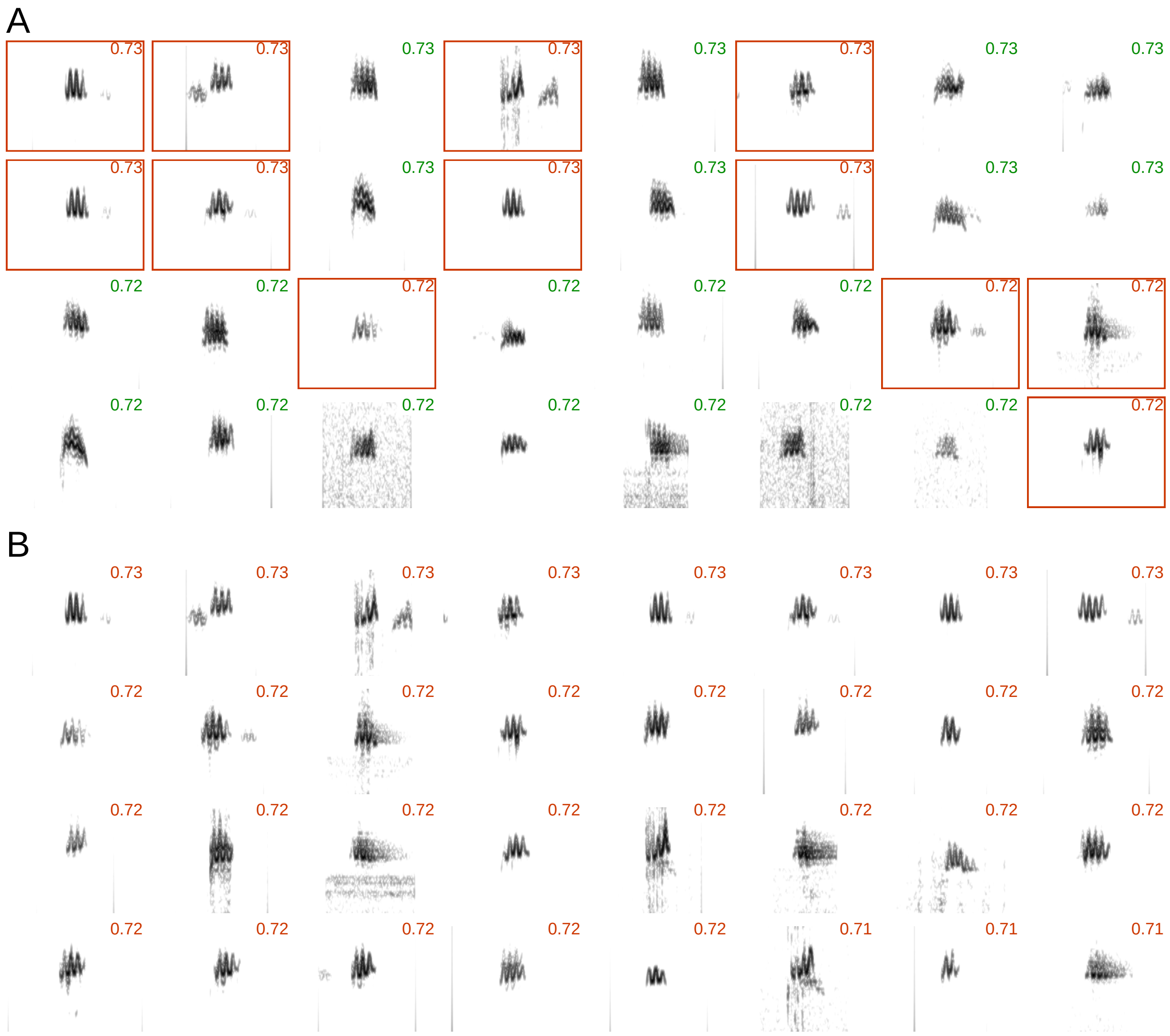


Figure S94: Classification examples for **magwar** drawn from the test dataset, shown as cropped spectrograms. Numbers in upper-right corners are calibrated probability values returned by the model. (A) Test examples scoring highest for this species. Any incorrect classifications are outlined in red. (B) Incorrectly-classified examples scoring highest for this species (i.e., the most confusing cases for the model). Spectrogram parameters: x-axis: 0-0.3 s; y-axis: 3-11 kHz; window type: Hanning; window length: 200 samples; hop size: 20 samples; dynamic range floor: -50 dB).

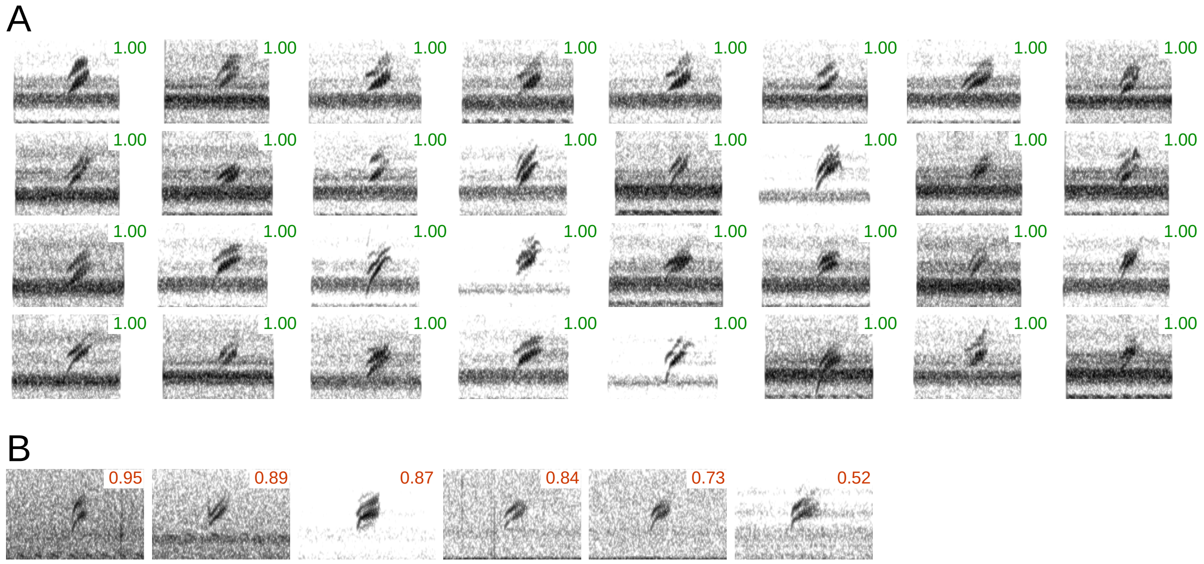


Figure S95: Classification examples for **mouwar** drawn from the test dataset, shown as cropped spectrograms. Numbers in upper-right corners are calibrated probability values returned by the model. (A) Test examples scoring highest for this species. Any incorrect classifications are outlined in red. (B) Incorrectly-classified examples scoring highest for this species (i.e., the most confusing cases for the model). Spectrogram parameters: x-axis: 0-0.3 s; y-axis: 3-10 kHz; window type: Hanning; window length: 200 samples; hop size: 20 samples; dynamic range floor: -45 dB).

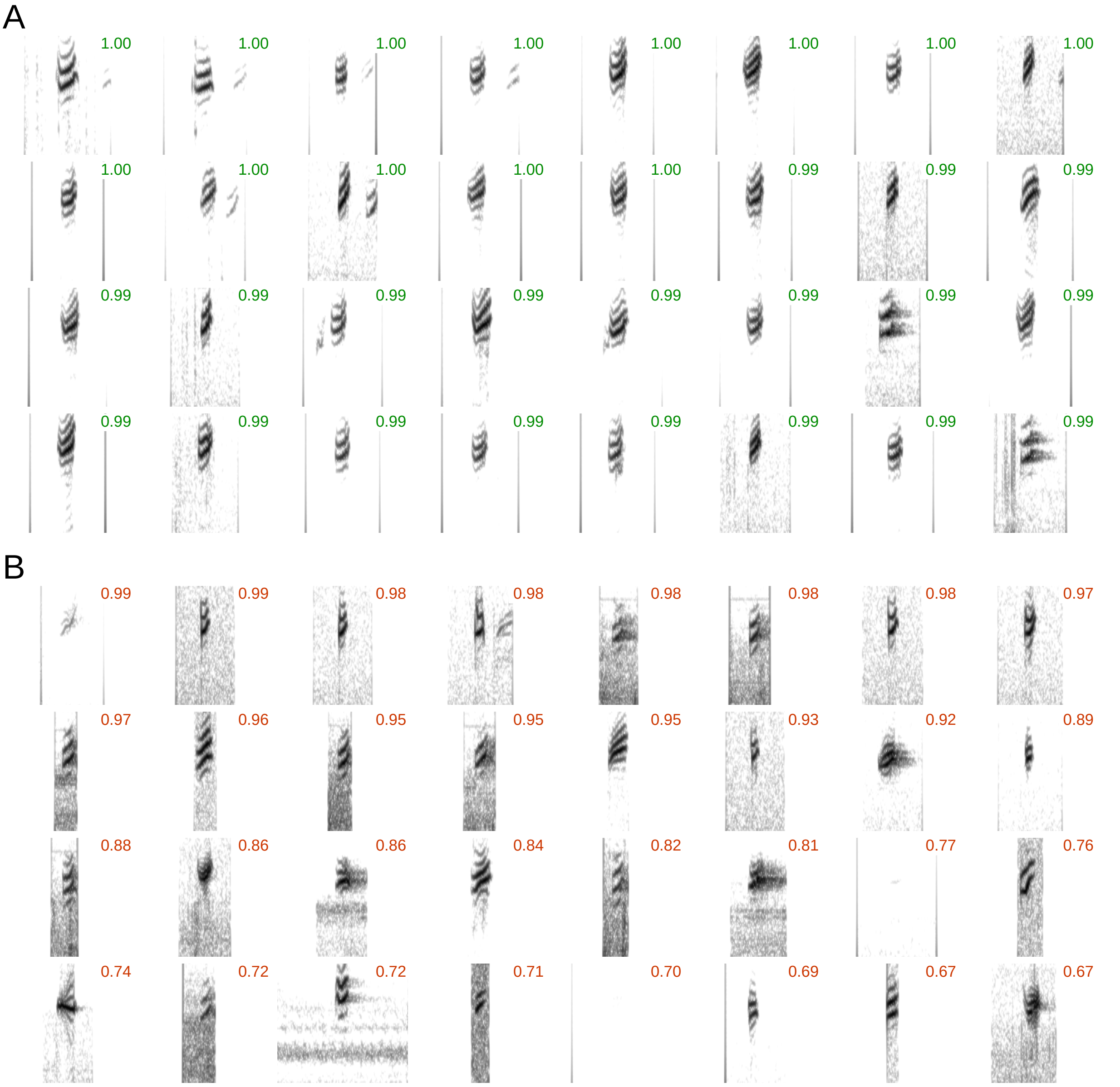


Figure S96: Classification examples for **naswar** drawn from the test dataset, shown as cropped spectrograms. Numbers in upper-right corners are calibrated probability values returned by the model. (A) Test examples scoring highest for this species. Any incorrect classifications are outlined in red. (B) Incorrectly-classified examples scoring highest for this species (i.e., the most confusing cases for the model). Spectrogram parameters: x-axis: 0-0.3 s; y-axis: 2-10 kHz; window type: Hanning; window length: 200 samples; hop size: 20 samples; dynamic range floor: -60 dB).

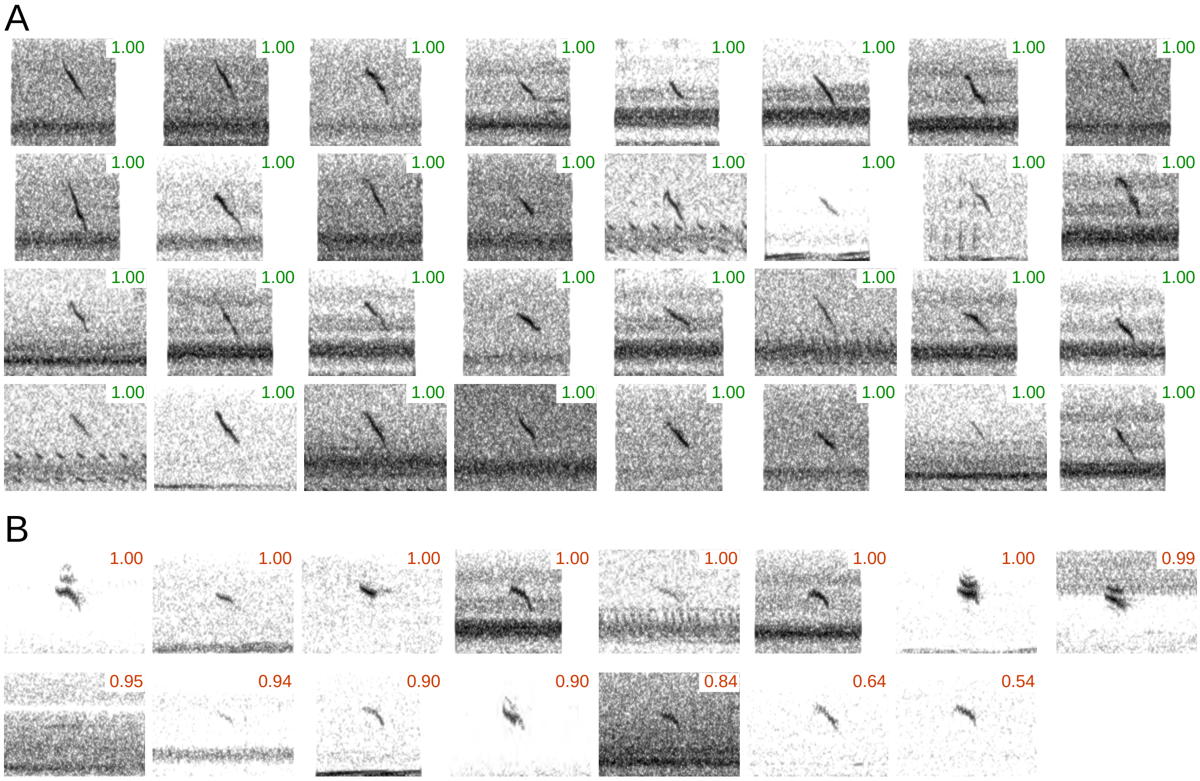


Figure S97: Classification examples for **norpar** drawn from the test dataset, shown as cropped spectrograms. Numbers in upper-right corners are calibrated probability values returned by the model. (A) Test examples scoring highest for this species. Any incorrect classifications are outlined in red. (B) Incorrectly-classified examples scoring highest for this species (i.e., the most confusing cases for the model). Spectrogram parameters: x-axis: 0-0.3 s; y-axis: 3-10 kHz; window type: Hanning; window length: 250 samples; hop size: 20 samples; dynamic range floor: -45 dB).

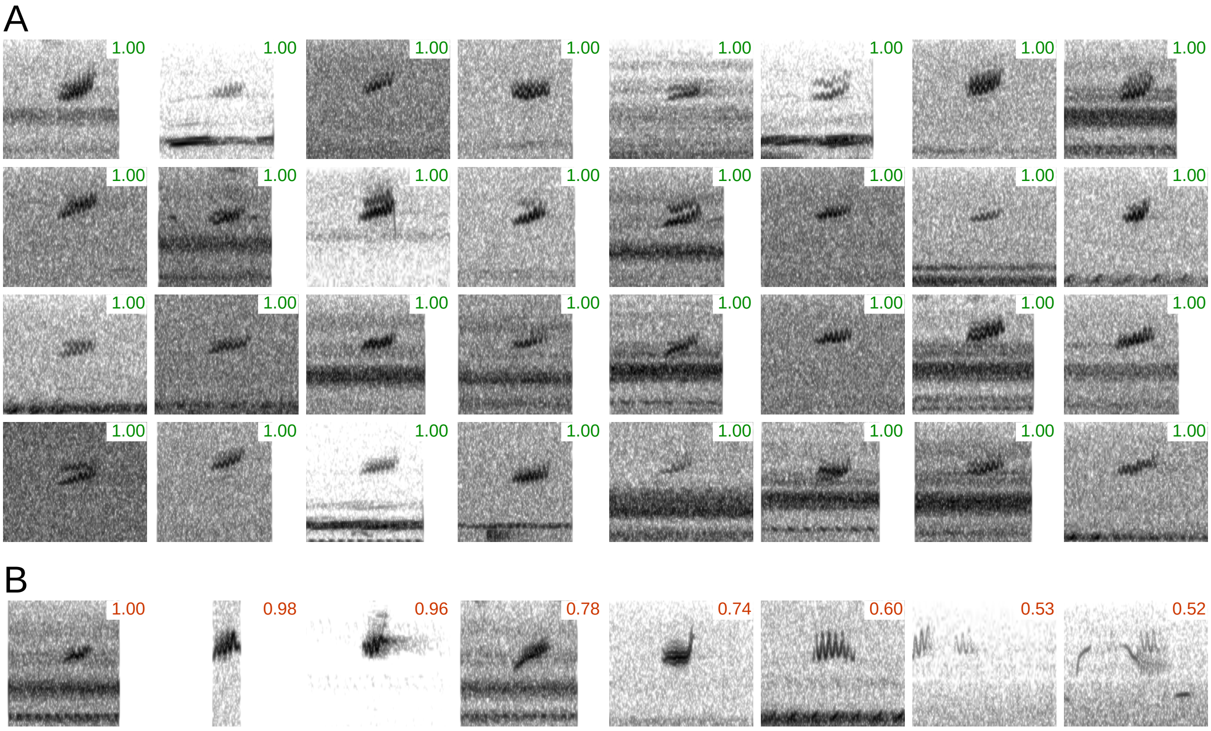


Figure S98: Classification examples for **norwat** drawn from the test dataset, shown as cropped spectrograms. Numbers in upper-right corners are calibrated probability values returned by the model. (A) Test examples scoring highest for this species. Any incorrect classifications are outlined in red. (B) Incorrectly-classified examples scoring highest for this species (i.e., the most confusing cases for the model). Spectrogram parameters: x-axis: 0-0.3 s; y-axis: 2-10 kHz; window type: Hanning; window length: 200 samples; hop size: 20 samples; dynamic range floor: -60 dB).

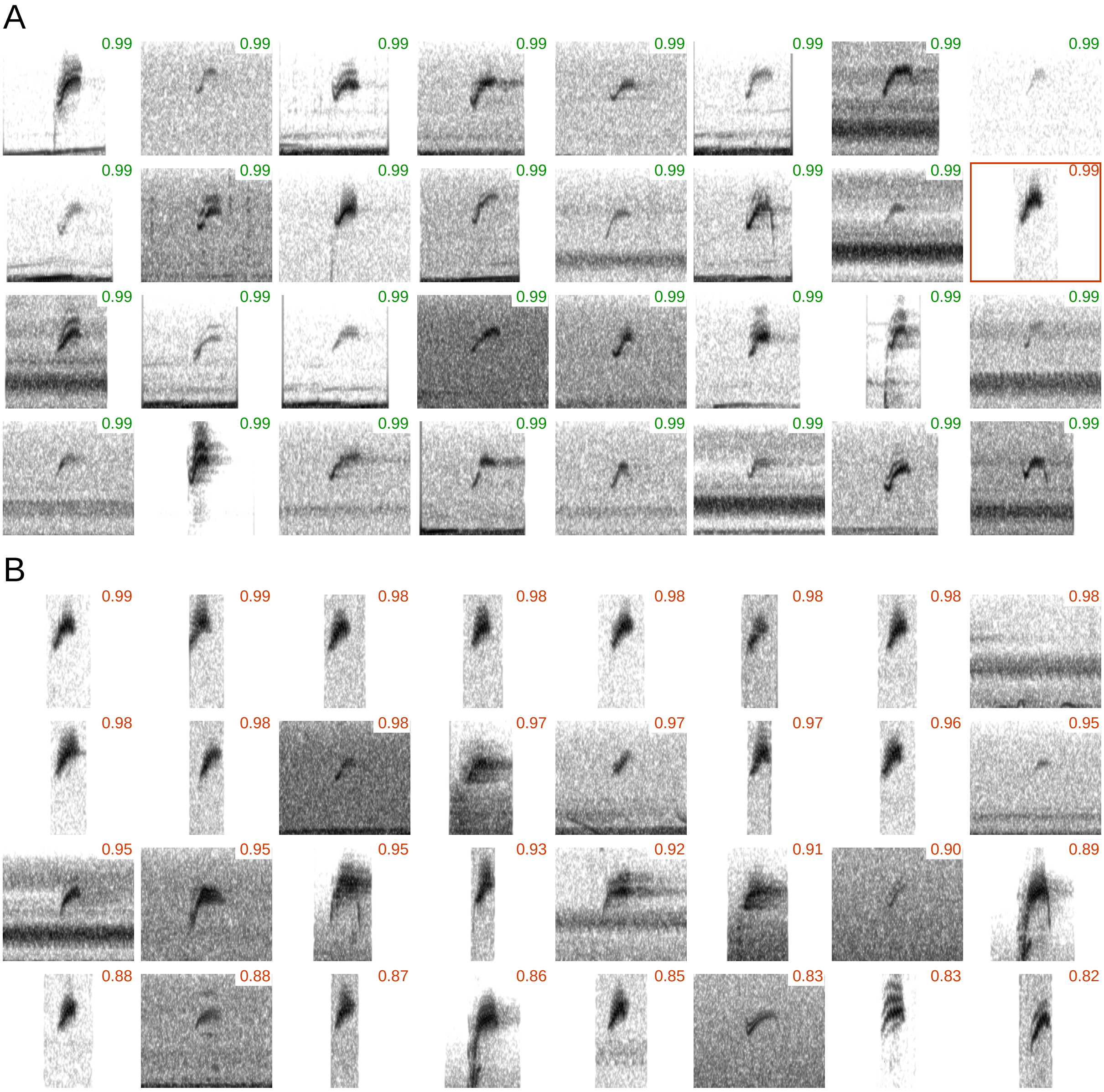


Figure S99: Classification examples for **ovenbi1** drawn from the test dataset, shown as cropped spectrograms. Numbers in upper-right corners are calibrated probability values returned by the model. (A) Test examples scoring highest for this species. Any incorrect classifications are outlined in red. (B) Incorrectly-classified examples scoring highest for this species (i.e., the most confusing cases for the model). Spectrogram parameters: x-axis: 0-0.3 s; y-axis: 3-11 kHz; window type: Hanning; window length: 200 samples; hop size: 20 samples; dynamic range floor: -60 dB).

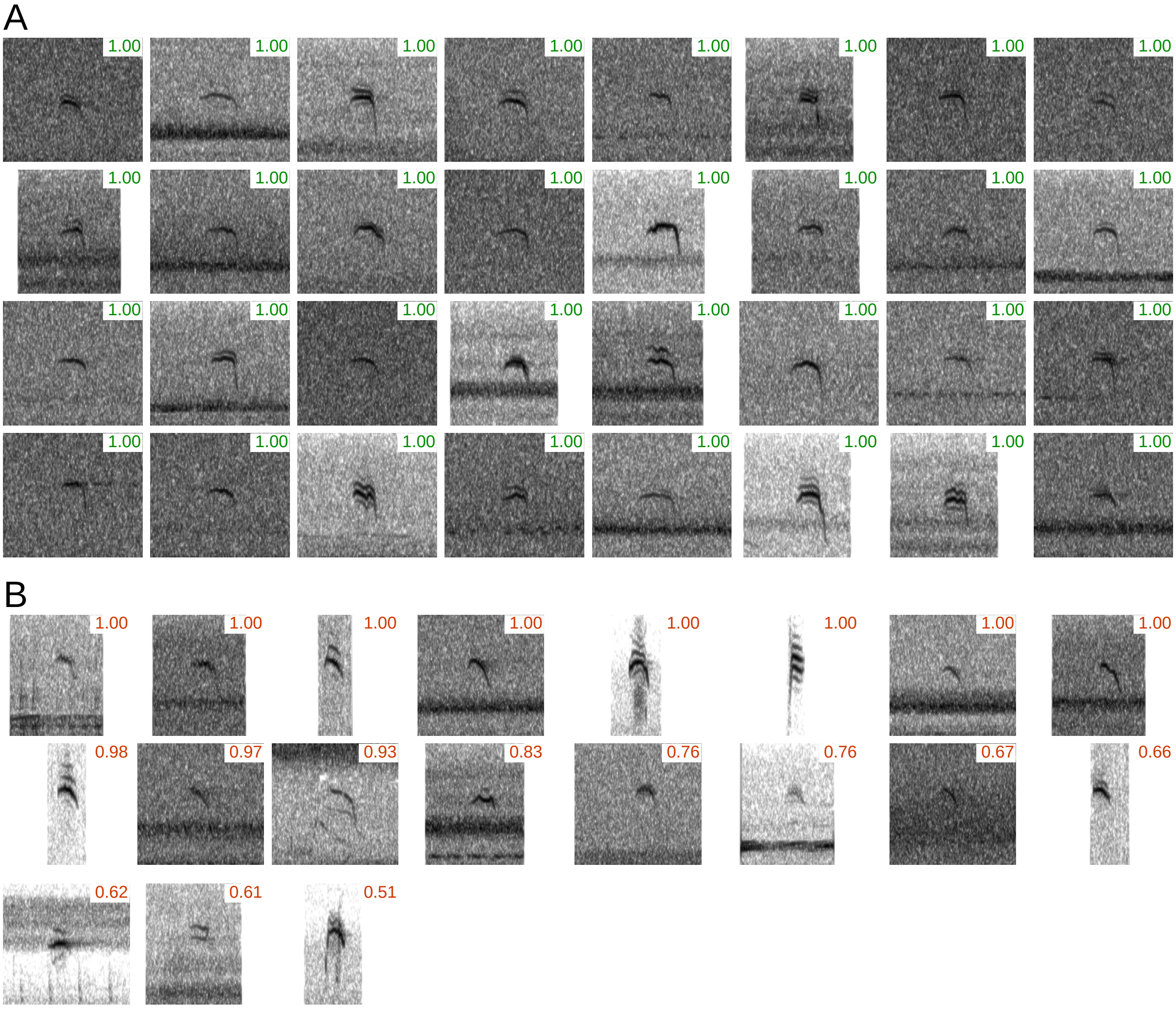


Figure S100: Classification examples for **palwar** drawn from the test dataset, shown as cropped spectrograms. Numbers in upper-right corners are calibrated probability values returned by the model. (A) Test examples scoring highest for this species. Any incorrect classifications are outlined in red. (B) Incorrectly-classified examples scoring highest for this species (i.e., the most confusing cases for the model). Spectrogram parameters: x-axis: 0-0.3 s; y-axis: 2-10 kHz; window type: Hanning; window length: 200 samples; hop size: 20 samples; dynamic range floor: -60 dB).

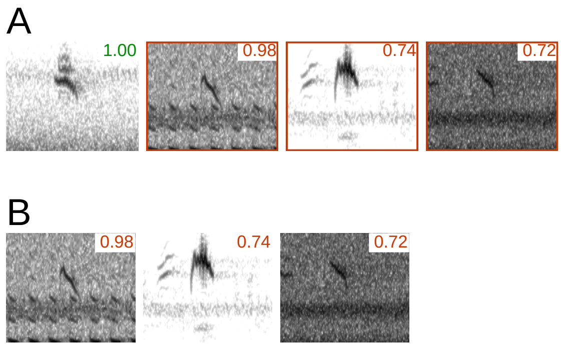


Figure S101: Classification examples for **pinwar** drawn from the test dataset, shown as cropped spectrograms. Numbers in upper-right corners are calibrated probability values returned by the model. (A) Test examples scoring highest for this species. Any incorrect classifications are outlined in red. (B) Incorrectly-classified examples scoring highest for this species (i.e., the most confusing cases for the model). Spectrogram parameters: x-axis: 0-0.3 s; y-axis: 2-10 kHz; window type: Hanning; window length: 200 samples; hop size: 20 samples; dynamic range floor: -60 dB).

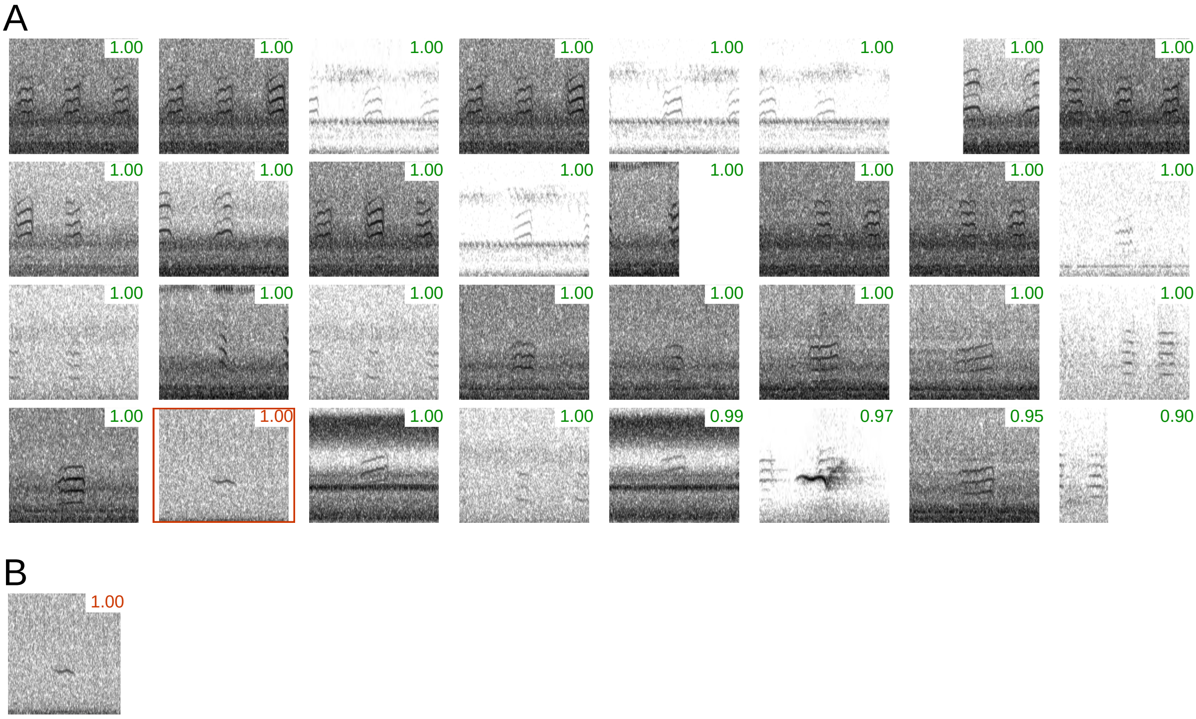


Figure S102: Classification examples for **rebnut** drawn from the test dataset, shown as cropped spectrograms. Numbers in upper-right corners are calibrated probability values returned by the model. (A) Test examples scoring highest for this species. Any incorrect classifications are outlined in red. (B) Incorrectly-classified examples scoring highest for this species (i.e., the most confusing cases for the model). Spectrogram parameters: x-axis: 0-1 s; y-axis: 1-6 kHz; window type: Hanning; window length: 420 samples; hop size: 35 samples; dynamic range floor: -50 dB).

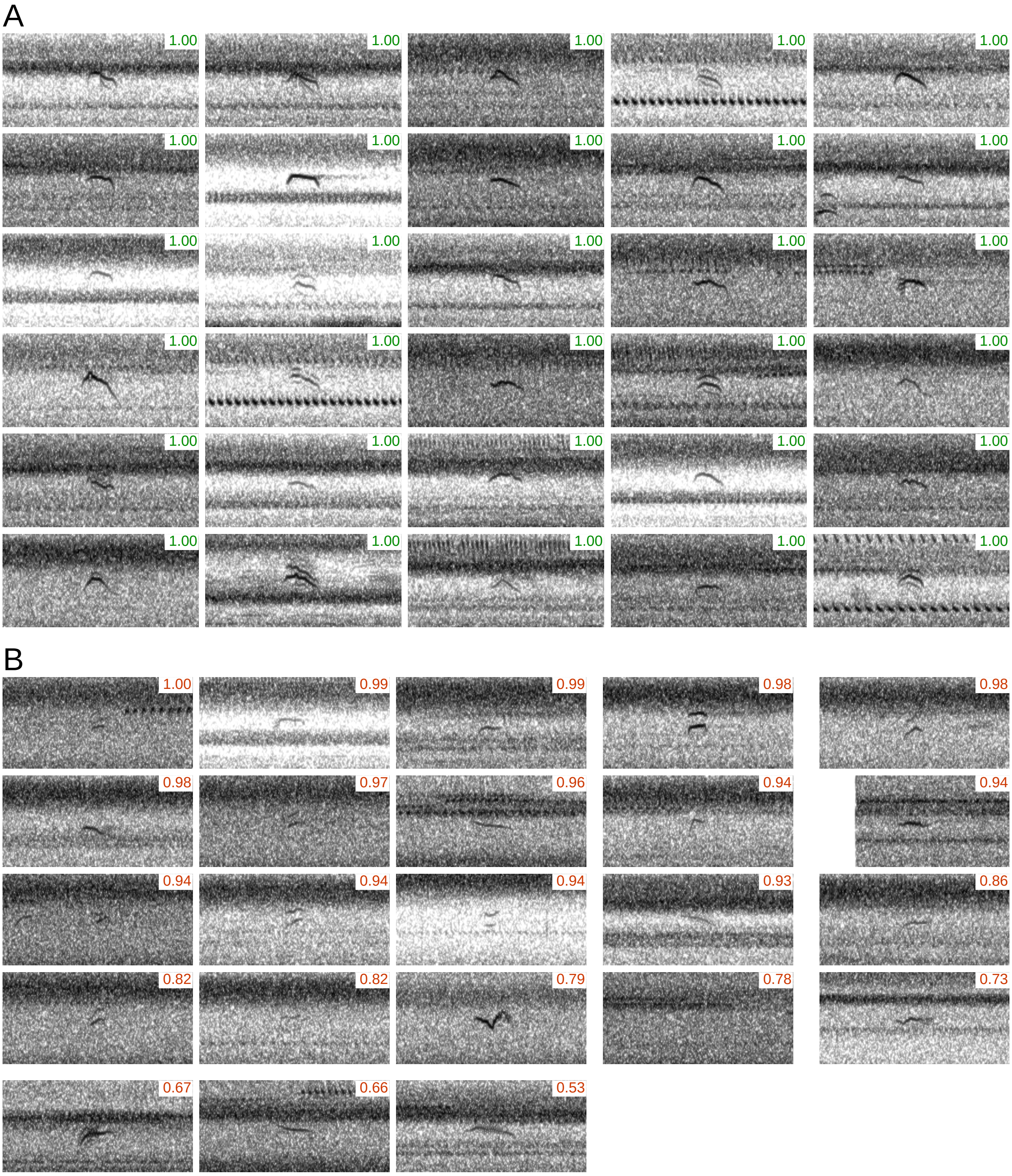


Figure S103: Classification examples for **robgro** drawn from the test dataset, shown as cropped spectrograms. Numbers in upper-right corners are calibrated probability values returned by the model. (A) Test examples scoring highest for this species. Any incorrect classifications are outlined in red. (B) Incorrectly-classified examples scoring highest for this species (i.e., the most confusing cases for the model). Spectrogram parameters: x-axis: 0-0.8 s; y-axis: 1-5 kHz; window type: Hanning; window length: 400 samples; hop size: 30 samples; dynamic range floor: -45 dB).

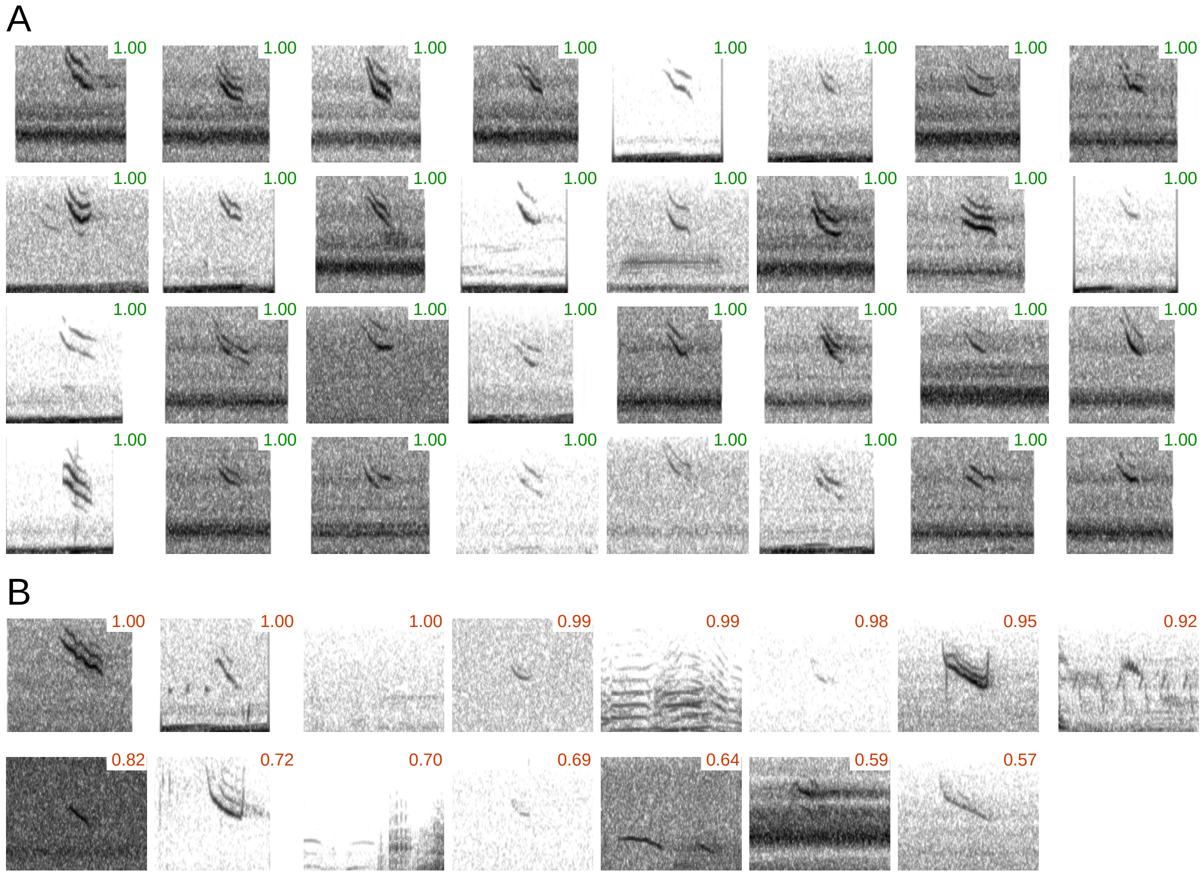


Figure S104: Classification examples for **savspa** drawn from the test dataset, shown as cropped spectrograms. Numbers in upper-right corners are calibrated probability values returned by the model. (A) Test examples scoring highest for this species. Any incorrect classifications are outlined in red. (B) Incorrectly-classified examples scoring highest for this species (i.e., the most confusing cases for the model). Spectrogram parameters: x-axis: 0-0.3 s; y-axis: 3-11 kHz; window type: Hanning; window length: 200 samples; hop size: 20 samples; dynamic range floor: -60 dB).

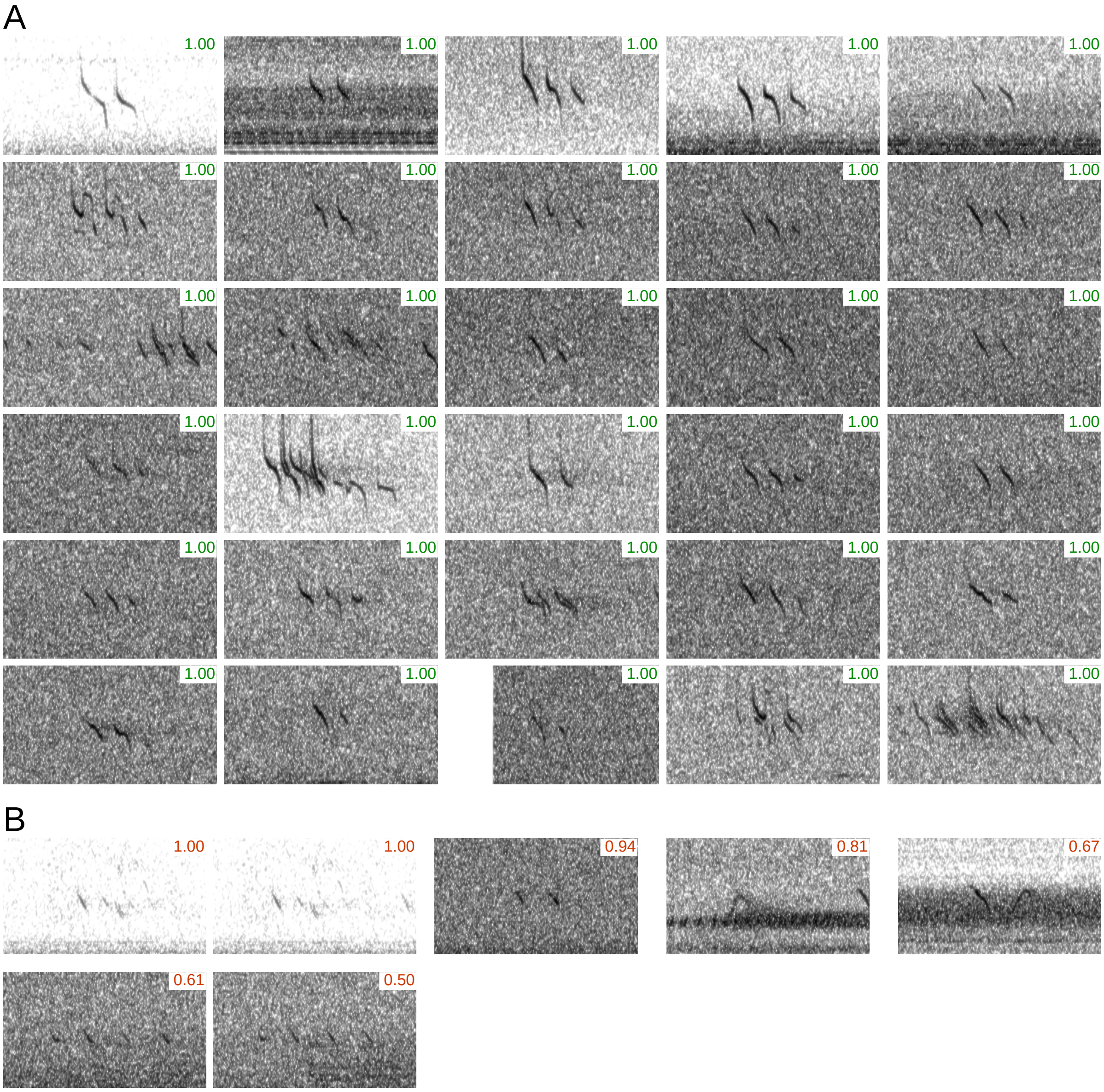


Figure S107: Classification examples for **shbdow** drawn from the test dataset, shown as cropped spectrograms. Numbers in upper-right corners are calibrated probability values returned by the model. (A) Test examples scoring highest for this species. Any incorrect classifications are outlined in red. (B) Incorrectly-classified examples scoring highest for this species (i.e., the most confusing cases for the model). Spectrogram parameters: x-axis: 0-0.8 s; y-axis: 1-5 kHz; window type: Hanning; window length: 400 samples; hop size: 30 samples; dynamic range floor: -45 dB).

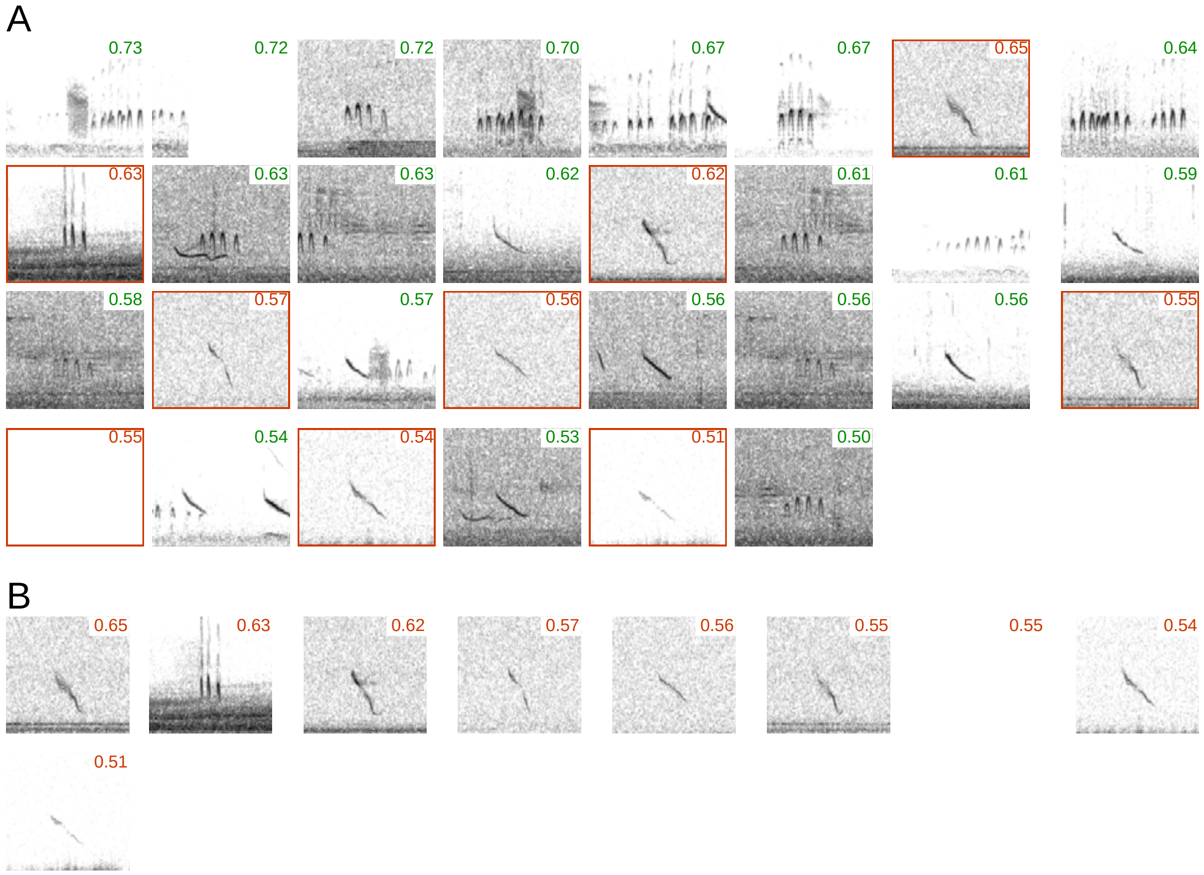


Figure S108: Classification examples for **snobun** drawn from the test dataset, shown as cropped spectrograms. Numbers in upper-right corners are calibrated probability values returned by the model. (A) Test examples scoring highest for this species. Any incorrect classifications are outlined in red. (B) Incorrectly-classified examples scoring highest for this species (i.e., the most confusing cases for the model). Spectrogram parameters: x-axis: 0-0.8 s; y-axis: 1-8 kHz; window type: Hanning; window length: 400 samples; hop size: 30 samples; dynamic range floor: -45 dB).

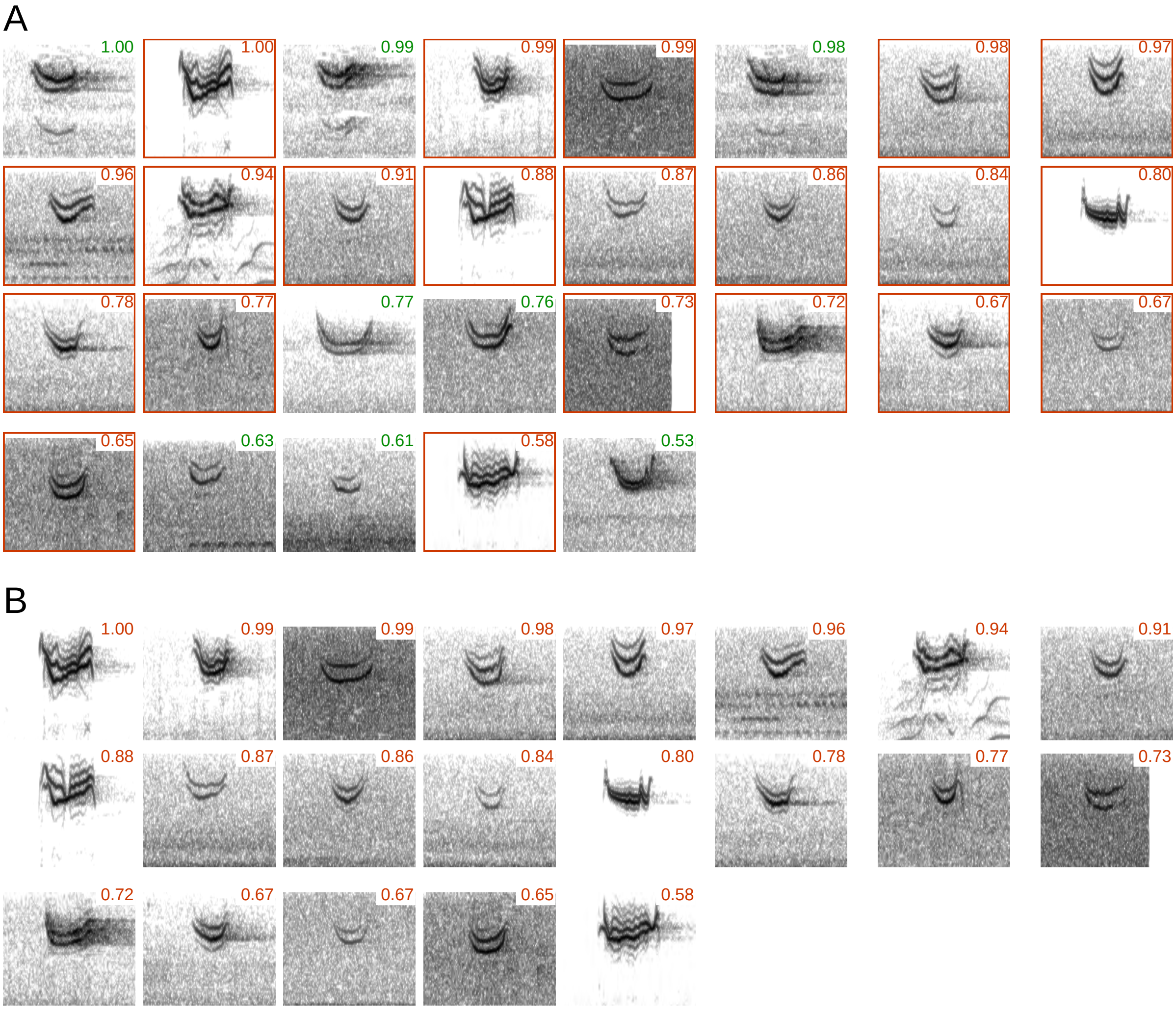


Figure S110: Classification examples for **sonspa** drawn from the test dataset, shown as cropped spectrograms. Numbers in upper-right corners are calibrated probability values returned by the model. (A) Test examples scoring highest for this species. Any incorrect classifications are outlined in red. (B) Incorrectly-classified examples scoring highest for this species (i.e., the most confusing cases for the model). Spectrogram parameters: x-axis: 0-0.3 s; y-axis: 3-11 kHz; window type: Hanning; window length: 200 samples; hop size: 20 samples; dynamic range floor: -60 dB).

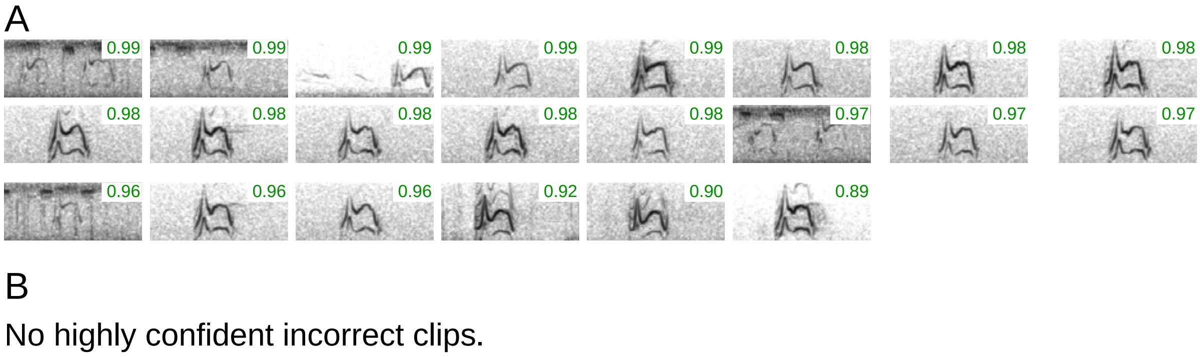


Figure S112: Classification examples for **sprpip** drawn from the test dataset, shown as cropped spectrograms. Numbers in upper-right corners are calibrated probability values returned by the model. (A) Test examples scoring highest for this species. Any incorrect classifications are outlined in red. (B) Incorrectly-classified examples scoring highest for this species (i.e., the most confusing cases for the model). Spectrogram parameters: x-axis: 0-0.3 s; y-axis: 2-10 kHz; window type: Hanning; window length: 200 samples; hop size: 20 samples; dynamic range floor: -60 dB).

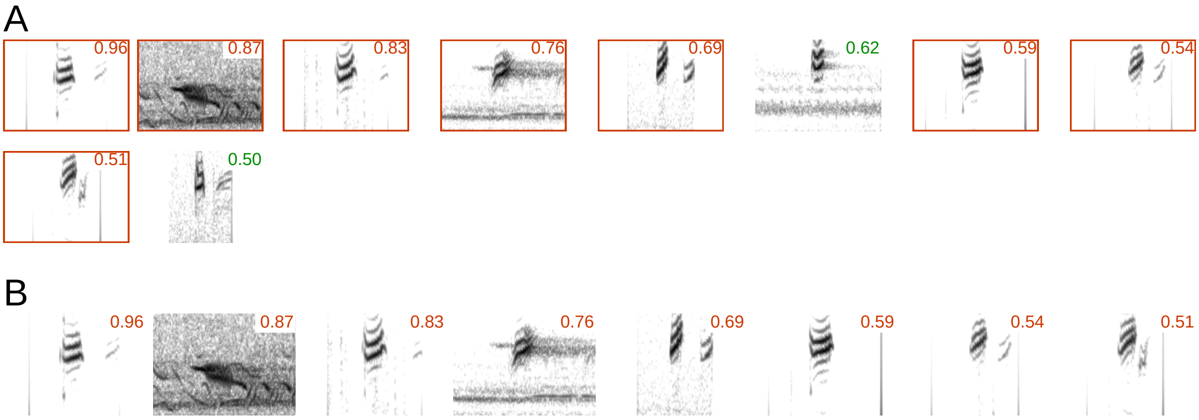


Figure S114: Classification examples for **tenwar** drawn from the test dataset, shown as cropped spectrograms. Numbers in upper-right corners are calibrated probability values returned by the model. (A) Test examples scoring highest for this species. Any incorrect classifications are outlined in red. (B) Incorrectly-classified examples scoring highest for this species (i.e., the most confusing cases for the model). Spectrogram parameters: x-axis: 0-0.3 s; y-axis: 2-10 kHz; window type: Hanning; window length: 200 samples; hop size: 20 samples; dynamic range floor: -60 dB).

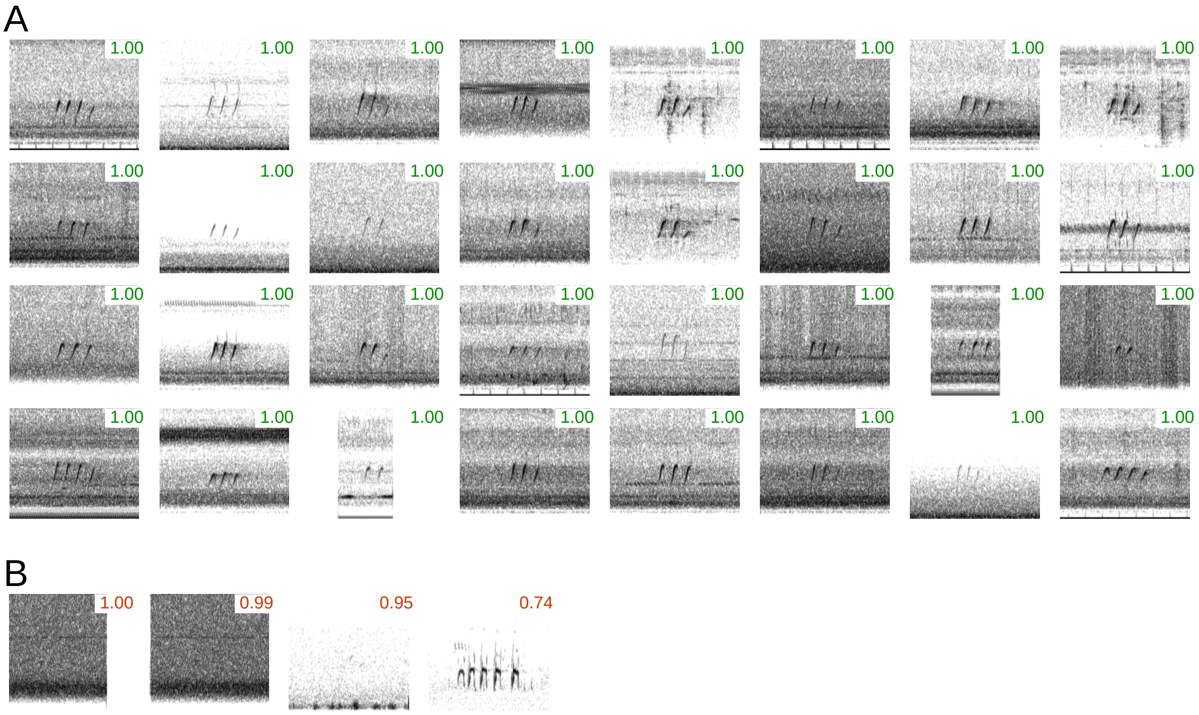


Figure S115: Classification examples for **uplsan** drawn from the test dataset, shown as cropped spectrograms. Numbers in upper-right corners are calibrated probability values returned by the model. (A) Test examples scoring highest for this species. Any incorrect classifications are outlined in red. (B) Incorrectly-classified examples scoring highest for this species (i.e., the most confusing cases for the model). Spectrogram parameters: x-axis: 0-1 s; y-axis: 0-6 kHz; window type: Hanning; window length: 420 samples; hop size: 35 samples; dynamic range floor: -50 dB).

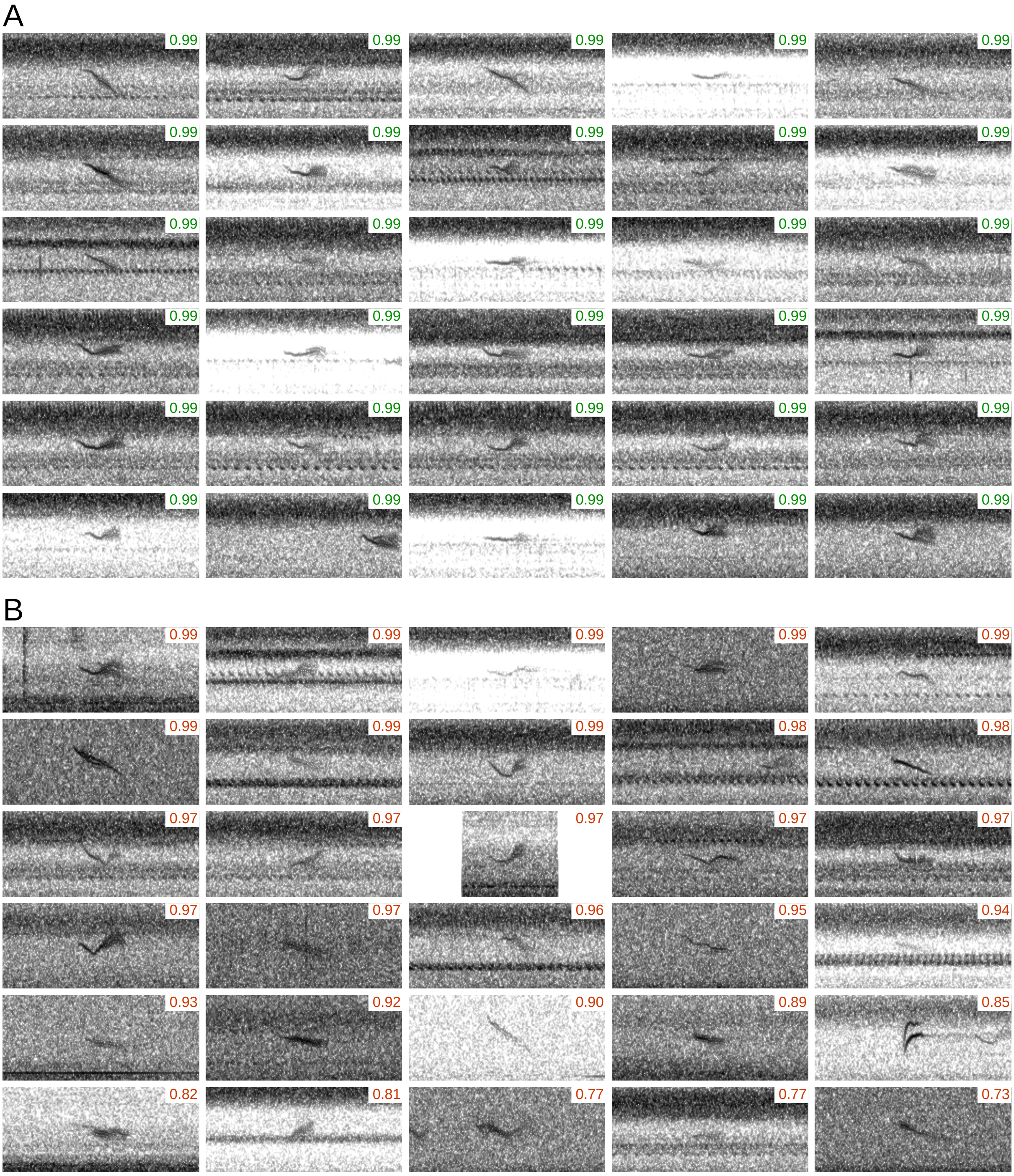


Figure S116: Classification examples for **veery** drawn from the test dataset, shown as cropped spectrograms. Numbers in upper-right corners are calibrated probability values returned by the model. (A) Test examples scoring highest for this species. Any incorrect classifications are outlined in red. (B) Incorrectly-classified examples scoring highest for this species (i.e., the most confusing cases for the model). Spectrogram parameters: x-axis: 0-0.8 s; y-axis: 1-5 kHz; window type: Hanning; window length: 400 samples; hop size: 30 samples; dynamic range floor: -45 dB).

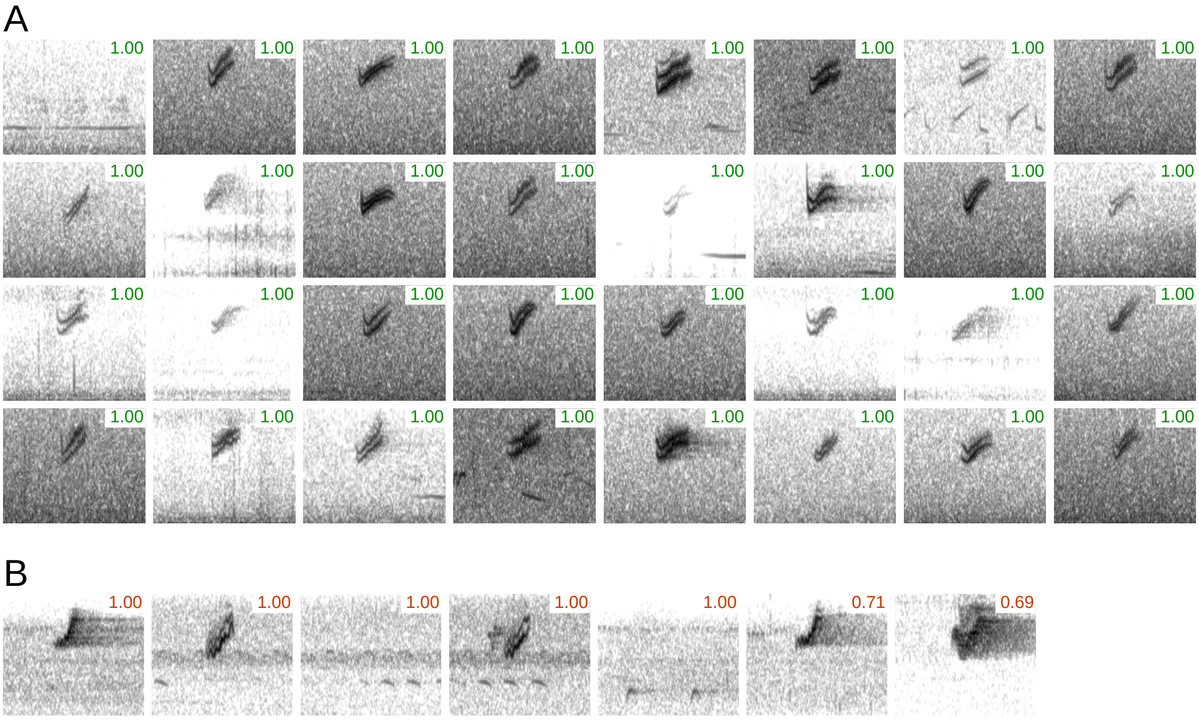


Figure S117: Classification examples for **vesspa** drawn from the test dataset, shown as cropped spectrograms. Numbers in upper-right corners are calibrated probability values returned by the model. (A) Test examples scoring highest for this species. Any incorrect classifications are outlined in red. (B) Incorrectly-classified examples scoring highest for this species (i.e., the most confusing cases for the model). Spectrogram parameters: x-axis: 0-0.3 s; y-axis: 2-10 kHz; window type: Hanning; window length: 200 samples; hop size: 20 samples; dynamic range floor: -60 dB).

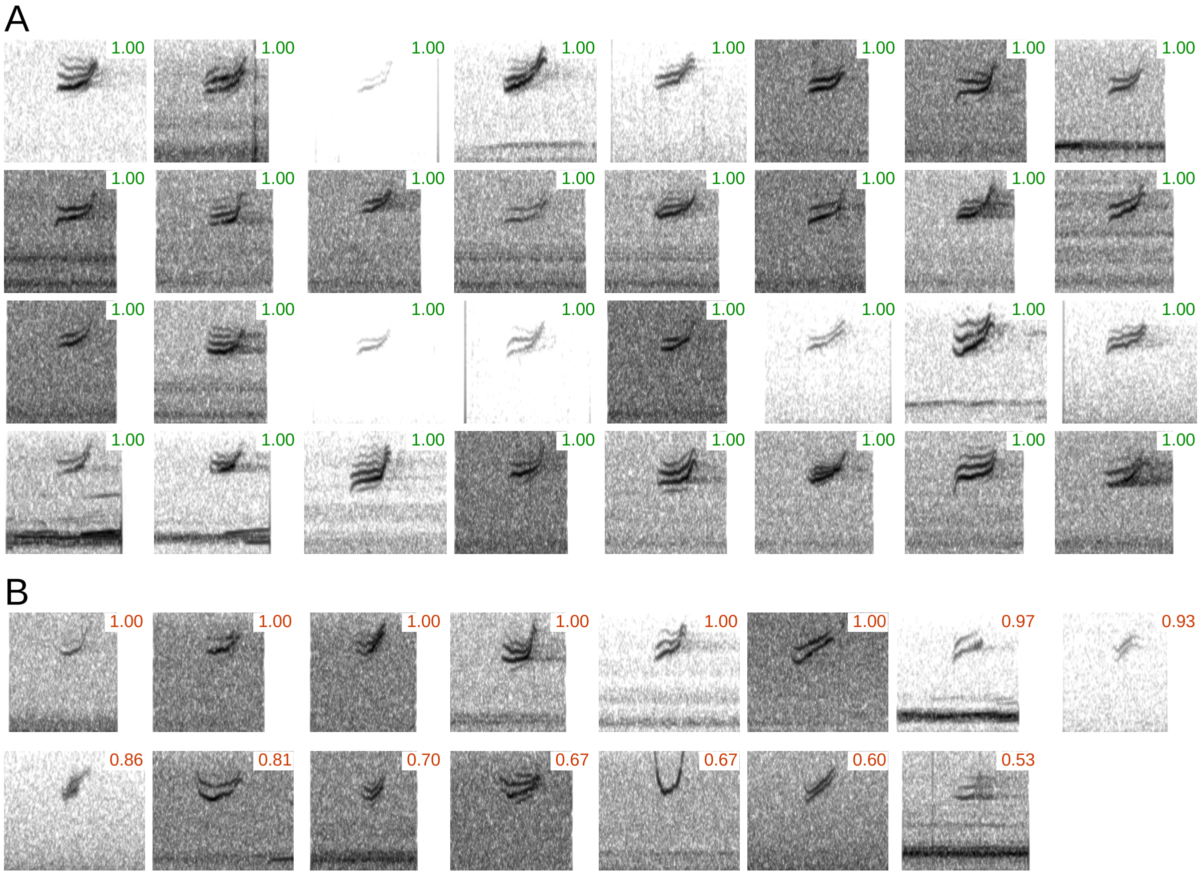


Figure S118: Classification examples for **whcspa** drawn from the test dataset, shown as cropped spectrograms. Numbers in upper-right corners are calibrated probability values returned by the model. (A) Test examples scoring highest for this species. Any incorrect classifications are outlined in red. (B) Incorrectly-classified examples scoring highest for this species (i.e., the most confusing cases for the model). Spectrogram parameters: x-axis: 0-0.3 s; y-axis: 2-10 kHz; window type: Hanning; window length: 200 samples; hop size: 20 samples; dynamic range floor: -60 dB).

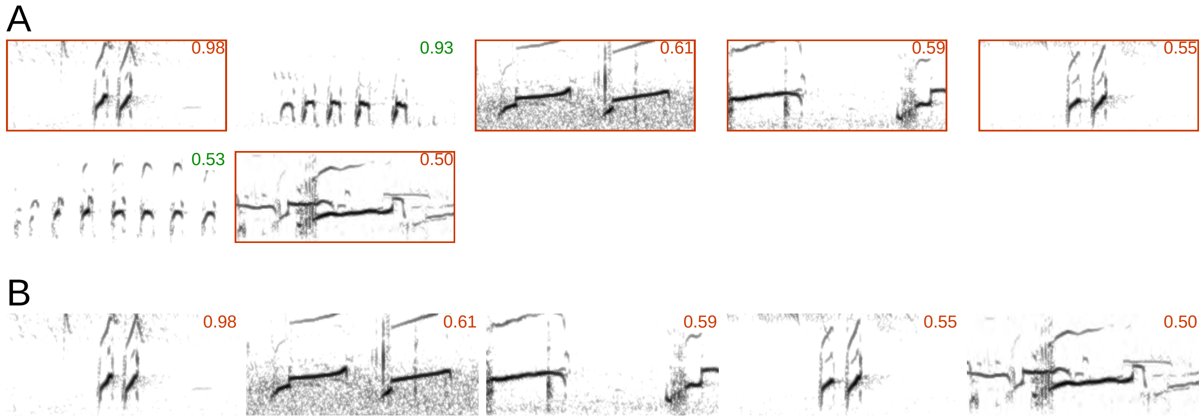


Figure S119: Classification examples for **whimbr** drawn from the test dataset, shown as cropped spectrograms. Numbers in upper-right corners are calibrated probability values returned by the model. (A) Test examples scoring highest for this species. Any incorrect classifications are outlined in red. (B) Incorrectly-classified examples scoring highest for this species (i.e., the most confusing cases for the model). Spectrogram parameters: x-axis: 0-0.8 s; y-axis: 1-5 kHz; window type: Hanning; window length: 400 samples; hop size: 30 samples; dynamic range floor: -45 dB).

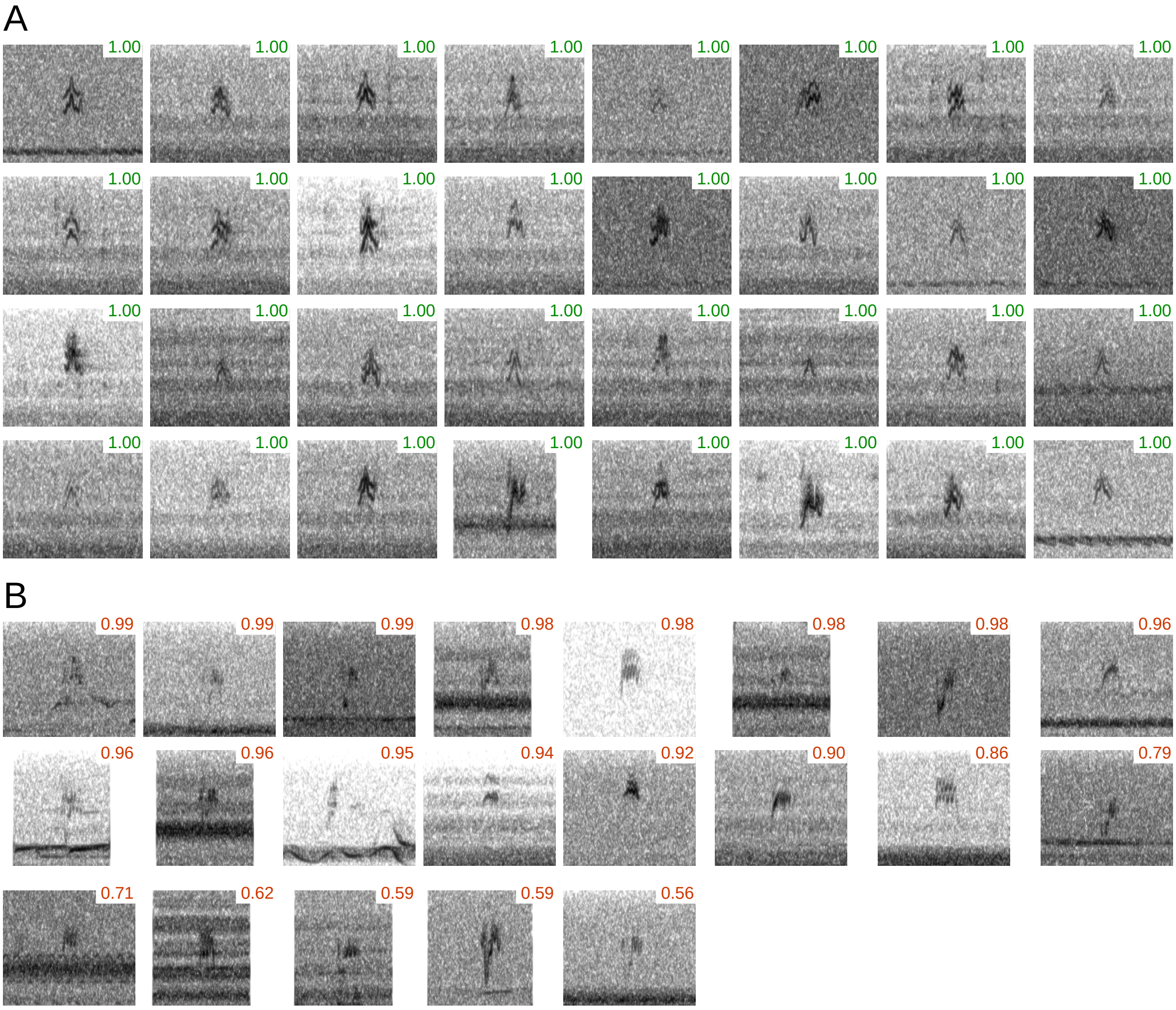


Figure S120: Classification examples for **wlswar** drawn from the test dataset, shown as cropped spectrograms. Numbers in upper-right corners are calibrated probability values returned by the model. (A) Test examples scoring highest for this species. Any incorrect classifications are outlined in red. (B) Incorrectly-classified examples scoring highest for this species (i.e., the most confusing cases for the model). Spectrogram parameters: x-axis: 0-0.3 s; y-axis: 2-11 kHz; window type: Hanning; window length: 200 samples; hop size: 20 samples; dynamic range floor: -60 dB).

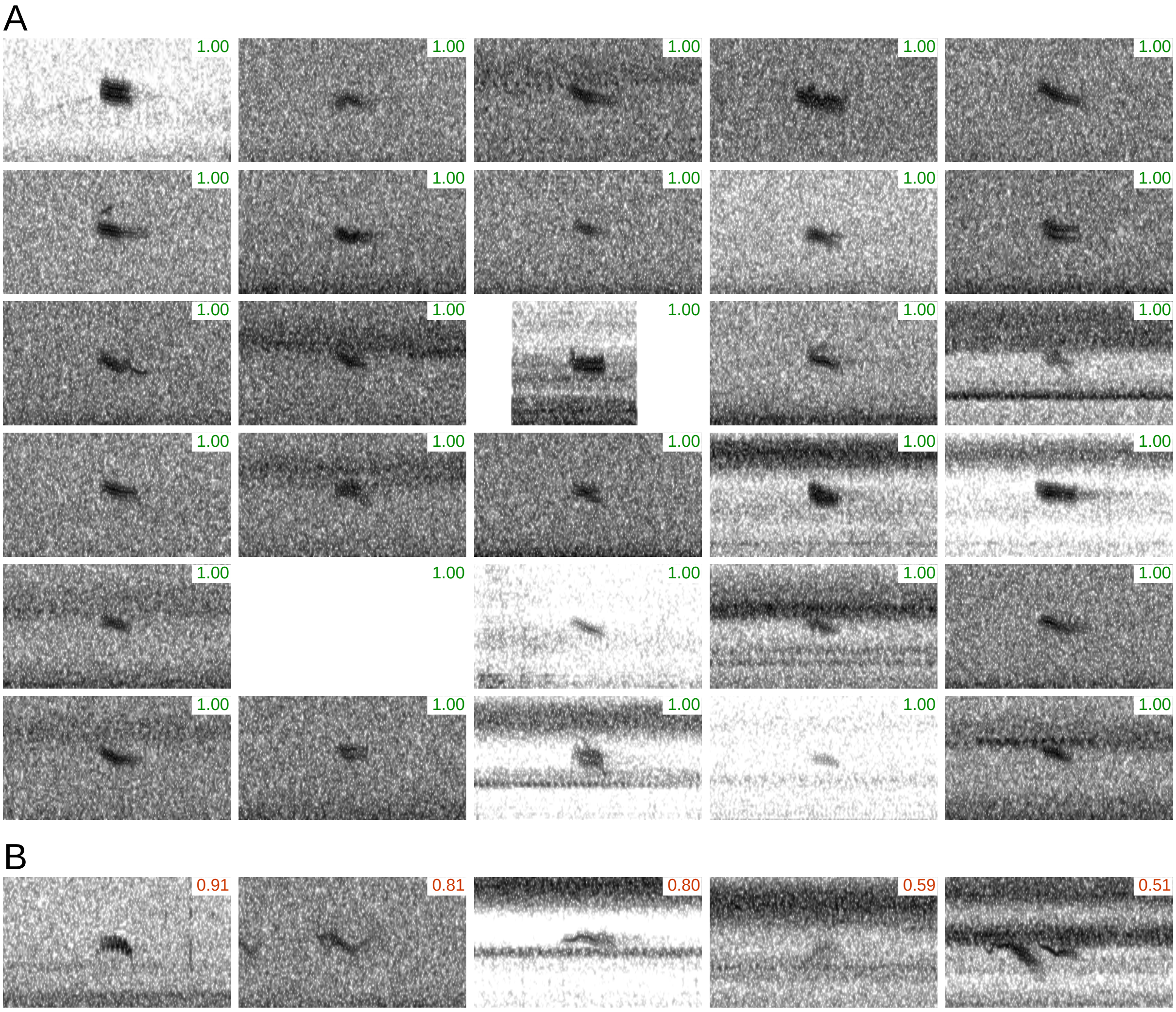


Figure S121: Classification examples for **woothr** drawn from the test dataset, shown as cropped spectrograms. Numbers in upper-right corners are calibrated probability values returned by the model. (A) Test examples scoring highest for this species. Any incorrect classifications are outlined in red. (B) Incorrectly-classified examples scoring highest for this species (i.e., the most confusing cases for the model). Spectrogram parameters: x-axis: 0-0.8 s; y-axis: 1-5 kHz; window type: Hanning; window length: 400 samples; hop size: 30 samples; dynamic range floor: -45 dB).

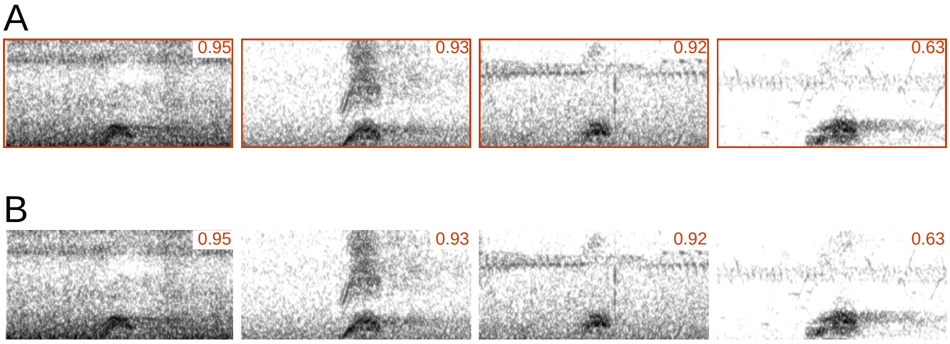


Figure S122: Classification examples for **ycnher** drawn from the test dataset, shown as cropped spectrograms. Numbers in upper-right corners are calibrated probability values returned by the model. (A) Test examples scoring highest for this species. Any incorrect classifications are outlined in red. (B) Incorrectly-classified examples scoring highest for this species (i.e., the most confusing cases for the model). Spectrogram parameters: x-axis: 0-0.8 s; y-axis: 1-5 kHz; window type: Hanning; window length: 400 samples; hop size: 30 samples; dynamic range floor: -45 dB).

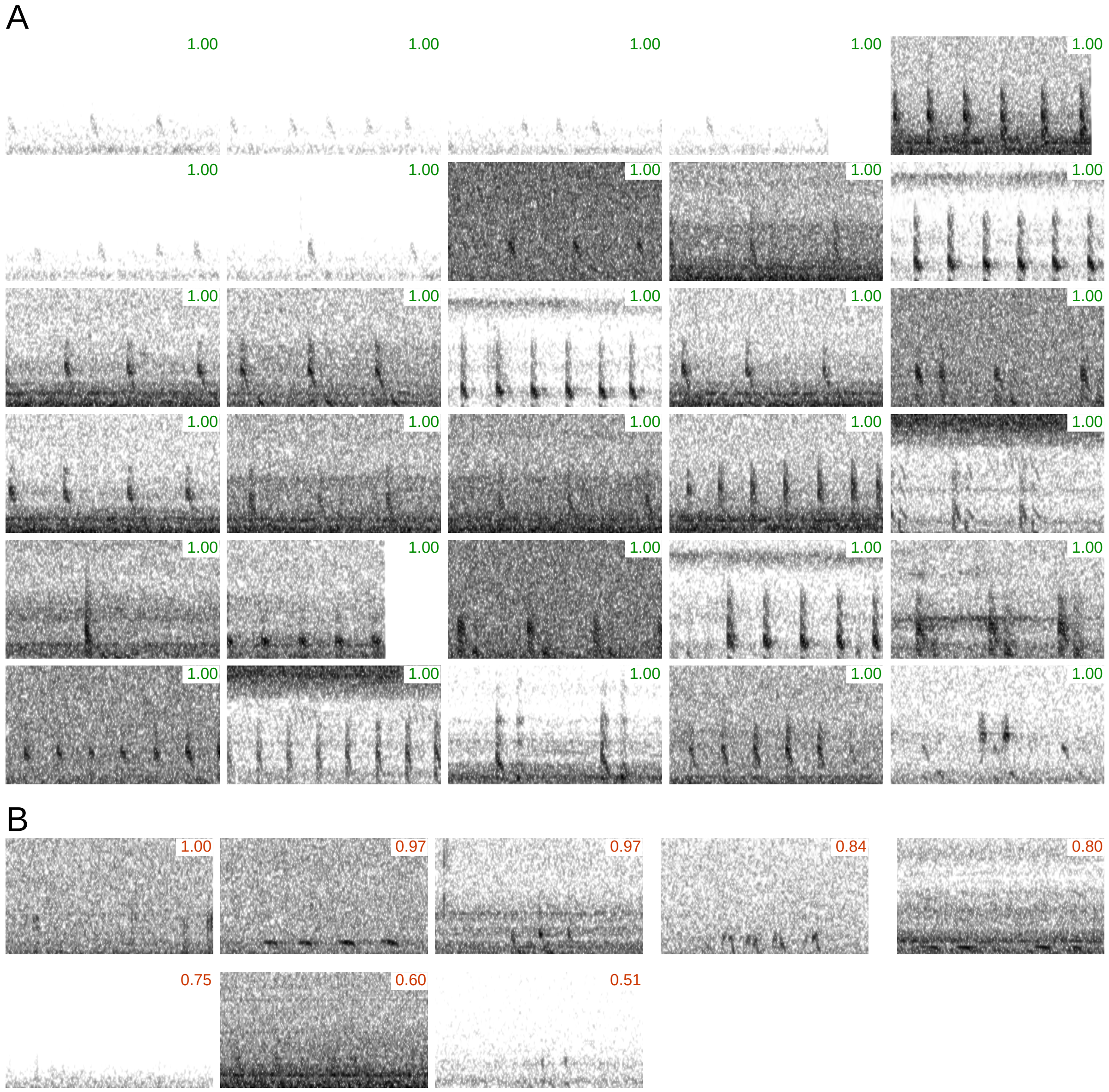


Figure S123: Classification examples for **yebcuc** drawn from the test dataset, shown as cropped spectrograms. Numbers in upper-right corners are calibrated probability values returned by the model. (A) Test examples scoring highest for this species. Any incorrect classifications are outlined in red. (B) Incorrectly-classified examples scoring highest for this species (i.e., the most confusing cases for the model). Spectrogram parameters: x-axis: 0-0.8 s; y-axis: 1-5 kHz; window type: Hanning; window length: 400 samples; hop size: 30 samples; dynamic range floor: -45 dB).

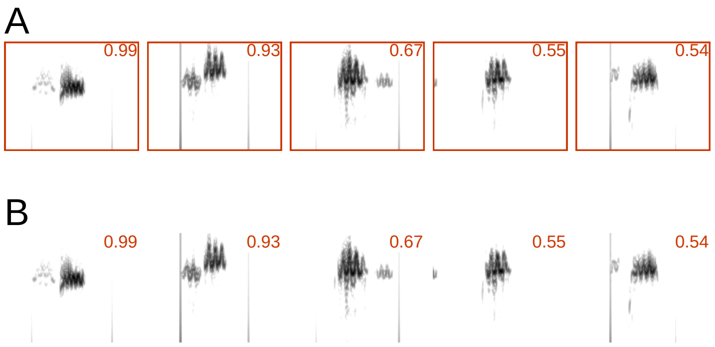


Figure S124: Classification examples for **yelwar** drawn from the test dataset, shown as cropped spectrograms. Numbers in upper-right corners are calibrated probability values returned by the model. (A) Test examples scoring highest for this species. Any incorrect classifications are outlined in red. (B) Incorrectly-classified examples scoring highest for this species (i.e., the most confusing cases for the model). Spectrogram parameters: x-axis: 0-0.3 s; y-axis: 2-10 kHz; window type: Hanning; window length: 200 samples; hop size: 20 samples; dynamic range floor: -60 dB).

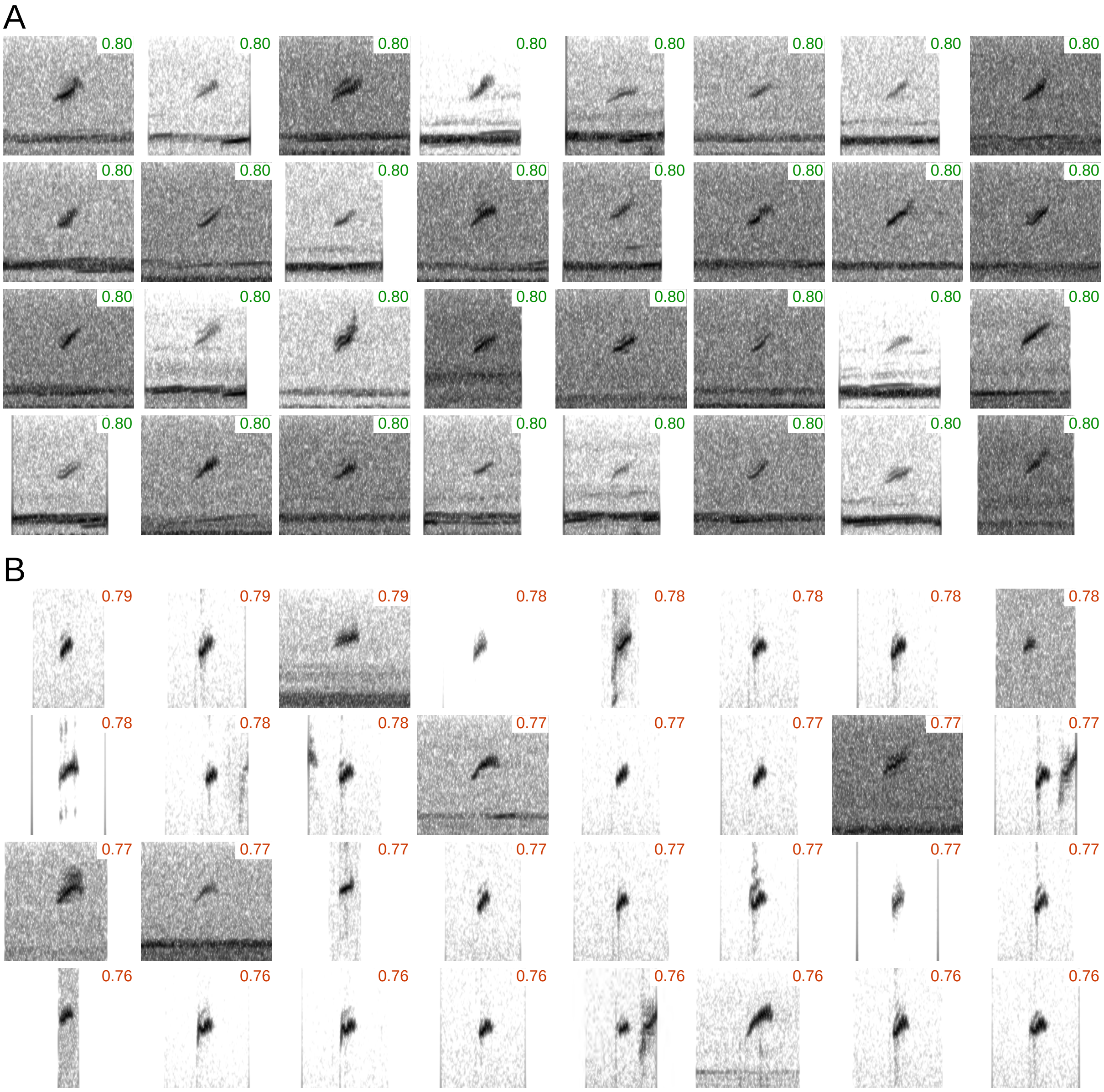


Figure S125: Classification examples for **yerwar** drawn from the test dataset, shown as cropped spectrograms. Numbers in upper-right corners are calibrated probability values returned by the model. (A) Test examples scoring highest for this species. Any incorrect classifications are outlined in red. (B) Incorrectly-classified examples scoring highest for this species (i.e., the most confusing cases for the model). Spectrogram parameters: x-axis: 0-0.3 s; y-axis: 2-10 kHz; window type: Hanning; window length: 200 samples; hop size: 20 samples; dynamic range floor: -60 dB).

### References


Farnsworth, A., Kelling, S., Lostanlen, V., Salamon, J., Cramer, A., Bello, J.P., 2021. Birdvox-296h: a large-scale dataset for detection and classification of flight calls URL: https://zenodo.org/record/4603643, doi:10.5281/zenodo.4603643. dOI: 10.5281/zenodo.4603643 Type: dataset.

Fonseca, E., Favory, X., Pons, J., Font, F., Serra, X., 2020. Fsd50k. Zenodo URL: https://zenodo.org/record/4060432, doi:10.5281/zenodo.4060432. dOI: 10.5281/zenodo.4060432.

Landsborough, B.J., Foote, J.R., Mennill, D.J., 2019. Decoding the 'zeep' complex: quantitative analysis of interspecific variation in the nocturnal flight calls of nine wood warbler species (Parulidae spp.). Bioacoustics 28, 555–574. URL: https://doi.org/10.1080/09524622.2018.1509373, doi:10.1080/09524622.2018.1509373.

Lanzone, M., Deleon, E., Grove, L., Farnsworth, A., 2009. Revealing undocumented or poorly known flight calls of warblers (Parulidae) using a novel method of recording birds in captivity. The Auk 126, 511–519. URL: https://doi.org/10.1525/auk.2009.08187, doi:10.1525/auk.2009.08187.

Lostanlen, V., Salamon, J., Farnsworth, A., Kelling, S., Bello, J.P., 2017. Birdvox-full-night: a dataset for avian flight call detection in continuous recordings. Zenodo URL: https://zenodo.org/record/1205569, doi:10.5281/zenodo.1205569. dOI: 10.5281/zenodo.1205569.

Lostanlen, V., Salamon, J., Farnsworth, A., Kelling, S., Bello, J.P., 2018. 2018 ieee international conference on acoustics, speech and signal processing (icassp), pp. 266–270. doi:10.1109/ICASSP.2018.8461410. iSSN: 2379-190X.

Morris, S.R., Horton, K.G., Tegeler, A.K., Lanzone, M., 2016. Individual flight-calling behaviour in wood warblers. Animal Behaviour 114, 241–247. URL: https://linkinghub.elsevier.com/retrieve/pii/S0003347216000385, doi:10.1016/j.anbehav.2016.01.027.

Piczak, K.J., 2015. Esc: Dataset for environmental sound classification, ACM Press. p. 1015–1018. URL: http://dl.acm.org/citation.cfm?doid=2733373.2806390, doi:10.1145/2733373.2806390. place: Brisbane, Australia Citation Key: piczak2015dataset.

Salamon, J., Jacoby, C., Bello, J.P., 2014. A dataset and taxonomy for urban sound research, Association for Computing Machinery, New York, NY, USA. p. 1041–1044. URL: https://doi.org/10.1145/2647868.2655045, doi:10.1145/2647868.2655045.

Van Doren, B.M., Sheldon, D., Geevarghese, J., Hochachka, W.M., Farnsworth, A., 2015. Autumn morning flights of migrant songbirds in the northeastern United States are linked to nocturnal migration and winds aloft. The Auk 132, 105–118. URL: http://www.bioone.org/doi/abs/10.1642/AUK-13-260.1, doi:10.1642/AUK-13-260.1.
